# Spatial compartmentalization of signalling imparts source-specific functions on secreted factors

**DOI:** 10.1101/2022.08.20.504649

**Authors:** E Groppa, P Martini, N Derakhshan, M Theret, M Ritso, LW Tung, YX Wang, H Soliman, M Hamer, L Stankiewicz, C Eisner, E Le Nevé, C Chang, L Yi, JH Yuan, S Kong, C Weng, J Adams, L Chang, A Peng, HM Blau, C Romualdi, FMV Rossi

**Affiliations:** School of Biomedical Engineering, University of British Columbia, 2222 Health Sciences Mall, Vancouver, British Columbia, Canada; Borea Therapeutics, Scuola Internazionale Superiore di Studi Avanzati, Via Bonomea 265, Trieste, Italy; Department of Molecular and Translational Medicine, University of Brescia, Brescia, Italy; Department of Biology, University of Padova, via U. Bassi 58B, Padova; Baxter Laboratory for Stem Cell Biology, Department of Microbiology and Immunology, Stanford University School of Medicine, Stanford, USA; Faculty of Pharmaceutical Sciences, Minia University, Minia, Egypt; Aspect Biosystems, 1781 W 75th Ave, Vancouver, BC, Canada; Department of Pediatrics, Université Laval, Quebec, Canada

**Keywords:** bioinformatics, intercellular signaling redundancy, muscle progenitor, FAP, VEGFA, skeletal muscle, angiogenesis

## Abstract

Efficient regeneration requires multiple cell types acting in a coordination. To better understand the intercellular networks involved and how they change when regeneration fails, we profiled the transcriptome of hematopoietic, stromal, myogenic, and endothelial cells over 14 days following acute muscle damage. A time-resolved computational model of interactions was generated, and VEGFA-driven endothelial engagement was identified as a key differentiating feature in models of successful and failed regeneration. In addition, it revealed that the majority of secreted signals, including VEGFA, are simultaneously produced by multiple cell types. To test whether the cellular source of a factor determines its function, we deleted VEGFA from two cell types residing in close proximity, stromal and myogenic progenitors. By comparing responses to different types of damage, we found that myogenic and stromal VEGFA have distinct functions in regeneration. This suggests that spatial compartmentalization of signaling plays a key role in intercellular communication networks.

**Highlights:**

- Ligand-receptor signaling redundancy during skeletal muscle regeneration
- Inflammatory cells, and muscle and fibro/adipogenic progenitors produce VEGFA
- VEGFA from muscle progenitors control their proliferation after muscle damage
- VEGFA from FAP controls angiogenesis only after ischemic damage

**eTOC blurb:** Groppa *et al*. performed a novel time-resolved bioinformatics analysis that revealed extensive ligand-receptor redundancy among the cell types contributing to skeletal muscle regeneration. They focused on one of these pathways, and showed that VEGFA from different cell types has distinct roles in regeneration.

## INTRODUCTION

Regeneration and development are among the many biological processes that require highly dynamic changes in cellular functions to take place in a coordinated fashion across multiple cell types. Such multicellular synchronization occurs via intercellular interactions mediated by a variety of signals, including a diverse number of secreted factors and their cognate receptors. Disruption of these cellular networks is likely implicated in pathogenic responses to damage such as fibrosis. Understanding the rules that govern intercellular communication and formulating predictive models capable of highlighting specific interactions have the potential to identify therapeutic intervention points.

To date, work towards the creation of such models has been mainly carried out in developmental systems. In RNAseq datasets, manual curation has been applied to define lists of relevant factors and receptors expressed by specific cell types. From these lists, interactions have been predicted computationally, generating models that represent ongoing interactions (Choi et al., 2015; Efremova et al., 2020; Kumar et al., 2018; Rezza et al., 2016; Verma et al., 2018; Zhou et al., 2017). In all cases, these models were created using data gathered at a single time point, and fail to capture the temporal dynamics of “cellular interactomes” as they exist *in vivo* over the course of a physiological response.

Skeletal Muscle is a highly regenerative tissue capable of complete functional restoration following damage (Mukund and Subramaniam, 2020). However, pathologies such as chronic damage (e.g. muscular dystrophies) and aging result in a loss of this regenerative capacity, providing a clinically compelling reason to identify strategies for restoring it (Babbs et al., 2020; Carosio et al., 2011). Here, we generated a dynamic model of skeletal muscle regeneration based on recurrent RNAseq sampling of multiple purified cell populations involved in the regenerative response to acute damage. We focused on communication across 1) muscle stem cells, also called satellite cells, 2) tissue resident mesenchymal fibro/adipogenic progenitors, 3) immune cells, 4) pericytes, and 5) endothelial cells. Previous studies have revealed a complex web of regulatory interactions among these cell types (Arnold et al., 2007; Chiristov et al., 2007; Dammone et al., 2018; Fiore et al., 2016; Juban et al., 2018; Joe et al., 2010; Kostallari et al., 2015; Latroche et al., 2017; Lemos et al., 2015; Mojumdar et al., 2014; Ochoa et al., 2007; Saclier et al., 2013; Uezumi et al., 2010; Villalta et al., 2014; Willenborg et al., 2012; Wosczyna et al., 2019; Zhang et al., 2020). This intercellular communication landscape is highly dynamic, with the same cell lineages carrying out drastically different and sometimes opposing functions at different times after damage (Scott et al., 2019; Sobral-Reyes and Lemos, 2019).

We built a new bioinformatic pipeline and used it to assemble a time-resolved interactome that connects the different skeletal muscle cell populations through either autocrine or paracrine ligand-receptor interactions. To identify the cellular interactions responsible in regeneration, we employed this pipeline to compare the responses in wild type mice to those of mice lacking C-C chemokine receptor 2 (CCR2KO) (Ochoa et al., 2007; Willenborg et al., 2012), a mouse strain in which circulating blood derived monocytes are unable to infiltrate the parenchyma of damaged tissue, thereby impairing muscle regeneration. We found that a key difference between the two models was the extent of engagement of endothelial cells, and we identified the lack of VEGFA from infiltrating macrophages as a causative factor.

Surprisingly, our analysis shows that VEGFA, as well as the vast majority of ligands and their cognate receptors, are expressed by multiple cell lineages within regenerating muscle. The pleiotropic activity of VEGFA signaling on a variety of cell types that are in close proximity, like endothelial and myogenic cells (Chiristov et al., 2007), prompted us to use this signaling as a benchmark to question whether ligands from different lineages function equally, and exist purely to provide redundancy and therefore robustness, or whether the cell of origin of a given secreted factor imparts specificity on its role during the regenerative process. To experimentally address this question, we deleted the VEGFA gene individually in each of its sources to ask if they are functionally equivalent.

Our results support the notion that signaling is spatially compartmentalized *in vivo*, and that the cellular origin of a ligand strongly impacts its function, with distinct cell lineages playing critical roles in response to specific types of damage. We conclude that the inclusion of spatial information and anatomical features in cellular interaction models will be critical to understand the ground rules underlying intercellular signaling *in vivo*.

## RESULTS

### A time- and lineage-resolved transcriptional analysis of regeneration

In order to describe the intercellular communication network that supports skeletal muscle regeneration, we purified fibro/adipogenic progenitors (FAP), endothelial cells (EC), myogenic progenitors (MP), inflammatory cells (IC), and pericytes (P) at steady state and at multiple time points after muscle damage. We induced damage by intramuscular injection of Notexin (NTX), a myotoxin, in the *Tibialis Anterior* muscle (TA) in both wild type (WT) and C-C chemokine receptor 2 knock out mice (CCR2KO), which provided models for efficient and delayed regeneration, respectively. In addition to purified populations from each genotype, whole TA bulk tissue samples from WT mice were also collected and analyzed at each time point. We subjected all samples to RNA sequencing and assembled a new multistep bioinformatic analysis workflow, part of which is schematically represented in Figure S1A. We first identified differentially expressed genes (DEG) in each cell subset (cell type specific analysis), followed by genes whose change in overall abundance is associated with the expansion of specific cell subsets rather than with changes in their transcriptional output per cell, and finally genes that are constitutively expressed by specific cell subsets. Next, based on the selected genes, we inferred crosstalk among specific cell subsets by pairing expression of ligands (Lig) and their cognate receptors (Rec); finally, we determined the biological processes associated with DEG and Lig-Rec pairs in each cell type.

Principal component analysis (PCA) on FAP, EC, MP, IC, and PER transcriptomes pooled from all the collected time points displayed clear separation among the cell subsets, confirming their distinct nature and a lack of ambiguity in their definition (Figure S1B-C). Next, we assessed transcriptome dynamics in each cell type during muscle regeneration by comparing gene expression across the different collection time points. We employed two types of bioinformatics analyses, silhouette and biological homogeneity index, to generate clusters grouping the different time points in a time-unbiased manner and estimate similarity across clusters. Samples clustering together across multiple timepoints in the PCA plots (Figure 1B and C, clusters are color-coded) indicate that the transcriptome, and presumably the function, of that specific cell type did not significantly change over time. Samples in which each cluster contains all samples from a subset of timepoints (Figure 1A and D) suggest that over time, cells moved through a discrete set of transcriptional states, likely indicating different functions. When performed on whole tissue samples, this analysis clearly showed that the regenerative process relies on well defined, temporally distinct stages (Figure S1D). Next, we investigated which of the tissue resident lineages are most dynamic, and therefore most likely to drive transitions across the different phases of regeneration. We initially focused on FAPs, which showed highly dynamic behavior and ECs, which changed very little over the time course (Figure 1A and C). FAP transcriptomes were grouped in four clusters based on silhouette analysis, suggesting they move through four main states during the repair process, with each of the clusters containing all replicates of the same time points, as shown by the high level of homogeneity index (0.6). Specifically, we found that FAP transitioned through two distinct stages early after damage (day 1 and day 2 time points formed two discrete clusters), followed by a third state (spanning days 3, 4, 5, and 6) and finally entered into a fourth steady-state shared by both the undamaged and late time points (day 0, 7, 10, and 14), as recently described by others (De Micheli et al., 2020). This ordered progression was disrupted in CCR2KO mice, in which clusters displayed a much lower homogeneity index (0.35), suggesting a key role for infiltrating inflammatory cells in the temporal organization of FAP transition states and of the overall regenerative process (Figure 1A-B). This is consistent with our previous description of the behavior of FAP in WT vs CCR2KO animals (Lemos et al., 2015).

**Figure 1:**
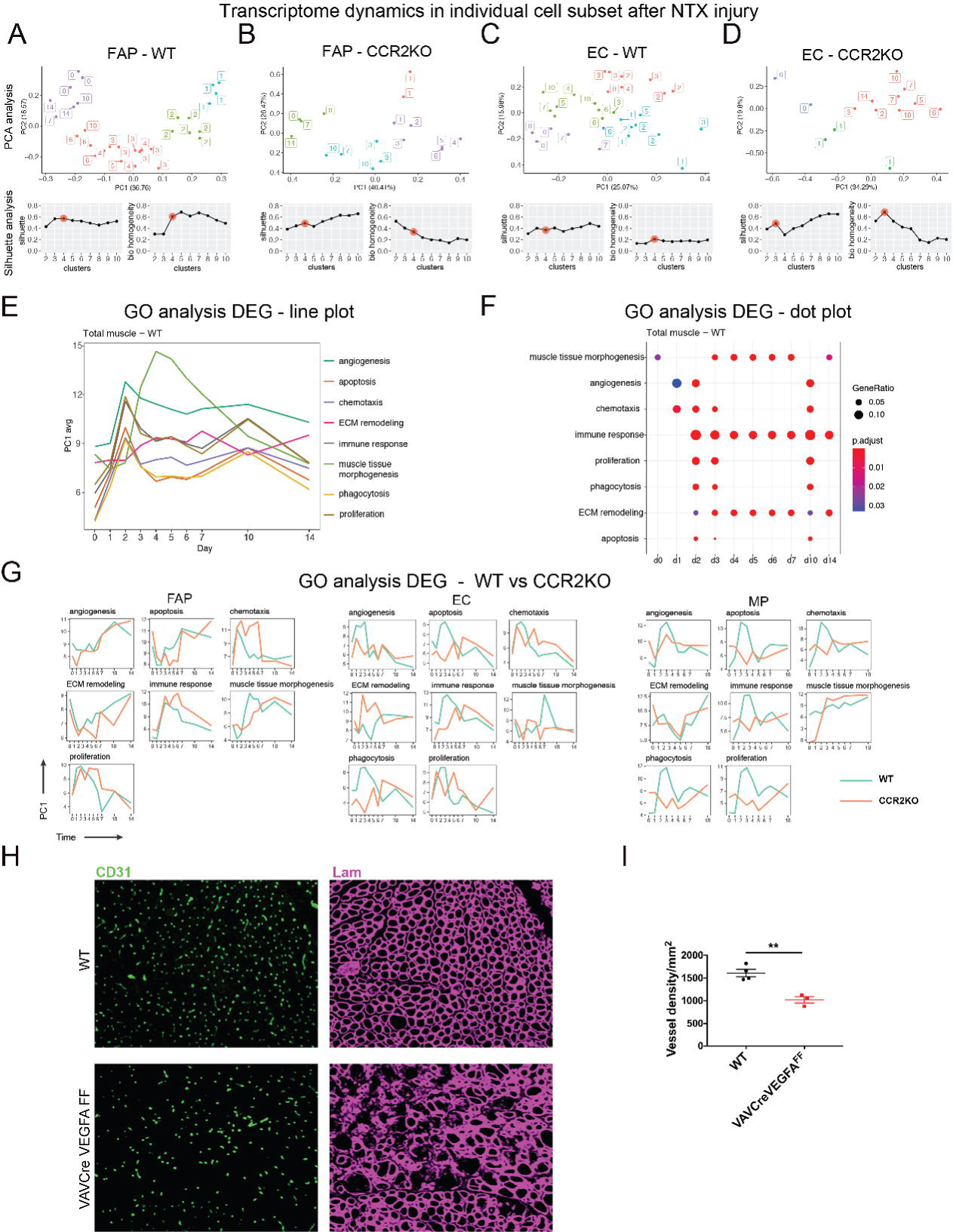
Distinct cellular activation in normal and delayed regeneration. **A-D)** Silhouette, biological homogeneity index and PCA analysis for FAP and EC RNAseq data collected throughout the response to myotoxin damage from both WT and CCR2KO mice. In the PCA plots, the number inside each box represents the time of harvest in days post damage, with zero representing undamaged samples. The color of the box indicates which samples cluster together. The representative values described in the main text are highlighted with red filled circles. **E)** Representation of the 8 main Gene Ontology (GO) categories over time, in whole WT muscle. **F)** GO enrichment analysis for the 8 main categories; dot color reflects adjusted p-value and dot size the ratio between category-associated DEGs and total number of genes in the category. **G)** Comparison of expression over time in each indicated cell type (FAP, EC, and MP) of the genes associated with the 8 GO category in WT and CCR2KO. **H)** Representative image of muscles collected from WT and VAVCre VEGFA^FF^ at 7 days after NTX damage. **I)** Quantification of vessel density based on CD31 staining of the samples shown in I (n=3-4, data represent the mean ± SEM, unpaired t test, **p< 0.01).

### Lack of inflammatory cell-derived VEGFA is responsible for disrupted regeneration in CCR2KO mice

In contrast, EC clusters were characterized by poor homogeneity index (0.21 with silhouette n = 4) in WT animals, which increased (0.7 with silhouette n = 3) in CCR2KO (Figure 1C and D). This suggests that NTX damage has only minor effects on the endothelium in WT mice, but that the lack of infiltrating inflammatory cells in the CCR2KO exacerbates the damage leading to increased involvement of the endothelium during the regenerative process. Histological analysis confirmed that only limited vascular remodeling, observed as a small increase in vascular density immediately following NTX damage, was taking place in WT animals (Figure S1E and G). In CCR2KO mice however, vessel disruption was much more evident and the response appeared biphasic, with the vessels’ diameter abnormally increasing early after damage, normalizing, and then increasing again 7 days post damage (Figure S1F and G). Increased vascular permeability, which was only observed on day 3 in controls, persisted until day 10 in CCR2KO (Figure S1H), consistently with the poor coverage by pericytes that regulate vascular integrity (Figure S1I).

To understand the biological activities associated with each cell lineage during normal and impaired muscle regeneration, we performed Gene Ontology (GO) Biological Process (BP) enrichment analysis based on DEG from whole tissue and individual cell populations. With the goal of both broadening as well as simplifying the outcomes of our analysis, we grouped GO BP terms in 8 main categories: muscle tissue morphogenesis, extracellular matrix (ECM) remodeling, immune response, apoptosis, chemotaxis, phagocytosis, proliferation, and angiogenesis. The results of our GO analysis were displayed as line plots, based on the top genes associated with PCA1, and dot plots, determined by the number of genes expressed by a cell type out of the total genes attributed to a given GO metaclass (defined in Material and Methods and Table S1). In the whole tissue, apoptosis, chemotaxis, phagocytosis, proliferation, and angiogenesis were predominant at early stages of regeneration (day1 to day 3 post-injury), while immune response, muscle tissue morphogenesis and ECM remodeling were first observed at 2 to 3 days after damage and remained active throughout the later stages of the repair process, with muscle tissue morphogenesis gradually returning towards baseline by the end point (Figure 1E-F).

GO enrichment analysis on DEGs showed that all cell types participate in the biological processes observed in whole muscle, differing from each other mainly in the timing and extent of their contribution (Figure S2A-B). Further analysis revealed that many biological processes, like apoptosis and muscle tissue morphogenesis, were delayed in CCR2KO FAPs compared to WT FAPs (Figure 1G). In contrast, proliferation and chemotaxis were prolonged, in line with the phenotype previously described in CCR2KO FAPs (Lemos et al., 2015). Similarly, we found that angiogenesis, chemotaxis, and proliferation were delayed in ECs from CCR2KO compared to ECs from WT, consistent with the abnormal vascular remodeling observed in CCR2KO vs WT mice after NTX damage (Figure 1G and S1E-I). Overall, WT MPs were more active than CCR2KO MPs, consistent with the impairment of the regenerative process in CCR2KO.

We tested the validity of the conclusions emerging from our GO enrichment analysis with additional bioinformatics and experimental investigations focused on cell proliferation. We determined the mitotic index (MI) from FAPs, ECs, and MPs in both WT and CCR2KO (Dmitrijeva et al., 2018). In parallel, we measured EdU incorporation in FAPs, ECs, and MPs at different time points after NTX injection. A notable concordance was observed between proliferation kinetics extracted from GO data and both MI as well as EdU experimental data (Figure S2C). While FAPs, ECs, and MPs in both WT and CCR2KO mice proliferate from day 1 to day 3 post-injury (early phase), they slowed drastically in WT mice after that point, but continue to expand in CCR2KO animals.

A key molecule produced by infiltrating macrophages is VEGFA, a well-known pro-angiogenic factor whose absence from CCR2KO mice might impair the endothelial response to damage. To test whether this was the case, we deleted VEGFA from all hematopoietic lineages by generating VAVCre VEGFA^FF^ mice. In these animals we found a reduction of vessel density consistent with the impaired vascular remodeling previously observed in our CCR2KO model, where the recruitment of VEGFA-producing monocytes from the bloodstream is absent (Figure 1H-I). This also resulted in disrupted muscle regeneration to a degree similar to that observed in CCR2KO, confirming that hematopoietic VEGFA was indeed a critical missing signal when inflammatory infiltration was prevented. In accordance with our previous work, we did not observe the typical fibrofatty infiltration triggered by damage in the absence of inflammatory cell recruitment, which is caused by lack of TNFα (Lemos et al., 2015) (Figure S2D).

In conclusion, our time-resolved transcriptomic analysis is capable of measuring the extent to which specific cell types are recruited by and contribute to regeneration. In addition, it pinpoints disruptions of the cells’ normal progress through distinct phases of the regenerative process. Together, this allowed the deconvolution of the complex phenotype of CCR2KO mice, and the identification of a functionally relevant biological process perturbed in this model.

### Intercellular communication networks are promiscuous

Most developmental processes, including adult tissue regeneration, are supported by a rich network of intercellular communications. With the goal to begin describing such networks in the context of muscle regeneration, we built a time-resolved map of both autocrine and paracrine signaling based on pairing manually curated lists of expressed receptors and ligands, and analyzed it through GO pathway enrichment. For each cell type and GO category analyzed, we subdivided the type of interaction as autocrine when the same cell type expressed both cognate ligand and receptor, paracrine when a cell type expressed one but not the other, and “shared” when a cell type engaged in autocrine or paracrine signaling but the ligand or receptor was also expressed by other cell types. We found each category to be represented in each GO and cell type, with shared interactions representing a surprisingly large proportion of the cellular interactome (Figure 2A and S3A). This suggests that each cell type interacts with multiple other cell populations through the same ligand-receptor interactions. In line with this finding, we observed that nearly 75% of all receptors and ligands are expressed at the same time by two or more cell types (Figure 2B).

**Figure 2:**
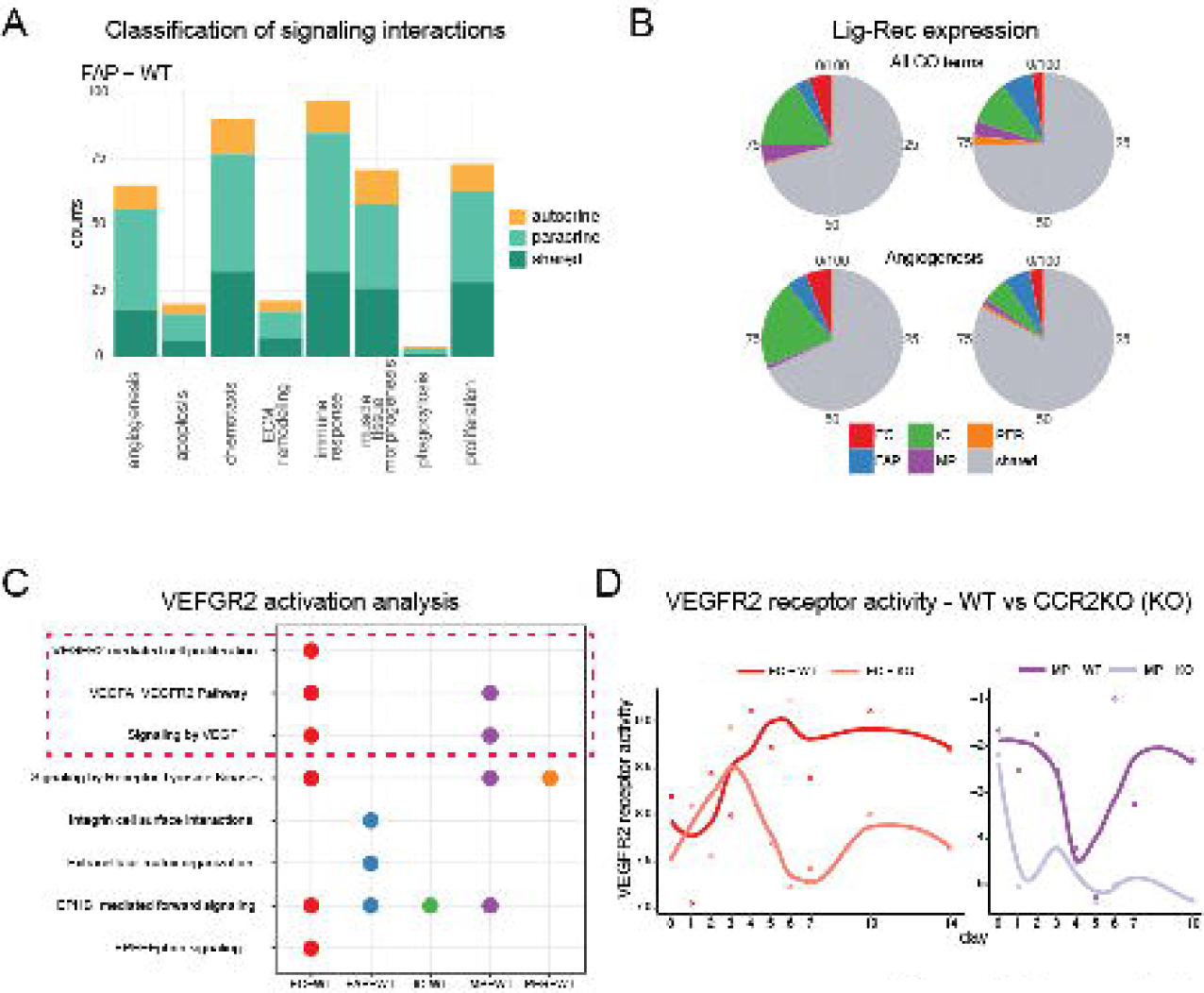
VEGFA, an example of promiscuous intercellular communication. **A)** Classification of signalling interactions of FAPs by GO category. Interactions are classified as autocrine if FAPs express both cognate ligand and receptor (in yellow), paracrine when FAP expresses one but not the other (in light green), and “shared” when FAP engages in autocrine or paracrine signaling but the ligand or receptor is also expressed by other cells (in dark green). **B)** Pie charts show the percentage of ligands and receptors expressed by only one cell type (colours) or shared by more than two cell types (grey). **C)** VEGFR2-related pathway activation in different cell types. **D)** Activation profile (Viper analysis) of *Kdr* (VEGFR2) in EC and MP from WT and CCRKO at different time points after NTX damage.

In summary, we found that during regeneration the vast majority of ligands have multiple cellular sources and cellular targets. This could represent true redundancy, in which ligands produced by different cells carry out the same function in the context of regenerating tissue, and multiple sources serve to confer robustness to the system. Alternatively, the same ligands could modulate specific tasks depending on their cell of origin.

### VEGFA from MP, but not from FAP, is important in skeletal muscle regeneration after acute damage

To assess whether ligands originating from different cell types are functionally equivalent, we again focused on VEGFA signaling, as it is vital for successful regeneration. Interestingly, beyond modulating angiogenesis, VEGFA has been proposed to have additional functions in healthy and diseased muscle (Verma et al., 2018, 2021). In both undamaged and damaged muscles, the ligand *Vegfa* can be produced by FAPs, MPs, and ICs (Figure S3B and Table S2). While its main receptor *Kdr* (VEGFR2) is restricted to MPs and ECs, co-receptors like *Nrp1*, *Nrp2*, and *EphB2* are also expressed by more than one cell type.

We first verified that the pathway is indeed functional during regeneration by applying two computational approaches: 1) regulatory network analysis, which determines the activity of a protein based on the number of co-expressed genes that belong to the same network unit; and 2) functional validation of receptor activity (TimeClip (Martini et al., 2014)), which elucidates the activation of the receptor downstream signaling over time. Based on the first analysis, we confirmed the activation of gold standard receptors for the different cell populations, such as PDGFRa for FAPs, PDGFRb for PERs, CCR2 for ICs, Tek for ECs, and Notch3 for MPs (Figure S4A). As expected, VEGFR2 was strongly active in ECs, but to a lesser extent also activated in ICs, MPs, and PERs. In the second approach, we integrated these results with the analysis of signaling downstream of VEGFR2 over time, which showed that VEGFA/VEGFR2 signaling is specifically active in ECs and MPs, but not in other cell subsets, consistent with the outputs of the intercellular network (Figure 2C). Interestingly, a parallel analysis showed reduced and delayed VEGFR2 activity in CCR2KO vs WT, in both EC and MPs (Figure 2D).

The effects of VEGFA depletion from inflammatory cells have already been reported in regenerative processes such as skin wound healing (Willenborg et al., 2012), but there is less known about the role of VEGFA produced by MPs and FAPs during muscle regeneration. First, we stained for VEGFA protein on cells sorted from muscle immediately following acute damage. As predicted in our bioinformatic analysis, and consistent with recently published results (Verma et al., 2018), we confirmed that it is produced by multiple cell types, including significant contribution from FAPs and MPs (Figure 3A). We next assessed the specific contribution of MPs and FAPs to overall VEGFA levels in the tissue by depleting it in different cell subsets. To this end we generated two mouse strains in which both alleles of VEGFA are floxed. The first strain carries the FAP-specific PDGFRa CT2+ (FAP^VEGFAKO^), while the second contains the MP-specific Pax7 CT2+ (MP^VEGFAKO^). After inducing Cre activation by tamoxifen treatment, we damaged the muscles with NTX and collected the tissues at 3, 7, and 10 days after injury. The efficiency of VEGFA deletion was similar (∼85%) in both Cre systems. Elisa for VEGFA on whole tissue revealed that its depletion in either MPs or FAPs reduced the total amount of this factor by a comparable amount, providing quantitative support that their contribution to VEGFA secretion is similar during muscle regeneration after NTX damage (Figure 3B). Inflammatory cell infiltration, comparable both quantitively and in terms of cell phenotypes across the three systems, was observed at 3 days after damage (Figure S4B). At 7 days post-injury, we observed no difference in gross muscle organization in FAP^VEGFAKO^ compared to control CT2-VEGFA floxed (WT), but found that the newly forming fibers were smaller and displayed disorganized morphology in tissues derived from MP^VEGFAKO^ mice (Figure 3C-E). Interestingly, there was no difference in vasculature density among the three groups (Figure 3F). Ten days after NTX, WT and FAP^VEGFAKO^ showed normal muscle regeneration represented by fibers with homogeneous diameter size, while MP^VEGFAKO^ was still characterized by smaller muscle fibers (Figure S4B).

**Figure 3:**
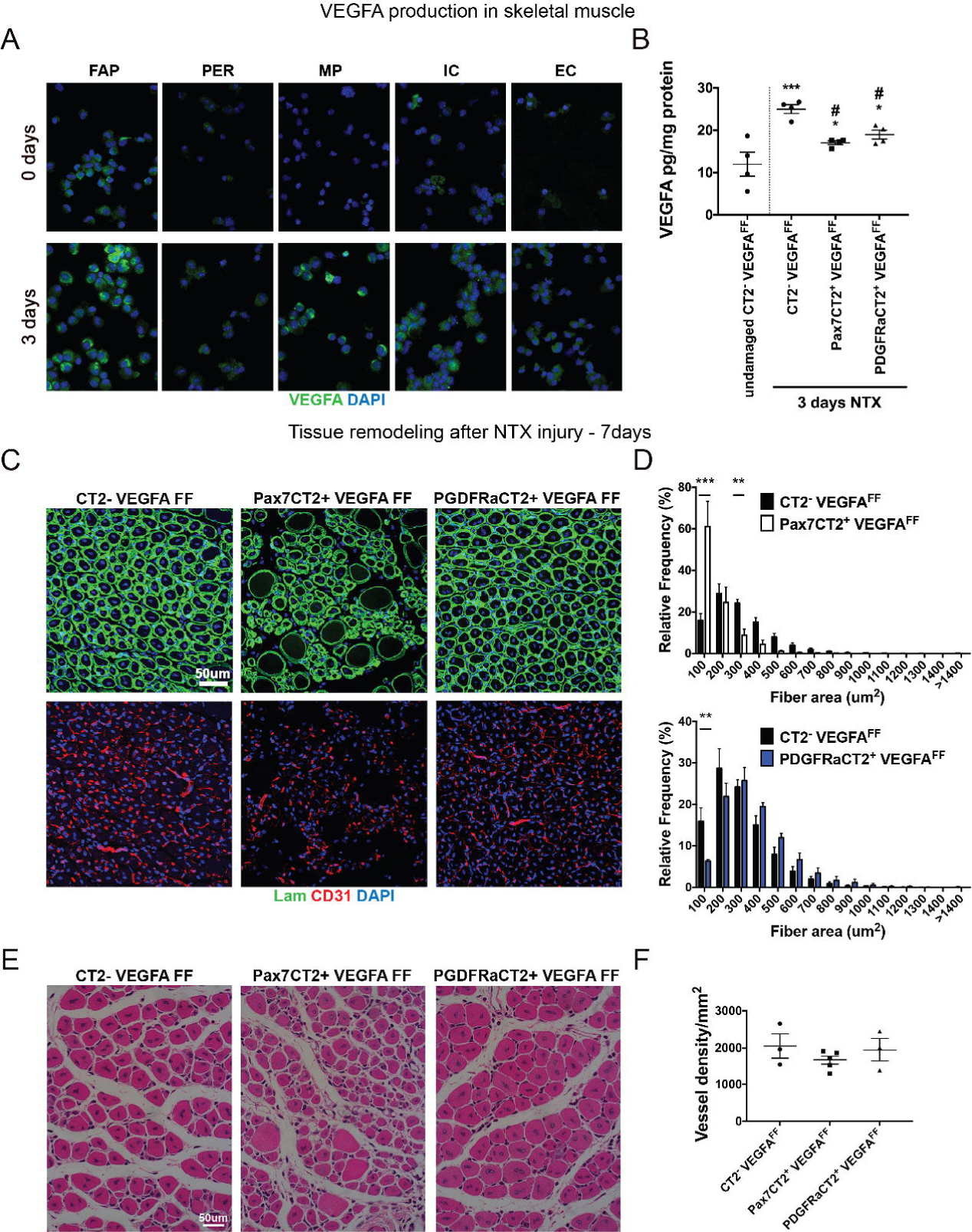
VEGFA from MP, but not from FAP, is important in skeletal muscle regeneration after acute damage. **A)** VEGFA staining on cytospinned FAP, EC, MP, IC, and PER purified from skeletal muscle before and after acute damage. **B)** Whole skeletal muscle VEGFA-Elisa (n=4, data represent the mean ± SEM, one-way ANOVA with multiple comparison, *p< 0.05, **p< 0.01, ***p< 0.001). **C)** Skeletal muscles collected from WT, MP^VEGFAKO^, and FAP^VEGFAKO^ 7 days after NTX damage and stained with laminin and CD31. **D)** Quantification of fiber size of the samples shown in C; n=3-4, data represent the mean ± SEM, two-way ANOVA with multiple comparison, **p< 0.01, ***p< 0.001. **E)** H&E of skeletal muscles collected from WT, MP^VEGFAKO^, and FAP^VEGFAKO^ at 7 days after NTX. **F)** Quantification of vessel density of the samples shown in C; n=3-5, data represent the mean ± SEM, one-way ANOVA with multiple comparison.

These data suggest that while VEGFA from MPs is critical for efficient skeletal muscle repair following acute myotoxin damage, VEGFA from FAPs is not only dispensable, but fails to rescue the phenotype resulting from ablation of VEGFA in MPs. Thus, autocrine VEGFA produced by MPs acts through a signaling network that paracrine VEGFA from FAPs is unable to activate, despite the close proximity of the two cell types in the tissue. Additionally, neither MP- nor FAP-derived VEGFA have an effect on regenerative angiogenesis in this setting.

### VEGFA controls MP proliferation, but not differentiation

To investigate the mechanism underlying the observed phenotype, we measured MP proliferation 3 and 7 days post-injury in mice that received a single EdU dose the day before harvest. At 3 days post-injury, MPs from MP^VEGFAKO^ mice were less proliferative and less abundant than in control animals, but overall TA muscle mass was comparable (Figure 4A). At 7 days post-injury, the observation had completely reversed; a higher percentage of MPs were proliferating in the MP^VEGFAKO^ mice than in control mice, and this compensatory expansion led to a normalization of the number of MPs (Figure 4B). However, TA muscle mass was reduced in the MP^VEGFAKO^ animals. Consistent with the lack of vessel density impairment in MP^VEGFAKO^ mice, we did not see any difference in EC proliferation (Figure S5A). Similar assays performed with MPs isolated from FAP^VEGFAKO^ mice showed that VEGFA depletion in FAPs does not reduce either MP or EC proliferation, and is consistent with the lack of phenotype observed in muscles of FAP^VEGFAKO^ mice.

**Figure 4:**
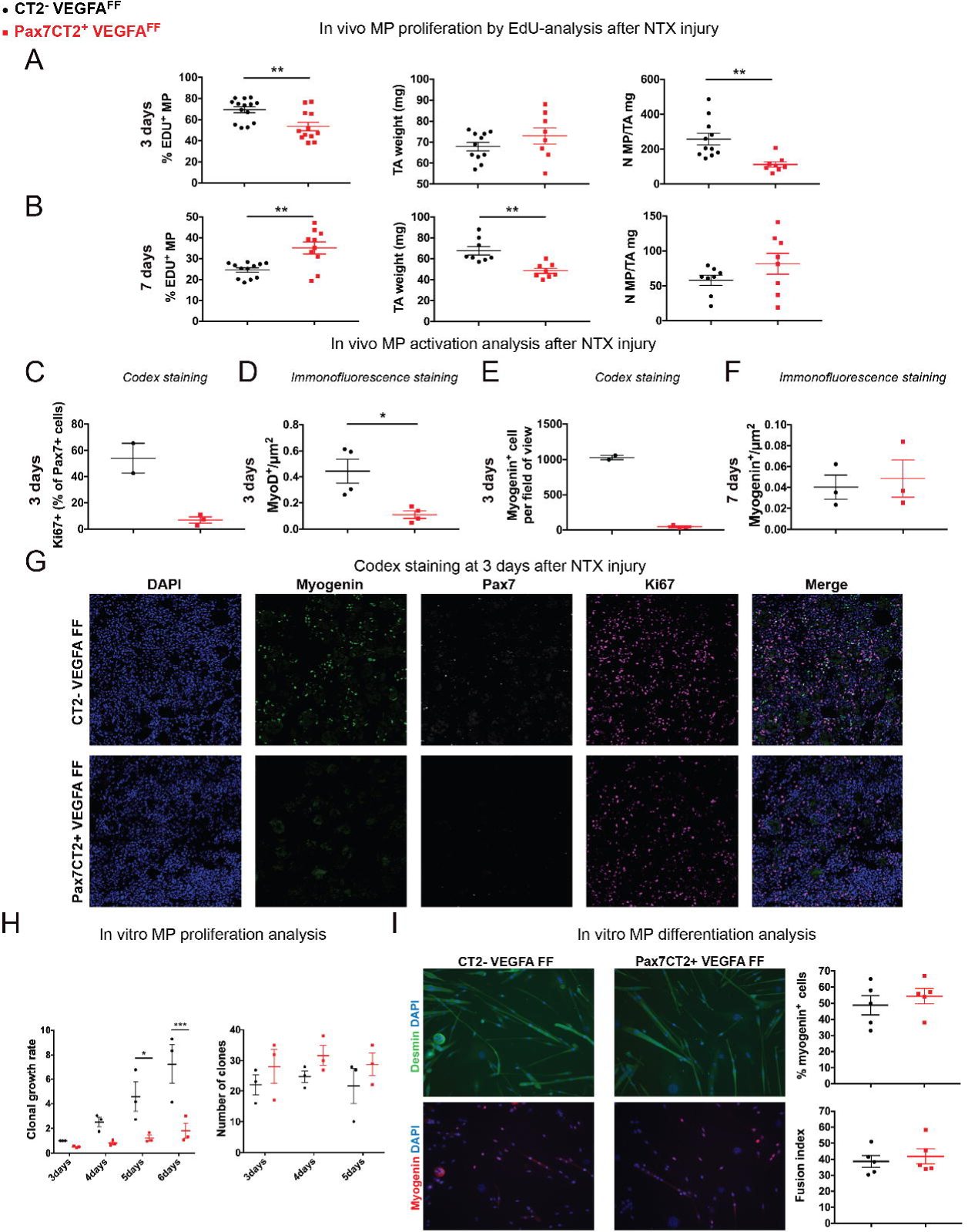
Autocrine VEGFA controls early MP-proliferation. **A-B)** Left: EdU incorporation in MPs from WT and MP^VEGFAKO^ mice, 3 and 7 days after NTX injury and 24 hrs after EdU treatment. Centre: TA muscle weight, and Right: absolute number of MP/mg of muscle in the same samples (n=8-14, data represent the mean ± SEM, unpaired t test or Mann Whitney test, **p< 0.01). **C-E)** Quantification of Ki67/Pax7, MyoD, and Myogenin positive cells from skeletal muscle collected at 3 days after NTX from wild type and MP^VEGFAKO^ animals; n=2-4, data represent the mean ± SEM, unpaired t test, *p< 0.05. **F)** Quantification of Myogenin positive cells from skeletal muscle collected at 7 days after NTX from wild type and MP^VEGFAKO^ animals; n=3, data represent the mean ± SEM, unpaired t test. **G)** Representative images of Codex panel from skeletal muscle collected at 3 days after NTX from wild type and MP^VEGFAKO^ animals. **H)** MP sorted from uninjured skeletal muscles of WT and MP^VEGFAKO^ mice were seeded at clonal density. Left: average number of cells forming one colony at different time points after cell seeding, normalized to the WT 3 days average. Right: absolute number of colonies (n=3, data represent the mean ± SEM, two-way ANOVA with multiple comparison, *p< 0.05, ***p< 0.001). **I)** Left: Representative pictures of myogenic differentiation in WT and MP^VEGFAKO^ cultures. Centre: frequency of myogenin positive cells (n=5, data represent the mean ± SEM, unpaired t test), and Right: fusion index (percentage of nuclei within syncytial structures) (n=5, data represent the mean ± SEM, unpaired t test).

To better understand how the deletion of VEGFA in MP affects muscle regeneration, we analyzed MP activation and differentiation by performing Pax7, Ki67, MyoD, and Myogenin staining at 3 and 7 days post injury. As expected from the previous results, MP^VEGFAKO^ mice displayed lower MP activation, shown by the decrease in the number of Ki67^+^/Pax7^+^ and MyoD^+^ cells at 3 days after damage (Figure 4C-D and G and S5B), which led to a dramatic decrease in the number of Myogenin^+^ cells (Figure 4E and G). However, consistent with the MP proliferation recovery in MP^VEGFAKO^, we did not notice any difference in the number of Myogenin^+^ cells between MP^VEGFAKO^ and WT at 7 days after injury (Figure 4F and S5C).

Recent studies have also described the role of VEGFA in supporting MP survival (Verma et al., 2021), thus we analysed the apoptosis in our systems and we noticed an increase in the number of MyoD^+^/Tunel^+^ MP in MP^VEGFAKO^ compared to WT animals (Figure S5D). This suggests that the reduction of number of differentiating MP at 3 days after injury, might be due to either, or both, an impairment of proliferation and increase of apoptosis caused by VEGFA depletion specifically in MPs.

The reduced MP proliferation in MP^VEGFAKO^ mice suggests that VEGFA may act through an autocrine loop in these cells. Indeed, MPs express VEGFA receptors *Kdr* (VEGFR2), and *Flt1* (VEGFR1), and *Nrp1* (Neuropilin-1) (Figure S5E). To confirm this hypothesis, we isolated MPs from MP^VEGFAKO^ and compared their bulk and clonal expansion as well as their myogenic differentiation against control cells. Impairment of MP proliferation was readily observed in bulk cultures at 7 days after cell seeding (Figure S5F). Clonal assays confirmed that VEGFA-depleted MPs were less capable of expansion than control cells, but displayed similar colony forming efficiency (Figure 4H), suggesting that the lack of VEGFA does not impair survival of quiescent MPs in vitro. When cells from MP^VEGFAKO^ and control mice were induced to differentiate, we observed no differences in either their myogenin expression (a measure of commitment to differentiation) or their ability to fuse into multinucleated fibers (Figure 4I), as indicated by the normal numbers of differentiating Myogenin^+^ cells counted at 7 days after NTX (Figure 4F). Thus, autocrine VEGFA acts directly on MPs by regulating their proliferative capacity early during regeneration.

### FAP-derived VEGFA is critical after ischemic damage

Our results suggest that FAP-derived VEGFA is dispensable in the regenerative response to myotoxin damage. However, we showed FAPs to be a robust source of VEGFA *in vivo*. Based on their proximity to vasculature and the recent report of a role for FAPs in ensuring vascular integrity following ischemic damage (Santini et al., 2020), we switched to a femoral artery ligation model (Fem Lig). Compared to NTX damage, this model is characterized by increased muscle fiber damage, vascular branching, endothelial proliferation, and activation of hypoxia related pathways, including the VEGFA signaling network (Figure S6A-E).

When compared to WT controls at 14 day post-injury, FAP^VEGFAKO^ displayed reduced adipose tissue formation and increased necrosis, while adipose tissue deposition increased with negligible necrosis in the MP^VEGFAKO^ (Figure 5A-F). Fiber size in WT and MP^VEGFAKO^ was quantified and revealed no significant difference (Figure 5G). Interestingly, and in line with the NTX damage model at 7days after injury (Figure 3D), fibers in the FAP^VEGFAKO^ model were larger than the control animals (Figure 5G and S7A). This divergent muscle repair across the three systems was observed already at 5 and 7 days after Fem Lig (Figure S7B). In summary, FAP-derived VEGFA plays a much more important role in muscle repair following ischemic compared to myotoxin damage, while the effects of MP-derived VEGFA are similar in the two models.

**Figure 5:**
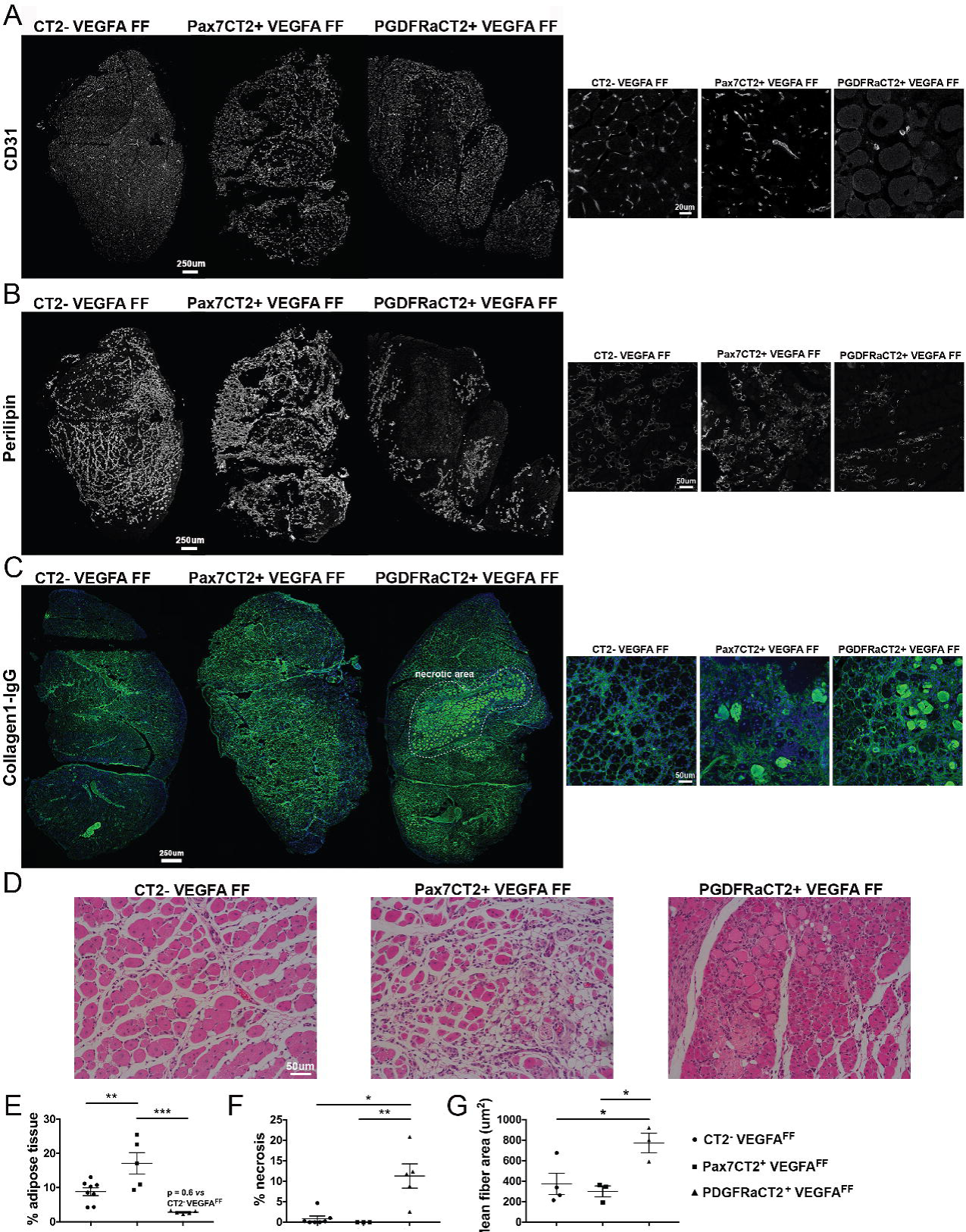
FAP-derived VEGFA is critical after ischemic damage. **A-C)** Representative images of vessels (CD31), adipocytes (perilipin), and both collagen (collagen1) and necrosis (IgG1) in skeletal muscle collected from WT, MP^VEGFAKO^, and FAP^VEGFAKO^ 14 days after femoral ligation (Fem Lig). **D)** H&E of skeletal muscles collected from WT, MP^VEGFAKO^, and FAP^VEGFAKO^ at 14 days after Fem Lig. **E-F)** Quantification of adipose tissue and extent of necrosis based on perilipin and IgG1 staining, respectively (n=3-8, data represent the mean ± SEM, one-way ANOVA with multiple comparison, *p< 0.05, **p< 0.01, ***p< 0.001). **G)** Quantification of muscle fiber cross sectional area 14 days after Fem Lig (for skeletal muscle fiber n=3-4, data represent the mean ± SEM, one-way ANOVA, *p< 0.05).

As ischemic damage results in extensive vascular remodeling, we focused on the effects of MP or FAP VEGFA depletion on endothelial cells. Similar to what was observed following NTX injury, EC proliferation was not affected by in MP^VEGFAKO^ mice after ischemic damage (Figure 6A). Conversely, VEGFA depletion from FAPs significantly impaired EC proliferation 4 days after surgery, and led to the appearance of large decellularized areas of muscle with sparse vasculature (Figure 6B-F).

**Figure 6:**
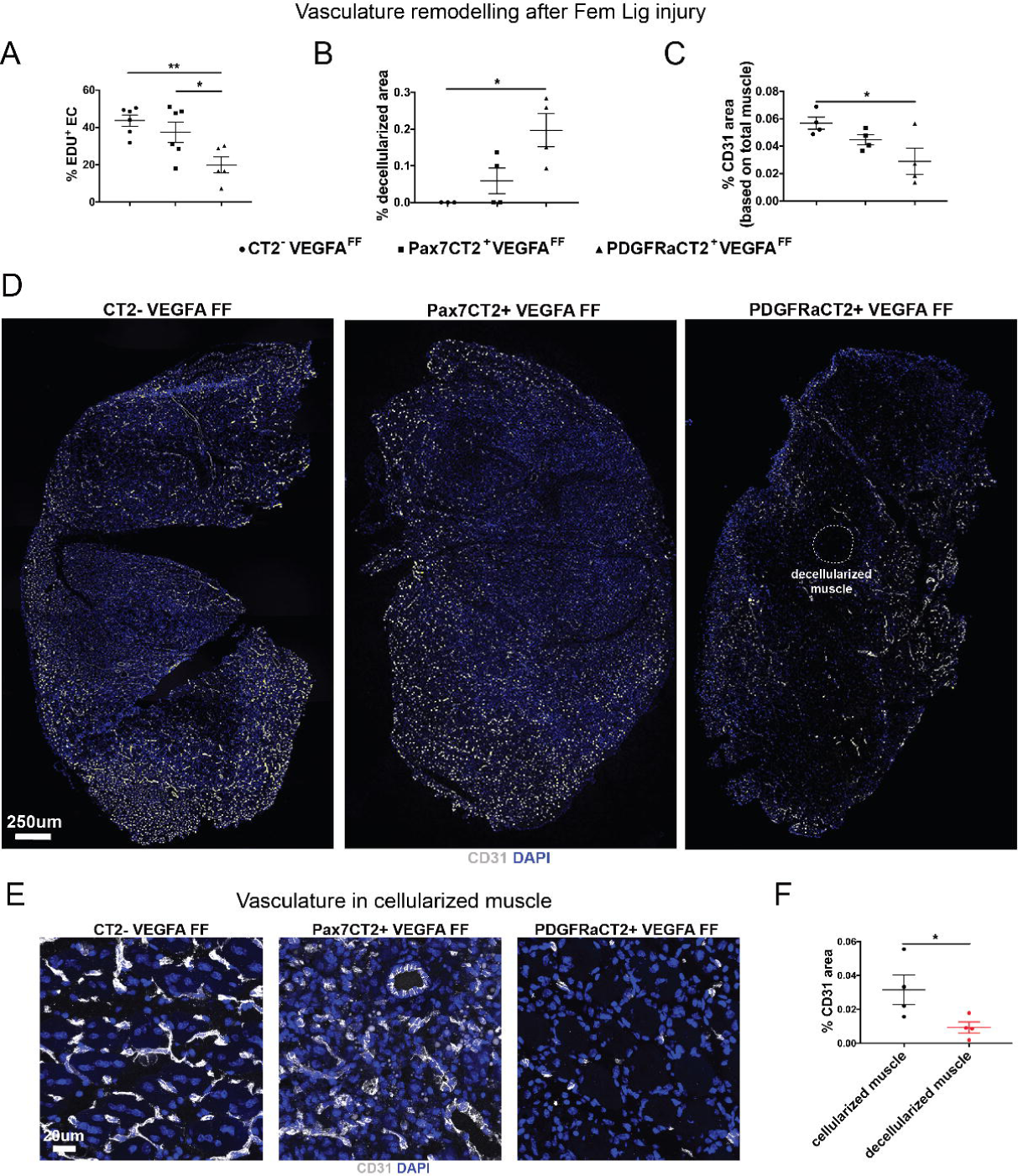
VEGFA depletion in FAP, but not in MP, affected vascular proliferation after ischemic damage. **A)** EdU incorporation in EC sorted from skeletal muscles of WT, MP^VEGFAKO^, and FAP^VEGFAKO^ mice 4 days after ischemic injury and 24 hrs after EdU treatment. (n=5-6, data represent the mean ± SEM, one-way ANOVA with multiple comparison, *p<0.05, **p<0.01). **B-C)** Histological quantification of decellularized and vascularized areas in skeletal muscles collected from WT, MP^VEGFAKO^, and FAP^VEGFAKO^ mice at 7 days after NTX (n=4, data represent the mean ± SEM, Kruskal-Wallis test or one-way ANOVA with multiple comparison, *p<0.05). **D-E)** Representative images of skeletal muscles collected from WT, MP^VEGFAKO^, and FAP^VEGFAKO^ mice 7 days after ischemic injury. **F)** Histological quantification of vasculature in cellularized and decellularized areas in muscles collected from FAP^VEGFAKO^ mice 7 days post damage (n=4, data represent the mean ± SEM, Kruskal-Wallis test or one-way ANOVA with multiple comparison, *p<0.05).

To test whether the vasculature was efficiently perfused we intravenously injected fluorescent lectin, which labels endothelial cells. Seven days after ligation surgery, the extent of labelling of CD31^+^ cells was assessed, revealing a strong damage-induced reduction in lectin labelling after ischemic damage, but no significant difference between FAP^VEGFAKO^ and WT mice (Figure S7C).

Next, we asked whether the same MP delayed proliferation phenotype that was evident in myotoxin damaged MP^VEGFAKO^ animals was also present after ischemic damage. Indeed, MP proliferation and total numbers were lower in MP^VEGFAKO^ vs WT two days after ischemic damage (Figure 7A). As with myotoxin damage, at later time points differences in MP proliferation were no longer detectable, and the difference in total MP numbers while still present, was significantly reduced. Thus, the phenotype in MP^VEGFAKO^ is the same independently of the type of damage.

**Figure 7:**
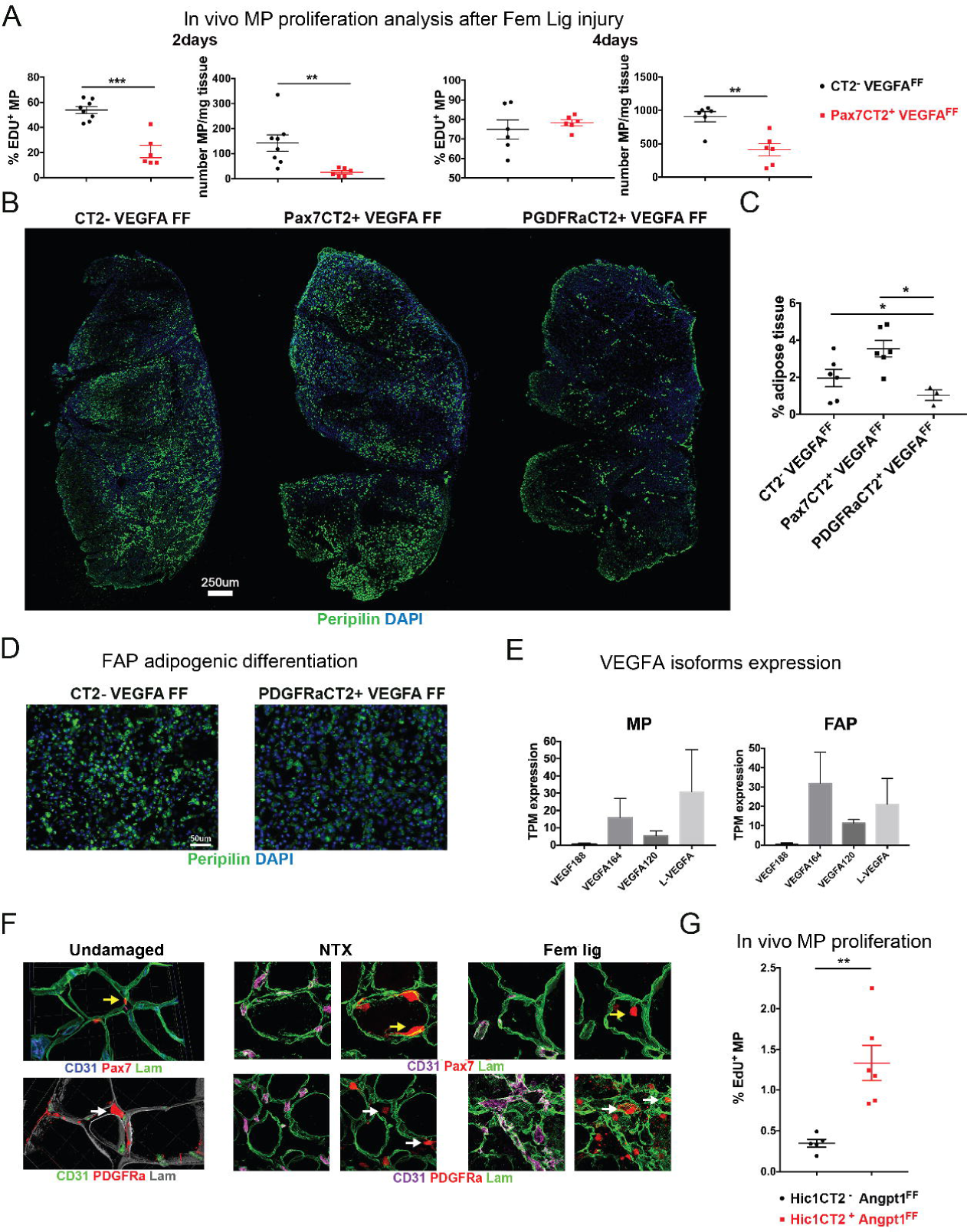
Spatial compartmentalization of signalling imparts source-specific functions on secreted factors. **A)** EdU incorporation in MP sorted from skeletal muscles of WT, MP^VEGFAKO^, and FAP^VEGFAKO^ mice 2 and 4 days after ischemic and 24 hrs after EdU treatment. (n=6-8, data represent the mean ± SEM, unpaired t test or Mann Whitney test, **p<0.01, ***p< 0.001). **B-C)** Histological staining of adipocytes (perilipin) and its quantification from skeletal muscles of WT, MP^VEGFAKO^, and FAP^VEGFAKO^ mice 5 days after ischemic injury (n=3-6, data represent the mean ± SEM, one-way ANOVA, *p<0.05). **D)** In vitro adipogenic differentiation of FAP sorted from uninjured WT and FAP*^VEGFAKO^* mice; the adipogenesis was identified with perilipin staining. **E)** Expression of VEGFA isoforms by MP and FAP purified from undamaged WT skeletal muscles. **F)** Histological staining of laminin (Lam), vessels (CD31), FAP (PDGFRa), and MP (Pax7) in uninjured, myotoxin- and ischemia-damaged WT skeletal muscles. **G)** EdU incorporation in MP purified from skeletal muscles of WT (Hic1CT2-Angpt1FF) and Angpt1 KO (Hic1CT2+Angpt1FF) animals at 3 weeks after Angpt1 depletion.

In contrast, FAP proliferation relative to WT control muscle was unchanged in FAP^VEGFAKO^ at all time points, suggesting that FAPs are not engaged in autocrine proliferative signaling through this factor (data not shown). However, we observed an increase in adipose tissue formation in TAs from MP^VEGFAKO^ mice and a decrease in FAP^VEGFAKO^ animals. These differences were already evident at 5 days (Figure 7B-C), and still present 14 days after surgery (Figure 5B and E). We asked whether a lack of VEGFA in FAPs may modulate their ability to differentiate. FAPs from WT and FAP^VEGFAKO^ were cultured and their proliferation and adipogenic potential compared. Consistent with the fact that FAPs do not express VEGFA receptors (Figure S5E), no differences were detected (Figure S7D and 7D).

### Basement membrane compartmentalizes skeletal muscle and controls intercellular networks

Based on our data, the ability of VEGFA to reach a nearby target cell appears limited by factors other than physical distance or the inherent ability of the target cell to respond to it. We hypothesized that these observations might be due to the presence of physical barriers that limit intercellular crosstalk by confining VEGFA availability. Interestingly, we found that both MPs and FAPs primarily express VEGFA*_164_*, an isoform with a high affinity for ECM, and its precursor L-VEGFA (Figure 7E) (Tee and Jaffe, 2001).

Thus, we focused our attention on the main ECM-rich structural barrier in skeletal muscle, the basement membrane (BM). At steady state, we found a hierarchy of compartmentalization with FAPs separated from ECs and MPs by a single layer of BM, while ECs were separated from MP by two layers (Figure 7F). In both myotoxin and ischemic damage, the integrity of the barrier between MPs and ECs was maintained, and MPs remained confined within the BM of the muscle fiber. Similarly, following NTX injury the positioning of ECs and FAPs relative to each other and the BM was left unperturbed, in agreement with the poor BM traversing by FAP described using spatial multiplexed imaging (Wang et al., 2022). In contrast, ischemic damage led to a remodeling of the BM, resulting in ECs and FAPs being embedded in the same BM layer. Such change would allow VEGFA from FAPs to be delivered directly to the endothelial BM, while VEGFA from MPs would adsorb to the BM ensheathing the myofiber and be prevented from reaching ECs.

While physical barriers like basement membrane might constrain specific ECM-bound ligands, like some VEGFA isoforms, they should not limit the freely diffusible ones. We investigated this focusing on another paracrine signal between FAP-produced Angiopoietin-1 (Angpt1), which lacks a matrix binding domain, and its receptor Tie2 (*Tek*), which is expressed by both MP and EC (Figure S3B and Table S2). We depleted Angpt1 in perivascular cells using a previously published Cre inducible system (Scott et al., 2019), and treated the mice with EdU for 3 weeks upon Angpt1 silencing. Interestingly, we observed an increased MP proliferation in mice whose FAPs lacked Angpt1 vs wild type animals, consistently with the known role of Angpt1/Tek in regulating MP quiescence (Figure 7G) (Abou-Khalil et al., 2009). This indicates that ECM represents a physical barrier for intercellular signaling occurring though ECM-ligands, like VEGFA isoforms, but not diffusible-ligand like Angpt1.

In summary, we have shown that despite the apparent high redundancy suggested by analyzing the expression of ligands and receptors, at least some signaling pathways are compartmentalized in adult tissues, and we propose that ECM-based structural barriers may play a role in determining who can signal to whom. Further work will be required to determine how widely this principle applies.

## DISCUSSION

The robustness of tissue regeneration depends on a web of interactions between different cells. Within this network, dysregulation or loss of function in just one of these cell types can lead to functional disruption of multiple other cell types, making it difficult to assign the observed phenotype to a specific molecular pathway. Macrophages in muscle regeneration are an excellent example, as they concomitantly signal to endothelium, stroma and myogenic cells (Arnold et al., 2007; Groppa et al., 2012; Lemos et al., 2015; Mojumdar et al., 2014; Ochoa et al., 2007; Villalta et al., 2014; Willenborg et al., 2012). Lack of infiltration by macrophages leads to regeneration failure, as observed in CCR2KO mice, but deconvolving the contribution of each of the target cell types to the overall phenotype and identifying its major driver has been daunted by the complexity of the processes involved. Here, by taking a time-resolved transcriptomic-based approach, we assessed the engagement of 5 key cell types in the regenerative process. We compared the kinetics of transcriptional change taking place during regeneration in wild type and CCR2KO mice and confirmed the disruption of normal FAP kinetics we previously reported. In addition, we found the lack of immune cell infiltration to have drastic impact on endothelial cells. A transcriptomic-based model of cellular interactions was built, and not surprisingly pointed to VEGFA as a key determinant of endothelial activity. These predictions, that vessel remodeling was disrupted in CCR2KO animals, and that specifically lack of VEGFA from inflammatory cells was a key determinant of this phenotype, were verified experimentally. Indeed, knocking out VEGFA from inflammatory cells was sufficient to lead to a phenotype vastly overlapping that of CCR2KO animals, including the muscle regeneration impairment but much reduced fibrosis/fat infiltration, which is likely due to a TNF-mediated direct effect on FAPs as we previously reported (Lemos et al., 2015).

However, our interactome model also raised new questions, by revealing that VEGFA was produced in comparable amounts by not just inflammatory cell, but also myogenic and stromal progenitors, all of which reside in close proximity to endothelial cells. This promiscuity was not limited to VEGFA, as more than three quarters of all examined ligands had multiple cellular sources within regenerating muscle, displaying a level of redundancy in specific pathways across different cell types analogous to those described in other recent studies (Rezza et al., 2016), and suggesting this redundancy underlies a general strategy in biology. This prompted the question of whether the observed VEGFA expression pattern in muscle regeneration represents true redundancy, in which a given factor carries out the same functions independently of which cell produced it, or conversely the cellular origin of a secreted molecule dictates its function in the context of the tissue. While the activity of redundant receptors can be validated bioinformatically (Alvarez et al., 2016; Martini et al., 2014), such as in our analysis of VEGFR2 (*Kdr*) in ECs and MPs, understanding the functionality of a ligand produced by multiple sources cannot be

investigated computationally. Thus, we generated additional mouse strains in which VEGFA was deleted specifically from MPs and FAPs, the two cell types, beyond inflammatory cells, that also produce it during regeneration.

Overall, our results show that VEGFA has multiple roles during muscle regeneration, and that these functions are tightly linked with the cell type that produces it. The type of damage inflicted determines which of these functions, and therefore sources, are critical. In particular, VEGFA from FAPs is dispensable in non-ischemic damage models, such as myotoxin injury, but it is critical for the vascular remodeling that takes place after an ischemic event. Likely, the reason for this is that myotoxin injury induces a relatively mild angiogenic response when compared to the ischemic trauma (Plant et al., 2006). Thus, VEGFA produced by ICs is sufficient to preserve the vasculature after NTX injection. However, in cases of severe ischemic damage, additional sources of VEGFA are required for a robust vascular response and FAP-derived VEGFA becomes crucial.

Most importantly, our data indicate that the target of VEGFA signaling is specified by the cell type of origin. Thus, VEGFA from MPs acts in an autocrine fashion to support the early proliferation of these cells in both ischemic and myotoxin damage models, but is dispensable for vascular remodeling. We did not investigate the mechanism with which VEGFA affects myogenic stem cell activity early after damage, but its ability to modulate symmetric vs asymmetric divisions, as described in Chen et al (Chen et al.), is likely to be involved. Surprisingly, despite the close proximity of myogenic and stromal cells in situ and the fact that they produce similar amounts of the protein, VEGFA from MPs cannot rescue the lack of VEGFA from FAPs and vice versa. Thus, signaling networks that appear to be shared across multiple cell types based on expression data, are in reality compartmentalized in adult tissues, and that the ability of one cell to signal to another cell is tightly controlled.

Hypothetically, this level of control could stem from the integration of antagonistic signaling pathways within the target cells, by which a given cell type may be unresponsive to a secreted factor even when the latter is present in the extracellular space and capable of engaging its receptors. However, we clearly show that MPs can respond to autocrine VEGFA, and a pro-angiogenic effect of MPs is readily observed when co-cultured with ECs (Mettouchi, 2012; Rhoads et al., 2013).

Alternatively, secreted factors could be prevented from reaching a specific target cell by anatomical barriers to its diffusion. Indeed, the importance of muscle architectural organization was elegantly demonstrated in the work of Webster *et al*. (Webster et al., 2016), where intravital imaging was used to show that the myofiber’s basement membrane remains intact after the damaged fiber itself has been removed (ghost fibers), in order to guide *de novo* myogenesis.

Activated MPs are in constant contact with the interior surface of these ghost fibers and are therefore essentially trapped in them. Our data suggest that such a barrier prevents MP-expressed VEGFA from reaching the endothelium, which is consistent with the notion that VEGFA can be locally constrained through anchoring to the ECM via its heparin binding domain (Ruhrberg et al., 2002). Indeed, previous works have described how VEGFA isoforms capable of freely diffusing within their microenvironment induce vascular remodeling that is markedly different compared to VEGFA isoforms that remain anchored in the ECM.

A limitation of our study is that we did not include factors expressed by myofibers, like VEGF, in our analysis. However, when this factor was specifically deleted in adult fibers, the only reported effect was mainly on the fiber’s ability to respond to exercise. No change in vascular density was seen, supporting our hypothesis that in adult tissue VEGF cannot act outside of the ensheathing basement membrane (Delavar et al., 2014; Knapp et al., 2016).

Using a MP-specific ad hoc VEGFA depletion model, Verma *et al*. described how VEGFA produced by MPs can interact with ECs, which in turn communicate via Notch signaling to regulate MP quiescence and vascular proximity (Verma et al., 2018). Though our data align on the importance of VEGFA signaling in regeneration, and its critical effect on the maintenance of proper MP activity and numbers, our conclusions regarding the signaling networks through which it acts differ. Indeed, while Verma *et al*. describe FAPs to be an equally viable source of VEGFA compared to MPs, in their experiments like in our hands FAP-derived VEGFA fails to rescue the phenotype observed in MP-derived VEGFA depletion, thus reinforcing our confidence in the critical role for signal compartmentalization in this system.

Though current bioinformatic approaches provide outstanding tools to predict ligand-receptor pairing, our work clearly shows that not all signaling interactions can be predicted based on expression data alone as additional factors should be kept in account when modelling muscle regeneration. Here, we describe how the presence of structural barriers and the type/severity of damage are all important elements. We propose that the interposition of stable, matrix-rich structures like the BM can limit the diffusion of signalling molecules, thus assigning distinct roles to cell subsets secreting the same ligand. In addition, the type of injury can also modify such structural barriers, thus shaping the cell signaling. First, the damage itself can have a direct impact on the re-organization of the BM as observed in our systems, where EC and FAP are separated by BM after toxic injury, but they are completely embedded in the same BM layer after ischemic damage. Second, the cellular target of the injury also dictates its consequences. In direct myotoxic damage (NTX), which affects only muscle fibers, circulating inflammatory cells are recruited to the tissue through unperturbed blood vessels, where they provide cytokines to regulate vascular remodeling and controlling FAP-activation (Theret et al., 2022). In contrast, in an ischemic injury, inflammatory cells infiltration might be reduced due to the vessel impairment (Germani et al., 2003). The different inflammatory cells recruitment itself could cause a lack of factors that leads to compensating signaling mechanisms. Back to our system, after NTX injury, VEGFA is provided by inflammatory cells to promote vascular remodeling whereas FAP-derived VEGFA is dispensable. The situation is completely different after Fem Lig damage, where damage to blood vessels is more extensive, and VEGFA by FAP becomes pivotal for muscle repair, probably because of an increase requirement for this factor.

Inclusion of these parameters, and surely others yet to be defined, will be required to more faithfully represent and predict the behavior of complex cellular systems in the future. Computational modeling is now part of many attempts to faithfully describe biological systems. However, it invariably considers biological networks in a homogeneous spatial environment. Incorporating structural features into these mathematical models is a new area of research (Getz et al., 2018). This could be integrated with innovative multiplex imaging able to simultaneously profile the spatial distribution of cell surface, intracellular and ECM proteins (Wang et al., 2022).

Finally, it is interesting to speculate that the compartmentalization of signaling described here is one of the factors that allows the same pathways to be re-utilized in multiple distinct biological processes.

## STAR⍰METHODS

### LEAD CONTACT AND MATERIAL AVAILIBILITY

Further information and requests for resources and reagents should be directed to and will be fulfilled by Dr. Elena Groppa and Dr. Fabio Rossi. This study did not generate new unique reagents.

### EXPERIMENTAL MODEL AND SUBJECT DETAILS

#### In vivo Mouse models

Either B6.Cg-Pax7^tm1(cre/ERT2)Gaka^/J (Jax stock number 017763) or Pdgfra^CreERT2^mice (kind gift from Dr. Brigid Hogan) or Tg(VAV1-cre)1Graf (Jax stock number 127936) were crossed with B6.Cg-Gt(ROSA)26Sor^tm14(CAG-tdTomato)Hze^/J (Jax stock number 007914) and VEGFa^tm2Gne^ (kindly provided by Dr. Richard Lang from Cincinnati Children’s Hospital) to generate Pax7 CT2 or Pdgfra CT2 VEGFa^flox/flox^ tdTomato mice. We used other mouse models like B6.129S4-Pdgfra^tm11(EGFP)Sor^/J (Jax stock number 007669; here in referred to as Pdgfra eGFP) and Tg(Cspg4-DsRed.T1)1Akik/J (Jax stock number 008241; herein referred to as NG2 DsRed), which were maintained either in a wild type B6 background (PDGFRa EGFP WT and NG2 DsRed WT) or bred with B6.129s4-ccr2^TM1lfc^/J (CCR2KO): (Jax stock number 004999; herein referred to as CCR2KO) to generate PDGFRa EGFP CCR2KO. Mice were housed, in a pathogen free facility, under standard conditions (12 hrs light/dark cycle). All experiments conducted in accordance to ethical treatment standards of the Animal Care Committee at the University of British Columbia. Allelic recombination was induced by daily injections of 0.1 mg/g tamoxifen (TAM) in 100 uL of corn oil for 5 consecutive days, administered intraperitoneally to 8 weeks old animals. To control TAM toxicity, all mice including controls were subjected to experimental procedures 2 weeks after TAM to allow a washout period. Male and female mice were equally mixed in our experiments.

#### Myogenic Progenitors primary culture

Myogenic progenitors (MPs) were extracted as described in the Method Details session and sorted as CD45-/CD31-/Sca1-/a7integrin+/VCAM+.

Clonal assay and Myogenic differentiation were assessed as previously described (Theret et al., 2017).

Cells were then cultured on a Matrigel®-coated (1:10, Corning 354234) 48 well-plate in proliferative media (Dulbecco’s modified eagle medium (DMEM/F12; Thermofisher), 20% fetal bovine serum (FBS), 0.75 ug/mL bFGF (PeproTech), 1% Penicillin/Streptomycin (Thermofisher). For clonal growth assay, sorted MPs were seeded at 50 cells per well in the proliferative media on a Matrigel®-coated 48 well plate and the number of clones and number of cells per each clone were quantified from 3 to 6 days after sort.

For differentiation assay, MPs were seeded at 3,000□cells/cm² for expansion for 5–7□days in proliferation media on Matrigel-Coated plates. After a passage, cells were seeded at 30,000□cells/cm² in proliferative media for 6□hours and then switched to a differentiation media (DMEM/F12, 2% Horse Serum (Sigma), 1% Penicillin/Streptomycin). The levels differentiation and fusions were assessed after 3 days. Cells were fixated in 4% paraformaldehyde (PFA) for 7 min, then incubated with blocking buffer (1X PBS, 3%Goat serum, 0.3% Triton X-100) for one hour before being incubated overnight at 4°C with Myogenin (AbLab, clone F5D) and Desmin (Abcam) antibodies. Then, cells were washed in PBS, incubated with secondaries antibodies (Invitrogen) and counterstained with 600nM DAPI for 5 minutes. The number of positive cells for Myogenin and number of nuclei per myotubes were then quantified.

### Fibro/Adipogenic Progenitors primary culture

For in vitro differentiation studies, skeletal muscle from WT and FAP^VEGFAKO^ mice were digested as described in the Method Details session and sorted as CD45-/CD31-/Sca1+/PDGFRa+. FAPs were seeded at a density of 10,000 cell/cm2 in high–glucose Dulbecco’s modified eagle medium (DMEM) (Invitrogen) supplemented with 10% FBS, 1M Sodium Pyruvate, and 2.5 ng/ml bFGF (Invitrogen). Once FAPs reached 80% confluency, adipogenic differentiation was induced with MesenCult^TM^ adipogenic differentiation media (STEM CELL) until day 5. Half of the media was changed every two days.

### METHOD DETAILS

#### Skeletal muscle injury models

Between 10-11 weeks of age, mice received an intramuscular injection of notexin (0.15 ug/TA, Latoxan) or underwent femoral ligation surgery. In femoral ligation, a skin incision was made over the femoral artery beginning at the inguinal ligament and continued caudally to the popliteal bifurcation. The ligation site on femoral artery was identified and the femoral vein and nerve along with connecting tissue were separated away from the artery of the ligation point with tip forceps.□Once the ligation site was clear of vein and nerve, an 8-0 silk suture was used to ligate around the femoral artery between epigastric artery and popliteal branch.□The skin incision was closed with simple interrupted pattern with 6-0 Vicarly suture (Padgett et al., 2016). The animals were placed in a clean cage with heating support to recover from anesthesia. Once fully recovered, the animals were returned to its original cage with easily accessible food and hydrogel on the cage bottom. Hindlimbs were collected 3-28 days after damage according to the experiment.

#### Tissue digestion and cell purification after notexin damage

Mice were sacrificed and their hindlimbs were cut into 2 mm pieces. A single cell suspension was made by a 30-minute incubation at 37°C in Collagenase type II solution (Millipore Sigma; 2.5 U/mL) activated by 10 mM CaCl_2_ followed by centrifugation at 360 G (5 minutes at 4°C). The cells were then incubated in a mixture of Collagenase D (Roche Biochemicals; 1.5 U/ml) and Dispase II (Roche Bio chemicals; 2.4 U/ml) and agitated every 15 minutes. Reaction was stopped by subsequent addition of FACS buffer (PBD, 2mM EDTA, 2% FBS) followed by centrifugation at 1700 rpm (4°C for 5 minutes). Red cells were lysed by Ammonium-Chloride-Potassium (ACK) buffer (Lonza; 1mL) followed by addition of FACS buffer to stop the reaction. Cells were incubated for 25-30 minutes at 4°C in the primary antibody mixture in FACS buffer ∼ 3 x 10^7^ cells/mL. Muscle mononucleated cells were stained with FITC anti-CD45 or APC anti-CD45 (AbLab; clone I3/2), FITC anti-CD31 or APC anti-CD31 (eBioscience; clone 390), PECy7 anti-Sca1 (eBioscience; clone D7), biotinylated anti-VCAM (AbLab; clone 429 MVCAM.A) followed by streptavadin-PE (Caltag Laboratories #SA1004-4), APC anti-α7integrin (AbLab; clone R2F2) and Hoechst 33342 (Sigma #B2261; 2.5ug/ml). Influx I or II cytometer (BD) sorters were used. From both PDGFRa EGFP WT and CCR2KO, FAP as CD45-/CD31-/EGFP+, inflammatory cells as CD45+/CD31-, myogenic progenitors as CD45-/CD31-/a7+/VCAM+, and endothelial cells as CD45-/CD31+ were sorted. From NG2 DsRed WT, pericytes as CD45-/CD31-/Sca1-/NG2+ were sorted. Whole skeletal muscle and cells were sorted at 0, 1, 2, 3, 4, 5, 6, 7, 10, 14 days after injury with notexin.

#### RNA extractions from sorted cell populations

Sorted cell populations were rinsed in diethyl pyrocarbonate in PBS (DEPC PBS; Invitrogen) and then, pelleted and resuspended (800G; 10 minutes; 4°C) twice. They were then resuspended in RNAzol (Sigma Aldrich; #R4533; 1 mL/1 x 10^7^ cells) followed by addition of diethyl pyrocarbonate (DEPC) water (400ul aqueous/1m RNAzol) and incubated for 15 minutes at room temperature. Samples were then centrifuged for 15 minutes (12000 G; 4°C) and supernatant was precipitated using an equal volume of isopropanol and 1 uL of linear acrylamide at -20°C. On the next day, samples were centrifuged (20000 G; 4°C) and the pellet was washed 3 times with 70% ethanol (diluted in DEPC water) and left to airdry. RNA solutions were stored at -80°C in DEPC water and Superase (Applied Biosystem #100021540; 1:20 dilution).

Flash frozen muscles were mixed in RNAzol (1 mL/TA muscle) and homogenized using a sonicator (Omni International). Lysates were agitated followed by addition of DEPC water (400 uL) and incubated for 15 minutes at room temperature. All samples were centrifuged for 15 minutes (12000 G; 4°C) and supernatant was precipitated using an equal volume of isopropanol at -20°C and incubated for 10 minutes at room temperature. Lysates were spun at 8000 G for 10 minutes at 4°C and washed and pelleted 3 times with 70% ethanol (diluted in DEPC water). Following airdrying, the pellet was resuspended in DEPC water (100 uL/TA) and Superase (1:20 dilution) and stored at -80°C until use.

#### Perfusion and tissue collection

Mice were anesthetized with 0.5 mg/g tribromoethanol (Avertin) and a horizontal incision was made above the sternum through the skin and the musculoskeletal layer, to expose the chest cavity and the heart. The right atrium was punctured prior to insertion of a 26 1/2-gauge needle (BD) into the left ventricle. Perfusion was performed using a controlled pressure of 120 mmHg for 3 minutes using freshly prepared 1% PFA for fixation, followed by 4 minutes of perfusion with phosphate buffered saline (PBS). Skin and fascia were removed from lower limb of the mice, and the hindlimb muscles were harvested from each leg. Whole muscles were stored in 0.5% PFA at 4°C overnight and then transferred into 40% sucrose solution in PBS at 4°C for 24 hours. Tissues were then embedded into blocks of O.C.T. Compound (Tissue-Tek) using isopentane cooled with liquid nitrogen. All tissues were then stored at -80□C. For histological analysis, embedded O.C.T blocks were equilibrated at -20°C and cut into 10 um sections, ensuring distribution of entire tissue on each slide. Slides were then stored at -80°C until use.

For vascular leakage analysis, each mouse was intravenously injected with 5mg of dextran-Texas Red (50mg/ml) (Invitrogen #D1864) 10 minutes before tissue harvesting.

#### Immunofluorescent staining on frozen sections

Previously cut samples, were brought into room temperature. Autofluorescence was quenched using Sodium Borohydride (Sigma 213462; 10mg/mL) for 1 hour and followed by blocking in 2% goat or donkey serum and 0.03% Triton X-100 (Sigma) in PBS for 1 hour. Sections were then incubated in primary antibodies diluted in the previously used blocking buffer for 1 hour or overnight. Slides were washed three times with 0.03% Triton X-100 (PBST) followed by a 1-hour incubation in Secondary antibody mixtures in PBST. Primary antibodies used were rabbit anti-laminin (Abcam #15575), mouse anti-collagen (Abcam #ab90395), rat and goat anti-CD31 (Biolegend clone 390 and R&D #AF3628, respectively) and rabbit anti-perilipin (Abcam #ab3526). Secondary antibodies included, donkey anti-rabbit 488 (Invitorgen #A21206), donkey anti-goat 488 (Invitrogen #A11055), donkey anti-rabbit 647 (Invitrogen #A31573), donkey anti-goat 647 (Life Technologies #A21447), as well as goat anti-rabbit 488 (Life Technologies #A11034), goat anti-rat 488 (Invitrogen A11006), goat anti-rabbit 647 (Invitrogen #A21245). Staining of damage/necrotic areas was performed with donkey anti-mouse IgG1 (Life Technologies #1305303) staining in combination with mouse anti mouse collagen 1. Tissue sections were washed with PBS 3 times for 5 minutes each and transferred to a beaker of PBS for 10 minutes followed by a 10-minute incubation in DAPI (Sigma; 1:1000). Cover slips were mounted on slides using Fluoromount-G (Southern Biotech) and stored at 4°C.

#### Cytospinned cells and staining

Each mouse was treated with 250ug Brefeldin A (Invitrogen) in 150ul of high glucose DMEM. At 12 hrs after the treatment, mice were sacrificed, muscle tissue collected and digested as previously indicated. Different cell types were purified based on the sorting strategy applied for generating bulk sequencing data, and cytospinned on slides. Staining for VEGFA was performed according to the immunofluorescence protocol above described, using primary antibody rabbit anti-VEGFA (Abcam #ab52917).

#### Myogenic cell immunofluorescence and TUNEL staining on frozen sections

Frozen mouse TA sections were washed briefly with PBS, fixed with 4% PFA in PBS, followed by quenching and permeabilization with 100mM glycine and 0.5% Triton X-100 in TBS. The buffer with 4% goat serum, 2% bovine serum albumin, 4% MOM blocking reagent (Vector Labs #MKB22131), 0.1% Tween-20 in TBS, was used for blocking the TA section. The same solution without the MOM reagent was used for primary and secondary antibody staining. Rabbit anti-MyoD (Abcam #ab133627), mouse anti-Pax7 (# 1/10 AbLab), and rabbit anti-laminin (Abcam #ab11575) antibodies were used for immunofluorescence staining overnight at 4°C, along with goat anti-rabbit IgG conjugated with either Alexa488 (ThermoFisher #A11034) or Alexa647 (ThermoFisher #A21245) and goat anti-mouse IgG1 Alexa488 (ThermoFisher #A21121) as secondary antibodies as the following step for 1h at room temperature the following day. Prior to mounting, nuclei on all sections were counterstained with 600nM DAPI in TBS for 5min at room temperature.

Apoptotic myogenic cell assays were performed with a TUNEL staining kit (ABP Biosciences A052). TdT catalysed ligation of biotin-conjugated dUTP was performed at 37°C for 90min as an additional step between permeabilization and blocking, with additional TBS wash steps before and after the reaction. Streptavidin-conjugated AndyFluor647 was added at the same time as secondary antibodies for immunofluorescence. Each time the experiment was conducted, one section was allocated for negative control staining where no TdT enzyme was added, but streptavidin-conjugated AndyFluor647 was still added at the later step. All fluorescence images were acquired with the Nikon Eclipse and Zeiss LSM900 microscopes, quantification was performed in Fiji software.

#### H&E staining

Muscles were harvested, sequentially fixed in 1% paraformaldehyde (PFA) and incubated in 70% ethanol. Muscles were embedded in paraffin and then sectioned at a thickness of 4 µm using a Leica microtome (RM 2255). Sections were stained with Hematoxylin and Eosin (H&E) (Waxit).

#### Image acquisition

Tissue sections were visualized on a Y-IFP fluorescent microscope (Nikon Instruments) and a C1 laser-scanning confocal microscope (Nikon Instruments; Eclipse Ti) equipped with lasers at 405, 488, 568, and 633 nm. NIS Elements software was used on both microscopes for multichannel image acquisition procedure. Single immunofluorescent images were captured using an auto-exposure feature and a hardware gain of 1. Large stitched images were acquired by a 10% overlap and manual refocusing in every 3 frames using the NIS Ar Elements Software (Nikon Instruments). Automatic post processing and shading correction was used to enhance the stitched images. In histology images white balance was set automatically using a white light background. High resolution images were taken using the confocal imaging (Zeiss; Axio Observer).

#### Histological quantification

Images chosen for analysis were selected based on representativeness in the whole data as well as quality of the tissue section. All quantifications were performed with the NIS Ar Elements software using 3 or more biological replicates for the respective experimental and control groups. Fiber size quantification was performed in notexin-damaged skeletal muscles, where regenerating fibers were identified by presence of center-located nuclei and infiltrating interstitial cells in the fiber, on a laminin stained section. All fibers in 10 images (20X; 300 um^2^) were quantified per animal. Vascular density quantification was performed on both damage models, where CD31 positive structures were quantified on an immunofluorescent-labeled slide. In notexin-damaged muscles, the number of vascular structures was divided by the total section area and 10 images (20X) were quantified from each animal. To characterize the phenotype of mice with femoral ligation, two whole-section stitched images/mouse were quantified. Here, due to vascular irregularities, the percentage of area positive for CD31 was calculated rather than using number of CD31+ vessels per area as applied for notexin injury model. Similarly, to quantify adipose tissue and necrotic areas in skeletal muscle collected from mice with femoral ligation, we calculated the percentage of area positive for perilipin and IgG staining, respectively. Last, the percentage of decellularized area was quantified based on low nuclei numbers (by DAPI staining). All quantifications were carried out using an automatic system where positive structures were masked by an intensity threshold.

#### Multiplexed immunofluorescence co-detection (CODEX)

Antibody conjugation, validation, tissue staining, image processing and data analysis were performed following Palla et al., 2020 (Palla et al., 2021).

#### Perilipin staining

After FAP-adipogenic differentiation, media was removed, cells were quickly rinsed with 1X PBS before being fixed with 4% pFA for 7min at room temperature. After 2 more wash, blocking buffer (1X PBS 0.3% Triton X100, 3% goat serum) was applied for an hour at room temperature and primary antibody (Perilipin Abcam #ab3526) was incubated overnight at 4◦C. Goat anti-rabbit IgG(H+L) Alexa 488 (Invitrogen #A11034) was used as a secondary antibody. DAPI was used to counterstain the cells. Plates were kept at 4◦C until imaging Cells were imaged using an ECHO Revolve microscope.

#### VEGFA ELISA experiment

Snap frozen muscles from wild type or knockout mice were collected at steady state or after injury. Tissue were homogenized (25 Hz; 30 minutes; Qiagen 85300) in 500ul PBS in proteinase inhibitor (1x). Protein quantification was performed using BCA assay kit (Thermofisher #23225). The ELISA assay was performed according to the instructions on the ELISA kit (R&D systems; MMV00) using 150ug for each sample.

#### Fluorescent Activated Cell Sorting (FACS) analysis

In vivo proliferation assays were performed by intraperitoneal injection of 0.5 mg (10 ug/mg) of Ethidium Bromide (EdU; Thermofisher #C10337) 12 and 24 hours before sacrificing the animals. Samples were digested as previously indicated. Surface and EdU staining was performed according to the manufacturer’s instructions. In vascular perfusion analysis, mice were treated with 50 uL of lectin-fluorescein isothiocyanate (Sigma #L0401) diluted with 50 uL of PBS. Mice were sacrificed after 10 minutes and tissues were collected, digested, and stained as previously indicated.

#### RNAseq bioinformatics analyses

##### Bulk-RNA sequencing

RNA quality control was performed with Agilent (Santa Clara, CA, USA) 2100 Bioanalyzer. Qualifying samples (n = 2–6) were then prepped following the standard protocol for the TruSeq stranded mRNA library kit (Illumina, San Diego, CA, USA) on the Illumina Neoprep automated nanofluidic library prep instrument or NEBnext Ultra ii Stranded mRNA (New England Biolabs, Ipswich, MA, USA). Sequencing was performed on the Illumina NextSeq 500 with Paired End 42bp × 42bp reads. Demultiplexed read sequences were then aligned to the Mus Musculus (PAR-masked)/mm10 reference sequence using TopHat splice junction mapper with Bowtie 2 (http://ccb.jhu.edu/ software/tophat/index.shtml) or STAR (https://www.ncbi.nlm. nih.gov/pubmed/23104886) aligners.

##### Filtering and normalization step

In our analysis, we discarded genes that did not have more than 100 raw counts in at least one time point. Additionally, we kept only genes with an expression average of 3 RPKM. With the remaining genes, we performed RUV normalization using RUVSeq R Package (version 1.14.0) to remove unwanted variation from our datasets (Risso et al., 2014). We used RUVg function with 3000 invariant genes and Table S3 summarizes the parameters applied in the RUV analysis for each dataset.

##### PCA and cluster analysis

PCA plots were made using plot PCA function from EDAseq R package [10.1186/1471-2105-12-480]. Using kmeans (stats R package) we clustered the samples in the PC score space. We empirically chose the optimal number of cluster k, with k={2, …, 10}, using two indexes: silhouette index as implemented in NbClust function from NbClust R package (Charrad et al., 2014) and the biological homogeneity index (BHI) as implemented in BHI function from clValid R package (Brock et al., 2008). The BHI was computed using the time/day. In brief, the higher the BHI the most homogeneous are the days within the cluster. When silhouette and BHI gives similar results, we preferred the lower number of clusters.

##### Differential Expression analysis

###### Time-wise analysis for cell-population and total muscle

Differential expression analysis was performed on all cell populations and in the total tissue. After filtering our datasets, we computed a set of time-invariant genes defined as the 3000 genes with the highest p-values. Using RUVSeq coefficients, we performed differential expression analysis using edgeR (version 3.22.3) (Robinson et al., 2010). We considered differentially expressed the genes with an adjusted p-value <= 0.01 and an absolute log2 Fold Change >= 2 in at least one comparison of the time points.

###### Cell-wise comparison

Differential expression analysis was performed among cells. After filtering our datasets, we computed a set of genes defined as 3000 genes with the highest p-value in the cell-wise comparison. Using RUVSeq coefficients, we performed differential expression analysis using edgeR. We considered differentially expressed the genes with an adjusted p-value <= 0.01 and an absolute log2 Fold Change >= 2 in any comparison among cell populations.

##### Time-wise clustering

Differentially expressed genes (DEG) from each cell population and whole tissue were clustered using TCseq R package (version 1.4.0) (http://bioconductor.org/packages/release/bioc/html/TCseq.html). We choose to organize DEGs into 9 clusters given the presence of 10 time points for nearly all cell populations sorted and whole tissue.

##### Identification of the “active” genes

Based on the centroids of the cluster, we identified the time-point were the cluster is activated using binarizeTimeSeries function from BoolNet R package (version 2.1.4) (Müssel et al., 2010). This method is based on kmeans and computes a cutoff according to the centroid values, thereby marking with active=1 those time points above the cutoff and active=0 otherwise. All the genes of the cluster inherited the activation pattern of the cluster centroid, therefore we defined a gene active in the population at day d (pop-active_d_) if it belongs to one cluster “active” at d.

We applied this for all cell population and the whole tissue. The pop-active genes in the whole tissue are used at point 1) below and for the GO enrichment (see *Gene Ontology Biological Process enrichment analysis)*

Along with pop-active_d_, we created a list of genes called “subset-specific” that potentially reflect cell subset expansion. These genes are 1) DEGs in the whole tissue time wise comparison and 2) significantly more expressed in one cell type than other two cell types (adjusted p-value<= 0.01 and an absolute log2 Fold Change >= 2). Like pop-active genes, subset-specific genes inherit the temporal activation pattern of the centroid of the cluster they belong in the whole tissue. Genes that were significantly more expressed in one cell type than other two cell types but not DEG in whole muscle, were defined as “constitutively-active” genes and assumed to be always active (active at all time points) because their functions are always required by the cell population.

##### Ligand-Receptor network analysis

The Ligand-Receptor Network (LRN) was built starting from a manually curated ligand receptor interaction dataset from Rezza *et al*. (Rezza et al., 2016). As described for all genes, we defined the ligands/receptors that are “active” along the time: pop-active_d_ are the ligands/receptors active at day d in the cell batch, subset-specific_d_ are the ligands/receptors that are mainly expressed by a cell type where they are either DEGs in the whole tissue or constitutively-active. At any day d, and for each cell type cell, we defined the autocrine cell network at day d (cell-AN_d_) as the induced subnetwork obtained from the LRN and active ligands/receptors (pop-active_d_cell_ + subset-specific_d_cell_ + constitutively-active_cell_ ligands and receptors). Paracrine cell networks (cell-PN_d_) were defined as the induced subnetwork obtained from LRN, ligands “active” at day d from all the cell and receptors active at day d in the cell (ligands: pop-active_d_ + subset-specific_d_ + constitutively-active; receptors: pop-active_d_cell_ + subset-specific_d_cell_ + constitutively-active_cell_).

##### Gene Ontology Biological Process enrichment analysis

To understand the function of the genes active at a given day, we performed a day by day GO enrichment analysis. The enrichment was performed on ‘pop-active’ genes at any day. We limited the analysis to GO Biological Process (GO BP) categories with 10 to 500 gene members (GO definition from org.Mm.eg.db version 3.8.2).

###### GO BP on whole muscle

Enrichment was made for each time point using the ‘pop-active’ genes in the whole tissue, i.e. those genes that belong to a cluster that is marked as active at a given day in the whole muscle. For this purpose, we used compareCluster from clusterProfiler R package (Yu et al., 2012), which allows to directly compare multiple list of genes. We compared the lists of ‘pop-active’ genes at any time point. Among significant GO BP (any GO at any time point with qvalueCutoff = 0.05) we manually selected the GO BP relevant for this study. We retrieved 582 significant BP of interest (Table S1). Given the terms redundancies in GO, we manually grouped the relevant GO BP in meta-category by their term similarity. For each group of related terms, we asked the GO BP term with the lowest corrected p-value to represent the meta-category. For the dot-plots (Figure 1F-G and S2A-B), we selected 8 relevant meta-category and used dotPlot function (clusterProfiler).

###### GO BP in cell population

For each cell population, GO BP enrichment was made for each time point using the pop-active genes. Using compareCluster function we compared the lists of all days simultaneously. Results has been filtered using the BP of interest emerged from whole tissue. Along with the list of interesting BPs, we borrowed from whole muscle analysis also the grouping of the terms in meta-category. As for whole muscle analysis, we asked the GO BP term with the lowest corrected p-value to represent the group. Plot were produced using dotPlot function (clusterProfiler).

Trend lines plot for the meta category for the different cell population (as well as WT vs CCR2KO profiles) were created using the average expression of the genes with loadings different from 0 in the PC1. PCS were computed using the Sparse PCA implementation from TimeClip R package (version 0.3.0) on the expression matrix of normalized counts obtained by averaging time replicates.

##### Gene Ontology biological process on ligand receptors

To better understand the processes driven by the ligand-receptors putative interactions, for each cell, we mapped GO BP on autocrine and paracrine networks. As previously described in whole muscle and cell population analysis, we filtered the results using the total muscle list of interesting BPs.

##### Protein activity inference

To infer protein activity of the receptors, we applied Viper (Viper R package) analysis to our data. Viper assumes that the expression of transcriptional targets of a protein (i.e. its regulon) can be a reporter of the protein activity itself.

Following an approach previously described (Alvarez et al., 2016), we created the regulatory networks from the cell population dataset (both WT and CCR2KO) and the total wild type tissue. Both datasets were normalized separately as describe earlier (see Filtering and Normalization Step).

We set up a candidate regulator list. A regulator that share two categories was assigned following this priority: ligands, receptors (Rezza et al., 2016), transcription factors (TF; GO:0003700, GO:0004677, GO:0030528, GO:0004677), transcriptional cofactors (COF; GO:0003712), and signaling pathway related genes (SP: GO:0007165, GO:0005622, GO:0005886).

To create the regulatory networks from the regulator list, we run ARACNE with 100 bootstrap and mutual information (MI) p-values threshold 10-8. Cell and total tissue regulatory networks were merged and regulators with a regulon size smaller than 25 co-expressed genes were excluded from the analysis. Viper function were used to create the protein activity matrix of the regulators and the cell population dataset.

##### TimeClip analysis of protein activity matrix

Time clip analysis (timeClip R package) was performed using the protein activity matrix. For each cell, day replicates were averaged (Martini et al., 2014). TimeClip was performed on the subset of Reactome pathways (from Graphite R package (Sales et al., 2019; Yu et al., 2012)) that contain the gene *Kdr*.

##### Mitotic Index

Mitotic Index was computed based on the work of Dmitrijeva and colleagues (Dmitrijeva et al., 2018) as the average mRNA expression of 9 genes previously described (Yang et al., 2016) (mouse orthologs were manually annotated). Days were grouped into broader categories as following: “steady” for day 0; “early” from 1 to 3 days after NTX; “middle” from 4 to 7 days after NTX; and “late” from 10 to 14 days after NTX.

## STATISTICAL ANALYSIS

All data is represented as mean ± standard error of the mean (SEM) and the sample number is indicated in the figure legends, where n indicates the number of animals used per group. Data in all figures were obtained from at least 3 independent experiments involving different mice. The significance of differences was assessed with the GraphPad Prism 6 software (GraphPad Software). The normal distribution of all data sets was tested and, depending on the results, multiple comparisons were performed with the parametric one- or two-way analysis of variance (ANOVA) or with the nonparametric Kruskal–Wallis test, while single comparisons were analyzed with the nonparametric Mann–Whitney test or the parametric one-tailed t-test. Results with p values of less than 0.05 were considered statistically significant.

## RESOURCE AVAILABILITY

The datasets supporting the current study are available in GEO with accession ID GSE210748.

## SUPPLEMENTAL INFORMATION

Supplemental information includes seven figures and three tables.

## ACKNOWLEDGEMENTS

This work was supported by CIHR Grant FDN-159908 TO FMVR. We thank the BRC-seq, UBC FACS, AbLab, BRC genotyping and transgenic units, David Guo and Lucas Rempel for technical assistance, as well as Simone Morandi for the graphical abstract.

## AUTHOR CONTRIBUTIONS

All authors were responsible for performing experiments and analysis. EG, PM, ND, MT, and FMVR were involved in experimental design, data interpretation and preparation of the manuscript. EG, PM, ND, MT, MH, CR, and FMVR were involved in editing the manuscript.

## REFERENCES

Abou-Khalil, R., Le Grand, F., Pallafacchina, G., Valable, S., Authier, F.J., Rudnicki, M.A., Gherardi, R.K., Germain, S., Chretien, F., Sotiropoulos, A., et al. (2009). Autocrine and Paracrine Angiopoietin 1/Tie-2 Signaling Promotes Muscle Satellite Cell Self-Renewal. Cell Stem Cell 5, 298–309.

Alvarez, M.J., Shen, Y., Giorgi, F.M., Lachmann, A., Ding, B.B., Hilda Ye, B., and Califano, A. (2016). Functional characterization of somatic mutations in cancer using network-based inference of protein activity. Nat. Genet. 48, 838–847.

Arnold, L., Henry, A., Poron, F., Baba-Amer, Y., van Rooijen, N., Plonquet, A., Gherardi, R.K., and Chazaud, B. (2007). Inflammatory monocytes recruited after skeletal muscle injury switch into antiinflammatory macrophages to support myogenesis. J. Exp. Med. 204, 1057–1069.

Babbs, A., Chatzopoulou, M., Edwards, B., Squire, S.E., Wilkinson, I.V.L., Wynne, G.M., Russell, A.J., and Davies, K.E. (2020). From diagnosis to therapy in Duchenne muscular dystrophy. Biochem. Soc. Trans. 48, 813–821.

Brock, G., Pihur, V., Datta, S., and Datta, S. (2008). ClValid: An R package for cluster validation. J. Stat. Softw. 25, 1–22.

Carosio, S., Berardinelli, M.G., Aucello, M., and Musarò, A. (2011). Impact of ageing on muscle cell regeneration. Ageing Res. Rev. 10, 35–42.

Charrad, M., Ghazzali, N., Boiteau, V., and Niknafs, A. (2014). Nbclust: An R package for determining the relevant number of clusters in a data set. J. Stat. Softw. 61, 1–36.

Chen, W., Wang, Y.X., Ritso, M., Perkins, T.J., and Rudnicki, M.A. KDR Signaling in Muscle Stem Cells Promotes Asymmetric Division and Progenitor Generation for Efficient Regeneration.

Chiristov, C., Chrétien, F., Abou-Khalil, R., Bassez, G., Vallet, G., Authier, F.J., Bassaglia, Y., Shinin, V., Tajbakhsh, S., Chazaud, B., et al. (2007). Muscle satellite cells and endothelial cells: Close neighbors and privileged partners. Mol. Biol. Cell 18, 1397–1409.

Choi, H., Sheng, J., Gao, D., Li, F., Durrans, A., Ryu, S., Lee, S.B., Narula, N., Rafii, S., Elemento, O., et al. (2015). Transcriptome Analysis of Individual Stromal Cell Populations Identifies Stroma-Tumor Crosstalk in Mouse Lung Cancer Model. Cell Rep. 10, 1187–1201.

Dammone, G., Karaz, S., Lukjanenko, L., Winkler, C., Sizzano, F., Jacot, G., Migliavacca, E., Palini, A., Desvergne, B., Gilardi, F., et al. (2018). PPARγ Controls Ectopic Adipogenesis and Cross-Talks with Myogenesis During Skeletal Muscle Regeneration. Int. J. Mol. Sci. Artic.

Delavar, H., Nogueira, L., Wagner, P.D., Hogan, M.C., Metzger, D., and Breen, E.C. (2014). Skeletal myofiber VEGF is essential for the exercise training response in adult mice. Am J Physiol Regul Integr Comp Physiol 306, 586–595.

Dmitrijeva, M., Ossowski, S., Serrano, L., and Schaefer, M.H. (2018). Tissue-specific DNA methylation loss during ageing and carcinogenesis is linked to chromosome structure, replication timing and cell division rates. Nucleic Acids Res. 46, 7022–7039.

Efremova, M., Vento-Tormo, M., Teichmann, S.A., and Vento-Tormo, R. (2020). CellPhoneDB: inferring cell–cell communication from combined expression of multi-subunit ligand–receptor complexes. Nat. Protoc.

Fiore, D., Judson, R.N., Low, M., Lee, S., Zhang, E., Hopkins, C., Xu, P., Lenzi, A., Rossi, F.M.V., and Lemos, D.R. (2016). Pharmacological blockage of fibro/adipogenic progenitor expansion and suppression of regenerative fibrogenesis is associated with impaired skeletal muscle regeneration. Stem Cell Res. 17, 161–169.

Gaë tan Juban, A., Saclier, M., Yacoub-Youssef, H., Gondin, J., and mi Mounier, R. (2018). AMPK Activation Regulates LTBP4-Dependent TGF-β1 Secretion by Pro-inflammatory Macrophages and Controls Fibrosis in Duchenne Muscular Dystrophy. Cell Rep. 25.

Germani, A., Di Carlo, A., Mangoni, A., Straino, S., Giacinti, C., Turrini, P., Biglioli, P., and Capogrossi, M.C. (2003). Vascular endothelial growth factor modulates skeletal myoblast function. Am. J. Pathol. 163, 1417–1428.

Getz, M.C., Nirody, J.A., and Rangamani, P. (2018). Stability analysis in spatial modeling of cell signaling. Wiley Interdiscip. Rev. Syst. Biol. Med. 10, e1395.

Groppa, E., Brkic, S., Bovo, E., Reginato, S., Sacchi, V., Maggio, N. Di, Muraro, M.G., Calabrese, D., Heberer, M., Gianni-Barrera, R., et al. VEGF dose regulates vascular stabilization through Semaphorin3A and the Neuropilin-1 + monocyte/TGF-b1 paracrine axis.

Joe, A.W.B., Yi, L., Natarajan, A., Le Grand, F., So, L., Wang, J., Rudnicki, M.A., and Rossi, F.M. V. (2010). Muscle injury activates resident fibro/adipogenic progenitors that facilitate myogenesis. Nat. Cell Biol. 12, 153–163.

Knapp, A.E., Goldberg, D., Delavar, H., Trisko, B.M., Tang, K., Hogan, M.C., Wagner, P.D., and Breen, E.C. (2016). Skeletal myofiber VEGF regulates contraction-induced perfusion and exercise capacity but not muscle capillarity in adult mice. Am. J. Physiol. Integr. Comp. Physiol. 311, R192–R199.

Kostallari, E., Baba-Amer, Y., Alonso-Martin, S., Ngoh, P., Relaix, F., Lafuste, P., and Gherardi, R.K. (2015). Pericytes in the myovascular niche promote post-natal myofiber growth and satellite cell quiescence.

Kumar, M.P., Du, J., Lagoudas, G., Jiao, Y., Sawyer, A., Drummond, D.C., Lauffenburger, D.A., and Raue, A. (2018). Analysis of Single-Cell RNA-Seq Identifies Cell-Cell Communication Associated with Tumor Characteristics. Cell Rep. 25, 1458–1468.e4.

Latroche, C., Le Weiss-Gayet, M., Muller, L., Gitiaux, C., Leblanc, P., Liot, S., Ben-Larbi, S., Abou-Khalil, R., Verger, N., Bardot, P., et al. (2017). Stem Cell Reports Ar ticle Coupling between Myogenesis and Angiogenesis during Skeletal Muscle Regeneration Is Stimulated by Restorative Macrophages. Stem Cell Reports 9, 2018–2033.

Lemos, D.R., Babaeijandaghi, F., Low, M., Chang, C.-K., Lee, S.T., Fiore, D., Zhang, R.-H., Natarajan, A., Nedospasov, S.A., and Rossi, F.M. V (2015). Nilotinib reduces muscle fibrosis in chronic muscle injury by promoting TNF-mediated apoptosis of fibro/adipogenic progenitors. Nat. Med. 21, 786–794.

Martini, P., Sales, G., Calura, E., Cagnin, S., Chiogna, M., and Romualdi, C. (2014). TimeClip: Pathway analysis for time course data without replicates. BMC Bioinformatics 15, S3.

Mettouchi, A. (2012). Cell Adhesion & Migration The role of extracellular matrix in vascular branching morphogenesis The role of extracellular matrix in vascular branching morphogenesis. De Micheli, A.J., Laurilliard, E.J., Heinke, C.L., Ravichandran, H., Fraczek, P., Soueid-Baumgarten, S., De Vlaminck, I., Elemento, O., and Cosgrove, B.D. (2020). Single-Cell Analysis of the Muscle Stem Cell Hierarchy Identifies Heterotypic Communication Signals Involved in Skeletal Muscle Regeneration. Cell Rep. 30, 3583–3595.e5.

Mojumdar, K., Liang, F., Giordano, C., Lemaire, C., Danialou, G., Okazaki, T., Bourdon, J., Rafei, M., Galipeau, J., Divangahi, M., et al. (2014). Inflammatory monocytes promote progression of Duchenne muscular dystrophy and can be therapeutically targeted via CCR 2 . EMBO Mol. Med. 6, 1476–1492.

Mukund, K., and Subramaniam, | Shankar (2020). Skeletal muscle: A review of molecular structure and function, in health and disease Health and Disease Models of Systems Properties and Processes > Cellular Models. WIREs Syst Biol Med 12.

Müssel, C., Hopfensitz, M., and Kestler, H.A. (2010). BoolNet-an R package for generation, reconstruction and analysis of Boolean networks. Bioinformatics 26, 1378–1380.

Ochoa, O., Sun, D., Reyes-Reyna, S.M., Waite, L.L., Michalek, J.E., McManus, L.M., and Shireman, P.K. (2007). Delayed angiogenesis and VEGF production in CCR2-/- mice during impaired skeletal muscle regeneration. Am. J. Physiol. - Regul. Integr. Comp. Physiol. 293.

Padgett, M.E., McCord, T.J., McClung, J.M., and Kontos, C.D. (2016). Methods for acute and subacute murine hindlimb ischemia. J. Vis. Exp. 2016, e54166.

Palla, A.R., Ravichandran, M., Wang, Y.X., Alexandrova, L., Yang, A. V., Kraft, P., Holbrook, C.A., Schürch, C.M., Ho, A.T.V., and Blau, H.M. (2021). Inhibition of prostaglandin-degrading enzyme 15-PGDH rejuvenates aged muscle mass and strength. Science (80-.). 371.

Plant, D.R., Colarossi, F.E., and Lynch, G.S. (2006). Notexin causes greater myotoxic damage and slower functional repair in mouse skeletal muscles than bupivacaine. Muscle and Nerve 34, 577–585.

Rezza, A., Wang, Z., Sennett, R., Qiao, W., Wang, D., Heitman, N., Mok, K.W., Clavel, C., Yi, R., Zandstra, P., et al. (2016). Signaling Networks among Stem Cell Precursors, Transit-Amplifying Progenitors, and their Niche in Developing Hair Follicles. Cell Rep. 14, 3001–3018.

Rhoads, R.P., Flann, K.L., Cardinal, T.R., Rathbone, C.R., Liu, X., and Allen, R.E. (2013). Satellite cells isolated from aged or dystrophic muscle exhibit a reduced capacity to promote angiogenesis in vitro. Biochem. Biophys. Res. Commun. 440, 399–404.

Risso, D., Ngai, J., Speed, T.P., and Dudoit, S. (2014). Normalization of RNA-seq data using factor analysis of control genes or samples. Nat. Biotechnol. 32, 896–902.

Robinson, M.D., Mccarthy, D.J., and Smyth, G.K. (2010). edgeR: a Bioconductor package for differential expression analysis of digital gene expression data. Bioinforma. Appl. NOTE 26, 139–140.

Ruhrberg, C., Gerhardt, H., Golding, M., Watson, R., Ioannidou, S., Fujisawa, H., Betsholtz, C., and Shima, D.T. (2002). Spatially restricted patterning cues provided by heparin-binding VEGF-A control blood vessel branching morphogenesis. Genes Dev. 16, 2684–2698.

Saclier, M., Yacoub-Youssef, H., Mackey, A.L., Arnold, L., Ardjoune, H., Magnan, M., Sailhan, F., Chelly, J., Pavlath, G.K., Mounier, R., et al. (2013). Differentially activated macrophages orchestrate myogenic precursor cell fate during human skeletal muscle regeneration. Stem Cells 31, 384–396.

Sales, G., Calura, E., and Romualdi, C. (2019). Meta Graphite-a new layer of pathway annotation to get metabolite networks. Bioinformatics 35, 1258–1260.

Santini, M.P., Malide, D., Hoffman, G., Pandey, G., D’Escamard, V., Nomura-Kitabayashi, A., Rovira, I., Kataoka, H., Ochando, J., Harvey, R.P., et al. (2020). Tissue-Resident PDGFRα + Progenitor Cells Contribute to Fibrosis versus Healing in a Context- and Spatiotemporally Dependent Manner. Cell Rep. 30, 555–570.e7.

Scott, R.W., Arostegui, M., Schweitzer, R., Rossi, F.M.V., and Underhill, T.M. (2019). Hic1 Defines Quiescent Mesenchymal Progenitor Subpopulations with Distinct Functions and Fates in Skeletal Muscle Regeneration. Cell Stem Cell 25, 797–813.e9.

Sobral-Reyes, M.F., and Lemos, D.R. (2019). Recapitulating human tissue damage, repair and fibrosis with hPSC-derived organoids: Concise review. Stem Cells 38, 318–329.

Tee, M.K., and Jaffe, R.B. (2001). A precursor form of vascular endothelial growth factor arises by initiation from an upstream in-frame CUG codon. Biochem. J. 359, 219–226.

Theret, M., Gsaier, L., Schaffer, B., Juban, G., Ben Larbi, S., Weiss-Gayet, M., Bultot, L., Collodet, C., Foretz, M., Desplanches, D., et al. (2017). ^AMPK^ α1- ^LDH^ pathway regulates muscle stem cell self-renewal by controlling metabolic homeostasis. EMBO J. 36, 1946–1962.

Theret, M., Saclier, M., Messina, G., and Rossi, F.M. V (2022). Macrophages in Skeletal Muscle Dystrophies, An Entangled Partner. J. Neuromuscul. Dis. 9, 1–23.

Uezumi, A., Fukada, S.I., Yamamoto, N., Takeda, S., and Tsuchida, K. (2010). Mesenchymal progenitors distinct from satellite cells contribute to ectopic fat cell formation in skeletal muscle. Nat. Cell Biol. 12, 143–152.

Verma, M., Asakura, Y., Murakonda, B.S.R., Pengo, T., Latroche, C., Chazaud, B., McLoon, L.K., and Asakura, A. (2018). Muscle Satellite Cell Cross-Talk with a Vascular Niche Maintains Quiescence via VEGF and Notch Signaling. Cell Stem Cell 23, 530–543.e9.

Verma, M., Asakura, Y., Wang, X., Zhou, K., Ünverdi, M., Kann, A.P., Krauss, R.S., and Asakura, A. (2021). Endothelial cell signature in muscle stem cells validated by VEGFA-FLT1-AKT1 axis promoting survival of muscle stem cell. BioRxiv 2021.08.28.458037.

Villalta, S.A., Rosenthal, W., Martinez, L., Kaur, A., Sparwasser, T., Tidball, J.G., Margeta, M., Spencer, M.J., and Bluestone, J.A. (2014). Regulatory T cells suppress muscle inflammation and injury in muscular dystrophy. Sci. Transl. Med. 6, 258ra142.

Wang, Y.X., Holbrook, C.A., Hamilton, J.N., Garoussian, J., Afshar, M., Su, S., Schürch, C.M., Lee, M.Y., Goltsev, Y., Kundaje, A., et al. (2022). A single cell spatial temporal atlas of skeletal muscle reveals cellular neighborhoods that orchestrate regeneration and become disrupted in aging. BioRxiv 2022.06.10.494732.

Webster, M.T., Manor, U., Lippincott-Schwartz, J., and Fan, C.M. (2016). Intravital Imaging Reveals Ghost Fibers as Architectural Units Guiding Myogenic Progenitors during Regeneration. Cell Stem Cell 18, 243–252.

Willenborg, S., Lucas, T., Van Loo, G., Knipper, J.A., Krieg, T., Haase, I., Brachvogel, B., Hammerschmidt, M., Nagy, A., Ferrara, N., et al. (2012). CCR2 recruits an inflammatory macrophage subpopulation critical for angiogenesis in tissue repair. Blood 120, 613–625.

Wosczyna, M.N., Konishi, C.T., Perez Carbajal, E.E., Wang, T.T., Walsh, R.A., Gan, Q., Wagner, M.W., and Rando, T.A. (2019). Mesenchymal Stromal Cells Are Required for Regeneration and Homeostatic Maintenance of Skeletal Muscle. Cell Rep. 27, 2029–2035.e5.

Yang, Z., Wong, A., Kuh, D., Paul, D.S., Rakyan, V.K., Leslie, R.D., Zheng, S.C., Widschwendter, M., Beck, S., and Teschendorff, A.E. (2016). Correlation of an epigenetic mitotic clock with cancer risk. Genome Biol. 17, 1–18.

Yu, G., Wang, L.G., Han, Y., and He, Q.Y. (2012). ClusterProfiler: An R package for comparing biological themes among gene clusters. Omi. A J. Integr. Biol. 16, 284–287.

Zhang, J., Muri, J., Fitzgerald, G., Gorski, T., Gianni-Barrera, R., Masschelein, E., D’Hulst, G., Gilardoni, P., Turiel, G., Fan, Z., et al. (2020). Endothelial Lactate Controls Muscle Regeneration from Ischemia by Inducing M2-like Macrophage Polarization. Cell Metab. 31, 1136–1153.e7.

Zhou, J.X., Taramelli, R., Pedrini, E., Knijnenburg, T., and Huang, S. (2017). Extracting Intercellular Signaling Network of Cancer Tissues using Ligand-Receptor Expression Patterns from Whole-tumor and Single-cell Transcriptomes. Sci. Rep. 7.

