## Supplementary material for "Spatial compartmentalization of signalling imparts source-specific functions on secreted factors": Supplemental Information_text.docx

**Figure S1 (Related to Figure 1): Distinct cellular activation in normal and delayed regeneration. A)** Schematic overview of the process used to identify genes modulated during the response to damage. Selected genes were either dynamically expressed in a specific cell type, changed in abundance because of the expansion of a specific cell subset or constitutively expressed by only one cell type. **B-C)** Principal Component Analysis (PCA) of FAP, EC, MP, IC, and PER from WT and CCR2KO mice at different time points after NTX injury shows minimal overlap between the transcriptomes of sorted subset. **D)** Silhouette and biological homogeneity index analysis for total WT muscles collected before and at different time points after myotoxin (NTX) damage. **E-F)** Representative images of the vasculature from WT and CCR2KO muscles at different time points after NTX injury. **G)** Vessel density over time and vessel diameter quantification at day 7 post damage (n=3, data represent the mean ± SEM, Kruskal-Wallis test with multiple comparison, ^*^p<0.05). **H)** Representative images of vascular leakage and **I)** pericyte coverage at different time points after NTX damage. Mice were intravenously injected with dextran-Texas Red 10 minutes before tissue harvesting.

**Figure S2 (Related to Figure 1):** **Kinetics of biological processes during skeletal muscle regeneration. A)** Trend line for the 8 main Gene Ontology (GO) categories carried out by FAP, EC, MP, IC, and PER during NTX-induced skeletal muscle regeneration in WT mice. **B)** GO enrichment analysis for the samples in A); dot color reflects adjusted p-value and dot size the ratio between category associated DEGs and total number of genes in the category. **C)** Mitotic index calculated on DEG of FAP, EC, and MP, and proliferation of FAP, EC, and MP from muscles of WT and CCR2KO mice after NTX injury. Twenty-four hours before tissue harvesting, mice were treated with EdU (n=3-10, data represent the mean ± SEM, Kruskal-Wallis test with multiple comparison, ^***^p<0.001). **D)** Representative images of skeletal muscles stained with perilipin (Per) and collagen (Col) 7 days after NTX damage.

**Figure S3 (Related to Figure 2): Redundancy of ligands and receptors in regenerating skeletal muscle after acute injury. A)** Bar plot displaying the main GO categories associated to ligand (Lig) - receptor (Rec) interactions emerged from our interactome analysis. We defined these interactions as autocrine when a cell type expresses both cognate ligand and receptor (in yellow), paracrine when a cell type expresses one but not the other (in light green), and “shared” when a cell type engages in autocrine or paracrine signaling but the ligand or receptor is also expressed by other cells (in dark green). **B)** Schematic representation of the interactome associated to *angiogenesis* GO term in undamaged skeletal muscles.

**Figure S4 (Related to Figure 2 and 3): A) Receptor activity in regenerating skeletal muscle after acute injury.** Activity of receptors used as gold standard markers of the different cell types. The activation was calculated for each time point using regulatory network analysis based on the expression of known receptor-targets. **B) Tissue remodeling after acute non-ischemic injury.** H&E of skeletal muscles collected from WT, MP^VEGFAKO^, and FAP^VEGFAKO^ at 3 and 10 days after NTX.

**Figure S5 (Related to Figure 4):** **VEGFA depletion in MP impairs their proliferation, but does not affect the endothelium. A)** Proliferation assay performed on EC and MP from skeletal muscles from WT and FAP^VEGFAKO^ at 3 days after NTX. **B)** Staining of MP undergoing myogenic differentiation by MyoD in muscles collected from WT and MP^VEGFAKO^ at 3 days after NTX. **C)** Staining of MP undergoing myogenic differentiation by Myogenin in muscles collected from WT and MP^VEGFAKO^ at 7 days after NTX. **D)** On the left, representative images of the TUNEL and MyoD staining. On the right, quantification of TUNEL and MyoD positive cells from skeletal muscle collected at 3 days after NTX from wild type and MP^VEGFAKO^ animals (n=6-8, data represent the mean ± SEM, unpaired t test, ^*^p< 0.05). **E)** Expression of VEGFA receptors in different cell subsets. **F)** Histological images representing *in vitro* expansion of MP sorted from WT and MP^VEGFAKO^ mice at 7 days after cell seeding.

**Figure S6 (Related to Figure 5):** **Differences in vascular response between myotoxin and ischemic damage. A)** Extent of skeletal muscle damage identified by detecting IgGs within the myofibers 5 days after myotoxin and ischemic injuries. **B)** Representative image of vasculature (CD31), basement membrane (laminin), and nuclei (DAPI) in muscle sections collected after NTX and femoral ligation injuries. **C)** Heatmap representing hypoxia and VEGFA related gene expression from RNA-bulk sequencing of skeletal muscles collected at day 1 after myotoxin and ischemic injuries. **D)** HIF-1 signaling pathway (KEGG) with hypoxia and VEGFA related genes highlighted in blue. **E)** Quantification of VEGFA in whole muscles collected 3 days after myotoxin and ischemic injuries. **F)** Proliferation of EC from skeletal muscles of WT mice 4 days after myotoxin and ischemic injuries.

**Figure S7 (Related to Figure 5, Figure 6, and Figure 7):** **A) Tissue remodelling after acute ischemic injury.** H&E of skeletal muscles collected from WT, MP^VEGFAKO^, and FAP^VEGFAKO^ at 5 and 7 days after Fem Lig. **B) VEGFA depletion in FAP does not impair vascular perfusion.** FACS analysis of EC (CD45-/CD31+) from WT and FAP^VEGFAKO^ mice, which were perfused with fluorescently labeled lectin before tissue harvesting and digestion. CD31 staining identifies all ECs, while lectin staining points to ECs in perfused vessels. **C) VEGFA depletion in FAP does not impair FAP proliferation.** Proliferation assay performed on FAP from undamaged skeletal muscles of WT and FAP^VEGFAKO^ mice.

**Table S3 (Related to Figure 1A-D and Figure S1D):** **RUV analysis.** **Cell**, the cell/tissue analyzed. **Type**, the condition either wild-type, wt, knock-out, ko. **RuvSeq K**, the RUVSeq parameter k used.

| **Cell** | **Type** | **RuvSeq K** |
| --- | --- | --- |
| ec | wt | 2 |
| fap | wt | 1 |
| mp | wt | 1 |
| ic | wt | 1 |
| per | wt | 1 |
| ec | ko | 3 |
| fap | ko | 1 |
| mp | ko | 1 |
| Total muscle | wt | 1 |
