## Supplementary material for "Spatial compartmentalization of signalling imparts source-specific functions on secreted factors": TableS1_GO_metacategory.docx

**Table S1 (Related to Figure 1E-G and S2A-B):**

**GO BP Description.** Gene Ontology Biological Process Description grouped in Meta-categories.

| **GO BP Description** | **Meta-category** |
| --- | --- |
| actin crosslink formation | cytoskeleton organization |
| actin cytoskeleton organization | cytoskeleton organization |
| actin cytoskeleton reorganization | cytoskeleton organization |
| actin filament bundle assembly | cytoskeleton organization |
| actin filament bundle organization | cytoskeleton organization |
| actin filament organization | cytoskeleton organization |
| actin filament polymerization | cytoskeleton organization |
| actin filament-based movement | cytoskeleton organization |
| actin filament-based process | cytoskeleton organization |
| actin polymerization or depolymerization | cytoskeleton organization |
| actin-mediated cell contraction | cytoskeleton organization |
| activated T cell proliferation | immune response |
| activation of cysteine-type endopeptidase activity involved in apoptotic process | apoptosis |
| activation of immune response | immune response |
| activation of innate immune response | immune response |
| actomyosin structure organization | cytoskeleton organization |
| acute inflammatory response | immune response |
| acute-phase response | immune response |
| adaptive immune response | immune response |
| adaptive immune response based on somatic recombination of immune receptors built from immunoglobulin superfamily domains | immune response |
| alpha-beta T cell activation | immune response |
| alpha-beta T cell activation involved in immune response | immune response |
| alpha-beta T cell differentiation | immune response |
| alpha-beta T cell differentiation involved in immune response | immune response |
| alpha-beta T cell proliferation | immune response |
| ameboidal-type cell migration | migration |
| angiogenesis | angiogenesis |
| angiogenesis involved in wound healing | angiogenesis |
| antigen processing and presentation | immune response |
| antigen processing and presentation of exogenous antigen | immune response |
| antigen processing and presentation of exogenous peptide antigen | immune response |
| antigen processing and presentation of exogenous peptide antigen via MHC class II | immune response |
| antigen processing and presentation of peptide antigen | immune response |
| antigen processing and presentation of peptide antigen via MHC class I | immune response |
| antigen processing and presentation of peptide antigen via MHC class II | immune response |
| antigen processing and presentation of peptide or polysaccharide antigen via MHC class II | immune response |
| antigen receptor-mediated signaling pathway | immune response |
| apoptotic cell clearance | apoptosis |
| apoptotic signaling pathway | apoptosis |
| B cell activation | immune response |
| B cell activation involved in immune response | immune response |
| B cell apoptotic process | immune response |
| B cell differentiation | immune response |
| B cell homeostasis | immune response |
| B cell mediated immunity | immune response |
| B cell proliferation | immune response |
| B cell receptor signaling pathway | immune response |
| blood circulation | blood circulation |
| blood coagulation | blood coagulation |
| blood vessel development | angiogenesis |
| blood vessel endothelial cell migration | angiogenesis |
| blood vessel morphogenesis | angiogenesis |
| brown fat cell differentiation | brown fat cell differentiation |
| calcium ion homeostasis | calcium ion regulation |
| calcium ion transmembrane import into cytosol | calcium ion regulation |
| calcium ion transmembrane transport | calcium ion regulation |
| calcium ion transport | calcium ion regulation |
| calcium ion transport into cytosol | calcium ion regulation |
| calcium-mediated signaling | calcium ion regulation |
| CD4-positive, alpha-beta T cell activation | immune response |
| CD4-positive, alpha-beta T cell differentiation | immune response |
| CD4-positive, alpha-beta T cell differentiation involved in immune response | immune response |
| CD8-positive, alpha-beta T cell activation | immune response |
| cell activation | cell activation |
| cell activation involved in immune response | immune response |
| cell adhesion mediated by integrin | cell adhesion |
| cell chemotaxis | chemotaxis |
| cell killing | apoptosis |
| cell maturation | cell maturation |
| cell-cell adhesion | cell adhesion |
| cell-cell adhesion via plasma-membrane adhesion molecules | cell adhesion |
| cell-matrix adhesion | cell adhesion |
| cell-substrate adhesion | cell adhesion |
| cellular extravasation | chemotaxis |
| cellular response to interferon-beta | cytokine production |
| cellular response to interferon-gamma | cytokine production |
| cellular response to interleukin-1 | cytokine production |
| cellular response to interleukin-4 | cytokine production |
| cellular response to tumor necrosis factor | cytokine production |
| cellular response to type I interferon | cytokine production |
| chemokine production | chemokine production |
| chemokine-mediated signaling pathway | chemotaxis |
| chemotaxis | chemotaxis |
| circulatory system process | blood circulation |
| coagulation | blood coagulation |
| connective tissue development | ECM remodeling |
| cytokine production | cytokine production |
| cytokine production involved in immune response | cytokine production |
| cytokine secretion | cytokine production |
| cytokine-mediated signaling pathway | cytokine production |
| cytokinesis | cytokine production |
| cytoskeleton-dependent cytokinesis | cytokine production |
| dendritic cell differentiation | immune response |
| developmental growth involved in morphogenesis | developmental growth |
| developmental maturation | cell maturation |
| endocrine process | endocrine process |
| endocytosis | endocytosis |
| endothelial cell proliferation | angiogenesis |
| eosinophil chemotaxis | immune response |
| eosinophil migration | immune response |
| epithelial cell apoptotic process | epithelial cell regulation |
| epithelial cell proliferation | epithelial cell regulation |
| ERK1 and ERK2 cascade | ERK1 and ERK2 cascade |
| erythrocyte homeostasis | erythrocyte regulation |
| extracellular matrix organization | ECM remodeling |
| extracellular structure organization | ECM remodeling |
| extrinsic apoptotic signaling pathway | apoptosis |
| extrinsic apoptotic signaling pathway via death domain receptors | apoptosis |
| Fc receptor mediated stimulatory signaling pathway | Fc receptor mediated stimulatory signaling pathway |
| Fc receptor signaling pathway | Fc receptor mediated stimulatory signaling pathway |
| hematopoietic progenitor cell differentiation | hematopoiesis |
| hemostasis | homeostasis |
| heterophilic cell-cell adhesion via plasma membrane cell adhesion molecules | cell adhesion |
| heterotypic cell-cell adhesion | cell adhesion |
| homotypic cell-cell adhesion | cell adhesion |
| humoral immune response | immune response |
| humoral immune response mediated by circulating immunoglobulin | immune response |
| I-kappaB kinase/NF-kappaB signaling | immune response |
| immune effector process | immune response |
| immune response-activating cell surface receptor signaling pathway | immune response |
| immune response-activating signal transduction | immune response |
| immune response-regulating cell surface receptor signaling pathway | immune response |
| immune response-regulating signaling pathway | immune response |
| immunoglobulin mediated immune response | immune response |
| immunoglobulin production | immune response |
| inflammatory cell apoptotic process | apoptosis |
| inflammatory response | immune response |
| inflammatory response to antigenic stimulus | immune response |
| innate immune response | immune response |
| innate immune response-activating signal transduction | immune response |
| integrin-mediated signaling pathway | cell adhesion |
| interferon-gamma production | cytokine production |
| interleukin-1 beta production | cytokine production |
| interleukin-1 beta secretion | cytokine production |
| interleukin-1 production | cytokine production |
| interleukin-1 secretion | cytokine production |
| interleukin-10 production | cytokine production |
| interleukin-12 production | cytokine production |
| interleukin-2 production | cytokine production |
| interleukin-4 production | cytokine production |
| interleukin-6 production | cytokine production |
| interleukin-6 secretion | cytokine production |
| interleukin-8 production | cytokine production |
| interleukin-8 secretion | cytokine production |
| intrinsic apoptotic signaling pathway | apoptosis |
| intrinsic apoptotic signaling pathway in response to DNA damage | apoptosis |
| JAK-STAT cascade | JAK-STAT cascade |
| leukocyte activation | immune response |
| leukocyte activation involved in immune response | immune response |
| leukocyte adhesion to vascular endothelial cell | immune response |
| leukocyte aggregation | immune response |
| leukocyte apoptotic process | immune response |
| leukocyte cell-cell adhesion | immune response |
| leukocyte chemotaxis | immune response |
| leukocyte degranulation | immune response |
| leukocyte differentiation | immune response |
| leukocyte homeostasis | immune response |
| leukocyte mediated cytotoxicity | immune response |
| leukocyte mediated immunity | immune response |
| leukocyte migration | immune response |
| leukocyte migration involved in inflammatory response | immune response |
| leukocyte proliferation | immune response |
| leukocyte tethering or rolling | immune response |
| lymph node development | lymph node development |
| lymphocyte activation | immune response |
| lymphocyte activation involved in immune response | immune response |
| lymphocyte apoptotic process | immune response |
| lymphocyte chemotaxis | immune response |
| lymphocyte differentiation | immune response |
| lymphocyte homeostasis | immune response |
| lymphocyte mediated immunity | immune response |
| lymphocyte migration | immune response |
| lymphocyte proliferation | immune response |
| macrophage activation | immune response |
| macrophage chemotaxis | immune response |
| macrophage derived foam cell differentiation | immune response |
| macrophage differentiation | immune response |
| macrophage migration | immune response |
| MAPK cascade | MAPK cascade |
| mast cell activation | immune response |
| mast cell activation involved in immune response | immune response |
| mast cell degranulation | immune response |
| mast cell mediated immunity | immune response |
| mature B cell differentiation | immune response |
| mature B cell differentiation involved in immune response | immune response |
| multicellular organismal homeostasis | homeostasis |
| muscle adaptation | muscle tissue morphogenesis |
| muscle cell development | muscle tissue morphogenesis |
| muscle cell differentiation | muscle tissue morphogenesis |
| muscle cell migration | muscle tissue morphogenesis |
| muscle contraction | muscle tissue morphogenesis |
| muscle organ development | muscle tissue morphogenesis |
| muscle structure development | muscle tissue morphogenesis |
| muscle system process | muscle tissue morphogenesis |
| muscle tissue development | muscle tissue morphogenesis |
| muscle tissue morphogenesis | muscle tissue morphogenesis |
| musculoskeletal movement | muscle tissue morphogenesis |
| MyD88-dependent toll-like receptor signaling pathway | MyD88-dependent toll-like receptor |
| myoblast differentiation | muscle tissue morphogenesis |
| myoblast fusion | muscle tissue morphogenesis |
| myofibril assembly | muscle tissue morphogenesis |
| myotube differentiation | muscle tissue morphogenesis |
| natural killer cell activation | immune response |
| natural killer cell mediated cytotoxicity | immune response |
| natural killer cell mediated immunity | immune response |
| negative regulation of alpha-beta T cell activation | immune response |
| negative regulation of angiogenesis | angiogenesis |
| negative regulation of antigen receptor-mediated signaling pathway | immune response |
| negative regulation of apoptotic signaling pathway | apoptosis |
| negative regulation of B cell activation | immune response |
| negative regulation of B cell proliferation | immune response |
| negative regulation of blood coagulation | blood coagulation |
| negative regulation of blood pressure | blood pressure |
| negative regulation of blood vessel morphogenesis | angiogenesis |
| negative regulation of cell adhesion | cell adhesion |
| negative regulation of cell development | cell differentiation |
| negative regulation of cell differentiation | cell differentiation |
| negative regulation of cell killing | apoptosis |
| negative regulation of cell migration | chemotaxis |
| negative regulation of cell motility | chemotaxis |
| negative regulation of cell proliferation | proliferation |
| negative regulation of cell-cell adhesion | cell adhesion |
| negative regulation of cytokine production | cytokine production |
| negative regulation of cytokine secretion | cytokine production |
| negative regulation of developmental growth | developmental growth |
| negative regulation of extrinsic apoptotic signaling pathway | apoptosis |
| negative regulation of extrinsic apoptotic signaling pathway via death domain receptors | apoptosis |
| negative regulation of hemopoiesis | hematopoiesis |
| negative regulation of hemostasis | homeostasis |
| negative regulation of immune effector process | immune response |
| negative regulation of immune response | immune response |
| negative regulation of immune system process | immune response |
| negative regulation of inflammatory response | immune response |
| negative regulation of innate immune response | immune response |
| negative regulation of interleukin-1 beta production | cytokine production |
| negative regulation of interleukin-1 production | cytokine production |
| negative regulation of interleukin-12 production | cytokine production |
| negative regulation of interleukin-2 production | cytokine production |
| negative regulation of interleukin-6 production | cytokine production |
| negative regulation of leukocyte activation | immune response |
| negative regulation of leukocyte apoptotic process | immune response |
| negative regulation of leukocyte cell-cell adhesion | immune response |
| negative regulation of leukocyte differentiation | immune response |
| negative regulation of leukocyte mediated cytotoxicity | immune response |
| negative regulation of leukocyte mediated immunity | immune response |
| negative regulation of leukocyte proliferation | immune response |
| negative regulation of lymphocyte mediated immunity | immune response |
| negative regulation of lymphocyte proliferation | immune response |
| negative regulation of myoblast differentiation | muscle tissue morphogenesis |
| negative regulation of natural killer cell mediated cytotoxicity | immune response |
| negative regulation of natural killer cell mediated immunity | immune response |
| negative regulation of response to wounding | wounding |
| negative regulation of T cell activation | immune response |
| negative regulation of T cell proliferation | immune response |
| negative regulation of tissue remodeling | growth |
| negative regulation of tumor necrosis factor production | immune response |
| negative regulation of tumor necrosis factor superfamily cytokine production | immune response |
| negative regulation of wound healing | wounding |
| neutrophil activation | immune response |
| neutrophil chemotaxis | immune response |
| neutrophil mediated immunity | immune response |
| neutrophil migration | immune response |
| non-canonical Wnt signaling pathway | non-canonical Wnt signaling pathway |
| phagocytosis | phagocytosis |
| phagocytosis, engulfment | phagocytosis |
| platelet activation | blood coagulation |
| platelet aggregation | blood coagulation |
| positive regulation of acute inflammatory response | immune response |
| positive regulation of adaptive immune response | immune response |
| positive regulation of adaptive immune response based on somatic recombination of immune receptors built from immunoglobulin superfamily domains | immune response |
| positive regulation of alpha-beta T cell activation | immune response |
| positive regulation of alpha-beta T cell differentiation | immune response |
| positive regulation of angiogenesis | angiogenesis |
| positive regulation of antigen receptor-mediated signaling pathway | immune response |
| positive regulation of apoptotic process | apoptosis |
| positive regulation of B cell activation | immune response |
| positive regulation of B cell differentiation | immune response |
| positive regulation of B cell mediated immunity | immune response |
| positive regulation of B cell proliferation | immune response |
| positive regulation of blood coagulation | blood coagulation |
| positive regulation of blood vessel diameter | angiogenesis |
| positive regulation of blood vessel endothelial cell migration | angiogenesis |
| positive regulation of CD4-positive, alpha-beta T cell activation | immune response |
| positive regulation of CD4-positive, alpha-beta T cell differentiation | immune response |
| positive regulation of cell adhesion | cell adhesion |
| positive regulation of cell adhesion mediated by integrin | cell adhesion |
| positive regulation of cell death | apoptosis |
| positive regulation of cell development | growth |
| positive regulation of cell killing | apoptosis |
| positive regulation of cell migration | chemotaxis |
| positive regulation of cell motility | chemotaxis |
| positive regulation of cell-cell adhesion | cell adhesion |
| positive regulation of cell-matrix adhesion | cell adhesion |
| positive regulation of cell-substrate adhesion | cell adhesion |
| positive regulation of cellular extravasation | chemotaxis |
| positive regulation of chemokine production | chemokine production |
| positive regulation of chemotaxis | chemokine production |
| positive regulation of coagulation | blood coagulation |
| positive regulation of cytokine production | cytokine production |
| positive regulation of cytokine production involved in immune response | cytokine production |
| positive regulation of cytokine secretion | cytokine production |
| positive regulation of cytokine-mediated signaling pathway | cytokine production |
| positive regulation of endocytosis | endocytosis |
| positive regulation of endothelial cell migration | angiogenesis |
| positive regulation of endothelial cell proliferation | angiogenesis |
| positive regulation of epithelial cell migration | epithelial cell regulation |
| positive regulation of epithelial cell proliferation | epithelial cell regulation |
| positive regulation of epithelial to mesenchymal transition | epithelial cell regulation |
| positive regulation of ERK1 and ERK2 cascade | ERK1 and ERK2 cascade |
| positive regulation of erythrocyte differentiation | erythrocyte regulation |
| positive regulation of extrinsic apoptotic signaling pathway | apoptosis |
| positive regulation of fibroblast proliferation | fibroblast regulation |
| positive regulation of I-kappaB kinase/NF-kappaB signaling | immune response |
| positive regulation of immune effector process | immune response |
| positive regulation of immune response | immune response |
| positive regulation of immune system process | immune response |
| positive regulation of immunoglobulin mediated immune response | immune response |
| positive regulation of inflammatory response | immune response |
| positive regulation of innate immune response | immune response |
| positive regulation of interferon-alpha production | cytokine production |
| positive regulation of interferon-gamma production | cytokine production |
| positive regulation of interleukin-1 beta production | cytokine production |
| positive regulation of interleukin-1 beta secretion | cytokine production |
| positive regulation of interleukin-1 production | cytokine production |
| positive regulation of interleukin-1 secretion | cytokine production |
| positive regulation of interleukin-10 production | cytokine production |
| positive regulation of interleukin-2 production | cytokine production |
| positive regulation of interleukin-6 production | cytokine production |
| positive regulation of interleukin-6 secretion | cytokine production |
| positive regulation of interleukin-8 production | cytokine production |
| positive regulation of JAK-STAT cascade | JAK-STAT cascade |
| positive regulation of JNK cascade | JAK-STAT cascade |
| positive regulation of leukocyte activation | immune response |
| positive regulation of leukocyte apoptotic process | immune response |
| positive regulation of leukocyte cell-cell adhesion | immune response |
| positive regulation of leukocyte chemotaxis | immune response |
| positive regulation of leukocyte degranulation | immune response |
| positive regulation of leukocyte differentiation | immune response |
| positive regulation of leukocyte mediated cytotoxicity | immune response |
| positive regulation of leukocyte mediated immunity | immune response |
| positive regulation of leukocyte migration | immune response |
| positive regulation of leukocyte proliferation | immune response |
| positive regulation of lymphocyte activation | immune response |
| positive regulation of lymphocyte apoptotic process | immune response |
| positive regulation of lymphocyte differentiation | immune response |
| positive regulation of lymphocyte mediated immunity | immune response |
| positive regulation of lymphocyte migration | immune response |
| positive regulation of lymphocyte proliferation | immune response |
| positive regulation of macrophage activation | immune response |
| positive regulation of macrophage chemotaxis | immune response |
| positive regulation of macrophage differentiation | immune response |
| positive regulation of macrophage migration | immune response |
| positive regulation of MAPK cascade | MAPK cascade |
| positive regulation of mast cell activation | immune response |
| positive regulation of mast cell activation involved in immune response | immune response |
| positive regulation of mast cell degranulation | immune response |
| positive regulation of myoblast differentiation | muscle tissue morphogenesis |
| positive regulation of myoblast fusion | muscle tissue morphogenesis |
| positive regulation of myotube differentiation | muscle tissue morphogenesis |
| positive regulation of neutrophil chemotaxis | immune response |
| positive regulation of neutrophil migration | immune response |
| positive regulation of NF-kappaB transcription factor activity | NFKB signaling |
| positive regulation of production of molecular mediator of immune response | immune response |
| positive regulation of programmed cell death | apoptosis |
| positive regulation of response to cytokine stimulus | cytokine production |
| positive regulation of response to wounding | wounding |
| positive regulation of smooth muscle cell migration | smooth muscle cell regulation |
| positive regulation of smooth muscle cell proliferation | smooth muscle cell regulation |
| positive regulation of STAT cascade | STAT cascade |
| positive regulation of striated muscle cell differentiation | muscle tissue morphogenesis |
| positive regulation of T cell activation | immune response |
| positive regulation of T cell differentiation | immune response |
| positive regulation of T cell mediated immunity | immune response |
| positive regulation of T cell migration | immune response |
| positive regulation of T cell proliferation | immune response |
| positive regulation of toll-like receptor signaling pathway | toll-like receptor signaling pathway |
| positive regulation of tumor necrosis factor production | cytokine production |
| positive regulation of tumor necrosis factor superfamily cytokine production | cytokine production |
| positive regulation of type I interferon production | cytokine production |
| positive regulation of vasculature development | angiogenesis |
| positive regulation of wound healing | wounding |
| positive T cell selection | immune response |
| production of molecular mediator involved in inflammatory response | immune response |
| production of molecular mediator of immune response | immune response |
| regulation of actin cytoskeleton organization | cytoskeleton organization |
| regulation of actin cytoskeleton reorganization | cytoskeleton organization |
| regulation of actin filament length | cytoskeleton organization |
| regulation of actin filament organization | cytoskeleton organization |
| regulation of actin filament polymerization | cytoskeleton organization |
| regulation of actin filament-based movement | cytoskeleton organization |
| regulation of actin filament-based process | cytoskeleton organization |
| regulation of actin polymerization or depolymerization | cytoskeleton organization |
| regulation of activated T cell proliferation | immune response |
| regulation of acute inflammatory response | immune response |
| regulation of adaptive immune response | immune response |
| regulation of adaptive immune response based on somatic recombination of immune receptors built from immunoglobulin superfamily domains | immune response |
| regulation of alpha-beta T cell activation | immune response |
| regulation of alpha-beta T cell differentiation | immune response |
| regulation of alpha-beta T cell proliferation | immune response |
| regulation of angiogenesis | angiogenesis |
| regulation of antigen processing and presentation | immune response |
| regulation of antigen receptor-mediated signaling pathway | immune response |
| regulation of apoptotic signaling pathway | apoptosis |
| regulation of B cell activation | immune response |
| regulation of B cell apoptotic process | immune response |
| regulation of B cell differentiation | immune response |
| regulation of B cell mediated immunity | immune response |
| regulation of B cell proliferation | immune response |
| regulation of B cell receptor signaling pathway | immune response |
| regulation of blood circulation | blood circulation |
| regulation of blood coagulation | blood coagulation |
| regulation of blood pressure | blood pressure |
| regulation of CD4-positive, alpha-beta T cell activation | immune response |
| regulation of CD4-positive, alpha-beta T cell differentiation | immune response |
| regulation of cell activation | cell activation |
| regulation of cell adhesion | cell adhesion |
| regulation of cell adhesion mediated by integrin | cell adhesion |
| regulation of cell division | proliferation |
| regulation of cell killing | apoptosis |
| regulation of cell maturation | cell maturation |
| regulation of cell morphogenesis | cell morphogenesis |
| regulation of cell-cell adhesion | cell adhesion |
| regulation of cell-matrix adhesion | cell adhesion |
| regulation of cell-substrate adhesion | cell adhesion |
| regulation of cellular extravasation | chemotaxis |
| regulation of chemokine production | chemokine production |
| regulation of chemotaxis | chemotaxis |
| regulation of coagulation | blood coagulation |
| regulation of cytokine production | cytokine production |
| regulation of cytokine production involved in immune response | cytokine production |
| regulation of cytokine secretion | cytokine production |
| regulation of cytokine-mediated signaling pathway | cytokine-mediated signaling pathway |
| regulation of cytoskeleton organization | cytoskeleton organization |
| regulation of developmental growth | growth |
| regulation of endocytosis | endocytosis |
| regulation of endothelial cell proliferation | angiogenesis |
| regulation of epithelial cell proliferation | epithelial cell regulation |
| regulation of ERK1 and ERK2 cascade | ERK1 and ERK2 cascade |
| regulation of erythrocyte differentiation | erythrocyte regulation |
| regulation of extrinsic apoptotic signaling pathway | apoptosis |
| regulation of extrinsic apoptotic signaling pathway via death domain receptors | apoptosis |
| regulation of fibroblast proliferation | fibroblast regulation |
| regulation of granulocyte chemotaxis | granulocyte regulation |
| regulation of granulocyte differentiation | granulocyte regulation |
| regulation of I-kappaB kinase/NF-kappaB signaling | NFKB signaling |
| regulation of immune effector process | immune response |
| regulation of immune response | immune response |
| regulation of immunoglobulin mediated immune response | immune response |
| regulation of immunoglobulin production | immune response |
| regulation of inflammatory response | immune response |
| regulation of inflammatory response to antigenic stimulus | immune response |
| regulation of innate immune response | immune response |
| regulation of interferon-alpha production | cytokine production |
| regulation of interferon-gamma production | cytokine production |
| regulation of interleukin-1 beta production | cytokine production |
| regulation of interleukin-1 beta secretion | cytokine production |
| regulation of interleukin-1 production | cytokine production |
| regulation of interleukin-1 secretion | cytokine production |
| regulation of interleukin-10 production | cytokine production |
| regulation of interleukin-12 production | cytokine production |
| regulation of interleukin-17 production | cytokine production |
| regulation of interleukin-2 production | cytokine production |
| regulation of interleukin-4 production | cytokine production |
| regulation of interleukin-6 production | cytokine production |
| regulation of interleukin-8 production | cytokine production |
| regulation of interleukin-8 secretion | cytokine production |
| regulation of JAK-STAT cascade | JAK-STAT cascade |
| regulation of leukocyte activation | immune response |
| regulation of leukocyte apoptotic process | immune response |
| regulation of leukocyte cell-cell adhesion | immune response |
| regulation of leukocyte chemotaxis | immune response |
| regulation of leukocyte degranulation | immune response |
| regulation of leukocyte differentiation | immune response |
| regulation of leukocyte mediated cytotoxicity | immune response |
| regulation of leukocyte mediated immunity | immune response |
| regulation of leukocyte migration | immune response |
| regulation of leukocyte proliferation | immune response |
| regulation of lymphocyte activation | immune response |
| regulation of lymphocyte apoptotic process | immune response |
| regulation of lymphocyte differentiation | immune response |
| regulation of lymphocyte mediated immunity | immune response |
| regulation of lymphocyte migration | immune response |
| regulation of lymphocyte proliferation | immune response |
| regulation of macrophage activation | immune response |
| regulation of macrophage chemotaxis | immune response |
| regulation of macrophage derived foam cell differentiation | immune response |
| regulation of macrophage differentiation | immune response |
| regulation of macrophage migration | immune response |
| regulation of MAPK cascade | MAPK cascade |
| regulation of mast cell activation | immune response |
| regulation of mast cell activation involved in immune response | immune response |
| regulation of mast cell degranulation | immune response |
| regulation of muscle contraction | muscle tissue morphogenesis |
| regulation of muscle system process | muscle tissue morphogenesis |
| regulation of myoblast differentiation | muscle tissue morphogenesis |
| regulation of myoblast fusion | muscle tissue morphogenesis |
| regulation of myotube differentiation | muscle tissue morphogenesis |
| regulation of natural killer cell mediated cytotoxicity | immune response |
| regulation of natural killer cell mediated immunity | immune response |
| regulation of neutrophil chemotaxis | immune response |
| regulation of neutrophil migration | immune response |
| regulation of phagocytosis | phagocytosis |
| regulation of phagocytosis, engulfment | phagocytosis |
| regulation of platelet activation | blood coagulation |
| regulation of regulatory T cell differentiation | immune response |
| regulation of response to cytokine stimulus | cytokine-mediated signaling pathway |
| regulation of response to wounding | wounding |
| regulation of smooth muscle cell migration | smooth muscle cell regulation |
| regulation of smooth muscle cell proliferation | smooth muscle cell regulation |
| regulation of STAT cascade | STAT cascade |
| regulation of striated muscle cell differentiation | muscle tissue morphogenesis |
| regulation of T cell activation | immune response |
| regulation of T cell apoptotic process | immune response |
| regulation of T cell cytokine production | immune response |
| regulation of T cell differentiation | immune response |
| regulation of T cell mediated cytotoxicity | immune response |
| regulation of T cell mediated immunity | immune response |
| regulation of T cell migration | immune response |
| regulation of T cell proliferation | immune response |
| regulation of T cell receptor signaling pathway | immune response |
| regulation of T-helper 1 type immune response | immune response |
| regulation of tissue remodeling | tissue remodelling |
| regulation of toll-like receptor signaling pathway | immune response/proliferation/toll-like receptor signaling pathway |
| regulation of tumor necrosis factor production | cytokine production |
| regulation of tumor necrosis factor superfamily cytokine production | cytokine production |
| regulation of type 2 immune response | immune response |
| regulation of type I interferon production | cytokine production |
| regulation of type I interferon-mediated signaling pathway | cytokine production |
| regulation of tyrosine phosphorylation of STAT protein | STAT cascade |
| regulation of vascular endothelial growth factor production | angiogenesis |
| regulation of vasculature development | angiogenesis |
| regulation of vesicle-mediated transport | regulation of vesicle-mediated transport |
| regulation of wound healing | wounding |
| regulatory T cell differentiation | immune response |
| response to interferon-alpha | cytokine production |
| response to interferon-beta | cytokine production |
| response to interferon-gamma | cytokine production |
| response to interleukin-1 | cytokine production |
| response to tumor necrosis factor | cytokine production |
| response to type I interferon | cytokine production |
| response to wounding | wounding |
| sarcomere organization | muscle tissue morphogenesis |
| skeletal muscle cell differentiation | muscle tissue morphogenesis |
| skeletal muscle contraction | muscle tissue morphogenesis |
| skeletal muscle tissue regeneration | muscle tissue morphogenesis |
| skeletal system development | muscle tissue morphogenesis |
| skeletal system morphogenesis | muscle tissue morphogenesis |
| smooth muscle cell migration | smooth muscle cell regulation |
| smooth muscle cell proliferation | smooth muscle cell regulation |
| STAT cascade | STAT cascade |
| striated muscle adaptation | muscle tissue morphogenesis |
| striated muscle cell development | muscle tissue morphogenesis |
| striated muscle cell differentiation | muscle tissue morphogenesis |
| striated muscle contraction | muscle tissue morphogenesis |
| T cell activation | immune response |
| T cell activation involved in immune response | immune response |
| T cell apoptotic process | immune response |
| T cell chemotaxis | immune response |
| T cell cytokine production | immune response |
| T cell differentiation | immune response |
| T cell differentiation involved in immune response | immune response |
| T cell homeostasis | immune response |
| T cell mediated cytotoxicity | immune response |
| T cell mediated immunity | immune response |
| T cell migration | immune response |
| T cell proliferation | immune response |
| T cell receptor signaling pathway | immune response |
| T cell selection | immune response |
| T-helper 1 type immune response | immune response |
| T-helper cell differentiation | immune response |
| tissue regeneration | tissue remodelling |
| tissue remodeling | tissue remodelling |
| toll-like receptor 2 signaling pathway | toll-like receptor signaling pathway |
| toll-like receptor signaling pathway | toll-like receptor signaling pathway |
| tumor necrosis factor production | cytokine production |
| tumor necrosis factor superfamily cytokine production | cytokine production |
| type 2 immune response | immune response |
| type I interferon production | cytokine production |
| type I interferon signaling pathway | cytokine production |
| tyrosine phosphorylation of STAT protein | STAT cascade |
| vasculature development | angiogenesis |
| wound healing | wounding |
