## Supplementary material for "Spatial compartmentalization of signalling imparts source-specific functions on secreted factors": TableS2_whole_interactome_table.docx

**Table S2 (Related to Figure S3B): Whole interactome table.** **Ligand**, the Ligand name. **Source**, the type of expression analysis used to identify the source. Can be batch, total, constitutive meaning cell-specific expression, putative cell-specific expansion and constitutively expressed from cell (see Figure S1A and Methods). **Lcell**, the cell expression the ligand. **Day**, when the ligand and receptor are expressed. **Receptor**, the Receptor name. **Rcell**, the cell expressing the receptor.

| **Ligand** | **Source** | **Lcell** | **Day** | **Receptor** | **Rcell** |
| --- | --- | --- | --- | --- | --- |
| Bmp2 | batch | fap | d0 | Eng | ec |
| Bmp2 | batch | mp | d0 | Eng | ec |
| Bmp2 | constitutive | per | d0 | Eng | ec |
| Bmp2 | batch | fap | d0 | Bmpr1a | ec |
| Bmp2 | batch | mp | d0 | Bmpr1a | ec |
| Bmp2 | constitutive | per | d0 | Bmpr1a | ec |
| Bmp2 | batch | fap | d0 | Acvr2a | ec |
| Bmp2 | batch | mp | d0 | Acvr2a | ec |
| Bmp2 | constitutive | per | d0 | Acvr2a | ec |
| Bmp2 | batch | fap | d0 | Bmpr2 | ec |
| Bmp2 | batch | mp | d0 | Bmpr2 | ec |
| Bmp2 | constitutive | per | d0 | Bmpr2 | ec |
| Bmp4 | batch | fap | d0 | Bmpr1a | ec |
| Bmp4 | batch | mp | d0 | Bmpr1a | ec |
| Bmp4 | constitutive | ec | d0 | Bmpr1a | ec |
| Bmp4 | constitutive | per | d0 | Bmpr1a | ec |
| Bmp4 | batch | fap | d0 | Bmpr2 | ec |
| Bmp4 | batch | mp | d0 | Bmpr2 | ec |
| Bmp4 | constitutive | ec | d0 | Bmpr2 | ec |
| Bmp4 | constitutive | per | d0 | Bmpr2 | ec |
| Bmp6 | batch | ec | d0 | Bmpr1a | ec |
| Bmp6 | batch | fap | d0 | Bmpr1a | ec |
| Bmp6 | batch | mp | d0 | Bmpr1a | ec |
| Bmp6 | constitutive | per | d0 | Bmpr1a | ec |
| Bmp6 | batch | ec | d0 | Acvr2a | ec |
| Bmp6 | batch | fap | d0 | Acvr2a | ec |
| Bmp6 | batch | mp | d0 | Acvr2a | ec |
| Bmp6 | constitutive | per | d0 | Acvr2a | ec |
| Bmp6 | batch | ec | d0 | Bmpr2 | ec |
| Bmp6 | batch | fap | d0 | Bmpr2 | ec |
| Bmp6 | batch | mp | d0 | Bmpr2 | ec |
| Bmp6 | constitutive | per | d0 | Bmpr2 | ec |
| Bmp7 | batch | fap | d0 | Eng | ec |
| Bmp7 | batch | fap | d0 | Bmpr1a | ec |
| Bmp7 | batch | fap | d0 | Acvr2a | ec |
| Bmp7 | batch | fap | d0 | Bmpr2 | ec |
| Clu | batch | ec | d0 | Vldlr | ec |
| Clu | batch | fap | d0 | Vldlr | ec |
| Clu | batch | ic | d0 | Vldlr | ec |
| Clu | batch | mp | d0 | Vldlr | ec |
| Crlf1 | batch | ec | d0 | Ctf1 | ec |
| Crlf1 | batch | fap | d0 | Ctf1 | ec |
| Crlf1 | batch | mp | d0 | Ctf1 | ec |
| Csf2 | batch | fap | d0 | Csf2rb | ec |
| Ctgf | batch | ec | d0 | Itga5 | ec |
| Ctgf | batch | fap | d0 | Itga5 | ec |
| Ctgf | batch | ic | d0 | Itga5 | ec |
| Ctgf | batch | mp | d0 | Itga5 | ec |
| Ctgf | constitutive | per | d0 | Itga5 | ec |
| Edn1 | batch | fap | d0 | Ednra | ec |
| Edn1 | constitutive | ec | d0 | Ednra | ec |
| Efnb1 | batch | fap | d0 | Ephb2 | ec |
| Efnb1 | batch | ic | d0 | Ephb2 | ec |
| Efnb1 | constitutive | ec | d0 | Ephb2 | ec |
| Efnb1 | constitutive | per | d0 | Ephb2 | ec |
| Fgf1 | batch | fap | d0 | Fgfr3 | ec |
| Fgf1 | constitutive | per | d0 | Fgfr3 | ec |
| Igf1 | batch | ec | d0 | Igfbp3 | ec |
| Igf1 | batch | fap | d0 | Igfbp3 | ec |
| Igf1 | batch | ic | d0 | Igfbp3 | ec |
| Igf1 | batch | ec | d0 | Igfbp7 | ec |
| Igf1 | batch | fap | d0 | Igfbp7 | ec |
| Igf1 | batch | ic | d0 | Igfbp7 | ec |
| Igf1 | batch | ec | d0 | Igfbp2 | ec |
| Igf1 | batch | fap | d0 | Igfbp2 | ec |
| Igf1 | batch | ic | d0 | Igfbp2 | ec |
| Kitl | batch | fap | d0 | Kit | ec |
| Kitl | batch | ic | d0 | Kit | ec |
| Kitl | batch | mp | d0 | Kit | ec |
| Kitl | constitutive | ec | d0 | Kit | ec |
| Kitl | constitutive | per | d0 | Kit | ec |
| Nov | batch | ec | d0 | Notch1 | ec |
| Nov | batch | fap | d0 | Notch1 | ec |
| Nov | batch | per | d0 | Notch1 | ec |
| Nov | batch | ec | d0 | Itga5 | ec |
| Nov | batch | fap | d0 | Itga5 | ec |
| Nov | batch | per | d0 | Itga5 | ec |
| Nov | batch | ec | d0 | Itgb3 | ec |
| Nov | batch | fap | d0 | Itgb3 | ec |
| Nov | batch | per | d0 | Itgb3 | ec |
| Ntf3 | batch | ec | d0 | Ntrk2 | ec |
| Ntf3 | batch | fap | d0 | Ntrk2 | ec |
| Ntf3 | batch | mp | d0 | Ntrk2 | ec |
| Ntf3 | constitutive | per | d0 | Ntrk2 | ec |
| Pgf | batch | fap | d0 | Flt1 | ec |
| Pgf | constitutive | per | d0 | Flt1 | ec |
| Pgf | batch | fap | d0 | Nrp2 | ec |
| Pgf | constitutive | per | d0 | Nrp2 | ec |
| Pgf | batch | fap | d0 | Nrp1 | ec |
| Pgf | constitutive | per | d0 | Nrp1 | ec |
| Sfrp4 | batch | ec | d0 | Fzd2 | ec |
| Sfrp4 | batch | fap | d0 | Fzd2 | ec |
| Sfrp4 | batch | mp | d0 | Fzd2 | ec |
| Vegfa | batch | fap | d0 | Flt1 | ec |
| Vegfa | batch | mp | d0 | Flt1 | ec |
| Vegfa | batch | fap | d0 | Kdr | ec |
| Vegfa | batch | mp | d0 | Kdr | ec |
| Vegfa | batch | fap | d0 | Nrp2 | ec |
| Vegfa | batch | mp | d0 | Nrp2 | ec |
| Vegfa | batch | fap | d0 | Nrp1 | ec |
| Vegfa | batch | mp | d0 | Nrp1 | ec |
| Vegfa | batch | fap | d0 | Ephb2 | ec |
| Vegfa | batch | mp | d0 | Ephb2 | ec |
| Vegfa | batch | fap | d0 | Vtn | ec |
| Vegfa | batch | mp | d0 | Vtn | ec |
| Vegfa | batch | fap | d0 | Itga9 | ec |
| Vegfa | batch | mp | d0 | Itga9 | ec |
| Wnt11 | batch | fap | d0 | Fzd4 | ec |
| Wnt11 | batch | ic | d0 | Fzd4 | ec |
| Wnt2 | batch | fap | d0 | Sfrp1 | ec |
| Dll1 | batch | ec | d0 | Notch1 | ec |
| Fgf9 | batch | ec | d0 | Fgfr3 | ec |
| Igf2 | batch | ec | d0 | Insr | ec |
| Igf2 | batch | ec | d0 | Vtn | ec |
| Il15 | batch | ec | d0 | Il15ra | ec |
| Il15 | batch | ic | d0 | Il15ra | ec |
| Sfrp5 | batch | ec | d0 | Fzd2 | ec |
| Sfrp5 | batch | mp | d0 | Fzd2 | ec |
| Sfrp5 | batch | per | d0 | Fzd2 | ec |
| Tnfsf10 | batch | ec | d0 | Tnfrsf10b | ec |
| Vegfc | batch | ec | d0 | Flt4 | ec |
| Vegfc | batch | mp | d0 | Flt4 | ec |
| Vegfc | batch | ec | d0 | Kdr | ec |
| Vegfc | batch | mp | d0 | Kdr | ec |
| Vegfc | batch | ec | d0 | Nrp2 | ec |
| Vegfc | batch | mp | d0 | Nrp2 | ec |
| Vegfc | batch | ec | d0 | Nrp1 | ec |
| Vegfc | batch | mp | d0 | Nrp1 | ec |
| Vegfc | batch | ec | d0 | Itga9 | ec |
| Vegfc | batch | mp | d0 | Itga9 | ec |
| Areg | batch | mp | d0 | Egfr | ec |
| Clcf1 | batch | ic | d0 | Crlf1 | ec |
| Clcf1 | batch | mp | d0 | Crlf1 | ec |
| Clcf1 | batch | ic | d0 | Cntfr | ec |
| Clcf1 | batch | mp | d0 | Cntfr | ec |
| Ctf1 | batch | mp | d0 | Il6st | ec |
| Edn3 | batch | mp | d0 | Ednra | ec |
| Gdnf | batch | mp | d0 | Ret | ec |
| Gdnf | batch | per | d0 | Ret | ec |
| Gmfb | batch | mp | d0 | Egfr | ec |
| Gmfb | batch | mp | d0 | Itpr3 | ec |
| Hbegf | batch | mp | d0 | Egfr | ec |
| Hbegf | constitutive | ec | d0 | Egfr | ec |
| Hbegf | constitutive | per | d0 | Egfr | ec |
| Inhbb | batch | mp | d0 | Acvr2a | ec |
| Inhbb | constitutive | fap | d0 | Acvr2a | ec |
| Inhbb | constitutive | per | d0 | Acvr2a | ec |
| Jag1 | batch | mp | d0 | Notch1 | ec |
| Jag1 | constitutive | per | d0 | Notch1 | ec |
| Jag2 | batch | mp | d0 | Notch1 | ec |
| Jag2 | constitutive | ec | d0 | Notch1 | ec |
| Nppc | batch | mp | d0 | Npr2 | ec |
| Rabep1 | batch | mp | d0 | Lsr | ec |
| Serpini1 | batch | mp | d0 | Plat | ec |
| Serpini1 | constitutive | per | d0 | Plat | ec |
| Tgfb3 | batch | mp | d0 | Acvrl1 | ec |
| Tnfsf12 | batch | mp | d0 | Tnfrsf25 | ec |
| Tnfsf12 | batch | mp | d0 | Tnfrsf1a | ec |
| Tnfsf12 | batch | mp | d0 | Fas | ec |
| Ifng | batch | ic | d0 | Ifngr1 | ec |
| Il6 | batch | ic | d0 | Il6st | ec |
| Ltb | batch | ic | d0 | Tnfrsf1a | ec |
| Mmp9 | batch | ic | d0 | Flt1 | ec |
| Pdgfb | batch | ic | d0 | Pdgfrb | ec |
| Pdgfb | constitutive | ec | d0 | Pdgfrb | ec |
| Adm | batch | per | d0 | Calcrl | ec |
| Angpt4 | batch | per | d0 | Tek | ec |
| Dll4 | total | ec | d0 | Notch1 | ec |
| Angpt1 | constitutive | fap | d0 | Tek | ec |
| Fgf7 | constitutive | fap | d0 | Fgfr3 | ec |
| Fgf7 | constitutive | fap | d0 | Nrp1 | ec |
| Tgfb2 | constitutive | ec | d0 | Tgfbr3 | ec |
| Tgfb2 | constitutive | fap | d0 | Tgfbr3 | ec |
| Tgfb2 | constitutive | per | d0 | Tgfbr3 | ec |
| Tgfb2 | constitutive | ec | d0 | Vtn | ec |
| Tgfb2 | constitutive | fap | d0 | Vtn | ec |
| Tgfb2 | constitutive | per | d0 | Vtn | ec |
| Bmp5 | constitutive | ec | d0 | Bmpr1a | ec |
| Bmp5 | constitutive | fap | d0 | Bmpr1a | ec |
| Bmp5 | constitutive | per | d0 | Bmpr1a | ec |
| Cmtm8 | constitutive | ec | d0 | Egfr | ec |
| Angpt2 | constitutive | ec | d0 | Tek | ec |
| Angpt2 | constitutive | ic | d0 | Tek | ec |
| Angpt2 | constitutive | per | d0 | Tek | ec |
| Lif | constitutive | mp | d0 | Il6st | ec |
| Lif | constitutive | mp | d0 | Lifr | ec |
| Ebi3 | constitutive | ic | d0 | Il27ra | ec |
| Nampt | constitutive | ic | d0 | Adora2a | ec |
| Sfrp1 | constitutive | per | d0 | Fzd2 | ec |
| Adm | batch | fap | d1 | Calcrl | ec |
| Adm | batch | per | d1 | Calcrl | ec |
| Bmp2 | batch | ec | d1 | Eng | ec |
| Bmp2 | batch | fap | d1 | Eng | ec |
| Bmp2 | batch | mp | d1 | Eng | ec |
| Bmp2 | constitutive | per | d1 | Eng | ec |
| Bmp2 | batch | ec | d1 | Bmpr1a | ec |
| Bmp2 | batch | fap | d1 | Bmpr1a | ec |
| Bmp2 | batch | mp | d1 | Bmpr1a | ec |
| Bmp2 | constitutive | per | d1 | Bmpr1a | ec |
| Bmp2 | batch | ec | d1 | Acvr2a | ec |
| Bmp2 | batch | fap | d1 | Acvr2a | ec |
| Bmp2 | batch | mp | d1 | Acvr2a | ec |
| Bmp2 | constitutive | per | d1 | Acvr2a | ec |
| Bmp2 | batch | ec | d1 | Bmpr2 | ec |
| Bmp2 | batch | fap | d1 | Bmpr2 | ec |
| Bmp2 | batch | mp | d1 | Bmpr2 | ec |
| Bmp2 | constitutive | per | d1 | Bmpr2 | ec |
| Bmp6 | batch | fap | d1 | Bmpr1a | ec |
| Bmp6 | constitutive | per | d1 | Bmpr1a | ec |
| Bmp6 | batch | fap | d1 | Acvr2a | ec |
| Bmp6 | constitutive | per | d1 | Acvr2a | ec |
| Bmp6 | batch | fap | d1 | Bmpr2 | ec |
| Bmp6 | constitutive | per | d1 | Bmpr2 | ec |
| Efnb1 | batch | fap | d1 | Ephb2 | ec |
| Efnb1 | batch | ic | d1 | Ephb2 | ec |
| Efnb1 | constitutive | ec | d1 | Ephb2 | ec |
| Efnb1 | constitutive | per | d1 | Ephb2 | ec |
| Hgf | batch | fap | d1 | Vtn | ec |
| Il6 | batch | ec | d1 | Il6st | ec |
| Il6 | batch | fap | d1 | Il6st | ec |
| Il6 | batch | per | d1 | Il6st | ec |
| Kitl | batch | fap | d1 | Kit | ec |
| Kitl | batch | ic | d1 | Kit | ec |
| Kitl | batch | mp | d1 | Kit | ec |
| Kitl | constitutive | ec | d1 | Kit | ec |
| Kitl | constitutive | per | d1 | Kit | ec |
| Lif | batch | fap | d1 | Il6st | ec |
| Lif | batch | ic | d1 | Il6st | ec |
| Lif | batch | per | d1 | Il6st | ec |
| Lif | constitutive | mp | d1 | Il6st | ec |
| Lif | batch | fap | d1 | Lifr | ec |
| Lif | batch | ic | d1 | Lifr | ec |
| Lif | batch | per | d1 | Lifr | ec |
| Lif | constitutive | mp | d1 | Lifr | ec |
| Pdgfc | batch | fap | d1 | Pdgfra | ec |
| Pdgfc | constitutive | ic | d1 | Pdgfra | ec |
| Pdgfc | constitutive | per | d1 | Pdgfra | ec |
| Pf4 | batch | fap | d1 | Ldlr | ec |
| Pf4 | batch | ic | d1 | Ldlr | ec |
| Pf4 | batch | per | d1 | Ldlr | ec |
| Pf4 | batch | fap | d1 | Thbd | ec |
| Pf4 | batch | ic | d1 | Thbd | ec |
| Pf4 | batch | per | d1 | Thbd | ec |
| Tnfrsf11b | batch | fap | d1 | Vtn | ec |
| Wnt2 | batch | fap | d1 | Sfrp1 | ec |
| Il1b | batch | ec | d1 | Il1r2 | ec |
| Il1b | batch | ic | d1 | Il1r2 | ec |
| Il1b | batch | per | d1 | Il1r2 | ec |
| Il1b | batch | ec | d1 | Adrb2 | ec |
| Il1b | batch | ic | d1 | Adrb2 | ec |
| Il1b | batch | per | d1 | Adrb2 | ec |
| Inhbb | batch | ec | d1 | Acvr2a | ec |
| Inhbb | constitutive | fap | d1 | Acvr2a | ec |
| Inhbb | constitutive | per | d1 | Acvr2a | ec |
| Pgf | batch | ec | d1 | Flt1 | ec |
| Pgf | constitutive | per | d1 | Flt1 | ec |
| Pgf | batch | ec | d1 | Nrp2 | ec |
| Pgf | constitutive | per | d1 | Nrp2 | ec |
| Pgf | batch | ec | d1 | Nrp1 | ec |
| Pgf | constitutive | per | d1 | Nrp1 | ec |
| Areg | batch | ic | d1 | Egfr | ec |
| Areg | batch | mp | d1 | Egfr | ec |
| Bmp4 | batch | mp | d1 | Bmpr1a | ec |
| Bmp4 | constitutive | ec | d1 | Bmpr1a | ec |
| Bmp4 | constitutive | per | d1 | Bmpr1a | ec |
| Bmp4 | batch | mp | d1 | Bmpr2 | ec |
| Bmp4 | constitutive | ec | d1 | Bmpr2 | ec |
| Bmp4 | constitutive | per | d1 | Bmpr2 | ec |
| Clcf1 | batch | mp | d1 | Crlf1 | ec |
| Clu | batch | ic | d1 | Vldlr | ec |
| Clu | batch | mp | d1 | Vldlr | ec |
| Ctgf | batch | ic | d1 | Itga5 | ec |
| Ctgf | batch | mp | d1 | Itga5 | ec |
| Ctgf | constitutive | per | d1 | Itga5 | ec |
| Gdnf | batch | mp | d1 | Ret | ec |
| Gdnf | batch | per | d1 | Ret | ec |
| Gmfb | batch | mp | d1 | Egfr | ec |
| Gmfb | batch | mp | d1 | Itpr3 | ec |
| Hbegf | batch | mp | d1 | Egfr | ec |
| Hbegf | constitutive | ec | d1 | Egfr | ec |
| Hbegf | constitutive | per | d1 | Egfr | ec |
| Il33 | batch | mp | d1 | Il1rl1 | ec |
| Jag2 | batch | mp | d1 | Notch1 | ec |
| Jag2 | constitutive | ec | d1 | Notch1 | ec |
| Nppc | batch | mp | d1 | Npr2 | ec |
| Pdap1 | batch | mp | d1 | Pdgfa | ec |
| Serpini1 | batch | mp | d1 | Plat | ec |
| Serpini1 | constitutive | per | d1 | Plat | ec |
| Sfrp4 | batch | mp | d1 | Fzd2 | ec |
| Sfrp5 | batch | mp | d1 | Fzd2 | ec |
| Spp1 | batch | mp | d1 | Itga5 | ec |
| Spp1 | batch | per | d1 | Itga5 | ec |
| Spp1 | batch | mp | d1 | Vtn | ec |
| Spp1 | batch | per | d1 | Vtn | ec |
| Spp1 | batch | mp | d1 | Itgb3 | ec |
| Spp1 | batch | per | d1 | Itgb3 | ec |
| Spp1 | batch | mp | d1 | Itga9 | ec |
| Spp1 | batch | per | d1 | Itga9 | ec |
| Vegfa | batch | ic | d1 | Flt1 | ec |
| Vegfa | batch | mp | d1 | Flt1 | ec |
| Vegfa | batch | ic | d1 | Kdr | ec |
| Vegfa | batch | mp | d1 | Kdr | ec |
| Vegfa | batch | ic | d1 | Nrp2 | ec |
| Vegfa | batch | mp | d1 | Nrp2 | ec |
| Vegfa | batch | ic | d1 | Nrp1 | ec |
| Vegfa | batch | mp | d1 | Nrp1 | ec |
| Vegfa | batch | ic | d1 | Ephb2 | ec |
| Vegfa | batch | mp | d1 | Ephb2 | ec |
| Vegfa | batch | ic | d1 | Vtn | ec |
| Vegfa | batch | mp | d1 | Vtn | ec |
| Vegfa | batch | ic | d1 | Itga9 | ec |
| Vegfa | batch | mp | d1 | Itga9 | ec |
| Vegfc | batch | mp | d1 | Flt4 | ec |
| Vegfc | batch | mp | d1 | Kdr | ec |
| Vegfc | batch | mp | d1 | Nrp2 | ec |
| Vegfc | batch | mp | d1 | Nrp1 | ec |
| Vegfc | batch | mp | d1 | Itga9 | ec |
| Il1a | batch | ic | d1 | Il1r2 | ec |
| Jag1 | batch | ic | d1 | Notch1 | ec |
| Jag1 | constitutive | per | d1 | Notch1 | ec |
| Osm | batch | ic | d1 | Osmr | ec |
| Osm | batch | per | d1 | Osmr | ec |
| Osm | batch | ic | d1 | Il6st | ec |
| Osm | batch | per | d1 | Il6st | ec |
| Osm | batch | ic | d1 | Lifr | ec |
| Osm | batch | per | d1 | Lifr | ec |
| Tnf | batch | ic | d1 | Tnfrsf1a | ec |
| Tnf | batch | per | d1 | Tnfrsf1a | ec |
| Tnf | batch | ic | d1 | Fas | ec |
| Tnf | batch | per | d1 | Fas | ec |
| Angpt4 | batch | per | d1 | Tek | ec |
| Btc | batch | per | d1 | Egfr | ec |
| Il11 | batch | per | d1 | Il6st | ec |
| Mmp9 | batch | per | d1 | Flt1 | ec |
| Nov | batch | per | d1 | Notch1 | ec |
| Nov | batch | per | d1 | Itga5 | ec |
| Nov | batch | per | d1 | Itgb3 | ec |
| Sfrp2 | batch | per | d1 | Fzd2 | ec |
| Angpt1 | constitutive | fap | d1 | Tek | ec |
| Fgf7 | constitutive | fap | d1 | Nrp1 | ec |
| Tgfb2 | constitutive | ec | d1 | Tgfbr3 | ec |
| Tgfb2 | constitutive | fap | d1 | Tgfbr3 | ec |
| Tgfb2 | constitutive | per | d1 | Tgfbr3 | ec |
| Tgfb2 | constitutive | ec | d1 | Vtn | ec |
| Tgfb2 | constitutive | fap | d1 | Vtn | ec |
| Tgfb2 | constitutive | per | d1 | Vtn | ec |
| Bmp5 | constitutive | ec | d1 | Bmpr1a | ec |
| Bmp5 | constitutive | fap | d1 | Bmpr1a | ec |
| Bmp5 | constitutive | per | d1 | Bmpr1a | ec |
| Pdgfb | constitutive | ec | d1 | Pdgfra | ec |
| Pdgfb | constitutive | ec | d1 | Pdgfrb | ec |
| Cmtm8 | constitutive | ec | d1 | Egfr | ec |
| Edn1 | constitutive | ec | d1 | Ednra | ec |
| Angpt2 | constitutive | ec | d1 | Tek | ec |
| Angpt2 | constitutive | ic | d1 | Tek | ec |
| Angpt2 | constitutive | per | d1 | Tek | ec |
| Pdgfa | constitutive | ec | d1 | Pdgfra | ec |
| Pdgfa | constitutive | mp | d1 | Pdgfra | ec |
| Pdgfa | constitutive | per | d1 | Pdgfra | ec |
| Ebi3 | constitutive | ic | d1 | Il27ra | ec |
| Nampt | constitutive | ic | d1 | Adora2a | ec |
| Sfrp1 | constitutive | per | d1 | Fzd2 | ec |
| Ntf3 | constitutive | per | d1 | Ntrk2 | ec |
| Adm | batch | fap | d2 | Calcrl | ec |
| Adm | batch | per | d2 | Calcrl | ec |
| Apln | batch | ec | d2 | Aplnr | ec |
| Apln | batch | fap | d2 | Aplnr | ec |
| Bmp2 | batch | ec | d2 | Eng | ec |
| Bmp2 | batch | fap | d2 | Eng | ec |
| Bmp2 | batch | mp | d2 | Eng | ec |
| Bmp2 | constitutive | per | d2 | Eng | ec |
| Bmp2 | batch | ec | d2 | Bmpr1a | ec |
| Bmp2 | batch | fap | d2 | Bmpr1a | ec |
| Bmp2 | batch | mp | d2 | Bmpr1a | ec |
| Bmp2 | constitutive | per | d2 | Bmpr1a | ec |
| Bmp2 | batch | ec | d2 | Acvr2a | ec |
| Bmp2 | batch | fap | d2 | Acvr2a | ec |
| Bmp2 | batch | mp | d2 | Acvr2a | ec |
| Bmp2 | constitutive | per | d2 | Acvr2a | ec |
| Bmp2 | batch | ec | d2 | Bmpr2 | ec |
| Bmp2 | batch | fap | d2 | Bmpr2 | ec |
| Bmp2 | batch | mp | d2 | Bmpr2 | ec |
| Bmp2 | constitutive | per | d2 | Bmpr2 | ec |
| Bmp6 | batch | fap | d2 | Bmpr1a | ec |
| Bmp6 | constitutive | per | d2 | Bmpr1a | ec |
| Bmp6 | batch | fap | d2 | Acvr2a | ec |
| Bmp6 | constitutive | per | d2 | Acvr2a | ec |
| Bmp6 | batch | fap | d2 | Bmpr2 | ec |
| Bmp6 | constitutive | per | d2 | Bmpr2 | ec |
| Ccl2 | batch | ec | d2 | Ccr5 | ec |
| Ccl2 | batch | fap | d2 | Ccr5 | ec |
| Ccl2 | batch | ic | d2 | Ccr5 | ec |
| Ccl2 | batch | mp | d2 | Ccr5 | ec |
| Ccl2 | batch | per | d2 | Ccr5 | ec |
| Ccl3 | batch | ec | d2 | Ccr5 | ec |
| Ccl3 | batch | fap | d2 | Ccr5 | ec |
| Ccl3 | batch | ic | d2 | Ccr5 | ec |
| Ccl3 | batch | mp | d2 | Ccr5 | ec |
| Ccl3 | batch | per | d2 | Ccr5 | ec |
| Ccl4 | batch | fap | d2 | Ccr5 | ec |
| Ccl4 | batch | ic | d2 | Ccr5 | ec |
| Ccl4 | batch | mp | d2 | Ccr5 | ec |
| Ccl4 | batch | per | d2 | Ccr5 | ec |
| Ccl5 | batch | ec | d2 | Ccr5 | ec |
| Ccl5 | batch | fap | d2 | Ccr5 | ec |
| Ccl7 | batch | ec | d2 | Ccr5 | ec |
| Ccl7 | batch | fap | d2 | Ccr5 | ec |
| Ccl7 | batch | ic | d2 | Ccr5 | ec |
| Ccl7 | batch | mp | d2 | Ccr5 | ec |
| Ccl7 | batch | per | d2 | Ccr5 | ec |
| Csf1 | batch | fap | d2 | H2-Bl | ec |
| Csf1 | batch | ic | d2 | H2-Bl | ec |
| Csf1 | batch | mp | d2 | H2-Bl | ec |
| Csf2 | batch | fap | d2 | Csf2ra | ec |
| Csf2 | batch | fap | d2 | Csf2rb | ec |
| Csf2 | batch | fap | d2 | Csf3r | ec |
| Csf2 | batch | fap | d2 | Il3ra | ec |
| Efnb1 | batch | fap | d2 | Ephb2 | ec |
| Efnb1 | constitutive | ec | d2 | Ephb2 | ec |
| Efnb1 | constitutive | per | d2 | Ephb2 | ec |
| Hgf | batch | fap | d2 | Vtn | ec |
| Hgf | batch | ic | d2 | Vtn | ec |
| Il1b | batch | ec | d2 | Il1r2 | ec |
| Il1b | batch | fap | d2 | Il1r2 | ec |
| Il1b | batch | ic | d2 | Il1r2 | ec |
| Il1b | batch | mp | d2 | Il1r2 | ec |
| Il1b | batch | per | d2 | Il1r2 | ec |
| Il1b | batch | ec | d2 | Adrb2 | ec |
| Il1b | batch | fap | d2 | Adrb2 | ec |
| Il1b | batch | ic | d2 | Adrb2 | ec |
| Il1b | batch | mp | d2 | Adrb2 | ec |
| Il1b | batch | per | d2 | Adrb2 | ec |
| Il33 | batch | fap | d2 | Il1rl1 | ec |
| Il33 | batch | mp | d2 | Il1rl1 | ec |
| Il6 | batch | ec | d2 | Il6st | ec |
| Il6 | batch | fap | d2 | Il6st | ec |
| Il6 | batch | per | d2 | Il6st | ec |
| Kitl | batch | fap | d2 | Kit | ec |
| Kitl | batch | mp | d2 | Kit | ec |
| Kitl | constitutive | ec | d2 | Kit | ec |
| Kitl | constitutive | per | d2 | Kit | ec |
| Lif | batch | fap | d2 | Il6st | ec |
| Lif | batch | ic | d2 | Il6st | ec |
| Lif | batch | per | d2 | Il6st | ec |
| Lif | constitutive | mp | d2 | Il6st | ec |
| Lif | batch | fap | d2 | Lifr | ec |
| Lif | batch | ic | d2 | Lifr | ec |
| Lif | batch | per | d2 | Lifr | ec |
| Lif | constitutive | mp | d2 | Lifr | ec |
| Mmp9 | batch | fap | d2 | Flt1 | ec |
| Mmp9 | batch | mp | d2 | Flt1 | ec |
| Mmp9 | batch | per | d2 | Flt1 | ec |
| Pdgfc | batch | fap | d2 | Pdgfra | ec |
| Pdgfc | constitutive | ic | d2 | Pdgfra | ec |
| Pdgfc | constitutive | per | d2 | Pdgfra | ec |
| Pf4 | batch | ec | d2 | Ldlr | ec |
| Pf4 | batch | fap | d2 | Ldlr | ec |
| Pf4 | batch | ic | d2 | Ldlr | ec |
| Pf4 | batch | mp | d2 | Ldlr | ec |
| Pf4 | batch | per | d2 | Ldlr | ec |
| Pf4 | batch | ec | d2 | Thbd | ec |
| Pf4 | batch | fap | d2 | Thbd | ec |
| Pf4 | batch | ic | d2 | Thbd | ec |
| Pf4 | batch | mp | d2 | Thbd | ec |
| Pf4 | batch | per | d2 | Thbd | ec |
| Sfrp2 | batch | ec | d2 | Fzd2 | ec |
| Sfrp2 | batch | fap | d2 | Fzd2 | ec |
| Sfrp2 | batch | per | d2 | Fzd2 | ec |
| Spp1 | batch | ec | d2 | Itga5 | ec |
| Spp1 | batch | fap | d2 | Itga5 | ec |
| Spp1 | batch | ic | d2 | Itga5 | ec |
| Spp1 | batch | mp | d2 | Itga5 | ec |
| Spp1 | batch | per | d2 | Itga5 | ec |
| Spp1 | batch | ec | d2 | Vtn | ec |
| Spp1 | batch | fap | d2 | Vtn | ec |
| Spp1 | batch | ic | d2 | Vtn | ec |
| Spp1 | batch | mp | d2 | Vtn | ec |
| Spp1 | batch | per | d2 | Vtn | ec |
| Spp1 | batch | ec | d2 | Itgb3 | ec |
| Spp1 | batch | fap | d2 | Itgb3 | ec |
| Spp1 | batch | ic | d2 | Itgb3 | ec |
| Spp1 | batch | mp | d2 | Itgb3 | ec |
| Spp1 | batch | per | d2 | Itgb3 | ec |
| Spp1 | batch | ec | d2 | Itga9 | ec |
| Spp1 | batch | fap | d2 | Itga9 | ec |
| Spp1 | batch | ic | d2 | Itga9 | ec |
| Spp1 | batch | mp | d2 | Itga9 | ec |
| Spp1 | batch | per | d2 | Itga9 | ec |
| Tnfrsf11b | batch | fap | d2 | Vtn | ec |
| Tslp | batch | fap | d2 | Crlf2 | ec |
| Tslp | constitutive | ec | d2 | Crlf2 | ec |
| Wnt2 | batch | fap | d2 | Sfrp1 | ec |
| Igf1 | batch | ec | d2 | Igfbp7 | ec |
| Igf1 | batch | ic | d2 | Igfbp7 | ec |
| Igf1 | batch | mp | d2 | Igfbp7 | ec |
| Igf1 | batch | ec | d2 | Igfbp2 | ec |
| Igf1 | batch | ic | d2 | Igfbp2 | ec |
| Igf1 | batch | mp | d2 | Igfbp2 | ec |
| Il15 | batch | ec | d2 | Il17ra | ec |
| Il15 | batch | ic | d2 | Il17ra | ec |
| Il15 | batch | ec | d2 | Il2rg | ec |
| Il15 | batch | ic | d2 | Il2rg | ec |
| Il15 | batch | ec | d2 | Il15ra | ec |
| Il15 | batch | ic | d2 | Il15ra | ec |
| Inhbb | batch | ec | d2 | Acvr2a | ec |
| Inhbb | constitutive | fap | d2 | Acvr2a | ec |
| Inhbb | constitutive | per | d2 | Acvr2a | ec |
| Lgals3 | batch | ec | d2 | Lgals3bp | ec |
| Lgals3 | batch | ic | d2 | Lgals3bp | ec |
| Lgals3 | batch | mp | d2 | Lgals3bp | ec |
| Lgals3 | batch | per | d2 | Lgals3bp | ec |
| Mmp12 | batch | ec | d2 | Plaur | ec |
| Mmp12 | batch | ic | d2 | Plaur | ec |
| Mmp12 | batch | mp | d2 | Plaur | ec |
| Mmp12 | batch | per | d2 | Plaur | ec |
| Mmp13 | batch | ec | d2 | F2r | ec |
| Mmp13 | batch | ic | d2 | F2r | ec |
| Mmp13 | batch | per | d2 | F2r | ec |
| Osm | batch | ec | d2 | Osmr | ec |
| Osm | batch | ic | d2 | Osmr | ec |
| Osm | batch | mp | d2 | Osmr | ec |
| Osm | batch | per | d2 | Osmr | ec |
| Osm | batch | ec | d2 | Il6st | ec |
| Osm | batch | ic | d2 | Il6st | ec |
| Osm | batch | mp | d2 | Il6st | ec |
| Osm | batch | per | d2 | Il6st | ec |
| Osm | batch | ec | d2 | Lifr | ec |
| Osm | batch | ic | d2 | Lifr | ec |
| Osm | batch | mp | d2 | Lifr | ec |
| Osm | batch | per | d2 | Lifr | ec |
| Pgf | batch | ec | d2 | Flt1 | ec |
| Pgf | constitutive | per | d2 | Flt1 | ec |
| Pgf | batch | ec | d2 | Nrp2 | ec |
| Pgf | constitutive | per | d2 | Nrp2 | ec |
| Pgf | batch | ec | d2 | Nrp1 | ec |
| Pgf | constitutive | per | d2 | Nrp1 | ec |
| Sfrp1 | batch | ec | d2 | Fzd2 | ec |
| Sfrp1 | constitutive | per | d2 | Fzd2 | ec |
| Tnfsf10 | batch | ec | d2 | Tnfrsf10b | ec |
| Areg | batch | ic | d2 | Egfr | ec |
| Areg | batch | mp | d2 | Egfr | ec |
| Bmp4 | batch | mp | d2 | Bmpr1a | ec |
| Bmp4 | constitutive | ec | d2 | Bmpr1a | ec |
| Bmp4 | constitutive | per | d2 | Bmpr1a | ec |
| Bmp4 | batch | mp | d2 | Bmpr2 | ec |
| Bmp4 | constitutive | ec | d2 | Bmpr2 | ec |
| Bmp4 | constitutive | per | d2 | Bmpr2 | ec |
| C4b | batch | mp | d2 | C3ar1 | ec |
| C4b | constitutive | per | d2 | C3ar1 | ec |
| Ccl11 | batch | mp | d2 | Ccr5 | ec |
| Ccl11 | batch | per | d2 | Ccr5 | ec |
| Ccl8 | batch | mp | d2 | Ccr5 | ec |
| Ccl8 | total | fap | d2 | Ccr5 | ec |
| Ccl8 | total | ic | d2 | Ccr5 | ec |
| Clcf1 | batch | mp | d2 | Crlf1 | ec |
| Clu | batch | mp | d2 | Vldlr | ec |
| Clu | total | per | d2 | Vldlr | ec |
| Ctgf | batch | mp | d2 | Itga5 | ec |
| Ctgf | constitutive | per | d2 | Itga5 | ec |
| Ctsg | batch | mp | d2 | F2rl3 | ec |
| Ebi3 | batch | mp | d2 | Il27ra | ec |
| Ebi3 | constitutive | ic | d2 | Il27ra | ec |
| Gdnf | batch | mp | d2 | Ret | ec |
| Gdnf | batch | per | d2 | Ret | ec |
| Gmfb | batch | mp | d2 | Egfr | ec |
| Gmfb | batch | mp | d2 | Itpr3 | ec |
| Hbegf | batch | mp | d2 | Egfr | ec |
| Hbegf | constitutive | ec | d2 | Egfr | ec |
| Hbegf | constitutive | per | d2 | Egfr | ec |
| Il10 | batch | ic | d2 | Il10ra | ec |
| Il10 | batch | mp | d2 | Il10ra | ec |
| Jag2 | batch | mp | d2 | Notch1 | ec |
| Jag2 | constitutive | ec | d2 | Notch1 | ec |
| Nppc | batch | mp | d2 | Npr2 | ec |
| Pdap1 | batch | mp | d2 | Pdgfa | ec |
| Serpini1 | batch | mp | d2 | Plat | ec |
| Serpini1 | constitutive | per | d2 | Plat | ec |
| Sfrp4 | batch | mp | d2 | Fzd2 | ec |
| Sfrp5 | batch | mp | d2 | Fzd2 | ec |
| Tgfb1 | batch | mp | d2 | Eng | ec |
| Tgfb1 | total | ec | d2 | Eng | ec |
| Tgfb1 | total | ic | d2 | Eng | ec |
| Tgfb1 | batch | mp | d2 | Acvrl1 | ec |
| Tgfb1 | total | ec | d2 | Acvrl1 | ec |
| Tgfb1 | total | ic | d2 | Acvrl1 | ec |
| Tgfb1 | batch | mp | d2 | Tgfbr3 | ec |
| Tgfb1 | total | ec | d2 | Tgfbr3 | ec |
| Tgfb1 | total | ic | d2 | Tgfbr3 | ec |
| Tgfb1 | batch | mp | d2 | Vtn | ec |
| Tgfb1 | total | ec | d2 | Vtn | ec |
| Tgfb1 | total | ic | d2 | Vtn | ec |
| Tnf | batch | ic | d2 | Tnfrsf1a | ec |
| Tnf | batch | mp | d2 | Tnfrsf1a | ec |
| Tnf | batch | per | d2 | Tnfrsf1a | ec |
| Tnf | batch | ic | d2 | Fas | ec |
| Tnf | batch | mp | d2 | Fas | ec |
| Tnf | batch | per | d2 | Fas | ec |
| Vegfa | batch | ic | d2 | Flt1 | ec |
| Vegfa | batch | mp | d2 | Flt1 | ec |
| Vegfa | batch | ic | d2 | Kdr | ec |
| Vegfa | batch | mp | d2 | Kdr | ec |
| Vegfa | batch | ic | d2 | Nrp2 | ec |
| Vegfa | batch | mp | d2 | Nrp2 | ec |
| Vegfa | batch | ic | d2 | Nrp1 | ec |
| Vegfa | batch | mp | d2 | Nrp1 | ec |
| Vegfa | batch | ic | d2 | Ephb2 | ec |
| Vegfa | batch | mp | d2 | Ephb2 | ec |
| Vegfa | batch | ic | d2 | Vtn | ec |
| Vegfa | batch | mp | d2 | Vtn | ec |
| Vegfa | batch | ic | d2 | Itga9 | ec |
| Vegfa | batch | mp | d2 | Itga9 | ec |
| Vegfc | batch | mp | d2 | Flt4 | ec |
| Vegfc | batch | mp | d2 | Kdr | ec |
| Vegfc | batch | mp | d2 | Nrp2 | ec |
| Vegfc | batch | mp | d2 | Nrp1 | ec |
| Vegfc | batch | mp | d2 | Itga9 | ec |
| Il1a | batch | ic | d2 | Il1r2 | ec |
| Jag1 | batch | ic | d2 | Notch1 | ec |
| Jag1 | constitutive | per | d2 | Notch1 | ec |
| Pdgfa | batch | ic | d2 | Pdgfra | ec |
| Pdgfa | constitutive | ec | d2 | Pdgfra | ec |
| Pdgfa | constitutive | mp | d2 | Pdgfra | ec |
| Pdgfa | constitutive | per | d2 | Pdgfra | ec |
| Tnfsf13 | batch | ic | d2 | Tnfrsf11b | ec |
| Tnfsf13 | batch | ic | d2 | Tnfrsf1a | ec |
| Tnfsf13 | batch | ic | d2 | Fas | ec |
| Angpt4 | batch | per | d2 | Tek | ec |
| Btc | batch | per | d2 | Egfr | ec |
| Il11 | batch | per | d2 | Il6st | ec |
| Nov | batch | per | d2 | Notch1 | ec |
| Nov | batch | per | d2 | Itga5 | ec |
| Nov | batch | per | d2 | Itgb3 | ec |
| Tnfsf12 | total | ec | d2 | Tnfrsf11b | ec |
| Tnfsf12 | total | ic | d2 | Tnfrsf11b | ec |
| Tnfsf12 | total | per | d2 | Tnfrsf11b | ec |
| Tnfsf12 | total | ec | d2 | Tnfrsf1a | ec |
| Tnfsf12 | total | ic | d2 | Tnfrsf1a | ec |
| Tnfsf12 | total | per | d2 | Tnfrsf1a | ec |
| Tnfsf12 | total | ec | d2 | Fas | ec |
| Tnfsf12 | total | ic | d2 | Fas | ec |
| Tnfsf12 | total | per | d2 | Fas | ec |
| Angpt1 | constitutive | fap | d2 | Tek | ec |
| Fgf7 | constitutive | fap | d2 | Nrp1 | ec |
| Tgfb2 | constitutive | ec | d2 | Tgfbr3 | ec |
| Tgfb2 | constitutive | fap | d2 | Tgfbr3 | ec |
| Tgfb2 | constitutive | per | d2 | Tgfbr3 | ec |
| Tgfb2 | constitutive | ec | d2 | Vtn | ec |
| Tgfb2 | constitutive | fap | d2 | Vtn | ec |
| Tgfb2 | constitutive | per | d2 | Vtn | ec |
| Bmp5 | constitutive | ec | d2 | Bmpr1a | ec |
| Bmp5 | constitutive | fap | d2 | Bmpr1a | ec |
| Bmp5 | constitutive | per | d2 | Bmpr1a | ec |
| Pdgfb | constitutive | ec | d2 | Pdgfra | ec |
| Pdgfb | constitutive | ec | d2 | Pdgfrb | ec |
| Cmtm8 | constitutive | ec | d2 | Egfr | ec |
| Edn1 | constitutive | ec | d2 | Ednra | ec |
| Angpt2 | constitutive | ec | d2 | Tek | ec |
| Angpt2 | constitutive | ic | d2 | Tek | ec |
| Angpt2 | constitutive | per | d2 | Tek | ec |
| Nampt | constitutive | ic | d2 | Adora2a | ec |
| Ntf3 | constitutive | per | d2 | Ntrk2 | ec |
| Apln | batch | ec | d3 | Aplnr | ec |
| Apln | batch | fap | d3 | Aplnr | ec |
| Bmp2 | batch | fap | d3 | Eng | ec |
| Bmp2 | constitutive | per | d3 | Eng | ec |
| Bmp2 | batch | fap | d3 | Bmpr1a | ec |
| Bmp2 | constitutive | per | d3 | Bmpr1a | ec |
| Bmp2 | batch | fap | d3 | Acvr2a | ec |
| Bmp2 | constitutive | per | d3 | Acvr2a | ec |
| Bmp2 | batch | fap | d3 | Bmpr2 | ec |
| Bmp2 | constitutive | per | d3 | Bmpr2 | ec |
| Bmp6 | batch | fap | d3 | Bmpr1a | ec |
| Bmp6 | constitutive | per | d3 | Bmpr1a | ec |
| Bmp6 | batch | fap | d3 | Acvr2a | ec |
| Bmp6 | constitutive | per | d3 | Acvr2a | ec |
| Bmp6 | batch | fap | d3 | Bmpr2 | ec |
| Bmp6 | constitutive | per | d3 | Bmpr2 | ec |
| C4b | batch | fap | d3 | C3ar1 | ec |
| C4b | constitutive | per | d3 | C3ar1 | ec |
| Ccl3 | batch | ec | d3 | Ccr5 | ec |
| Ccl3 | batch | fap | d3 | Ccr5 | ec |
| Ccl3 | batch | mp | d3 | Ccr5 | ec |
| Ccl3 | batch | per | d3 | Ccr5 | ec |
| Ccl4 | batch | fap | d3 | Ccr5 | ec |
| Ccl4 | batch | mp | d3 | Ccr5 | ec |
| Ccl4 | batch | per | d3 | Ccr5 | ec |
| Ccl5 | batch | ec | d3 | Ccr5 | ec |
| Ccl5 | batch | fap | d3 | Ccr5 | ec |
| Csf2 | batch | fap | d3 | Csf2ra | ec |
| Csf2 | batch | fap | d3 | Csf2rb | ec |
| Csf2 | batch | fap | d3 | Csf3r | ec |
| Csf2 | batch | fap | d3 | Il3ra | ec |
| Ctgf | batch | fap | d3 | Itga5 | ec |
| Ctgf | constitutive | per | d3 | Itga5 | ec |
| Efnb1 | batch | fap | d3 | Ephb2 | ec |
| Efnb1 | constitutive | ec | d3 | Ephb2 | ec |
| Efnb1 | constitutive | per | d3 | Ephb2 | ec |
| Fgf1 | batch | fap | d3 | Fgfr3 | ec |
| Fgf1 | constitutive | per | d3 | Fgfr3 | ec |
| Hbegf | batch | fap | d3 | Egfr | ec |
| Hbegf | constitutive | ec | d3 | Egfr | ec |
| Hbegf | constitutive | per | d3 | Egfr | ec |
| Hgf | batch | fap | d3 | Vtn | ec |
| Hgf | batch | ic | d3 | Vtn | ec |
| Igf1 | batch | ec | d3 | Igfbp5 | ec |
| Igf1 | batch | fap | d3 | Igfbp5 | ec |
| Igf1 | batch | ic | d3 | Igfbp5 | ec |
| Igf1 | batch | mp | d3 | Igfbp5 | ec |
| Igf1 | batch | ec | d3 | Igfbp7 | ec |
| Igf1 | batch | fap | d3 | Igfbp7 | ec |
| Igf1 | batch | ic | d3 | Igfbp7 | ec |
| Igf1 | batch | mp | d3 | Igfbp7 | ec |
| Igf1 | batch | ec | d3 | Igfbp2 | ec |
| Igf1 | batch | fap | d3 | Igfbp2 | ec |
| Igf1 | batch | ic | d3 | Igfbp2 | ec |
| Igf1 | batch | mp | d3 | Igfbp2 | ec |
| Igf2 | batch | fap | d3 | Insr | ec |
| Igf2 | batch | fap | d3 | Vtn | ec |
| Il1b | batch | fap | d3 | Adrb2 | ec |
| Il1b | batch | mp | d3 | Adrb2 | ec |
| Il1b | batch | per | d3 | Adrb2 | ec |
| Il33 | batch | fap | d3 | Il1rl1 | ec |
| Il33 | batch | mp | d3 | Il1rl1 | ec |
| Kitl | batch | fap | d3 | Kit | ec |
| Kitl | constitutive | ec | d3 | Kit | ec |
| Kitl | constitutive | per | d3 | Kit | ec |
| Mmp13 | batch | ec | d3 | F2r | ec |
| Mmp13 | batch | fap | d3 | F2r | ec |
| Mmp13 | batch | ic | d3 | F2r | ec |
| Mmp13 | batch | per | d3 | F2r | ec |
| Mmp9 | batch | fap | d3 | Flt1 | ec |
| Mmp9 | batch | mp | d3 | Flt1 | ec |
| Mmp9 | batch | per | d3 | Flt1 | ec |
| Ntf3 | batch | ec | d3 | Ntrk2 | ec |
| Ntf3 | batch | fap | d3 | Ntrk2 | ec |
| Ntf3 | constitutive | per | d3 | Ntrk2 | ec |
| Sfrp1 | batch | ec | d3 | Fzd2 | ec |
| Sfrp1 | batch | fap | d3 | Fzd2 | ec |
| Sfrp1 | constitutive | per | d3 | Fzd2 | ec |
| Sfrp2 | batch | ec | d3 | Fzd2 | ec |
| Sfrp2 | batch | fap | d3 | Fzd2 | ec |
| Sfrp2 | batch | per | d3 | Fzd2 | ec |
| Sfrp4 | batch | fap | d3 | Fzd2 | ec |
| Spp1 | batch | ec | d3 | Itga5 | ec |
| Spp1 | batch | fap | d3 | Itga5 | ec |
| Spp1 | batch | ic | d3 | Itga5 | ec |
| Spp1 | batch | mp | d3 | Itga5 | ec |
| Spp1 | batch | per | d3 | Itga5 | ec |
| Spp1 | batch | ec | d3 | Vtn | ec |
| Spp1 | batch | fap | d3 | Vtn | ec |
| Spp1 | batch | ic | d3 | Vtn | ec |
| Spp1 | batch | mp | d3 | Vtn | ec |
| Spp1 | batch | per | d3 | Vtn | ec |
| Spp1 | batch | ec | d3 | Itgb3 | ec |
| Spp1 | batch | fap | d3 | Itgb3 | ec |
| Spp1 | batch | ic | d3 | Itgb3 | ec |
| Spp1 | batch | mp | d3 | Itgb3 | ec |
| Spp1 | batch | per | d3 | Itgb3 | ec |
| Spp1 | batch | ec | d3 | Itga9 | ec |
| Spp1 | batch | fap | d3 | Itga9 | ec |
| Spp1 | batch | ic | d3 | Itga9 | ec |
| Spp1 | batch | mp | d3 | Itga9 | ec |
| Spp1 | batch | per | d3 | Itga9 | ec |
| Tgfb3 | batch | fap | d3 | Acvrl1 | ec |
| Tgfb3 | total | ec | d3 | Acvrl1 | ec |
| Tgfb3 | total | per | d3 | Acvrl1 | ec |
| Tnfrsf11b | batch | fap | d3 | Vtn | ec |
| Wnt2 | batch | fap | d3 | Sfrp1 | ec |
| Fgf9 | batch | ec | d3 | Fgfr3 | ec |
| Il15 | batch | ec | d3 | Il17ra | ec |
| Il15 | batch | ic | d3 | Il17ra | ec |
| Il15 | batch | ec | d3 | Il2rg | ec |
| Il15 | batch | ic | d3 | Il2rg | ec |
| Il15 | batch | ec | d3 | Il15ra | ec |
| Il15 | batch | ic | d3 | Il15ra | ec |
| Il6 | batch | ec | d3 | Il6st | ec |
| Il6 | batch | per | d3 | Il6st | ec |
| Inhbb | batch | ec | d3 | Acvr2a | ec |
| Inhbb | constitutive | fap | d3 | Acvr2a | ec |
| Inhbb | constitutive | per | d3 | Acvr2a | ec |
| Lgals3 | batch | ec | d3 | Lgals3bp | ec |
| Lgals3 | batch | ic | d3 | Lgals3bp | ec |
| Lgals3 | batch | mp | d3 | Lgals3bp | ec |
| Lgals3 | batch | per | d3 | Lgals3bp | ec |
| Osm | batch | ec | d3 | Osmr | ec |
| Osm | batch | mp | d3 | Osmr | ec |
| Osm | batch | per | d3 | Osmr | ec |
| Osm | batch | ec | d3 | Il6st | ec |
| Osm | batch | mp | d3 | Il6st | ec |
| Osm | batch | per | d3 | Il6st | ec |
| Osm | batch | ec | d3 | Lifr | ec |
| Osm | batch | mp | d3 | Lifr | ec |
| Osm | batch | per | d3 | Lifr | ec |
| Tnfsf10 | batch | ec | d3 | Tnfrsf10b | ec |
| Vegfc | batch | ec | d3 | Flt4 | ec |
| Vegfc | batch | ec | d3 | Kdr | ec |
| Vegfc | batch | ec | d3 | Nrp2 | ec |
| Vegfc | batch | ec | d3 | Nrp1 | ec |
| Vegfc | batch | ec | d3 | Itga9 | ec |
| Ccl2 | batch | mp | d3 | Ccr5 | ec |
| Ccl2 | batch | per | d3 | Ccr5 | ec |
| Ccl7 | batch | mp | d3 | Ccr5 | ec |
| Ccl7 | batch | per | d3 | Ccr5 | ec |
| Ccl8 | batch | mp | d3 | Ccr5 | ec |
| Ccl8 | total | fap | d3 | Ccr5 | ec |
| Ccl8 | total | ic | d3 | Ccr5 | ec |
| Csf1 | batch | ic | d3 | H2-Bl | ec |
| Csf1 | batch | mp | d3 | H2-Bl | ec |
| Ebi3 | batch | mp | d3 | Il27ra | ec |
| Ebi3 | constitutive | ic | d3 | Il27ra | ec |
| Il10 | batch | ic | d3 | Il10ra | ec |
| Il10 | batch | mp | d3 | Il10ra | ec |
| Pdap1 | batch | mp | d3 | Pdgfa | ec |
| Tgfb1 | batch | mp | d3 | Eng | ec |
| Tgfb1 | total | ec | d3 | Eng | ec |
| Tgfb1 | total | ic | d3 | Eng | ec |
| Tgfb1 | batch | mp | d3 | Acvrl1 | ec |
| Tgfb1 | total | ec | d3 | Acvrl1 | ec |
| Tgfb1 | total | ic | d3 | Acvrl1 | ec |
| Tgfb1 | batch | mp | d3 | Tgfbr3 | ec |
| Tgfb1 | total | ec | d3 | Tgfbr3 | ec |
| Tgfb1 | total | ic | d3 | Tgfbr3 | ec |
| Tgfb1 | batch | mp | d3 | Vtn | ec |
| Tgfb1 | total | ec | d3 | Vtn | ec |
| Tgfb1 | total | ic | d3 | Vtn | ec |
| Tnf | batch | mp | d3 | Tnfrsf1a | ec |
| Tnf | batch | per | d3 | Tnfrsf1a | ec |
| Tnf | batch | mp | d3 | Fas | ec |
| Tnf | batch | per | d3 | Fas | ec |
| Tnfsf13 | batch | ic | d3 | Tnfrsf11b | ec |
| Tnfsf13 | batch | ic | d3 | Tnfrsf25 | ec |
| Tnfsf13 | batch | ic | d3 | Tnfrsf1a | ec |
| Tnfsf13 | batch | ic | d3 | Fas | ec |
| Btc | batch | per | d3 | Erbb2 | ec |
| Btc | batch | per | d3 | Egfr | ec |
| Il11 | batch | per | d3 | Il6st | ec |
| Lif | batch | per | d3 | Il6st | ec |
| Lif | constitutive | mp | d3 | Il6st | ec |
| Lif | batch | per | d3 | Lifr | ec |
| Lif | constitutive | mp | d3 | Lifr | ec |
| Dll1 | total | mp | d3 | Notch1 | ec |
| Angpt1 | constitutive | fap | d3 | Tek | ec |
| Fgf7 | constitutive | fap | d3 | Fgfr3 | ec |
| Fgf7 | constitutive | fap | d3 | Nrp1 | ec |
| Tgfb2 | constitutive | ec | d3 | Tgfbr3 | ec |
| Tgfb2 | constitutive | fap | d3 | Tgfbr3 | ec |
| Tgfb2 | constitutive | per | d3 | Tgfbr3 | ec |
| Tgfb2 | constitutive | ec | d3 | Vtn | ec |
| Tgfb2 | constitutive | fap | d3 | Vtn | ec |
| Tgfb2 | constitutive | per | d3 | Vtn | ec |
| Bmp5 | constitutive | ec | d3 | Bmpr1a | ec |
| Bmp5 | constitutive | fap | d3 | Bmpr1a | ec |
| Bmp5 | constitutive | per | d3 | Bmpr1a | ec |
| Jag2 | constitutive | ec | d3 | Notch1 | ec |
| Pdgfb | constitutive | ec | d3 | Pdgfrb | ec |
| Cmtm8 | constitutive | ec | d3 | Egfr | ec |
| Edn1 | constitutive | ec | d3 | Ednra | ec |
| Angpt2 | constitutive | ec | d3 | Tek | ec |
| Angpt2 | constitutive | ic | d3 | Tek | ec |
| Angpt2 | constitutive | per | d3 | Tek | ec |
| Bmp4 | constitutive | ec | d3 | Bmpr1a | ec |
| Bmp4 | constitutive | per | d3 | Bmpr1a | ec |
| Bmp4 | constitutive | ec | d3 | Bmpr2 | ec |
| Bmp4 | constitutive | per | d3 | Bmpr2 | ec |
| Nampt | constitutive | ic | d3 | Adora2a | ec |
| Serpini1 | constitutive | per | d3 | Plat | ec |
| Pgf | constitutive | per | d3 | Flt1 | ec |
| Pgf | constitutive | per | d3 | Nrp2 | ec |
| Pgf | constitutive | per | d3 | Nrp1 | ec |
| Jag1 | constitutive | per | d3 | Notch1 | ec |
| Apln | batch | fap | d4 | Aplnr | ec |
| C4b | batch | fap | d4 | C3ar1 | ec |
| C4b | batch | ic | d4 | C3ar1 | ec |
| C4b | constitutive | per | d4 | C3ar1 | ec |
| Ccl3 | batch | ec | d4 | Ccr5 | ec |
| Ccl3 | batch | fap | d4 | Ccr5 | ec |
| Ccl3 | batch | mp | d4 | Ccr5 | ec |
| Ccl4 | batch | fap | d4 | Ccr5 | ec |
| Ccl4 | batch | mp | d4 | Ccr5 | ec |
| Ccl5 | batch | ec | d4 | Ccr5 | ec |
| Ccl5 | batch | fap | d4 | Ccr5 | ec |
| Ctgf | batch | fap | d4 | Itga5 | ec |
| Ctgf | constitutive | per | d4 | Itga5 | ec |
| Efnb1 | batch | fap | d4 | Ephb2 | ec |
| Efnb1 | batch | mp | d4 | Ephb2 | ec |
| Efnb1 | constitutive | ec | d4 | Ephb2 | ec |
| Efnb1 | constitutive | per | d4 | Ephb2 | ec |
| Fgf1 | batch | fap | d4 | Fgfr3 | ec |
| Fgf1 | constitutive | per | d4 | Fgfr3 | ec |
| Hbegf | batch | fap | d4 | Egfr | ec |
| Hbegf | constitutive | ec | d4 | Egfr | ec |
| Hbegf | constitutive | per | d4 | Egfr | ec |
| Hgf | batch | fap | d4 | Vtn | ec |
| Hgf | batch | ic | d4 | Vtn | ec |
| Igf1 | batch | ec | d4 | Igfbp5 | ec |
| Igf1 | batch | fap | d4 | Igfbp5 | ec |
| Igf1 | batch | ic | d4 | Igfbp5 | ec |
| Igf1 | batch | ec | d4 | Igfbp7 | ec |
| Igf1 | batch | fap | d4 | Igfbp7 | ec |
| Igf1 | batch | ic | d4 | Igfbp7 | ec |
| Igf1 | batch | ec | d4 | Igfbp2 | ec |
| Igf1 | batch | fap | d4 | Igfbp2 | ec |
| Igf1 | batch | ic | d4 | Igfbp2 | ec |
| Igf2 | batch | ec | d4 | Insr | ec |
| Igf2 | batch | fap | d4 | Insr | ec |
| Igf2 | batch | mp | d4 | Insr | ec |
| Igf2 | batch | ec | d4 | Vtn | ec |
| Igf2 | batch | fap | d4 | Vtn | ec |
| Igf2 | batch | mp | d4 | Vtn | ec |
| Il1b | batch | fap | d4 | Adrb2 | ec |
| Il33 | batch | fap | d4 | Il1rl1 | ec |
| Il33 | batch | mp | d4 | Il1rl1 | ec |
| Kitl | batch | fap | d4 | Kit | ec |
| Kitl | batch | mp | d4 | Kit | ec |
| Kitl | constitutive | ec | d4 | Kit | ec |
| Kitl | constitutive | per | d4 | Kit | ec |
| Mmp13 | batch | fap | d4 | F2r | ec |
| Mmp13 | batch | ic | d4 | F2r | ec |
| Mmp13 | batch | mp | d4 | F2r | ec |
| Mmp9 | batch | fap | d4 | Flt1 | ec |
| Mmp9 | batch | mp | d4 | Flt1 | ec |
| Ntf3 | batch | ec | d4 | Ntrk2 | ec |
| Ntf3 | batch | fap | d4 | Ntrk2 | ec |
| Ntf3 | constitutive | per | d4 | Ntrk2 | ec |
| Sfrp1 | batch | ec | d4 | Fzd2 | ec |
| Sfrp1 | batch | fap | d4 | Fzd2 | ec |
| Sfrp1 | constitutive | per | d4 | Fzd2 | ec |
| Sfrp2 | batch | ec | d4 | Fzd2 | ec |
| Sfrp2 | batch | fap | d4 | Fzd2 | ec |
| Sfrp4 | batch | fap | d4 | Fzd2 | ec |
| Spp1 | batch | ec | d4 | Itga5 | ec |
| Spp1 | batch | fap | d4 | Itga5 | ec |
| Spp1 | batch | ic | d4 | Itga5 | ec |
| Spp1 | batch | mp | d4 | Itga5 | ec |
| Spp1 | batch | ec | d4 | Vtn | ec |
| Spp1 | batch | fap | d4 | Vtn | ec |
| Spp1 | batch | ic | d4 | Vtn | ec |
| Spp1 | batch | mp | d4 | Vtn | ec |
| Spp1 | batch | ec | d4 | Itgb3 | ec |
| Spp1 | batch | fap | d4 | Itgb3 | ec |
| Spp1 | batch | ic | d4 | Itgb3 | ec |
| Spp1 | batch | mp | d4 | Itgb3 | ec |
| Spp1 | batch | ec | d4 | Itga9 | ec |
| Spp1 | batch | fap | d4 | Itga9 | ec |
| Spp1 | batch | ic | d4 | Itga9 | ec |
| Spp1 | batch | mp | d4 | Itga9 | ec |
| Tgfb3 | batch | fap | d4 | Acvrl1 | ec |
| Tgfb3 | total | ec | d4 | Acvrl1 | ec |
| Tgfb3 | total | per | d4 | Acvrl1 | ec |
| Tnfrsf11b | batch | fap | d4 | Vtn | ec |
| Clu | batch | ec | d4 | Vldlr | ec |
| Csf3 | batch | ec | d4 | Csf3r | ec |
| Dll1 | batch | ec | d4 | Notch1 | ec |
| Dll1 | total | mp | d4 | Notch1 | ec |
| Fgf9 | batch | ec | d4 | Fgfr3 | ec |
| Il15 | batch | ec | d4 | Il17ra | ec |
| Il15 | batch | ic | d4 | Il17ra | ec |
| Il15 | batch | ec | d4 | Il15ra | ec |
| Il15 | batch | ic | d4 | Il15ra | ec |
| Lgals3 | batch | ec | d4 | Lgals3bp | ec |
| Lgals3 | batch | ic | d4 | Lgals3bp | ec |
| Lgals3 | batch | mp | d4 | Lgals3bp | ec |
| Osm | batch | ec | d4 | Osmr | ec |
| Osm | batch | ec | d4 | Il6st | ec |
| Osm | batch | ec | d4 | Lifr | ec |
| Tnfsf10 | batch | ec | d4 | Tnfrsf10b | ec |
| Vegfc | batch | ec | d4 | Flt4 | ec |
| Vegfc | batch | mp | d4 | Flt4 | ec |
| Vegfc | batch | ec | d4 | Kdr | ec |
| Vegfc | batch | mp | d4 | Kdr | ec |
| Vegfc | batch | ec | d4 | Nrp2 | ec |
| Vegfc | batch | mp | d4 | Nrp2 | ec |
| Vegfc | batch | ec | d4 | Nrp1 | ec |
| Vegfc | batch | mp | d4 | Nrp1 | ec |
| Vegfc | batch | ec | d4 | Itga9 | ec |
| Vegfc | batch | mp | d4 | Itga9 | ec |
| Areg | batch | mp | d4 | Egfr | ec |
| Ccl8 | batch | mp | d4 | Ccr5 | ec |
| Ccl8 | total | fap | d4 | Ccr5 | ec |
| Ccl8 | total | ic | d4 | Ccr5 | ec |
| Ebi3 | batch | mp | d4 | Il27ra | ec |
| Ebi3 | constitutive | ic | d4 | Il27ra | ec |
| Gdnf | batch | mp | d4 | Ret | ec |
| Gmfb | batch | mp | d4 | Egfr | ec |
| Gmfb | batch | mp | d4 | Itpr3 | ec |
| Il10 | batch | ic | d4 | Il10ra | ec |
| Il10 | batch | mp | d4 | Il10ra | ec |
| Pdap1 | batch | mp | d4 | Pdgfa | ec |
| Rabep1 | batch | mp | d4 | Lsr | ec |
| Tgfb1 | batch | mp | d4 | Eng | ec |
| Tgfb1 | batch | mp | d4 | Acvrl1 | ec |
| Tgfb1 | batch | mp | d4 | Tgfbr3 | ec |
| Tgfb1 | batch | mp | d4 | Vtn | ec |
| Tgfb2 | batch | mp | d4 | Tgfbr3 | ec |
| Tgfb2 | constitutive | ec | d4 | Tgfbr3 | ec |
| Tgfb2 | constitutive | fap | d4 | Tgfbr3 | ec |
| Tgfb2 | constitutive | per | d4 | Tgfbr3 | ec |
| Tgfb2 | batch | mp | d4 | Vtn | ec |
| Tgfb2 | constitutive | ec | d4 | Vtn | ec |
| Tgfb2 | constitutive | fap | d4 | Vtn | ec |
| Tgfb2 | constitutive | per | d4 | Vtn | ec |
| Pdgfb | batch | ic | d4 | Pdgfrb | ec |
| Pdgfb | constitutive | ec | d4 | Pdgfrb | ec |
| Tnfsf13 | batch | ic | d4 | Tnfrsf11b | ec |
| Tnfsf13 | batch | ic | d4 | Tnfrsf25 | ec |
| Tnfsf13 | batch | ic | d4 | Tnfrsf1a | ec |
| Tnfsf13 | batch | ic | d4 | Fas | ec |
| Angpt1 | constitutive | fap | d4 | Tek | ec |
| Fgf7 | constitutive | fap | d4 | Fgfr3 | ec |
| Fgf7 | constitutive | fap | d4 | Nrp1 | ec |
| Bmp5 | constitutive | ec | d4 | Bmpr1a | ec |
| Bmp5 | constitutive | fap | d4 | Bmpr1a | ec |
| Bmp5 | constitutive | per | d4 | Bmpr1a | ec |
| Inhbb | constitutive | fap | d4 | Acvr2a | ec |
| Inhbb | constitutive | per | d4 | Acvr2a | ec |
| Jag2 | constitutive | ec | d4 | Notch1 | ec |
| Cmtm8 | constitutive | ec | d4 | Egfr | ec |
| Edn1 | constitutive | ec | d4 | Ednra | ec |
| Angpt2 | constitutive | ec | d4 | Tek | ec |
| Angpt2 | constitutive | ic | d4 | Tek | ec |
| Angpt2 | constitutive | per | d4 | Tek | ec |
| Bmp4 | constitutive | ec | d4 | Bmpr1a | ec |
| Bmp4 | constitutive | per | d4 | Bmpr1a | ec |
| Bmp4 | constitutive | ec | d4 | Bmpr2 | ec |
| Bmp4 | constitutive | per | d4 | Bmpr2 | ec |
| Lif | constitutive | mp | d4 | Il6st | ec |
| Lif | constitutive | mp | d4 | Lifr | ec |
| Nampt | constitutive | ic | d4 | Adora2a | ec |
| Serpini1 | constitutive | per | d4 | Plat | ec |
| Bmp6 | constitutive | per | d4 | Bmpr1a | ec |
| Bmp6 | constitutive | per | d4 | Acvr2a | ec |
| Bmp6 | constitutive | per | d4 | Bmpr2 | ec |
| Pgf | constitutive | per | d4 | Flt1 | ec |
| Pgf | constitutive | per | d4 | Nrp2 | ec |
| Pgf | constitutive | per | d4 | Nrp1 | ec |
| Jag1 | constitutive | per | d4 | Notch1 | ec |
| Bmp2 | constitutive | per | d4 | Eng | ec |
| Bmp2 | constitutive | per | d4 | Bmpr1a | ec |
| Bmp2 | constitutive | per | d4 | Acvr2a | ec |
| Bmp2 | constitutive | per | d4 | Bmpr2 | ec |
| Apln | batch | ec | d5 | Aplnr | ec |
| Apln | batch | fap | d5 | Aplnr | ec |
| Ctgf | batch | fap | d5 | Itga5 | ec |
| Ctgf | batch | ic | d5 | Itga5 | ec |
| Ctgf | constitutive | per | d5 | Itga5 | ec |
| Efnb1 | batch | fap | d5 | Ephb2 | ec |
| Efnb1 | batch | ic | d5 | Ephb2 | ec |
| Efnb1 | batch | mp | d5 | Ephb2 | ec |
| Efnb1 | constitutive | ec | d5 | Ephb2 | ec |
| Efnb1 | constitutive | per | d5 | Ephb2 | ec |
| Fgf1 | batch | fap | d5 | Fgfr3 | ec |
| Fgf1 | constitutive | per | d5 | Fgfr3 | ec |
| Fgf1 | batch | fap | d5 | Fgfr4 | ec |
| Fgf1 | constitutive | per | d5 | Fgfr4 | ec |
| Hbegf | batch | fap | d5 | Egfr | ec |
| Hbegf | constitutive | ec | d5 | Egfr | ec |
| Hbegf | constitutive | per | d5 | Egfr | ec |
| Hgf | batch | fap | d5 | Vtn | ec |
| Hgf | batch | ic | d5 | Vtn | ec |
| Igf1 | batch | ec | d5 | Igfbp5 | ec |
| Igf1 | batch | fap | d5 | Igfbp5 | ec |
| Igf1 | batch | ic | d5 | Igfbp5 | ec |
| Igf1 | batch | per | d5 | Igfbp5 | ec |
| Igf1 | batch | ec | d5 | Igfbp7 | ec |
| Igf1 | batch | fap | d5 | Igfbp7 | ec |
| Igf1 | batch | ic | d5 | Igfbp7 | ec |
| Igf1 | batch | per | d5 | Igfbp7 | ec |
| Igf1 | batch | ec | d5 | Igfbp2 | ec |
| Igf1 | batch | fap | d5 | Igfbp2 | ec |
| Igf1 | batch | ic | d5 | Igfbp2 | ec |
| Igf1 | batch | per | d5 | Igfbp2 | ec |
| Igf2 | batch | ec | d5 | Insr | ec |
| Igf2 | batch | fap | d5 | Insr | ec |
| Igf2 | batch | ic | d5 | Insr | ec |
| Igf2 | batch | mp | d5 | Insr | ec |
| Igf2 | batch | ec | d5 | Vtn | ec |
| Igf2 | batch | fap | d5 | Vtn | ec |
| Igf2 | batch | ic | d5 | Vtn | ec |
| Igf2 | batch | mp | d5 | Vtn | ec |
| Il1b | batch | fap | d5 | Adrb2 | ec |
| Il33 | batch | fap | d5 | Il1rl1 | ec |
| Il33 | batch | mp | d5 | Il1rl1 | ec |
| Il33 | batch | per | d5 | Il1rl1 | ec |
| Kitl | batch | fap | d5 | Kit | ec |
| Kitl | batch | ic | d5 | Kit | ec |
| Kitl | batch | mp | d5 | Kit | ec |
| Kitl | constitutive | ec | d5 | Kit | ec |
| Kitl | constitutive | per | d5 | Kit | ec |
| Mmp13 | batch | ec | d5 | F2r | ec |
| Mmp13 | batch | fap | d5 | F2r | ec |
| Mmp13 | batch | ic | d5 | F2r | ec |
| Mmp13 | batch | mp | d5 | F2r | ec |
| Mmp13 | batch | per | d5 | F2r | ec |
| Mmp9 | batch | fap | d5 | Flt1 | ec |
| Mmp9 | batch | per | d5 | Flt1 | ec |
| Ntf3 | batch | ec | d5 | Ntrk2 | ec |
| Ntf3 | batch | fap | d5 | Ntrk2 | ec |
| Ntf3 | constitutive | per | d5 | Ntrk2 | ec |
| Sfrp1 | batch | fap | d5 | Fzd2 | ec |
| Sfrp1 | constitutive | per | d5 | Fzd2 | ec |
| Sfrp2 | batch | fap | d5 | Fzd2 | ec |
| Sfrp2 | batch | per | d5 | Fzd2 | ec |
| Sfrp4 | batch | fap | d5 | Fzd2 | ec |
| Sfrp4 | batch | per | d5 | Fzd2 | ec |
| Spp1 | batch | fap | d5 | Itga5 | ec |
| Spp1 | batch | ic | d5 | Itga5 | ec |
| Spp1 | batch | mp | d5 | Itga5 | ec |
| Spp1 | batch | fap | d5 | Vtn | ec |
| Spp1 | batch | ic | d5 | Vtn | ec |
| Spp1 | batch | mp | d5 | Vtn | ec |
| Spp1 | batch | fap | d5 | Itgb3 | ec |
| Spp1 | batch | ic | d5 | Itgb3 | ec |
| Spp1 | batch | mp | d5 | Itgb3 | ec |
| Spp1 | batch | fap | d5 | Itga9 | ec |
| Spp1 | batch | ic | d5 | Itga9 | ec |
| Spp1 | batch | mp | d5 | Itga9 | ec |
| Tgfb3 | batch | fap | d5 | Acvrl1 | ec |
| Tgfb3 | total | ec | d5 | Acvrl1 | ec |
| Tgfb3 | total | per | d5 | Acvrl1 | ec |
| Tnfrsf11b | batch | fap | d5 | Vtn | ec |
| Clu | batch | ec | d5 | Vldlr | ec |
| Clu | batch | ic | d5 | Vldlr | ec |
| Dll1 | batch | ec | d5 | Notch1 | ec |
| Dll1 | total | mp | d5 | Notch1 | ec |
| Fgf9 | batch | ec | d5 | Fgfr3 | ec |
| Fgf9 | batch | ec | d5 | Fgfr4 | ec |
| Il15 | batch | ec | d5 | Il15ra | ec |
| Il15 | batch | ic | d5 | Il15ra | ec |
| Il6 | batch | ec | d5 | Il6st | ec |
| Il6 | batch | per | d5 | Il6st | ec |
| Inhbb | batch | ec | d5 | Acvr2a | ec |
| Inhbb | constitutive | fap | d5 | Acvr2a | ec |
| Inhbb | constitutive | per | d5 | Acvr2a | ec |
| Tnfsf10 | batch | ec | d5 | Tnfrsf10b | ec |
| Vegfc | batch | ec | d5 | Flt4 | ec |
| Vegfc | batch | mp | d5 | Flt4 | ec |
| Vegfc | batch | ec | d5 | Kdr | ec |
| Vegfc | batch | mp | d5 | Kdr | ec |
| Vegfc | batch | ec | d5 | Nrp2 | ec |
| Vegfc | batch | mp | d5 | Nrp2 | ec |
| Vegfc | batch | ec | d5 | Nrp1 | ec |
| Vegfc | batch | mp | d5 | Nrp1 | ec |
| Vegfc | batch | ec | d5 | Itga9 | ec |
| Vegfc | batch | mp | d5 | Itga9 | ec |
| Areg | batch | mp | d5 | Egfr | ec |
| Gdnf | batch | mp | d5 | Ret | ec |
| Gdnf | batch | per | d5 | Ret | ec |
| Gmfb | batch | mp | d5 | Egfr | ec |
| Gmfb | batch | mp | d5 | Itpr3 | ec |
| Pdap1 | batch | mp | d5 | Pdgfa | ec |
| Rabep1 | batch | mp | d5 | Lsr | ec |
| Tgfb2 | batch | mp | d5 | Tgfbr3 | ec |
| Tgfb2 | constitutive | ec | d5 | Tgfbr3 | ec |
| Tgfb2 | constitutive | fap | d5 | Tgfbr3 | ec |
| Tgfb2 | constitutive | per | d5 | Tgfbr3 | ec |
| Tgfb2 | batch | mp | d5 | Vtn | ec |
| Tgfb2 | constitutive | ec | d5 | Vtn | ec |
| Tgfb2 | constitutive | fap | d5 | Vtn | ec |
| Tgfb2 | constitutive | per | d5 | Vtn | ec |
| Csf1 | batch | ic | d5 | H2-Bl | ec |
| Il10 | batch | ic | d5 | Il10ra | ec |
| Pdgfb | batch | ic | d5 | Pdgfrb | ec |
| Pdgfb | constitutive | ec | d5 | Pdgfrb | ec |
| Tnfsf13 | batch | ic | d5 | Tnfrsf11b | ec |
| Tnfsf13 | batch | ic | d5 | Tnfrsf25 | ec |
| Tnfsf13 | batch | ic | d5 | Tnfrsf1a | ec |
| Tnfsf13 | batch | ic | d5 | Fas | ec |
| Adm | batch | per | d5 | Calcrl | ec |
| Angpt4 | batch | per | d5 | Tek | ec |
| Btc | batch | per | d5 | Erbb2 | ec |
| Btc | batch | per | d5 | Egfr | ec |
| Fgf18 | batch | per | d5 | Fgfr4 | ec |
| Il11 | batch | per | d5 | Il6st | ec |
| Lif | batch | per | d5 | Il6st | ec |
| Lif | constitutive | mp | d5 | Il6st | ec |
| Lif | batch | per | d5 | Lifr | ec |
| Lif | constitutive | mp | d5 | Lifr | ec |
| Nov | batch | per | d5 | Notch1 | ec |
| Nov | batch | per | d5 | Itga5 | ec |
| Nov | batch | per | d5 | Itgb3 | ec |
| Sfrp5 | batch | per | d5 | Fzd2 | ec |
| Angpt1 | constitutive | fap | d5 | Tek | ec |
| Fgf7 | constitutive | fap | d5 | Fgfr3 | ec |
| Fgf7 | constitutive | fap | d5 | Fgfr4 | ec |
| Fgf7 | constitutive | fap | d5 | Nrp1 | ec |
| Bmp5 | constitutive | ec | d5 | Bmpr1a | ec |
| Bmp5 | constitutive | fap | d5 | Bmpr1a | ec |
| Bmp5 | constitutive | per | d5 | Bmpr1a | ec |
| Jag2 | constitutive | ec | d5 | Notch1 | ec |
| Cmtm8 | constitutive | ec | d5 | Egfr | ec |
| Edn1 | constitutive | ec | d5 | Ednra | ec |
| Angpt2 | constitutive | ec | d5 | Tek | ec |
| Angpt2 | constitutive | ic | d5 | Tek | ec |
| Angpt2 | constitutive | per | d5 | Tek | ec |
| Bmp4 | constitutive | ec | d5 | Bmpr1a | ec |
| Bmp4 | constitutive | per | d5 | Bmpr1a | ec |
| Bmp4 | constitutive | ec | d5 | Bmpr2 | ec |
| Bmp4 | constitutive | per | d5 | Bmpr2 | ec |
| Ebi3 | constitutive | ic | d5 | Il27ra | ec |
| Nampt | constitutive | ic | d5 | Adora2a | ec |
| Serpini1 | constitutive | per | d5 | Plat | ec |
| Bmp6 | constitutive | per | d5 | Bmpr1a | ec |
| Bmp6 | constitutive | per | d5 | Acvr2a | ec |
| Bmp6 | constitutive | per | d5 | Bmpr2 | ec |
| Pgf | constitutive | per | d5 | Flt1 | ec |
| Pgf | constitutive | per | d5 | Nrp2 | ec |
| Pgf | constitutive | per | d5 | Nrp1 | ec |
| Jag1 | constitutive | per | d5 | Notch1 | ec |
| Bmp2 | constitutive | per | d5 | Eng | ec |
| Bmp2 | constitutive | per | d5 | Bmpr1a | ec |
| Bmp2 | constitutive | per | d5 | Acvr2a | ec |
| Bmp2 | constitutive | per | d5 | Bmpr2 | ec |
| Crlf1 | batch | fap | d6 | Ctf1 | ec |
| Crlf1 | batch | mp | d6 | Ctf1 | ec |
| Ctgf | batch | fap | d6 | Itga5 | ec |
| Ctgf | batch | ic | d6 | Itga5 | ec |
| Ctgf | batch | mp | d6 | Itga5 | ec |
| Ctgf | constitutive | per | d6 | Itga5 | ec |
| Efnb1 | batch | fap | d6 | Ephb2 | ec |
| Efnb1 | batch | ic | d6 | Ephb2 | ec |
| Efnb1 | batch | mp | d6 | Ephb2 | ec |
| Efnb1 | constitutive | ec | d6 | Ephb2 | ec |
| Efnb1 | constitutive | per | d6 | Ephb2 | ec |
| Fgf1 | batch | fap | d6 | Fgfr3 | ec |
| Fgf1 | constitutive | per | d6 | Fgfr3 | ec |
| Fgf1 | batch | fap | d6 | Fgfr4 | ec |
| Fgf1 | constitutive | per | d6 | Fgfr4 | ec |
| Hbegf | batch | fap | d6 | Egfr | ec |
| Hbegf | batch | mp | d6 | Egfr | ec |
| Hbegf | constitutive | ec | d6 | Egfr | ec |
| Hbegf | constitutive | per | d6 | Egfr | ec |
| Igf1 | batch | ec | d6 | Igfbp5 | ec |
| Igf1 | batch | fap | d6 | Igfbp5 | ec |
| Igf1 | batch | ic | d6 | Igfbp5 | ec |
| Igf1 | batch | ec | d6 | Igfbp7 | ec |
| Igf1 | batch | fap | d6 | Igfbp7 | ec |
| Igf1 | batch | ic | d6 | Igfbp7 | ec |
| Igf1 | batch | ec | d6 | Igfbp2 | ec |
| Igf1 | batch | fap | d6 | Igfbp2 | ec |
| Igf1 | batch | ic | d6 | Igfbp2 | ec |
| Igf2 | batch | ec | d6 | Insr | ec |
| Igf2 | batch | fap | d6 | Insr | ec |
| Igf2 | batch | ic | d6 | Insr | ec |
| Igf2 | batch | mp | d6 | Insr | ec |
| Igf2 | batch | ec | d6 | Vtn | ec |
| Igf2 | batch | fap | d6 | Vtn | ec |
| Igf2 | batch | ic | d6 | Vtn | ec |
| Igf2 | batch | mp | d6 | Vtn | ec |
| Il1b | batch | fap | d6 | Adrb2 | ec |
| Il33 | batch | fap | d6 | Il1rl1 | ec |
| Il33 | batch | mp | d6 | Il1rl1 | ec |
| Kitl | batch | fap | d6 | Kit | ec |
| Kitl | batch | ic | d6 | Kit | ec |
| Kitl | batch | mp | d6 | Kit | ec |
| Kitl | constitutive | ec | d6 | Kit | ec |
| Kitl | constitutive | per | d6 | Kit | ec |
| Mmp13 | batch | ec | d6 | F2r | ec |
| Mmp13 | batch | fap | d6 | F2r | ec |
| Mmp13 | batch | ic | d6 | F2r | ec |
| Mmp13 | batch | mp | d6 | F2r | ec |
| Mmp9 | batch | fap | d6 | Flt1 | ec |
| Mmp9 | batch | mp | d6 | Flt1 | ec |
| Ntf3 | batch | ec | d6 | Ntrk2 | ec |
| Ntf3 | batch | fap | d6 | Ntrk2 | ec |
| Ntf3 | batch | mp | d6 | Ntrk2 | ec |
| Ntf3 | constitutive | per | d6 | Ntrk2 | ec |
| Sfrp1 | batch | fap | d6 | Fzd2 | ec |
| Sfrp1 | constitutive | per | d6 | Fzd2 | ec |
| Sfrp2 | batch | fap | d6 | Fzd2 | ec |
| Sfrp4 | batch | fap | d6 | Fzd2 | ec |
| Sfrp4 | batch | mp | d6 | Fzd2 | ec |
| Spp1 | batch | fap | d6 | Itga5 | ec |
| Spp1 | batch | ic | d6 | Itga5 | ec |
| Spp1 | batch | mp | d6 | Itga5 | ec |
| Spp1 | batch | fap | d6 | Vtn | ec |
| Spp1 | batch | ic | d6 | Vtn | ec |
| Spp1 | batch | mp | d6 | Vtn | ec |
| Spp1 | batch | fap | d6 | Itgb3 | ec |
| Spp1 | batch | ic | d6 | Itgb3 | ec |
| Spp1 | batch | mp | d6 | Itgb3 | ec |
| Spp1 | batch | fap | d6 | Itga9 | ec |
| Spp1 | batch | ic | d6 | Itga9 | ec |
| Spp1 | batch | mp | d6 | Itga9 | ec |
| Tgfb3 | batch | fap | d6 | Acvrl1 | ec |
| Tgfb3 | batch | mp | d6 | Acvrl1 | ec |
| Tgfb3 | total | ec | d6 | Acvrl1 | ec |
| Tgfb3 | total | per | d6 | Acvrl1 | ec |
| Apln | batch | ec | d6 | Aplnr | ec |
| Clu | batch | ec | d6 | Vldlr | ec |
| Clu | batch | ic | d6 | Vldlr | ec |
| Clu | batch | mp | d6 | Vldlr | ec |
| Dll1 | batch | ec | d6 | Notch1 | ec |
| Dll1 | total | mp | d6 | Notch1 | ec |
| Fgf9 | batch | ec | d6 | Fgfr3 | ec |
| Fgf9 | batch | ec | d6 | Fgfr4 | ec |
| Il15 | batch | ec | d6 | Il15ra | ec |
| Il15 | batch | ic | d6 | Il15ra | ec |
| Il6 | batch | ec | d6 | Il6st | ec |
| Inhbb | batch | ec | d6 | Acvr2a | ec |
| Inhbb | batch | mp | d6 | Acvr2a | ec |
| Inhbb | constitutive | fap | d6 | Acvr2a | ec |
| Inhbb | constitutive | per | d6 | Acvr2a | ec |
| Tnfsf10 | batch | ec | d6 | Tnfrsf10b | ec |
| Vegfc | batch | ec | d6 | Flt4 | ec |
| Vegfc | batch | mp | d6 | Flt4 | ec |
| Vegfc | batch | ec | d6 | Kdr | ec |
| Vegfc | batch | mp | d6 | Kdr | ec |
| Vegfc | batch | ec | d6 | Nrp2 | ec |
| Vegfc | batch | mp | d6 | Nrp2 | ec |
| Vegfc | batch | ec | d6 | Nrp1 | ec |
| Vegfc | batch | mp | d6 | Nrp1 | ec |
| Vegfc | batch | ec | d6 | Itga9 | ec |
| Vegfc | batch | mp | d6 | Itga9 | ec |
| Bmp2 | batch | mp | d6 | Eng | ec |
| Bmp2 | constitutive | per | d6 | Eng | ec |
| Bmp2 | batch | mp | d6 | Bmpr1a | ec |
| Bmp2 | constitutive | per | d6 | Bmpr1a | ec |
| Bmp2 | batch | mp | d6 | Acvr2a | ec |
| Bmp2 | constitutive | per | d6 | Acvr2a | ec |
| Bmp2 | batch | mp | d6 | Bmpr2 | ec |
| Bmp2 | constitutive | per | d6 | Bmpr2 | ec |
| Bmp4 | batch | mp | d6 | Bmpr1a | ec |
| Bmp4 | constitutive | ec | d6 | Bmpr1a | ec |
| Bmp4 | constitutive | per | d6 | Bmpr1a | ec |
| Bmp4 | batch | mp | d6 | Bmpr2 | ec |
| Bmp4 | constitutive | ec | d6 | Bmpr2 | ec |
| Bmp4 | constitutive | per | d6 | Bmpr2 | ec |
| Bmp6 | batch | mp | d6 | Bmpr1a | ec |
| Bmp6 | constitutive | per | d6 | Bmpr1a | ec |
| Bmp6 | batch | mp | d6 | Acvr2a | ec |
| Bmp6 | constitutive | per | d6 | Acvr2a | ec |
| Bmp6 | batch | mp | d6 | Bmpr2 | ec |
| Bmp6 | constitutive | per | d6 | Bmpr2 | ec |
| Clcf1 | batch | mp | d6 | Crlf1 | ec |
| Ctf1 | batch | mp | d6 | Il6st | ec |
| Ctsg | batch | mp | d6 | F2rl3 | ec |
| Ebi3 | batch | mp | d6 | Il27ra | ec |
| Ebi3 | constitutive | ic | d6 | Il27ra | ec |
| Edn3 | batch | mp | d6 | Ednra | ec |
| Il10 | batch | ic | d6 | Il10ra | ec |
| Il10 | batch | mp | d6 | Il10ra | ec |
| Jag1 | batch | mp | d6 | Notch1 | ec |
| Jag1 | constitutive | per | d6 | Notch1 | ec |
| Jag2 | batch | mp | d6 | Notch1 | ec |
| Jag2 | constitutive | ec | d6 | Notch1 | ec |
| Nppc | batch | mp | d6 | Npr2 | ec |
| Pdap1 | batch | mp | d6 | Pdgfa | ec |
| Serpini1 | batch | mp | d6 | Plat | ec |
| Serpini1 | constitutive | per | d6 | Plat | ec |
| Sfrp5 | batch | mp | d6 | Fzd2 | ec |
| Tgfb1 | batch | mp | d6 | Eng | ec |
| Tgfb1 | batch | mp | d6 | Acvrl1 | ec |
| Tgfb1 | batch | mp | d6 | Tgfbr3 | ec |
| Tgfb1 | batch | mp | d6 | Vtn | ec |
| Tgfb2 | batch | mp | d6 | Tgfbr3 | ec |
| Tgfb2 | constitutive | ec | d6 | Tgfbr3 | ec |
| Tgfb2 | constitutive | fap | d6 | Tgfbr3 | ec |
| Tgfb2 | constitutive | per | d6 | Tgfbr3 | ec |
| Tgfb2 | batch | mp | d6 | Vtn | ec |
| Tgfb2 | constitutive | ec | d6 | Vtn | ec |
| Tgfb2 | constitutive | fap | d6 | Vtn | ec |
| Tgfb2 | constitutive | per | d6 | Vtn | ec |
| Tnfsf12 | batch | mp | d6 | Tnfrsf25 | ec |
| Tnfsf12 | batch | mp | d6 | Tnfrsf1a | ec |
| Tnfsf12 | batch | mp | d6 | Fas | ec |
| Vegfa | batch | mp | d6 | Flt1 | ec |
| Vegfa | batch | mp | d6 | Kdr | ec |
| Vegfa | batch | mp | d6 | Nrp2 | ec |
| Vegfa | batch | mp | d6 | Nrp1 | ec |
| Vegfa | batch | mp | d6 | Ephb2 | ec |
| Vegfa | batch | mp | d6 | Vtn | ec |
| Vegfa | batch | mp | d6 | Itga9 | ec |
| Csf1 | batch | ic | d6 | H2-Bl | ec |
| Hgf | batch | ic | d6 | Vtn | ec |
| Pdgfb | batch | ic | d6 | Pdgfrb | ec |
| Pdgfb | constitutive | ec | d6 | Pdgfrb | ec |
| Tnfsf13 | batch | ic | d6 | Tnfrsf25 | ec |
| Tnfsf13 | batch | ic | d6 | Tnfrsf1a | ec |
| Tnfsf13 | batch | ic | d6 | Fas | ec |
| Angpt1 | constitutive | fap | d6 | Tek | ec |
| Fgf7 | constitutive | fap | d6 | Fgfr3 | ec |
| Fgf7 | constitutive | fap | d6 | Fgfr4 | ec |
| Fgf7 | constitutive | fap | d6 | Nrp1 | ec |
| Bmp5 | constitutive | ec | d6 | Bmpr1a | ec |
| Bmp5 | constitutive | fap | d6 | Bmpr1a | ec |
| Bmp5 | constitutive | per | d6 | Bmpr1a | ec |
| Cmtm8 | constitutive | ec | d6 | Egfr | ec |
| Edn1 | constitutive | ec | d6 | Ednra | ec |
| Angpt2 | constitutive | ec | d6 | Tek | ec |
| Angpt2 | constitutive | ic | d6 | Tek | ec |
| Angpt2 | constitutive | per | d6 | Tek | ec |
| Lif | constitutive | mp | d6 | Il6st | ec |
| Lif | constitutive | mp | d6 | Lifr | ec |
| Nampt | constitutive | ic | d6 | Adora2a | ec |
| Pgf | constitutive | per | d6 | Flt1 | ec |
| Pgf | constitutive | per | d6 | Nrp2 | ec |
| Pgf | constitutive | per | d6 | Nrp1 | ec |
| Bmp2 | batch | fap | d7 | Eng | ec |
| Bmp2 | batch | mp | d7 | Eng | ec |
| Bmp2 | constitutive | per | d7 | Eng | ec |
| Bmp2 | batch | fap | d7 | Bmpr1a | ec |
| Bmp2 | batch | mp | d7 | Bmpr1a | ec |
| Bmp2 | constitutive | per | d7 | Bmpr1a | ec |
| Bmp2 | batch | fap | d7 | Acvr2a | ec |
| Bmp2 | batch | mp | d7 | Acvr2a | ec |
| Bmp2 | constitutive | per | d7 | Acvr2a | ec |
| Bmp2 | batch | fap | d7 | Bmpr2 | ec |
| Bmp2 | batch | mp | d7 | Bmpr2 | ec |
| Bmp2 | constitutive | per | d7 | Bmpr2 | ec |
| Bmp4 | batch | fap | d7 | Bmpr1a | ec |
| Bmp4 | batch | mp | d7 | Bmpr1a | ec |
| Bmp4 | constitutive | ec | d7 | Bmpr1a | ec |
| Bmp4 | constitutive | per | d7 | Bmpr1a | ec |
| Bmp4 | batch | fap | d7 | Bmpr2 | ec |
| Bmp4 | batch | mp | d7 | Bmpr2 | ec |
| Bmp4 | constitutive | ec | d7 | Bmpr2 | ec |
| Bmp4 | constitutive | per | d7 | Bmpr2 | ec |
| Bmp6 | batch | ec | d7 | Bmpr1a | ec |
| Bmp6 | batch | fap | d7 | Bmpr1a | ec |
| Bmp6 | batch | mp | d7 | Bmpr1a | ec |
| Bmp6 | constitutive | per | d7 | Bmpr1a | ec |
| Bmp6 | batch | ec | d7 | Acvr2a | ec |
| Bmp6 | batch | fap | d7 | Acvr2a | ec |
| Bmp6 | batch | mp | d7 | Acvr2a | ec |
| Bmp6 | constitutive | per | d7 | Acvr2a | ec |
| Bmp6 | batch | ec | d7 | Bmpr2 | ec |
| Bmp6 | batch | fap | d7 | Bmpr2 | ec |
| Bmp6 | batch | mp | d7 | Bmpr2 | ec |
| Bmp6 | constitutive | per | d7 | Bmpr2 | ec |
| Bmp7 | batch | fap | d7 | Eng | ec |
| Bmp7 | batch | fap | d7 | Bmpr1a | ec |
| Bmp7 | batch | fap | d7 | Acvr2a | ec |
| Bmp7 | batch | fap | d7 | Bmpr2 | ec |
| Clu | batch | ec | d7 | Vldlr | ec |
| Clu | batch | fap | d7 | Vldlr | ec |
| Clu | batch | ic | d7 | Vldlr | ec |
| Clu | batch | mp | d7 | Vldlr | ec |
| Crlf1 | batch | ec | d7 | Ctf1 | ec |
| Crlf1 | batch | fap | d7 | Ctf1 | ec |
| Crlf1 | batch | mp | d7 | Ctf1 | ec |
| Csf2 | batch | fap | d7 | Csf2rb | ec |
| Ctgf | batch | ec | d7 | Itga5 | ec |
| Ctgf | batch | fap | d7 | Itga5 | ec |
| Ctgf | batch | ic | d7 | Itga5 | ec |
| Ctgf | batch | mp | d7 | Itga5 | ec |
| Ctgf | constitutive | per | d7 | Itga5 | ec |
| Edn1 | batch | fap | d7 | Ednra | ec |
| Edn1 | constitutive | ec | d7 | Ednra | ec |
| Fgf1 | batch | fap | d7 | Fgfr3 | ec |
| Fgf1 | constitutive | per | d7 | Fgfr3 | ec |
| Fgf1 | batch | fap | d7 | Fgfr4 | ec |
| Fgf1 | constitutive | per | d7 | Fgfr4 | ec |
| Fgf18 | batch | fap | d7 | Fgfr4 | ec |
| Hbegf | batch | fap | d7 | Egfr | ec |
| Hbegf | batch | mp | d7 | Egfr | ec |
| Hbegf | constitutive | ec | d7 | Egfr | ec |
| Hbegf | constitutive | per | d7 | Egfr | ec |
| Igf1 | batch | ec | d7 | Igfbp3 | ec |
| Igf1 | batch | fap | d7 | Igfbp3 | ec |
| Igf1 | batch | ic | d7 | Igfbp3 | ec |
| Igf1 | batch | ec | d7 | Igfbp5 | ec |
| Igf1 | batch | fap | d7 | Igfbp5 | ec |
| Igf1 | batch | ic | d7 | Igfbp5 | ec |
| Igf1 | batch | ec | d7 | Igfbp7 | ec |
| Igf1 | batch | fap | d7 | Igfbp7 | ec |
| Igf1 | batch | ic | d7 | Igfbp7 | ec |
| Igf1 | batch | ec | d7 | Igfbp2 | ec |
| Igf1 | batch | fap | d7 | Igfbp2 | ec |
| Igf1 | batch | ic | d7 | Igfbp2 | ec |
| Igf2 | batch | ec | d7 | Insr | ec |
| Igf2 | batch | fap | d7 | Insr | ec |
| Igf2 | batch | ic | d7 | Insr | ec |
| Igf2 | batch | mp | d7 | Insr | ec |
| Igf2 | batch | ec | d7 | Vtn | ec |
| Igf2 | batch | fap | d7 | Vtn | ec |
| Igf2 | batch | ic | d7 | Vtn | ec |
| Igf2 | batch | mp | d7 | Vtn | ec |
| Mmp13 | batch | ec | d7 | F2r | ec |
| Mmp13 | batch | fap | d7 | F2r | ec |
| Mmp13 | batch | mp | d7 | F2r | ec |
| Nov | batch | ec | d7 | Notch1 | ec |
| Nov | batch | fap | d7 | Notch1 | ec |
| Nov | batch | ec | d7 | Itga5 | ec |
| Nov | batch | fap | d7 | Itga5 | ec |
| Nov | batch | ec | d7 | Itgb3 | ec |
| Nov | batch | fap | d7 | Itgb3 | ec |
| Ntf3 | batch | ec | d7 | Ntrk2 | ec |
| Ntf3 | batch | fap | d7 | Ntrk2 | ec |
| Ntf3 | batch | mp | d7 | Ntrk2 | ec |
| Ntf3 | constitutive | per | d7 | Ntrk2 | ec |
| Pgf | batch | fap | d7 | Flt1 | ec |
| Pgf | constitutive | per | d7 | Flt1 | ec |
| Pgf | batch | fap | d7 | Nrp2 | ec |
| Pgf | constitutive | per | d7 | Nrp2 | ec |
| Pgf | batch | fap | d7 | Nrp1 | ec |
| Pgf | constitutive | per | d7 | Nrp1 | ec |
| Sfrp1 | batch | fap | d7 | Fzd2 | ec |
| Sfrp1 | constitutive | per | d7 | Fzd2 | ec |
| Sfrp4 | batch | ec | d7 | Fzd2 | ec |
| Sfrp4 | batch | fap | d7 | Fzd2 | ec |
| Sfrp4 | batch | mp | d7 | Fzd2 | ec |
| Tgfb3 | batch | fap | d7 | Acvrl1 | ec |
| Tgfb3 | batch | mp | d7 | Acvrl1 | ec |
| Tgfb3 | total | ec | d7 | Acvrl1 | ec |
| Tgfb3 | total | per | d7 | Acvrl1 | ec |
| Vegfa | batch | fap | d7 | Flt1 | ec |
| Vegfa | batch | mp | d7 | Flt1 | ec |
| Vegfa | batch | fap | d7 | Kdr | ec |
| Vegfa | batch | mp | d7 | Kdr | ec |
| Vegfa | batch | fap | d7 | Nrp2 | ec |
| Vegfa | batch | mp | d7 | Nrp2 | ec |
| Vegfa | batch | fap | d7 | Nrp1 | ec |
| Vegfa | batch | mp | d7 | Nrp1 | ec |
| Vegfa | batch | fap | d7 | Ephb2 | ec |
| Vegfa | batch | mp | d7 | Ephb2 | ec |
| Vegfa | batch | fap | d7 | Vtn | ec |
| Vegfa | batch | mp | d7 | Vtn | ec |
| Vegfa | batch | fap | d7 | Itga9 | ec |
| Vegfa | batch | mp | d7 | Itga9 | ec |
| Wnt2 | batch | fap | d7 | Sfrp1 | ec |
| Dll1 | batch | ec | d7 | Notch1 | ec |
| Dll1 | total | mp | d7 | Notch1 | ec |
| Fgf9 | batch | ec | d7 | Fgfr3 | ec |
| Fgf9 | batch | ec | d7 | Fgfr4 | ec |
| Il15 | batch | ec | d7 | Il15ra | ec |
| Il15 | batch | ic | d7 | Il15ra | ec |
| Sfrp5 | batch | ec | d7 | Fzd2 | ec |
| Sfrp5 | batch | mp | d7 | Fzd2 | ec |
| Tnfsf10 | batch | ec | d7 | Tnfrsf10b | ec |
| Vegfc | batch | ec | d7 | Flt4 | ec |
| Vegfc | batch | mp | d7 | Flt4 | ec |
| Vegfc | batch | ec | d7 | Kdr | ec |
| Vegfc | batch | mp | d7 | Kdr | ec |
| Vegfc | batch | ec | d7 | Nrp2 | ec |
| Vegfc | batch | mp | d7 | Nrp2 | ec |
| Vegfc | batch | ec | d7 | Nrp1 | ec |
| Vegfc | batch | mp | d7 | Nrp1 | ec |
| Vegfc | batch | ec | d7 | Itga9 | ec |
| Vegfc | batch | mp | d7 | Itga9 | ec |
| Areg | batch | mp | d7 | Egfr | ec |
| Clcf1 | batch | ic | d7 | Crlf1 | ec |
| Clcf1 | batch | mp | d7 | Crlf1 | ec |
| Ctf1 | batch | mp | d7 | Il6st | ec |
| Ctsg | batch | ic | d7 | F2rl3 | ec |
| Ctsg | batch | mp | d7 | F2rl3 | ec |
| Ebi3 | batch | mp | d7 | Il27ra | ec |
| Ebi3 | constitutive | ic | d7 | Il27ra | ec |
| Edn3 | batch | mp | d7 | Ednra | ec |
| Efnb1 | batch | ic | d7 | Ephb2 | ec |
| Efnb1 | batch | mp | d7 | Ephb2 | ec |
| Efnb1 | constitutive | ec | d7 | Ephb2 | ec |
| Efnb1 | constitutive | per | d7 | Ephb2 | ec |
| Gdnf | batch | mp | d7 | Ret | ec |
| Gmfb | batch | mp | d7 | Egfr | ec |
| Gmfb | batch | mp | d7 | Itpr3 | ec |
| Il10 | batch | ic | d7 | Il10ra | ec |
| Il10 | batch | mp | d7 | Il10ra | ec |
| Il33 | batch | mp | d7 | Il1rl1 | ec |
| Inhbb | batch | mp | d7 | Acvr2a | ec |
| Inhbb | constitutive | fap | d7 | Acvr2a | ec |
| Inhbb | constitutive | per | d7 | Acvr2a | ec |
| Jag1 | batch | mp | d7 | Notch1 | ec |
| Jag1 | constitutive | per | d7 | Notch1 | ec |
| Jag2 | batch | mp | d7 | Notch1 | ec |
| Jag2 | constitutive | ec | d7 | Notch1 | ec |
| Kitl | batch | ic | d7 | Kit | ec |
| Kitl | batch | mp | d7 | Kit | ec |
| Kitl | constitutive | ec | d7 | Kit | ec |
| Kitl | constitutive | per | d7 | Kit | ec |
| Mmp9 | batch | ic | d7 | Flt1 | ec |
| Mmp9 | batch | mp | d7 | Flt1 | ec |
| Nppc | batch | mp | d7 | Npr2 | ec |
| Pdap1 | batch | mp | d7 | Pdgfa | ec |
| Rabep1 | batch | mp | d7 | Lsr | ec |
| Serpini1 | batch | mp | d7 | Plat | ec |
| Serpini1 | constitutive | per | d7 | Plat | ec |
| Spp1 | batch | mp | d7 | Itga5 | ec |
| Spp1 | batch | mp | d7 | Vtn | ec |
| Spp1 | batch | mp | d7 | Itgb3 | ec |
| Spp1 | batch | mp | d7 | Itga9 | ec |
| Tgfb1 | batch | mp | d7 | Eng | ec |
| Tgfb1 | batch | mp | d7 | Acvrl1 | ec |
| Tgfb1 | batch | mp | d7 | Tgfbr3 | ec |
| Tgfb1 | batch | mp | d7 | Vtn | ec |
| Tgfb2 | batch | mp | d7 | Tgfbr3 | ec |
| Tgfb2 | constitutive | ec | d7 | Tgfbr3 | ec |
| Tgfb2 | constitutive | fap | d7 | Tgfbr3 | ec |
| Tgfb2 | constitutive | per | d7 | Tgfbr3 | ec |
| Tgfb2 | batch | mp | d7 | Vtn | ec |
| Tgfb2 | constitutive | ec | d7 | Vtn | ec |
| Tgfb2 | constitutive | fap | d7 | Vtn | ec |
| Tgfb2 | constitutive | per | d7 | Vtn | ec |
| Tnfsf12 | batch | mp | d7 | Tnfrsf25 | ec |
| Tnfsf12 | batch | mp | d7 | Tnfrsf1a | ec |
| Tnfsf12 | batch | mp | d7 | Fas | ec |
| Csf1 | batch | ic | d7 | H2-Bl | ec |
| Ifng | batch | ic | d7 | Ifngr1 | ec |
| Il6 | batch | ic | d7 | Il6st | ec |
| Ltb | batch | ic | d7 | Tnfrsf1a | ec |
| Pdgfb | batch | ic | d7 | Pdgfrb | ec |
| Pdgfb | constitutive | ec | d7 | Pdgfrb | ec |
| Angpt1 | constitutive | fap | d7 | Tek | ec |
| Fgf7 | constitutive | fap | d7 | Fgfr3 | ec |
| Fgf7 | constitutive | fap | d7 | Fgfr4 | ec |
| Fgf7 | constitutive | fap | d7 | Nrp1 | ec |
| Bmp5 | constitutive | ec | d7 | Bmpr1a | ec |
| Bmp5 | constitutive | fap | d7 | Bmpr1a | ec |
| Bmp5 | constitutive | per | d7 | Bmpr1a | ec |
| Cmtm8 | constitutive | ec | d7 | Egfr | ec |
| Angpt2 | constitutive | ec | d7 | Tek | ec |
| Angpt2 | constitutive | ic | d7 | Tek | ec |
| Angpt2 | constitutive | per | d7 | Tek | ec |
| Lif | constitutive | mp | d7 | Il6st | ec |
| Lif | constitutive | mp | d7 | Lifr | ec |
| Nampt | constitutive | ic | d7 | Adora2a | ec |
| Bmp2 | batch | fap | d10 | Eng | ec |
| Bmp2 | batch | mp | d10 | Eng | ec |
| Bmp2 | constitutive | per | d10 | Eng | ec |
| Bmp2 | batch | fap | d10 | Bmpr1a | ec |
| Bmp2 | batch | mp | d10 | Bmpr1a | ec |
| Bmp2 | constitutive | per | d10 | Bmpr1a | ec |
| Bmp2 | batch | fap | d10 | Acvr2a | ec |
| Bmp2 | batch | mp | d10 | Acvr2a | ec |
| Bmp2 | constitutive | per | d10 | Acvr2a | ec |
| Bmp2 | batch | fap | d10 | Bmpr2 | ec |
| Bmp2 | batch | mp | d10 | Bmpr2 | ec |
| Bmp2 | constitutive | per | d10 | Bmpr2 | ec |
| Bmp4 | batch | fap | d10 | Bmpr1a | ec |
| Bmp4 | batch | mp | d10 | Bmpr1a | ec |
| Bmp4 | constitutive | ec | d10 | Bmpr1a | ec |
| Bmp4 | constitutive | per | d10 | Bmpr1a | ec |
| Bmp4 | batch | fap | d10 | Bmpr2 | ec |
| Bmp4 | batch | mp | d10 | Bmpr2 | ec |
| Bmp4 | constitutive | ec | d10 | Bmpr2 | ec |
| Bmp4 | constitutive | per | d10 | Bmpr2 | ec |
| Bmp6 | batch | fap | d10 | Bmpr1a | ec |
| Bmp6 | batch | mp | d10 | Bmpr1a | ec |
| Bmp6 | constitutive | per | d10 | Bmpr1a | ec |
| Bmp6 | batch | fap | d10 | Acvr2a | ec |
| Bmp6 | batch | mp | d10 | Acvr2a | ec |
| Bmp6 | constitutive | per | d10 | Acvr2a | ec |
| Bmp6 | batch | fap | d10 | Bmpr2 | ec |
| Bmp6 | batch | mp | d10 | Bmpr2 | ec |
| Bmp6 | constitutive | per | d10 | Bmpr2 | ec |
| Bmp7 | batch | fap | d10 | Eng | ec |
| Bmp7 | batch | fap | d10 | Bmpr1a | ec |
| Bmp7 | batch | fap | d10 | Acvr2a | ec |
| Bmp7 | batch | fap | d10 | Bmpr2 | ec |
| Clu | batch | ec | d10 | Vldlr | ec |
| Clu | batch | fap | d10 | Vldlr | ec |
| Clu | batch | mp | d10 | Vldlr | ec |
| Clu | total | per | d10 | Vldlr | ec |
| Crlf1 | batch | fap | d10 | Ctf1 | ec |
| Crlf1 | batch | mp | d10 | Ctf1 | ec |
| Csf2 | batch | fap | d10 | Csf2rb | ec |
| Csf2 | batch | fap | d10 | Il3ra | ec |
| Ctgf | batch | fap | d10 | Itga5 | ec |
| Ctgf | batch | mp | d10 | Itga5 | ec |
| Ctgf | constitutive | per | d10 | Itga5 | ec |
| Edn1 | batch | fap | d10 | Ednra | ec |
| Edn1 | constitutive | ec | d10 | Ednra | ec |
| Efnb1 | batch | fap | d10 | Ephb2 | ec |
| Efnb1 | batch | mp | d10 | Ephb2 | ec |
| Efnb1 | constitutive | ec | d10 | Ephb2 | ec |
| Efnb1 | constitutive | per | d10 | Ephb2 | ec |
| Fgf1 | batch | fap | d10 | Fgfr3 | ec |
| Fgf1 | batch | mp | d10 | Fgfr3 | ec |
| Fgf1 | constitutive | per | d10 | Fgfr3 | ec |
| Hbegf | batch | fap | d10 | Egfr | ec |
| Hbegf | batch | mp | d10 | Egfr | ec |
| Hbegf | constitutive | ec | d10 | Egfr | ec |
| Hbegf | constitutive | per | d10 | Egfr | ec |
| Igf1 | batch | ec | d10 | Igfbp7 | ec |
| Igf1 | batch | fap | d10 | Igfbp7 | ec |
| Igf1 | batch | ic | d10 | Igfbp7 | ec |
| Igf1 | batch | ec | d10 | Igfbp2 | ec |
| Igf1 | batch | fap | d10 | Igfbp2 | ec |
| Igf1 | batch | ic | d10 | Igfbp2 | ec |
| Igf2 | batch | ec | d10 | Insr | ec |
| Igf2 | batch | fap | d10 | Insr | ec |
| Igf2 | batch | mp | d10 | Insr | ec |
| Igf2 | batch | ec | d10 | Vtn | ec |
| Igf2 | batch | fap | d10 | Vtn | ec |
| Igf2 | batch | mp | d10 | Vtn | ec |
| Kitl | batch | fap | d10 | Kit | ec |
| Kitl | constitutive | ec | d10 | Kit | ec |
| Kitl | constitutive | per | d10 | Kit | ec |
| Mmp13 | batch | ec | d10 | F2r | ec |
| Mmp13 | batch | fap | d10 | F2r | ec |
| Mmp13 | batch | mp | d10 | F2r | ec |
| Nov | batch | fap | d10 | Notch1 | ec |
| Nov | batch | per | d10 | Notch1 | ec |
| Nov | batch | fap | d10 | Itga5 | ec |
| Nov | batch | per | d10 | Itga5 | ec |
| Nov | batch | fap | d10 | Itgb3 | ec |
| Nov | batch | per | d10 | Itgb3 | ec |
| Ntf3 | batch | ec | d10 | Ntrk2 | ec |
| Ntf3 | batch | fap | d10 | Ntrk2 | ec |
| Ntf3 | batch | mp | d10 | Ntrk2 | ec |
| Ntf3 | constitutive | per | d10 | Ntrk2 | ec |
| Pgf | batch | fap | d10 | Flt1 | ec |
| Pgf | batch | mp | d10 | Flt1 | ec |
| Pgf | constitutive | per | d10 | Flt1 | ec |
| Pgf | batch | fap | d10 | Nrp2 | ec |
| Pgf | batch | mp | d10 | Nrp2 | ec |
| Pgf | constitutive | per | d10 | Nrp2 | ec |
| Pgf | batch | fap | d10 | Nrp1 | ec |
| Pgf | batch | mp | d10 | Nrp1 | ec |
| Pgf | constitutive | per | d10 | Nrp1 | ec |
| Sfrp1 | batch | fap | d10 | Fzd2 | ec |
| Sfrp1 | constitutive | per | d10 | Fzd2 | ec |
| Sfrp4 | batch | fap | d10 | Fzd2 | ec |
| Sfrp4 | batch | mp | d10 | Fzd2 | ec |
| Tgfb3 | batch | fap | d10 | Acvrl1 | ec |
| Tgfb3 | batch | mp | d10 | Acvrl1 | ec |
| Tslp | batch | fap | d10 | Crlf2 | ec |
| Tslp | constitutive | ec | d10 | Crlf2 | ec |
| Vegfa | batch | fap | d10 | Flt1 | ec |
| Vegfa | batch | ic | d10 | Flt1 | ec |
| Vegfa | batch | mp | d10 | Flt1 | ec |
| Vegfa | batch | fap | d10 | Kdr | ec |
| Vegfa | batch | ic | d10 | Kdr | ec |
| Vegfa | batch | mp | d10 | Kdr | ec |
| Vegfa | batch | fap | d10 | Nrp2 | ec |
| Vegfa | batch | ic | d10 | Nrp2 | ec |
| Vegfa | batch | mp | d10 | Nrp2 | ec |
| Vegfa | batch | fap | d10 | Nrp1 | ec |
| Vegfa | batch | ic | d10 | Nrp1 | ec |
| Vegfa | batch | mp | d10 | Nrp1 | ec |
| Vegfa | batch | fap | d10 | Ephb2 | ec |
| Vegfa | batch | ic | d10 | Ephb2 | ec |
| Vegfa | batch | mp | d10 | Ephb2 | ec |
| Vegfa | batch | fap | d10 | Vtn | ec |
| Vegfa | batch | ic | d10 | Vtn | ec |
| Vegfa | batch | mp | d10 | Vtn | ec |
| Vegfa | batch | fap | d10 | Itga9 | ec |
| Vegfa | batch | ic | d10 | Itga9 | ec |
| Vegfa | batch | mp | d10 | Itga9 | ec |
| Wnt2 | batch | fap | d10 | Sfrp1 | ec |
| Dll1 | batch | ec | d10 | Notch1 | ec |
| Fgf9 | batch | ec | d10 | Fgfr3 | ec |
| Il15 | batch | ec | d10 | Il2rg | ec |
| Il15 | batch | ic | d10 | Il2rg | ec |
| Il15 | batch | ec | d10 | Il15ra | ec |
| Il15 | batch | ic | d10 | Il15ra | ec |
| Tnfsf10 | batch | ec | d10 | Tnfrsf10b | ec |
| Vegfc | batch | ec | d10 | Flt4 | ec |
| Vegfc | batch | ec | d10 | Kdr | ec |
| Vegfc | batch | ec | d10 | Nrp2 | ec |
| Vegfc | batch | ec | d10 | Nrp1 | ec |
| Vegfc | batch | ec | d10 | Itga9 | ec |
| Clcf1 | batch | ic | d10 | Crlf1 | ec |
| Clcf1 | batch | mp | d10 | Crlf1 | ec |
| Ctf1 | batch | mp | d10 | Il6st | ec |
| Ctsg | batch | ic | d10 | F2rl3 | ec |
| Ctsg | batch | mp | d10 | F2rl3 | ec |
| Edn3 | batch | mp | d10 | Ednra | ec |
| Inhbb | batch | mp | d10 | Acvr2a | ec |
| Inhbb | constitutive | fap | d10 | Acvr2a | ec |
| Inhbb | constitutive | per | d10 | Acvr2a | ec |
| Jag1 | batch | ic | d10 | Notch1 | ec |
| Jag1 | batch | mp | d10 | Notch1 | ec |
| Jag1 | constitutive | per | d10 | Notch1 | ec |
| Jag2 | batch | mp | d10 | Notch1 | ec |
| Jag2 | constitutive | ec | d10 | Notch1 | ec |
| Nppc | batch | mp | d10 | Npr2 | ec |
| Serpini1 | batch | mp | d10 | Plat | ec |
| Serpini1 | constitutive | per | d10 | Plat | ec |
| Sfrp2 | batch | mp | d10 | Fzd2 | ec |
| Sfrp5 | batch | mp | d10 | Fzd2 | ec |
| Sfrp5 | batch | per | d10 | Fzd2 | ec |
| Tgfb2 | batch | mp | d10 | Tgfbr3 | ec |
| Tgfb2 | constitutive | ec | d10 | Tgfbr3 | ec |
| Tgfb2 | constitutive | fap | d10 | Tgfbr3 | ec |
| Tgfb2 | constitutive | per | d10 | Tgfbr3 | ec |
| Tgfb2 | batch | mp | d10 | Vtn | ec |
| Tgfb2 | constitutive | ec | d10 | Vtn | ec |
| Tgfb2 | constitutive | fap | d10 | Vtn | ec |
| Tgfb2 | constitutive | per | d10 | Vtn | ec |
| Tnfsf12 | batch | mp | d10 | Tnfrsf25 | ec |
| Tnfsf12 | total | ec | d10 | Tnfrsf25 | ec |
| Tnfsf12 | total | ic | d10 | Tnfrsf25 | ec |
| Tnfsf12 | total | per | d10 | Tnfrsf25 | ec |
| Tnfsf12 | batch | mp | d10 | Tnfrsf1a | ec |
| Tnfsf12 | total | ec | d10 | Tnfrsf1a | ec |
| Tnfsf12 | total | ic | d10 | Tnfrsf1a | ec |
| Tnfsf12 | total | per | d10 | Tnfrsf1a | ec |
| Tnfsf12 | batch | mp | d10 | Fas | ec |
| Tnfsf12 | total | ec | d10 | Fas | ec |
| Tnfsf12 | total | ic | d10 | Fas | ec |
| Tnfsf12 | total | per | d10 | Fas | ec |
| Vegfb | batch | mp | d10 | Flt1 | ec |
| Vegfb | batch | mp | d10 | Nrp1 | ec |
| Areg | batch | ic | d10 | Egfr | ec |
| Csf1 | batch | ic | d10 | H2-Bl | ec |
| Ifng | batch | ic | d10 | Ifngr1 | ec |
| Il10 | batch | ic | d10 | Il10ra | ec |
| Il1b | batch | ic | d10 | Adrb2 | ec |
| Il6 | batch | ic | d10 | Il6st | ec |
| Lif | batch | ic | d10 | Il6st | ec |
| Lif | constitutive | mp | d10 | Il6st | ec |
| Lif | batch | ic | d10 | Lifr | ec |
| Lif | constitutive | mp | d10 | Lifr | ec |
| Ltb | batch | ic | d10 | Tnfrsf1a | ec |
| Mmp9 | batch | ic | d10 | Flt1 | ec |
| Osm | batch | ic | d10 | Osmr | ec |
| Osm | batch | ic | d10 | Il6st | ec |
| Osm | batch | ic | d10 | Lifr | ec |
| Pdgfb | batch | ic | d10 | Pdgfrb | ec |
| Pdgfb | constitutive | ec | d10 | Pdgfrb | ec |
| Pf4 | batch | ic | d10 | Thbd | ec |
| Ppbp | batch | ic | d10 | Gabbr1 | ec |
| Tnf | batch | ic | d10 | Tnfrsf1a | ec |
| Tnf | batch | ic | d10 | Fas | ec |
| Adm | batch | per | d10 | Calcrl | ec |
| Angpt4 | batch | per | d10 | Tek | ec |
| Gdnf | batch | per | d10 | Ret | ec |
| Tgfb1 | total | ec | d10 | Eng | ec |
| Tgfb1 | total | ic | d10 | Eng | ec |
| Tgfb1 | total | ec | d10 | Acvrl1 | ec |
| Tgfb1 | total | ic | d10 | Acvrl1 | ec |
| Tgfb1 | total | ec | d10 | Tgfbr3 | ec |
| Tgfb1 | total | ic | d10 | Tgfbr3 | ec |
| Tgfb1 | total | ec | d10 | Vtn | ec |
| Tgfb1 | total | ic | d10 | Vtn | ec |
| Angpt1 | constitutive | fap | d10 | Tek | ec |
| Fgf7 | constitutive | fap | d10 | Fgfr3 | ec |
| Fgf7 | constitutive | fap | d10 | Nrp1 | ec |
| Bmp5 | constitutive | ec | d10 | Bmpr1a | ec |
| Bmp5 | constitutive | fap | d10 | Bmpr1a | ec |
| Bmp5 | constitutive | per | d10 | Bmpr1a | ec |
| Cmtm8 | constitutive | ec | d10 | Egfr | ec |
| Angpt2 | constitutive | ec | d10 | Tek | ec |
| Angpt2 | constitutive | ic | d10 | Tek | ec |
| Angpt2 | constitutive | per | d10 | Tek | ec |
| Ebi3 | constitutive | ic | d10 | Il27ra | ec |
| Nampt | constitutive | ic | d10 | Adora2a | ec |
| Bmp2 | batch | fap | d14 | Eng | ec |
| Bmp2 | constitutive | per | d14 | Eng | ec |
| Bmp2 | batch | fap | d14 | Bmpr1a | ec |
| Bmp2 | constitutive | per | d14 | Bmpr1a | ec |
| Bmp2 | batch | fap | d14 | Acvr2a | ec |
| Bmp2 | constitutive | per | d14 | Acvr2a | ec |
| Bmp2 | batch | fap | d14 | Bmpr2 | ec |
| Bmp2 | constitutive | per | d14 | Bmpr2 | ec |
| Bmp4 | batch | fap | d14 | Bmpr1a | ec |
| Bmp4 | constitutive | ec | d14 | Bmpr1a | ec |
| Bmp4 | constitutive | per | d14 | Bmpr1a | ec |
| Bmp4 | batch | fap | d14 | Bmpr2 | ec |
| Bmp4 | constitutive | ec | d14 | Bmpr2 | ec |
| Bmp4 | constitutive | per | d14 | Bmpr2 | ec |
| Bmp6 | batch | fap | d14 | Bmpr1a | ec |
| Bmp6 | constitutive | per | d14 | Bmpr1a | ec |
| Bmp6 | batch | fap | d14 | Acvr2a | ec |
| Bmp6 | constitutive | per | d14 | Acvr2a | ec |
| Bmp6 | batch | fap | d14 | Bmpr2 | ec |
| Bmp6 | constitutive | per | d14 | Bmpr2 | ec |
| Bmp7 | batch | fap | d14 | Eng | ec |
| Bmp7 | batch | fap | d14 | Bmpr1a | ec |
| Bmp7 | batch | fap | d14 | Acvr2a | ec |
| Bmp7 | batch | fap | d14 | Bmpr2 | ec |
| Clu | batch | ec | d14 | Vldlr | ec |
| Clu | batch | fap | d14 | Vldlr | ec |
| Clu | batch | ic | d14 | Vldlr | ec |
| Csf2 | batch | fap | d14 | Csf2rb | ec |
| Ctgf | batch | fap | d14 | Itga5 | ec |
| Ctgf | batch | ic | d14 | Itga5 | ec |
| Ctgf | constitutive | per | d14 | Itga5 | ec |
| Edn1 | batch | fap | d14 | Ednra | ec |
| Edn1 | constitutive | ec | d14 | Ednra | ec |
| Efnb1 | batch | fap | d14 | Ephb2 | ec |
| Efnb1 | batch | ic | d14 | Ephb2 | ec |
| Efnb1 | constitutive | ec | d14 | Ephb2 | ec |
| Efnb1 | constitutive | per | d14 | Ephb2 | ec |
| Fgf1 | batch | fap | d14 | Fgfr3 | ec |
| Fgf1 | constitutive | per | d14 | Fgfr3 | ec |
| Hbegf | batch | fap | d14 | Egfr | ec |
| Hbegf | constitutive | ec | d14 | Egfr | ec |
| Hbegf | constitutive | per | d14 | Egfr | ec |
| Igf1 | batch | ec | d14 | Igfbp5 | ec |
| Igf1 | batch | fap | d14 | Igfbp5 | ec |
| Igf1 | batch | ic | d14 | Igfbp5 | ec |
| Igf1 | batch | ec | d14 | Igfbp7 | ec |
| Igf1 | batch | fap | d14 | Igfbp7 | ec |
| Igf1 | batch | ic | d14 | Igfbp7 | ec |
| Igf1 | batch | ec | d14 | Igfbp2 | ec |
| Igf1 | batch | fap | d14 | Igfbp2 | ec |
| Igf1 | batch | ic | d14 | Igfbp2 | ec |
| Igf2 | batch | ec | d14 | Insr | ec |
| Igf2 | batch | fap | d14 | Insr | ec |
| Igf2 | batch | ic | d14 | Insr | ec |
| Igf2 | batch | ec | d14 | Vtn | ec |
| Igf2 | batch | fap | d14 | Vtn | ec |
| Igf2 | batch | ic | d14 | Vtn | ec |
| Kitl | batch | fap | d14 | Kit | ec |
| Kitl | batch | ic | d14 | Kit | ec |
| Kitl | constitutive | ec | d14 | Kit | ec |
| Kitl | constitutive | per | d14 | Kit | ec |
| Mmp13 | batch | fap | d14 | F2r | ec |
| Nov | batch | fap | d14 | Notch1 | ec |
| Nov | batch | fap | d14 | Itga5 | ec |
| Nov | batch | fap | d14 | Itgb3 | ec |
| Ntf3 | batch | ec | d14 | Ntrk2 | ec |
| Ntf3 | batch | fap | d14 | Ntrk2 | ec |
| Ntf3 | constitutive | per | d14 | Ntrk2 | ec |
| Pgf | batch | fap | d14 | Flt1 | ec |
| Pgf | constitutive | per | d14 | Flt1 | ec |
| Pgf | batch | fap | d14 | Nrp2 | ec |
| Pgf | constitutive | per | d14 | Nrp2 | ec |
| Pgf | batch | fap | d14 | Nrp1 | ec |
| Pgf | constitutive | per | d14 | Nrp1 | ec |
| Sfrp1 | batch | fap | d14 | Fzd2 | ec |
| Sfrp1 | constitutive | per | d14 | Fzd2 | ec |
| Sfrp4 | batch | fap | d14 | Fzd2 | ec |
| Tgfb3 | batch | fap | d14 | Acvrl1 | ec |
| Vegfa | batch | fap | d14 | Flt1 | ec |
| Vegfa | batch | fap | d14 | Kdr | ec |
| Vegfa | batch | fap | d14 | Nrp2 | ec |
| Vegfa | batch | fap | d14 | Nrp1 | ec |
| Vegfa | batch | fap | d14 | Ephb2 | ec |
| Vegfa | batch | fap | d14 | Vtn | ec |
| Vegfa | batch | fap | d14 | Itga9 | ec |
| Wnt11 | batch | fap | d14 | Fzd4 | ec |
| Wnt2 | batch | fap | d14 | Sfrp1 | ec |
| Dll1 | batch | ec | d14 | Notch1 | ec |
| Fgf9 | batch | ec | d14 | Fgfr3 | ec |
| Il15 | batch | ec | d14 | Il15ra | ec |
| Il15 | batch | ic | d14 | Il15ra | ec |
| Tnfsf10 | batch | ec | d14 | Tnfrsf10b | ec |
| Vegfc | batch | ec | d14 | Flt4 | ec |
| Vegfc | batch | ec | d14 | Kdr | ec |
| Vegfc | batch | ec | d14 | Nrp2 | ec |
| Vegfc | batch | ec | d14 | Nrp1 | ec |
| Vegfc | batch | ec | d14 | Itga9 | ec |
| Il10 | batch | ic | d14 | Il10ra | ec |
| Pdgfb | batch | ic | d14 | Pdgfrb | ec |
| Pdgfb | constitutive | ec | d14 | Pdgfrb | ec |
| Dll4 | total | ec | d14 | Notch1 | ec |
| Angpt1 | constitutive | fap | d14 | Tek | ec |
| Fgf7 | constitutive | fap | d14 | Fgfr3 | ec |
| Fgf7 | constitutive | fap | d14 | Nrp1 | ec |
| Tgfb2 | constitutive | ec | d14 | Tgfbr3 | ec |
| Tgfb2 | constitutive | fap | d14 | Tgfbr3 | ec |
| Tgfb2 | constitutive | per | d14 | Tgfbr3 | ec |
| Tgfb2 | constitutive | ec | d14 | Vtn | ec |
| Tgfb2 | constitutive | fap | d14 | Vtn | ec |
| Tgfb2 | constitutive | per | d14 | Vtn | ec |
| Bmp5 | constitutive | ec | d14 | Bmpr1a | ec |
| Bmp5 | constitutive | fap | d14 | Bmpr1a | ec |
| Bmp5 | constitutive | per | d14 | Bmpr1a | ec |
| Inhbb | constitutive | fap | d14 | Acvr2a | ec |
| Inhbb | constitutive | per | d14 | Acvr2a | ec |
| Jag2 | constitutive | ec | d14 | Notch1 | ec |
| Cmtm8 | constitutive | ec | d14 | Egfr | ec |
| Angpt2 | constitutive | ec | d14 | Tek | ec |
| Angpt2 | constitutive | ic | d14 | Tek | ec |
| Angpt2 | constitutive | per | d14 | Tek | ec |
| Lif | constitutive | mp | d14 | Il6st | ec |
| Lif | constitutive | mp | d14 | Lifr | ec |
| Ebi3 | constitutive | ic | d14 | Il27ra | ec |
| Nampt | constitutive | ic | d14 | Adora2a | ec |
| Serpini1 | constitutive | per | d14 | Plat | ec |
| Jag1 | constitutive | per | d14 | Notch1 | ec |
| Bmp2 | batch | fap | d0 | Bmpr1a | fap |
| Bmp2 | batch | mp | d0 | Bmpr1a | fap |
| Bmp2 | constitutive | per | d0 | Bmpr1a | fap |
| Bmp2 | batch | fap | d0 | Acvr2a | fap |
| Bmp2 | batch | mp | d0 | Acvr2a | fap |
| Bmp2 | constitutive | per | d0 | Acvr2a | fap |
| Bmp4 | batch | fap | d0 | Bmpr1a | fap |
| Bmp4 | batch | mp | d0 | Bmpr1a | fap |
| Bmp4 | constitutive | ec | d0 | Bmpr1a | fap |
| Bmp4 | constitutive | per | d0 | Bmpr1a | fap |
| Bmp6 | batch | ec | d0 | Bmpr1a | fap |
| Bmp6 | batch | fap | d0 | Bmpr1a | fap |
| Bmp6 | batch | mp | d0 | Bmpr1a | fap |
| Bmp6 | constitutive | per | d0 | Bmpr1a | fap |
| Bmp6 | batch | ec | d0 | Acvr2a | fap |
| Bmp6 | batch | fap | d0 | Acvr2a | fap |
| Bmp6 | batch | mp | d0 | Acvr2a | fap |
| Bmp6 | constitutive | per | d0 | Acvr2a | fap |
| Bmp7 | batch | fap | d0 | Bmpr1a | fap |
| Bmp7 | batch | fap | d0 | Acvr2a | fap |
| Bmp7 | batch | fap | d0 | Acvr1 | fap |
| Ccl11 | batch | fap | d0 | Dpp4 | fap |
| Ccl11 | batch | mp | d0 | Dpp4 | fap |
| Ccl11 | batch | per | d0 | Dpp4 | fap |
| Clu | batch | ec | d0 | Vldlr | fap |
| Clu | batch | fap | d0 | Vldlr | fap |
| Clu | batch | ic | d0 | Vldlr | fap |
| Clu | batch | mp | d0 | Vldlr | fap |
| Crlf1 | batch | ec | d0 | Ctf1 | fap |
| Crlf1 | batch | fap | d0 | Ctf1 | fap |
| Crlf1 | batch | mp | d0 | Ctf1 | fap |
| Cst3 | batch | fap | d0 | Tgfbr2 | fap |
| Cst3 | constitutive | ic | d0 | Tgfbr2 | fap |
| Cst3 | constitutive | per | d0 | Tgfbr2 | fap |
| Efnb1 | batch | fap | d0 | Ephb2 | fap |
| Efnb1 | batch | ic | d0 | Ephb2 | fap |
| Efnb1 | constitutive | ec | d0 | Ephb2 | fap |
| Efnb1 | constitutive | per | d0 | Ephb2 | fap |
| Fgf1 | batch | fap | d0 | Fgfr2 | fap |
| Fgf1 | constitutive | per | d0 | Fgfr2 | fap |
| Fgf1 | batch | fap | d0 | Fgfr1 | fap |
| Fgf1 | constitutive | per | d0 | Fgfr1 | fap |
| Gas6 | batch | fap | d0 | Axl | fap |
| Gas6 | batch | mp | d0 | Axl | fap |
| Igf1 | batch | ec | d0 | Igfbp3 | fap |
| Igf1 | batch | fap | d0 | Igfbp3 | fap |
| Igf1 | batch | ic | d0 | Igfbp3 | fap |
| Igf1 | batch | ec | d0 | Igfbp5 | fap |
| Igf1 | batch | fap | d0 | Igfbp5 | fap |
| Igf1 | batch | ic | d0 | Igfbp5 | fap |
| Igf1 | batch | ec | d0 | Igfbp6 | fap |
| Igf1 | batch | fap | d0 | Igfbp6 | fap |
| Igf1 | batch | ic | d0 | Igfbp6 | fap |
| Igf1 | batch | ec | d0 | Igfbp2 | fap |
| Igf1 | batch | fap | d0 | Igfbp2 | fap |
| Igf1 | batch | ic | d0 | Igfbp2 | fap |
| Il18 | batch | fap | d0 | Il1rl2 | fap |
| Il18 | constitutive | ic | d0 | Il1rl2 | fap |
| Il18 | constitutive | mp | d0 | Il1rl2 | fap |
| Nov | batch | ec | d0 | Itgb3 | fap |
| Nov | batch | fap | d0 | Itgb3 | fap |
| Nov | batch | per | d0 | Itgb3 | fap |
| Nov | batch | ec | d0 | Itgav | fap |
| Nov | batch | fap | d0 | Itgav | fap |
| Nov | batch | per | d0 | Itgav | fap |
| Ntf3 | batch | ec | d0 | Ntrk2 | fap |
| Ntf3 | batch | fap | d0 | Ntrk2 | fap |
| Ntf3 | batch | mp | d0 | Ntrk2 | fap |
| Ntf3 | constitutive | per | d0 | Ntrk2 | fap |
| Ntf3 | batch | ec | d0 | Ngfr | fap |
| Ntf3 | batch | fap | d0 | Ngfr | fap |
| Ntf3 | batch | mp | d0 | Ngfr | fap |
| Ntf3 | constitutive | per | d0 | Ngfr | fap |
| Pdgfc | batch | fap | d0 | Pdgfra | fap |
| Pdgfc | constitutive | ic | d0 | Pdgfra | fap |
| Pdgfc | constitutive | per | d0 | Pdgfra | fap |
| Pgf | batch | fap | d0 | Nrp1 | fap |
| Pgf | constitutive | per | d0 | Nrp1 | fap |
| Ptn | batch | fap | d0 | Ptprs | fap |
| Ptn | constitutive | per | d0 | Ptprs | fap |
| S100b | batch | fap | d0 | Fgfr1 | fap |
| S100b | batch | mp | d0 | Fgfr1 | fap |
| S100b | constitutive | per | d0 | Fgfr1 | fap |
| Sfrp4 | batch | ec | d0 | Fzd2 | fap |
| Sfrp4 | batch | fap | d0 | Fzd2 | fap |
| Sfrp4 | batch | mp | d0 | Fzd2 | fap |
| Tslp | batch | fap | d0 | Crlf2 | fap |
| Tslp | constitutive | ec | d0 | Crlf2 | fap |
| Vegfa | batch | fap | d0 | Nrp1 | fap |
| Vegfa | batch | mp | d0 | Nrp1 | fap |
| Vegfa | batch | fap | d0 | Ephb2 | fap |
| Vegfa | batch | mp | d0 | Ephb2 | fap |
| Vegfa | batch | fap | d0 | Itga9 | fap |
| Vegfa | batch | mp | d0 | Itga9 | fap |
| Wnt11 | batch | fap | d0 | Fzd4 | fap |
| Wnt11 | batch | ic | d0 | Fzd4 | fap |
| Wnt2 | batch | fap | d0 | Sfrp1 | fap |
| Wnt2 | batch | fap | d0 | Fzd1 | fap |
| Cxcl10 | batch | ec | d0 | Dpp4 | fap |
| Cxcl9 | batch | ec | d0 | Dpp4 | fap |
| Cxcl9 | batch | ic | d0 | Dpp4 | fap |
| Dll1 | batch | ec | d0 | Notch2 | fap |
| Fgf9 | batch | ec | d0 | Fgfr2 | fap |
| Il15 | batch | ec | d0 | Il15ra | fap |
| Il15 | batch | ic | d0 | Il15ra | fap |
| Sfrp5 | batch | ec | d0 | Fzd2 | fap |
| Sfrp5 | batch | mp | d0 | Fzd2 | fap |
| Sfrp5 | batch | per | d0 | Fzd2 | fap |
| Vegfc | batch | ec | d0 | Nrp1 | fap |
| Vegfc | batch | mp | d0 | Nrp1 | fap |
| Vegfc | batch | ec | d0 | Itga9 | fap |
| Vegfc | batch | mp | d0 | Itga9 | fap |
| Areg | batch | mp | d0 | Egfr | fap |
| Clcf1 | batch | ic | d0 | Crlf1 | fap |
| Clcf1 | batch | mp | d0 | Crlf1 | fap |
| Clcf1 | batch | ic | d0 | Cntfr | fap |
| Clcf1 | batch | mp | d0 | Cntfr | fap |
| Clcf1 | batch | ic | d0 | Esr1 | fap |
| Clcf1 | batch | mp | d0 | Esr1 | fap |
| Ctf1 | batch | mp | d0 | Il6st | fap |
| Cxcl12 | batch | ic | d0 | Cxcr4 | fap |
| Cxcl12 | batch | mp | d0 | Cxcr4 | fap |
| Cxcl12 | constitutive | ec | d0 | Cxcr4 | fap |
| Cxcl12 | constitutive | per | d0 | Cxcr4 | fap |
| Cxcl12 | batch | ic | d0 | Dpp4 | fap |
| Cxcl12 | batch | mp | d0 | Dpp4 | fap |
| Cxcl12 | constitutive | ec | d0 | Dpp4 | fap |
| Cxcl12 | constitutive | per | d0 | Dpp4 | fap |
| Gdnf | batch | mp | d0 | Ret | fap |
| Gdnf | batch | per | d0 | Ret | fap |
| Gdnf | batch | mp | d0 | Gfra2 | fap |
| Gdnf | batch | per | d0 | Gfra2 | fap |
| Gmfb | batch | mp | d0 | Egfr | fap |
| Hbegf | batch | mp | d0 | Egfr | fap |
| Hbegf | constitutive | ec | d0 | Egfr | fap |
| Hbegf | constitutive | per | d0 | Egfr | fap |
| Inhbb | batch | mp | d0 | Acvr2a | fap |
| Inhbb | constitutive | fap | d0 | Acvr2a | fap |
| Inhbb | constitutive | per | d0 | Acvr2a | fap |
| Inhbb | batch | mp | d0 | Acvr1 | fap |
| Inhbb | constitutive | fap | d0 | Acvr1 | fap |
| Inhbb | constitutive | per | d0 | Acvr1 | fap |
| Jag1 | batch | mp | d0 | Notch2 | fap |
| Jag1 | constitutive | per | d0 | Notch2 | fap |
| Jag2 | batch | mp | d0 | Notch2 | fap |
| Jag2 | constitutive | ec | d0 | Notch2 | fap |
| Ngf | batch | mp | d0 | Ngfr | fap |
| Ngf | constitutive | per | d0 | Ngfr | fap |
| Nppc | batch | mp | d0 | Npr2 | fap |
| Serpini1 | batch | mp | d0 | Plat | fap |
| Serpini1 | constitutive | per | d0 | Plat | fap |
| Tgfb3 | batch | mp | d0 | Acvrl1 | fap |
| Tgfb3 | batch | mp | d0 | Tgfbr2 | fap |
| Tgfb3 | batch | mp | d0 | Itgav | fap |
| Tnfsf12 | batch | mp | d0 | Tnfrsf12a | fap |
| Tnfsf12 | batch | mp | d0 | Tnfrsf1a | fap |
| Ccl22 | batch | ic | d0 | Dpp4 | fap |
| Ccl5 | batch | ic | d0 | Sdc4 | fap |
| Ifng | batch | ic | d0 | Ifngr1 | fap |
| Il6 | batch | ic | d0 | Il6st | fap |
| Ltb | batch | ic | d0 | Tnfrsf1a | fap |
| Pdgfb | batch | ic | d0 | Pdgfra | fap |
| Pdgfb | constitutive | ec | d0 | Pdgfra | fap |
| Pdgfb | batch | ic | d0 | Pdgfrb | fap |
| Pdgfb | constitutive | ec | d0 | Pdgfrb | fap |
| Adm | batch | per | d0 | Calcrl | fap |
| Angpt4 | batch | per | d0 | Tek | fap |
| Angpt1 | constitutive | fap | d0 | Tek | fap |
| Fgf7 | constitutive | fap | d0 | Fgfr2 | fap |
| Fgf7 | constitutive | fap | d0 | Nrp1 | fap |
| Tgfb2 | constitutive | ec | d0 | Tgfbr3 | fap |
| Tgfb2 | constitutive | fap | d0 | Tgfbr3 | fap |
| Tgfb2 | constitutive | per | d0 | Tgfbr3 | fap |
| Tgfb2 | constitutive | ec | d0 | Tgfbr2 | fap |
| Tgfb2 | constitutive | fap | d0 | Tgfbr2 | fap |
| Tgfb2 | constitutive | per | d0 | Tgfbr2 | fap |
| Bmp5 | constitutive | ec | d0 | Bmpr1a | fap |
| Bmp5 | constitutive | fap | d0 | Bmpr1a | fap |
| Bmp5 | constitutive | per | d0 | Bmpr1a | fap |
| Cmtm8 | constitutive | ec | d0 | Egfr | fap |
| Angpt2 | constitutive | ec | d0 | Tek | fap |
| Angpt2 | constitutive | ic | d0 | Tek | fap |
| Angpt2 | constitutive | per | d0 | Tek | fap |
| Pdgfa | constitutive | ec | d0 | Pdgfra | fap |
| Pdgfa | constitutive | mp | d0 | Pdgfra | fap |
| Pdgfa | constitutive | per | d0 | Pdgfra | fap |
| Lif | constitutive | mp | d0 | Il6st | fap |
| Sfrp1 | constitutive | per | d0 | Fzd2 | fap |
| Bmp2 | batch | ec | d1 | Bmpr1a | fap |
| Bmp2 | batch | fap | d1 | Bmpr1a | fap |
| Bmp2 | batch | mp | d1 | Bmpr1a | fap |
| Bmp2 | constitutive | per | d1 | Bmpr1a | fap |
| Bmp2 | batch | ec | d1 | Acvr2a | fap |
| Bmp2 | batch | fap | d1 | Acvr2a | fap |
| Bmp2 | batch | mp | d1 | Acvr2a | fap |
| Bmp2 | constitutive | per | d1 | Acvr2a | fap |
| Bmp6 | batch | fap | d1 | Bmpr1a | fap |
| Bmp6 | constitutive | per | d1 | Bmpr1a | fap |
| Bmp6 | batch | fap | d1 | Acvr2a | fap |
| Bmp6 | constitutive | per | d1 | Acvr2a | fap |
| Cxcl2 | batch | ec | d1 | Dpp4 | fap |
| Cxcl2 | batch | fap | d1 | Dpp4 | fap |
| Cxcl2 | batch | ic | d1 | Dpp4 | fap |
| Cxcl2 | batch | per | d1 | Dpp4 | fap |
| Efnb1 | batch | fap | d1 | Ephb2 | fap |
| Efnb1 | batch | ic | d1 | Ephb2 | fap |
| Efnb1 | constitutive | ec | d1 | Ephb2 | fap |
| Efnb1 | constitutive | per | d1 | Ephb2 | fap |
| Il18 | batch | fap | d1 | Il1rl2 | fap |
| Il18 | constitutive | ic | d1 | Il1rl2 | fap |
| Il18 | constitutive | mp | d1 | Il1rl2 | fap |
| Il6 | batch | ec | d1 | Il6st | fap |
| Il6 | batch | fap | d1 | Il6st | fap |
| Il6 | batch | per | d1 | Il6st | fap |
| Lif | batch | fap | d1 | Il6st | fap |
| Lif | batch | ic | d1 | Il6st | fap |
| Lif | batch | per | d1 | Il6st | fap |
| Lif | constitutive | mp | d1 | Il6st | fap |
| Pdgfc | batch | fap | d1 | Pdgfra | fap |
| Pdgfc | constitutive | ic | d1 | Pdgfra | fap |
| Pdgfc | constitutive | per | d1 | Pdgfra | fap |
| Pf4 | batch | fap | d1 | Ldlr | fap |
| Pf4 | batch | ic | d1 | Ldlr | fap |
| Pf4 | batch | per | d1 | Ldlr | fap |
| Pf4 | batch | fap | d1 | Thbd | fap |
| Pf4 | batch | ic | d1 | Thbd | fap |
| Pf4 | batch | per | d1 | Thbd | fap |
| Tslp | batch | fap | d1 | Crlf2 | fap |
| Tslp | constitutive | ec | d1 | Crlf2 | fap |
| Wnt2 | batch | fap | d1 | Sfrp1 | fap |
| Wnt2 | batch | fap | d1 | Fzd1 | fap |
| Il1b | batch | ec | d1 | Il1r1 | fap |
| Il1b | batch | ic | d1 | Il1r1 | fap |
| Il1b | batch | per | d1 | Il1r1 | fap |
| Il1b | batch | ec | d1 | Il1r2 | fap |
| Il1b | batch | ic | d1 | Il1r2 | fap |
| Il1b | batch | per | d1 | Il1r2 | fap |
| Inhbb | batch | ec | d1 | Acvr2a | fap |
| Inhbb | constitutive | fap | d1 | Acvr2a | fap |
| Inhbb | constitutive | per | d1 | Acvr2a | fap |
| Inhbb | batch | ec | d1 | Acvr1 | fap |
| Inhbb | constitutive | fap | d1 | Acvr1 | fap |
| Inhbb | constitutive | per | d1 | Acvr1 | fap |
| Pgf | batch | ec | d1 | Nrp1 | fap |
| Pgf | constitutive | per | d1 | Nrp1 | fap |
| Areg | batch | ic | d1 | Egfr | fap |
| Areg | batch | mp | d1 | Egfr | fap |
| Bmp4 | batch | mp | d1 | Bmpr1a | fap |
| Bmp4 | constitutive | ec | d1 | Bmpr1a | fap |
| Bmp4 | constitutive | per | d1 | Bmpr1a | fap |
| Ccl11 | batch | mp | d1 | Dpp4 | fap |
| Ccl11 | batch | per | d1 | Dpp4 | fap |
| Clcf1 | batch | mp | d1 | Crlf1 | fap |
| Clcf1 | batch | mp | d1 | Cntfr | fap |
| Clcf1 | batch | mp | d1 | Esr1 | fap |
| Ctgf | batch | ic | d1 | Itga5 | fap |
| Ctgf | batch | mp | d1 | Itga5 | fap |
| Ctgf | constitutive | per | d1 | Itga5 | fap |
| Cxcl12 | batch | ic | d1 | Dpp4 | fap |
| Cxcl12 | batch | mp | d1 | Dpp4 | fap |
| Cxcl12 | constitutive | ec | d1 | Dpp4 | fap |
| Cxcl12 | constitutive | per | d1 | Dpp4 | fap |
| Gas6 | batch | mp | d1 | Axl | fap |
| Gdnf | batch | mp | d1 | Gfra2 | fap |
| Gdnf | batch | per | d1 | Gfra2 | fap |
| Gmfb | batch | mp | d1 | Egfr | fap |
| Hbegf | batch | mp | d1 | Cd44 | fap |
| Hbegf | constitutive | ec | d1 | Cd44 | fap |
| Hbegf | constitutive | per | d1 | Cd44 | fap |
| Hbegf | batch | mp | d1 | Egfr | fap |
| Hbegf | constitutive | ec | d1 | Egfr | fap |
| Hbegf | constitutive | per | d1 | Egfr | fap |
| Il33 | batch | mp | d1 | Il1rl1 | fap |
| Jag2 | batch | mp | d1 | Notch2 | fap |
| Jag2 | constitutive | ec | d1 | Notch2 | fap |
| Ngf | batch | mp | d1 | Ngfr | fap |
| Ngf | constitutive | per | d1 | Ngfr | fap |
| Nppc | batch | mp | d1 | Npr2 | fap |
| Pdap1 | batch | mp | d1 | Pdgfa | fap |
| S100b | batch | mp | d1 | Fgfr1 | fap |
| S100b | constitutive | per | d1 | Fgfr1 | fap |
| Serpini1 | batch | mp | d1 | Plat | fap |
| Serpini1 | constitutive | per | d1 | Plat | fap |
| Sfrp4 | batch | mp | d1 | Fzd2 | fap |
| Sfrp5 | batch | mp | d1 | Fzd2 | fap |
| Spp1 | batch | mp | d1 | Itga5 | fap |
| Spp1 | batch | per | d1 | Itga5 | fap |
| Spp1 | batch | mp | d1 | Itgb5 | fap |
| Spp1 | batch | per | d1 | Itgb5 | fap |
| Spp1 | batch | mp | d1 | Itgb3 | fap |
| Spp1 | batch | per | d1 | Itgb3 | fap |
| Spp1 | batch | mp | d1 | Itgav | fap |
| Spp1 | batch | per | d1 | Itgav | fap |
| Spp1 | batch | mp | d1 | Itga9 | fap |
| Spp1 | batch | per | d1 | Itga9 | fap |
| Vegfa | batch | ic | d1 | Nrp1 | fap |
| Vegfa | batch | mp | d1 | Nrp1 | fap |
| Vegfa | batch | ic | d1 | Ephb2 | fap |
| Vegfa | batch | mp | d1 | Ephb2 | fap |
| Vegfa | batch | ic | d1 | Itga9 | fap |
| Vegfa | batch | mp | d1 | Itga9 | fap |
| Vegfc | batch | mp | d1 | Nrp1 | fap |
| Vegfc | batch | mp | d1 | Itga9 | fap |
| Cxcl10 | batch | ic | d1 | Dpp4 | fap |
| Il1a | batch | ic | d1 | Il1r1 | fap |
| Il1a | batch | ic | d1 | Il1r2 | fap |
| Jag1 | batch | ic | d1 | Notch2 | fap |
| Jag1 | constitutive | per | d1 | Notch2 | fap |
| Osm | batch | ic | d1 | Osmr | fap |
| Osm | batch | per | d1 | Osmr | fap |
| Osm | batch | ic | d1 | Il6st | fap |
| Osm | batch | per | d1 | Il6st | fap |
| Ppbp | batch | ic | d1 | Slc1a5 | fap |
| Ppbp | batch | ic | d1 | Itgb5 | fap |
| Tnf | batch | ic | d1 | Tnfrsf1a | fap |
| Tnf | batch | per | d1 | Tnfrsf1a | fap |
| Btc | batch | per | d1 | Egfr | fap |
| Il11 | batch | per | d1 | Il11ra1 | fap |
| Il11 | batch | per | d1 | Il6st | fap |
| Nov | batch | per | d1 | Itga5 | fap |
| Nov | batch | per | d1 | Itgb3 | fap |
| Nov | batch | per | d1 | Itgav | fap |
| Sfrp2 | batch | per | d1 | Fzd2 | fap |
| Fgf7 | constitutive | fap | d1 | Nrp1 | fap |
| Tgfb2 | constitutive | ec | d1 | Tgfbr3 | fap |
| Tgfb2 | constitutive | fap | d1 | Tgfbr3 | fap |
| Tgfb2 | constitutive | per | d1 | Tgfbr3 | fap |
| Tgfb2 | constitutive | ec | d1 | Tgfbr2 | fap |
| Tgfb2 | constitutive | fap | d1 | Tgfbr2 | fap |
| Tgfb2 | constitutive | per | d1 | Tgfbr2 | fap |
| Bmp5 | constitutive | ec | d1 | Bmpr1a | fap |
| Bmp5 | constitutive | fap | d1 | Bmpr1a | fap |
| Bmp5 | constitutive | per | d1 | Bmpr1a | fap |
| Pdgfb | constitutive | ec | d1 | Pdgfra | fap |
| Pdgfb | constitutive | ec | d1 | Pdgfrb | fap |
| Cmtm8 | constitutive | ec | d1 | Egfr | fap |
| Pdgfa | constitutive | ec | d1 | Pdgfra | fap |
| Pdgfa | constitutive | mp | d1 | Pdgfra | fap |
| Pdgfa | constitutive | per | d1 | Pdgfra | fap |
| Cst3 | constitutive | ic | d1 | Tgfbr2 | fap |
| Cst3 | constitutive | per | d1 | Tgfbr2 | fap |
| Fgf1 | constitutive | per | d1 | Fgfr1 | fap |
| Ptn | constitutive | per | d1 | Ptprs | fap |
| Sfrp1 | constitutive | per | d1 | Fzd2 | fap |
| Ntf3 | constitutive | per | d1 | Ngfr | fap |
| Bmp2 | batch | ec | d2 | Bmpr1a | fap |
| Bmp2 | batch | fap | d2 | Bmpr1a | fap |
| Bmp2 | batch | mp | d2 | Bmpr1a | fap |
| Bmp2 | constitutive | per | d2 | Bmpr1a | fap |
| Bmp2 | batch | ec | d2 | Acvr2a | fap |
| Bmp2 | batch | fap | d2 | Acvr2a | fap |
| Bmp2 | batch | mp | d2 | Acvr2a | fap |
| Bmp2 | constitutive | per | d2 | Acvr2a | fap |
| Bmp6 | batch | fap | d2 | Bmpr1a | fap |
| Bmp6 | constitutive | per | d2 | Bmpr1a | fap |
| Bmp6 | batch | fap | d2 | Acvr2a | fap |
| Bmp6 | constitutive | per | d2 | Acvr2a | fap |
| Ccl5 | batch | ec | d2 | Sdc4 | fap |
| Ccl5 | batch | fap | d2 | Sdc4 | fap |
| Csf1 | batch | fap | d2 | Csf1r | fap |
| Csf1 | batch | ic | d2 | Csf1r | fap |
| Csf1 | batch | mp | d2 | Csf1r | fap |
| Csf2 | batch | fap | d2 | Il3ra | fap |
| Cxcl10 | batch | ec | d2 | Dpp4 | fap |
| Cxcl10 | batch | fap | d2 | Dpp4 | fap |
| Cxcl10 | batch | ic | d2 | Dpp4 | fap |
| Cxcl10 | batch | per | d2 | Dpp4 | fap |
| Cxcl12 | batch | fap | d2 | Dpp4 | fap |
| Cxcl12 | batch | mp | d2 | Dpp4 | fap |
| Cxcl12 | constitutive | ec | d2 | Dpp4 | fap |
| Cxcl12 | constitutive | per | d2 | Dpp4 | fap |
| Cxcl2 | batch | ec | d2 | Dpp4 | fap |
| Cxcl2 | batch | fap | d2 | Dpp4 | fap |
| Cxcl2 | batch | ic | d2 | Dpp4 | fap |
| Cxcl2 | batch | mp | d2 | Dpp4 | fap |
| Cxcl2 | batch | per | d2 | Dpp4 | fap |
| Cxcl9 | batch | ec | d2 | Dpp4 | fap |
| Cxcl9 | batch | fap | d2 | Dpp4 | fap |
| Cxcl9 | batch | ic | d2 | Dpp4 | fap |
| Efnb1 | batch | fap | d2 | Ephb2 | fap |
| Efnb1 | constitutive | ec | d2 | Ephb2 | fap |
| Efnb1 | constitutive | per | d2 | Ephb2 | fap |
| Il18 | batch | fap | d2 | Il1rl2 | fap |
| Il18 | constitutive | ic | d2 | Il1rl2 | fap |
| Il18 | constitutive | mp | d2 | Il1rl2 | fap |
| Il1b | batch | ec | d2 | Il1r1 | fap |
| Il1b | batch | fap | d2 | Il1r1 | fap |
| Il1b | batch | ic | d2 | Il1r1 | fap |
| Il1b | batch | mp | d2 | Il1r1 | fap |
| Il1b | batch | per | d2 | Il1r1 | fap |
| Il1b | batch | ec | d2 | Il1r2 | fap |
| Il1b | batch | fap | d2 | Il1r2 | fap |
| Il1b | batch | ic | d2 | Il1r2 | fap |
| Il1b | batch | mp | d2 | Il1r2 | fap |
| Il1b | batch | per | d2 | Il1r2 | fap |
| Il33 | batch | fap | d2 | Il1rl1 | fap |
| Il33 | batch | mp | d2 | Il1rl1 | fap |
| Il6 | batch | ec | d2 | Il6st | fap |
| Il6 | batch | fap | d2 | Il6st | fap |
| Il6 | batch | per | d2 | Il6st | fap |
| Lif | batch | fap | d2 | Il6st | fap |
| Lif | batch | ic | d2 | Il6st | fap |
| Lif | batch | per | d2 | Il6st | fap |
| Lif | constitutive | mp | d2 | Il6st | fap |
| Pdgfc | batch | fap | d2 | Pdgfra | fap |
| Pdgfc | constitutive | ic | d2 | Pdgfra | fap |
| Pdgfc | constitutive | per | d2 | Pdgfra | fap |
| Pf4 | batch | ec | d2 | Ldlr | fap |
| Pf4 | batch | fap | d2 | Ldlr | fap |
| Pf4 | batch | ic | d2 | Ldlr | fap |
| Pf4 | batch | mp | d2 | Ldlr | fap |
| Pf4 | batch | per | d2 | Ldlr | fap |
| Pf4 | batch | ec | d2 | Thbd | fap |
| Pf4 | batch | fap | d2 | Thbd | fap |
| Pf4 | batch | ic | d2 | Thbd | fap |
| Pf4 | batch | mp | d2 | Thbd | fap |
| Pf4 | batch | per | d2 | Thbd | fap |
| Sfrp2 | batch | ec | d2 | Fzd2 | fap |
| Sfrp2 | batch | fap | d2 | Fzd2 | fap |
| Sfrp2 | batch | per | d2 | Fzd2 | fap |
| Spp1 | batch | ec | d2 | Itga5 | fap |
| Spp1 | batch | fap | d2 | Itga5 | fap |
| Spp1 | batch | ic | d2 | Itga5 | fap |
| Spp1 | batch | mp | d2 | Itga5 | fap |
| Spp1 | batch | per | d2 | Itga5 | fap |
| Spp1 | batch | ec | d2 | Itgb5 | fap |
| Spp1 | batch | fap | d2 | Itgb5 | fap |
| Spp1 | batch | ic | d2 | Itgb5 | fap |
| Spp1 | batch | mp | d2 | Itgb5 | fap |
| Spp1 | batch | per | d2 | Itgb5 | fap |
| Spp1 | batch | ec | d2 | Itgb3 | fap |
| Spp1 | batch | fap | d2 | Itgb3 | fap |
| Spp1 | batch | ic | d2 | Itgb3 | fap |
| Spp1 | batch | mp | d2 | Itgb3 | fap |
| Spp1 | batch | per | d2 | Itgb3 | fap |
| Spp1 | batch | ec | d2 | Itgav | fap |
| Spp1 | batch | fap | d2 | Itgav | fap |
| Spp1 | batch | ic | d2 | Itgav | fap |
| Spp1 | batch | mp | d2 | Itgav | fap |
| Spp1 | batch | per | d2 | Itgav | fap |
| Spp1 | batch | ec | d2 | Itga9 | fap |
| Spp1 | batch | fap | d2 | Itga9 | fap |
| Spp1 | batch | ic | d2 | Itga9 | fap |
| Spp1 | batch | mp | d2 | Itga9 | fap |
| Spp1 | batch | per | d2 | Itga9 | fap |
| Tslp | batch | fap | d2 | Crlf2 | fap |
| Tslp | constitutive | ec | d2 | Crlf2 | fap |
| Wnt2 | batch | fap | d2 | Sfrp1 | fap |
| Wnt2 | batch | fap | d2 | Fzd1 | fap |
| Igf1 | batch | ec | d2 | Igfbp2 | fap |
| Igf1 | batch | ic | d2 | Igfbp2 | fap |
| Igf1 | batch | mp | d2 | Igfbp2 | fap |
| Il15 | batch | ec | d2 | Il2rg | fap |
| Il15 | batch | ic | d2 | Il2rg | fap |
| Il15 | batch | ec | d2 | Il17ra | fap |
| Il15 | batch | ic | d2 | Il17ra | fap |
| Il15 | batch | ec | d2 | Il15ra | fap |
| Il15 | batch | ic | d2 | Il15ra | fap |
| Inhbb | batch | ec | d2 | Acvr2a | fap |
| Inhbb | constitutive | fap | d2 | Acvr2a | fap |
| Inhbb | constitutive | per | d2 | Acvr2a | fap |
| Inhbb | batch | ec | d2 | Acvr1 | fap |
| Inhbb | constitutive | fap | d2 | Acvr1 | fap |
| Inhbb | constitutive | per | d2 | Acvr1 | fap |
| Lgals3 | batch | ec | d2 | Lgals3bp | fap |
| Lgals3 | batch | ic | d2 | Lgals3bp | fap |
| Lgals3 | batch | mp | d2 | Lgals3bp | fap |
| Lgals3 | batch | per | d2 | Lgals3bp | fap |
| Mmp12 | batch | ec | d2 | Plaur | fap |
| Mmp12 | batch | ic | d2 | Plaur | fap |
| Mmp12 | batch | mp | d2 | Plaur | fap |
| Mmp12 | batch | per | d2 | Plaur | fap |
| Mmp13 | batch | ec | d2 | F2r | fap |
| Mmp13 | batch | ic | d2 | F2r | fap |
| Mmp13 | batch | per | d2 | F2r | fap |
| Osm | batch | ec | d2 | Osmr | fap |
| Osm | batch | ic | d2 | Osmr | fap |
| Osm | batch | mp | d2 | Osmr | fap |
| Osm | batch | per | d2 | Osmr | fap |
| Osm | batch | ec | d2 | Il6st | fap |
| Osm | batch | ic | d2 | Il6st | fap |
| Osm | batch | mp | d2 | Il6st | fap |
| Osm | batch | per | d2 | Il6st | fap |
| Pgf | batch | ec | d2 | Nrp1 | fap |
| Pgf | constitutive | per | d2 | Nrp1 | fap |
| Sfrp1 | batch | ec | d2 | Fzd2 | fap |
| Sfrp1 | constitutive | per | d2 | Fzd2 | fap |
| Tnfsf10 | batch | ec | d2 | Tnfrsf10b | fap |
| Areg | batch | ic | d2 | Egfr | fap |
| Areg | batch | mp | d2 | Egfr | fap |
| Bmp4 | batch | mp | d2 | Bmpr1a | fap |
| Bmp4 | constitutive | ec | d2 | Bmpr1a | fap |
| Bmp4 | constitutive | per | d2 | Bmpr1a | fap |
| Ccl11 | batch | mp | d2 | Dpp4 | fap |
| Ccl11 | batch | per | d2 | Dpp4 | fap |
| Clcf1 | batch | mp | d2 | Crlf1 | fap |
| Clcf1 | batch | mp | d2 | Cntfr | fap |
| Clcf1 | batch | mp | d2 | Esr1 | fap |
| Cst3 | batch | mp | d2 | Tgfbr2 | fap |
| Cst3 | constitutive | ic | d2 | Tgfbr2 | fap |
| Cst3 | constitutive | per | d2 | Tgfbr2 | fap |
| Ctgf | batch | mp | d2 | Itga5 | fap |
| Ctgf | constitutive | per | d2 | Itga5 | fap |
| Gas6 | batch | mp | d2 | Axl | fap |
| Gdnf | batch | mp | d2 | Gfra2 | fap |
| Gdnf | batch | per | d2 | Gfra2 | fap |
| Gmfb | batch | mp | d2 | Egfr | fap |
| Hbegf | batch | mp | d2 | Cd44 | fap |
| Hbegf | constitutive | ec | d2 | Cd44 | fap |
| Hbegf | constitutive | per | d2 | Cd44 | fap |
| Hbegf | batch | mp | d2 | Egfr | fap |
| Hbegf | constitutive | ec | d2 | Egfr | fap |
| Hbegf | constitutive | per | d2 | Egfr | fap |
| Jag2 | batch | mp | d2 | Notch1 | fap |
| Jag2 | constitutive | ec | d2 | Notch1 | fap |
| Jag2 | batch | mp | d2 | Notch3 | fap |
| Jag2 | constitutive | ec | d2 | Notch3 | fap |
| Jag2 | batch | mp | d2 | Notch2 | fap |
| Jag2 | constitutive | ec | d2 | Notch2 | fap |
| Ngf | batch | mp | d2 | Ngfr | fap |
| Ngf | constitutive | per | d2 | Ngfr | fap |
| Nppc | batch | mp | d2 | Npr2 | fap |
| Pdap1 | batch | mp | d2 | Pdgfa | fap |
| S100b | batch | mp | d2 | Fgfr1 | fap |
| S100b | constitutive | per | d2 | Fgfr1 | fap |
| Serpini1 | batch | mp | d2 | Plat | fap |
| Serpini1 | constitutive | per | d2 | Plat | fap |
| Sfrp4 | batch | mp | d2 | Fzd2 | fap |
| Sfrp5 | batch | mp | d2 | Fzd2 | fap |
| Tgfb1 | batch | mp | d2 | Acvrl1 | fap |
| Tgfb1 | total | ec | d2 | Acvrl1 | fap |
| Tgfb1 | total | ic | d2 | Acvrl1 | fap |
| Tgfb1 | batch | mp | d2 | Tgfbr3 | fap |
| Tgfb1 | total | ec | d2 | Tgfbr3 | fap |
| Tgfb1 | total | ic | d2 | Tgfbr3 | fap |
| Tgfb1 | batch | mp | d2 | Tgfbr2 | fap |
| Tgfb1 | total | ec | d2 | Tgfbr2 | fap |
| Tgfb1 | total | ic | d2 | Tgfbr2 | fap |
| Tgfb1 | batch | mp | d2 | Itgav | fap |
| Tgfb1 | total | ec | d2 | Itgav | fap |
| Tgfb1 | total | ic | d2 | Itgav | fap |
| Tnf | batch | ic | d2 | Tnfrsf1a | fap |
| Tnf | batch | mp | d2 | Tnfrsf1a | fap |
| Tnf | batch | per | d2 | Tnfrsf1a | fap |
| Vegfa | batch | ic | d2 | Nrp1 | fap |
| Vegfa | batch | mp | d2 | Nrp1 | fap |
| Vegfa | batch | ic | d2 | Ephb2 | fap |
| Vegfa | batch | mp | d2 | Ephb2 | fap |
| Vegfa | batch | ic | d2 | Itga9 | fap |
| Vegfa | batch | mp | d2 | Itga9 | fap |
| Vegfc | batch | mp | d2 | Nrp1 | fap |
| Vegfc | batch | mp | d2 | Itga9 | fap |
| Il1a | batch | ic | d2 | Il1r1 | fap |
| Il1a | batch | ic | d2 | Il1r2 | fap |
| Jag1 | batch | ic | d2 | Notch1 | fap |
| Jag1 | constitutive | per | d2 | Notch1 | fap |
| Jag1 | batch | ic | d2 | Notch3 | fap |
| Jag1 | constitutive | per | d2 | Notch3 | fap |
| Jag1 | batch | ic | d2 | Notch2 | fap |
| Jag1 | constitutive | per | d2 | Notch2 | fap |
| Pdgfa | batch | ic | d2 | Pdgfra | fap |
| Pdgfa | constitutive | ec | d2 | Pdgfra | fap |
| Pdgfa | constitutive | mp | d2 | Pdgfra | fap |
| Pdgfa | constitutive | per | d2 | Pdgfra | fap |
| Ppbp | batch | ic | d2 | Slc1a5 | fap |
| Ppbp | batch | ic | d2 | Itgb5 | fap |
| Tnfsf13 | batch | ic | d2 | Tnfrsf11b | fap |
| Tnfsf13 | batch | ic | d2 | Tnfrsf12a | fap |
| Tnfsf13 | batch | ic | d2 | Tnfrsf1a | fap |
| Btc | batch | per | d2 | Egfr | fap |
| Il11 | batch | per | d2 | Il11ra1 | fap |
| Il11 | batch | per | d2 | Il6st | fap |
| Nov | batch | per | d2 | Itga5 | fap |
| Nov | batch | per | d2 | Notch1 | fap |
| Nov | batch | per | d2 | Itgb3 | fap |
| Nov | batch | per | d2 | Itgav | fap |
| Tnfsf12 | total | ec | d2 | Tnfrsf11b | fap |
| Tnfsf12 | total | ic | d2 | Tnfrsf11b | fap |
| Tnfsf12 | total | per | d2 | Tnfrsf11b | fap |
| Tnfsf12 | total | ec | d2 | Tnfrsf12a | fap |
| Tnfsf12 | total | ic | d2 | Tnfrsf12a | fap |
| Tnfsf12 | total | per | d2 | Tnfrsf12a | fap |
| Tnfsf12 | total | ec | d2 | Tnfrsf1a | fap |
| Tnfsf12 | total | ic | d2 | Tnfrsf1a | fap |
| Tnfsf12 | total | per | d2 | Tnfrsf1a | fap |
| Fgf7 | constitutive | fap | d2 | Nrp1 | fap |
| Tgfb2 | constitutive | ec | d2 | Tgfbr3 | fap |
| Tgfb2 | constitutive | fap | d2 | Tgfbr3 | fap |
| Tgfb2 | constitutive | per | d2 | Tgfbr3 | fap |
| Tgfb2 | constitutive | ec | d2 | Tgfbr2 | fap |
| Tgfb2 | constitutive | fap | d2 | Tgfbr2 | fap |
| Tgfb2 | constitutive | per | d2 | Tgfbr2 | fap |
| Bmp5 | constitutive | ec | d2 | Bmpr1a | fap |
| Bmp5 | constitutive | fap | d2 | Bmpr1a | fap |
| Bmp5 | constitutive | per | d2 | Bmpr1a | fap |
| Pdgfb | constitutive | ec | d2 | Lrp1 | fap |
| Pdgfb | constitutive | ec | d2 | Pdgfra | fap |
| Pdgfb | constitutive | ec | d2 | Pdgfrb | fap |
| Cmtm8 | constitutive | ec | d2 | Egfr | fap |
| Fgf1 | constitutive | per | d2 | Fgfr1 | fap |
| Ptn | constitutive | per | d2 | Ptprs | fap |
| Ntf3 | constitutive | per | d2 | Ngfr | fap |
| Apln | batch | ec | d3 | Aplnr | fap |
| Apln | batch | fap | d3 | Aplnr | fap |
| Bmp2 | batch | fap | d3 | Bmpr1a | fap |
| Bmp2 | constitutive | per | d3 | Bmpr1a | fap |
| Bmp2 | batch | fap | d3 | Acvr2a | fap |
| Bmp2 | constitutive | per | d3 | Acvr2a | fap |
| Bmp6 | batch | fap | d3 | Bmpr1a | fap |
| Bmp6 | constitutive | per | d3 | Bmpr1a | fap |
| Bmp6 | batch | fap | d3 | Acvr2a | fap |
| Bmp6 | constitutive | per | d3 | Acvr2a | fap |
| Ccl5 | batch | ec | d3 | Sdc4 | fap |
| Ccl5 | batch | fap | d3 | Sdc4 | fap |
| Csf2 | batch | fap | d3 | Il3ra | fap |
| Cst3 | batch | fap | d3 | Tgfbr2 | fap |
| Cst3 | batch | mp | d3 | Tgfbr2 | fap |
| Cst3 | constitutive | ic | d3 | Tgfbr2 | fap |
| Cst3 | constitutive | per | d3 | Tgfbr2 | fap |
| Ctgf | batch | fap | d3 | Itga5 | fap |
| Ctgf | constitutive | per | d3 | Itga5 | fap |
| Cxcl10 | batch | ec | d3 | Dpp4 | fap |
| Cxcl10 | batch | fap | d3 | Dpp4 | fap |
| Cxcl10 | batch | ic | d3 | Dpp4 | fap |
| Cxcl10 | batch | per | d3 | Dpp4 | fap |
| Cxcl12 | batch | fap | d3 | Cxcr4 | fap |
| Cxcl12 | constitutive | ec | d3 | Cxcr4 | fap |
| Cxcl12 | constitutive | per | d3 | Cxcr4 | fap |
| Cxcl12 | batch | fap | d3 | Dpp4 | fap |
| Cxcl12 | constitutive | ec | d3 | Dpp4 | fap |
| Cxcl12 | constitutive | per | d3 | Dpp4 | fap |
| Cxcl9 | batch | ec | d3 | Dpp4 | fap |
| Cxcl9 | batch | fap | d3 | Dpp4 | fap |
| Cxcl9 | batch | ic | d3 | Dpp4 | fap |
| Efnb1 | batch | fap | d3 | Ephb2 | fap |
| Efnb1 | constitutive | ec | d3 | Ephb2 | fap |
| Efnb1 | constitutive | per | d3 | Ephb2 | fap |
| Fgf1 | batch | fap | d3 | Fgfr2 | fap |
| Fgf1 | constitutive | per | d3 | Fgfr2 | fap |
| Fgf1 | batch | fap | d3 | Fgfr1 | fap |
| Fgf1 | constitutive | per | d3 | Fgfr1 | fap |
| Gas6 | batch | fap | d3 | Axl | fap |
| Hbegf | batch | fap | d3 | Cd44 | fap |
| Hbegf | constitutive | ec | d3 | Cd44 | fap |
| Hbegf | constitutive | per | d3 | Cd44 | fap |
| Hbegf | batch | fap | d3 | Egfr | fap |
| Hbegf | constitutive | ec | d3 | Egfr | fap |
| Hbegf | constitutive | per | d3 | Egfr | fap |
| Hgf | batch | fap | d3 | Vtn | fap |
| Hgf | batch | ic | d3 | Vtn | fap |
| Igf1 | batch | ec | d3 | Igfbp3 | fap |
| Igf1 | batch | fap | d3 | Igfbp3 | fap |
| Igf1 | batch | ic | d3 | Igfbp3 | fap |
| Igf1 | batch | mp | d3 | Igfbp3 | fap |
| Igf1 | batch | ec | d3 | Igfbp4 | fap |
| Igf1 | batch | fap | d3 | Igfbp4 | fap |
| Igf1 | batch | ic | d3 | Igfbp4 | fap |
| Igf1 | batch | mp | d3 | Igfbp4 | fap |
| Igf1 | batch | ec | d3 | Igfbp5 | fap |
| Igf1 | batch | fap | d3 | Igfbp5 | fap |
| Igf1 | batch | ic | d3 | Igfbp5 | fap |
| Igf1 | batch | mp | d3 | Igfbp5 | fap |
| Igf1 | batch | ec | d3 | Igfbp7 | fap |
| Igf1 | batch | fap | d3 | Igfbp7 | fap |
| Igf1 | batch | ic | d3 | Igfbp7 | fap |
| Igf1 | batch | mp | d3 | Igfbp7 | fap |
| Igf1 | batch | ec | d3 | Igfbp2 | fap |
| Igf1 | batch | fap | d3 | Igfbp2 | fap |
| Igf1 | batch | ic | d3 | Igfbp2 | fap |
| Igf1 | batch | mp | d3 | Igfbp2 | fap |
| Igf2 | batch | fap | d3 | Vtn | fap |
| Il18 | batch | fap | d3 | Il1rl2 | fap |
| Il18 | constitutive | ic | d3 | Il1rl2 | fap |
| Il18 | constitutive | mp | d3 | Il1rl2 | fap |
| Il1b | batch | fap | d3 | Il1r1 | fap |
| Il1b | batch | mp | d3 | Il1r1 | fap |
| Il1b | batch | per | d3 | Il1r1 | fap |
| Il1b | batch | fap | d3 | Il1r2 | fap |
| Il1b | batch | mp | d3 | Il1r2 | fap |
| Il1b | batch | per | d3 | Il1r2 | fap |
| Il33 | batch | fap | d3 | Il1rl1 | fap |
| Il33 | batch | mp | d3 | Il1rl1 | fap |
| Mif | batch | fap | d3 | Cd74 | fap |
| Mif | batch | mp | d3 | Cd74 | fap |
| Mmp13 | batch | ec | d3 | F2r | fap |
| Mmp13 | batch | fap | d3 | F2r | fap |
| Mmp13 | batch | ic | d3 | F2r | fap |
| Mmp13 | batch | per | d3 | F2r | fap |
| Ntf3 | batch | ec | d3 | Ngfr | fap |
| Ntf3 | batch | fap | d3 | Ngfr | fap |
| Ntf3 | constitutive | per | d3 | Ngfr | fap |
| Pdgfa | batch | fap | d3 | Pdgfra | fap |
| Pdgfa | batch | ic | d3 | Pdgfra | fap |
| Pdgfa | constitutive | ec | d3 | Pdgfra | fap |
| Pdgfa | constitutive | mp | d3 | Pdgfra | fap |
| Pdgfa | constitutive | per | d3 | Pdgfra | fap |
| Pdgfc | batch | fap | d3 | Pdgfra | fap |
| Pdgfc | constitutive | ic | d3 | Pdgfra | fap |
| Pdgfc | constitutive | per | d3 | Pdgfra | fap |
| Pthlh | batch | fap | d3 | Pth1r | fap |
| Ptn | batch | fap | d3 | Ptprs | fap |
| Ptn | constitutive | per | d3 | Ptprs | fap |
| S100b | batch | fap | d3 | Fgfr1 | fap |
| S100b | constitutive | per | d3 | Fgfr1 | fap |
| Sfrp1 | batch | ec | d3 | Fzd2 | fap |
| Sfrp1 | batch | fap | d3 | Fzd2 | fap |
| Sfrp1 | constitutive | per | d3 | Fzd2 | fap |
| Sfrp2 | batch | ec | d3 | Fzd2 | fap |
| Sfrp2 | batch | fap | d3 | Fzd2 | fap |
| Sfrp2 | batch | per | d3 | Fzd2 | fap |
| Sfrp4 | batch | fap | d3 | Fzd2 | fap |
| Spp1 | batch | ec | d3 | Itga5 | fap |
| Spp1 | batch | fap | d3 | Itga5 | fap |
| Spp1 | batch | ic | d3 | Itga5 | fap |
| Spp1 | batch | mp | d3 | Itga5 | fap |
| Spp1 | batch | per | d3 | Itga5 | fap |
| Spp1 | batch | ec | d3 | Vtn | fap |
| Spp1 | batch | fap | d3 | Vtn | fap |
| Spp1 | batch | ic | d3 | Vtn | fap |
| Spp1 | batch | mp | d3 | Vtn | fap |
| Spp1 | batch | per | d3 | Vtn | fap |
| Spp1 | batch | ec | d3 | Itgb5 | fap |
| Spp1 | batch | fap | d3 | Itgb5 | fap |
| Spp1 | batch | ic | d3 | Itgb5 | fap |
| Spp1 | batch | mp | d3 | Itgb5 | fap |
| Spp1 | batch | per | d3 | Itgb5 | fap |
| Spp1 | batch | ec | d3 | Itgb3 | fap |
| Spp1 | batch | fap | d3 | Itgb3 | fap |
| Spp1 | batch | ic | d3 | Itgb3 | fap |
| Spp1 | batch | mp | d3 | Itgb3 | fap |
| Spp1 | batch | per | d3 | Itgb3 | fap |
| Spp1 | batch | ec | d3 | Itgav | fap |
| Spp1 | batch | fap | d3 | Itgav | fap |
| Spp1 | batch | ic | d3 | Itgav | fap |
| Spp1 | batch | mp | d3 | Itgav | fap |
| Spp1 | batch | per | d3 | Itgav | fap |
| Spp1 | batch | ec | d3 | Itga9 | fap |
| Spp1 | batch | fap | d3 | Itga9 | fap |
| Spp1 | batch | ic | d3 | Itga9 | fap |
| Spp1 | batch | mp | d3 | Itga9 | fap |
| Spp1 | batch | per | d3 | Itga9 | fap |
| Tgfb3 | batch | fap | d3 | Acvrl1 | fap |
| Tgfb3 | total | ec | d3 | Acvrl1 | fap |
| Tgfb3 | total | per | d3 | Acvrl1 | fap |
| Tgfb3 | batch | fap | d3 | Tgfbr2 | fap |
| Tgfb3 | total | ec | d3 | Tgfbr2 | fap |
| Tgfb3 | total | per | d3 | Tgfbr2 | fap |
| Tgfb3 | batch | fap | d3 | Itgav | fap |
| Tgfb3 | total | ec | d3 | Itgav | fap |
| Tgfb3 | total | per | d3 | Itgav | fap |
| Tnfrsf11b | batch | fap | d3 | Vtn | fap |
| Tslp | batch | fap | d3 | Crlf2 | fap |
| Tslp | constitutive | ec | d3 | Crlf2 | fap |
| Wnt2 | batch | fap | d3 | Sfrp1 | fap |
| Wnt2 | batch | fap | d3 | Fzd1 | fap |
| Fgf9 | batch | ec | d3 | Fgfr2 | fap |
| Il15 | batch | ec | d3 | Il2rg | fap |
| Il15 | batch | ic | d3 | Il2rg | fap |
| Il15 | batch | ec | d3 | Il17ra | fap |
| Il15 | batch | ic | d3 | Il17ra | fap |
| Il15 | batch | ec | d3 | Il15ra | fap |
| Il15 | batch | ic | d3 | Il15ra | fap |
| Il6 | batch | ec | d3 | Il6st | fap |
| Il6 | batch | per | d3 | Il6st | fap |
| Inhbb | batch | ec | d3 | Acvr2a | fap |
| Inhbb | constitutive | fap | d3 | Acvr2a | fap |
| Inhbb | constitutive | per | d3 | Acvr2a | fap |
| Inhbb | batch | ec | d3 | Acvr1 | fap |
| Inhbb | constitutive | fap | d3 | Acvr1 | fap |
| Inhbb | constitutive | per | d3 | Acvr1 | fap |
| Lgals3 | batch | ec | d3 | Lgals3bp | fap |
| Lgals3 | batch | ic | d3 | Lgals3bp | fap |
| Lgals3 | batch | mp | d3 | Lgals3bp | fap |
| Lgals3 | batch | per | d3 | Lgals3bp | fap |
| Mmp12 | batch | ec | d3 | Plaur | fap |
| Mmp12 | batch | ic | d3 | Plaur | fap |
| Mmp12 | batch | mp | d3 | Plaur | fap |
| Mmp12 | batch | per | d3 | Plaur | fap |
| Osm | batch | ec | d3 | Osmr | fap |
| Osm | batch | mp | d3 | Osmr | fap |
| Osm | batch | per | d3 | Osmr | fap |
| Osm | batch | ec | d3 | Il6st | fap |
| Osm | batch | mp | d3 | Il6st | fap |
| Osm | batch | per | d3 | Il6st | fap |
| Tnfsf10 | batch | ec | d3 | Tnfrsf10b | fap |
| Vegfc | batch | ec | d3 | Nrp1 | fap |
| Vegfc | batch | ec | d3 | Itga9 | fap |
| Csf1 | batch | ic | d3 | Csf1r | fap |
| Csf1 | batch | mp | d3 | Csf1r | fap |
| Cxcl2 | batch | mp | d3 | Dpp4 | fap |
| Cxcl2 | batch | per | d3 | Dpp4 | fap |
| Pdap1 | batch | mp | d3 | Pdgfa | fap |
| Tgfb1 | batch | mp | d3 | Vtn | fap |
| Tgfb1 | total | ec | d3 | Vtn | fap |
| Tgfb1 | total | ic | d3 | Vtn | fap |
| Tgfb1 | batch | mp | d3 | Acvrl1 | fap |
| Tgfb1 | total | ec | d3 | Acvrl1 | fap |
| Tgfb1 | total | ic | d3 | Acvrl1 | fap |
| Tgfb1 | batch | mp | d3 | Tgfbr3 | fap |
| Tgfb1 | total | ec | d3 | Tgfbr3 | fap |
| Tgfb1 | total | ic | d3 | Tgfbr3 | fap |
| Tgfb1 | batch | mp | d3 | Tgfbr2 | fap |
| Tgfb1 | total | ec | d3 | Tgfbr2 | fap |
| Tgfb1 | total | ic | d3 | Tgfbr2 | fap |
| Tgfb1 | batch | mp | d3 | Itgav | fap |
| Tgfb1 | total | ec | d3 | Itgav | fap |
| Tgfb1 | total | ic | d3 | Itgav | fap |
| Tnf | batch | mp | d3 | Tnfrsf1a | fap |
| Tnf | batch | per | d3 | Tnfrsf1a | fap |
| Tnfsf13 | batch | ic | d3 | Tnfrsf11b | fap |
| Tnfsf13 | batch | ic | d3 | Tnfrsf12a | fap |
| Tnfsf13 | batch | ic | d3 | Tnfrsf1a | fap |
| Btc | batch | per | d3 | Erbb2 | fap |
| Btc | batch | per | d3 | Egfr | fap |
| Il11 | batch | per | d3 | Il11ra1 | fap |
| Il11 | batch | per | d3 | Il6st | fap |
| Lif | batch | per | d3 | Il6st | fap |
| Lif | constitutive | mp | d3 | Il6st | fap |
| Dll1 | total | mp | d3 | Notch1 | fap |
| Dll1 | total | mp | d3 | Notch3 | fap |
| Dll1 | total | mp | d3 | Notch2 | fap |
| Fgf7 | constitutive | fap | d3 | Fgfr2 | fap |
| Fgf7 | constitutive | fap | d3 | Nrp1 | fap |
| Tgfb2 | constitutive | ec | d3 | Vtn | fap |
| Tgfb2 | constitutive | fap | d3 | Vtn | fap |
| Tgfb2 | constitutive | per | d3 | Vtn | fap |
| Tgfb2 | constitutive | ec | d3 | Tgfbr3 | fap |
| Tgfb2 | constitutive | fap | d3 | Tgfbr3 | fap |
| Tgfb2 | constitutive | per | d3 | Tgfbr3 | fap |
| Tgfb2 | constitutive | ec | d3 | Tgfbr2 | fap |
| Tgfb2 | constitutive | fap | d3 | Tgfbr2 | fap |
| Tgfb2 | constitutive | per | d3 | Tgfbr2 | fap |
| Bmp5 | constitutive | ec | d3 | Bmpr1a | fap |
| Bmp5 | constitutive | fap | d3 | Bmpr1a | fap |
| Bmp5 | constitutive | per | d3 | Bmpr1a | fap |
| Jag2 | constitutive | ec | d3 | Notch1 | fap |
| Jag2 | constitutive | ec | d3 | Notch3 | fap |
| Jag2 | constitutive | ec | d3 | Notch2 | fap |
| Pdgfb | constitutive | ec | d3 | Pdgfra | fap |
| Pdgfb | constitutive | ec | d3 | Pdgfrb | fap |
| Cmtm8 | constitutive | ec | d3 | Egfr | fap |
| Edn1 | constitutive | ec | d3 | Ednra | fap |
| Bmp4 | constitutive | ec | d3 | Bmpr1a | fap |
| Bmp4 | constitutive | per | d3 | Bmpr1a | fap |
| Serpini1 | constitutive | per | d3 | Plat | fap |
| Ngf | constitutive | per | d3 | Ngfr | fap |
| Pgf | constitutive | per | d3 | Nrp1 | fap |
| Jag1 | constitutive | per | d3 | Notch1 | fap |
| Jag1 | constitutive | per | d3 | Notch3 | fap |
| Jag1 | constitutive | per | d3 | Notch2 | fap |
| Apln | batch | fap | d4 | Aplnr | fap |
| Ccl5 | batch | ec | d4 | Sdc4 | fap |
| Ccl5 | batch | fap | d4 | Sdc4 | fap |
| Cst3 | batch | fap | d4 | Tgfbr2 | fap |
| Cst3 | constitutive | ic | d4 | Tgfbr2 | fap |
| Cst3 | constitutive | per | d4 | Tgfbr2 | fap |
| Ctgf | batch | fap | d4 | Itga5 | fap |
| Ctgf | constitutive | per | d4 | Itga5 | fap |
| Cxcl12 | batch | fap | d4 | Cxcr4 | fap |
| Cxcl12 | constitutive | ec | d4 | Cxcr4 | fap |
| Cxcl12 | constitutive | per | d4 | Cxcr4 | fap |
| Efnb1 | batch | fap | d4 | Ephb2 | fap |
| Efnb1 | batch | mp | d4 | Ephb2 | fap |
| Efnb1 | constitutive | ec | d4 | Ephb2 | fap |
| Efnb1 | constitutive | per | d4 | Ephb2 | fap |
| Fgf1 | batch | fap | d4 | Fgfr2 | fap |
| Fgf1 | constitutive | per | d4 | Fgfr2 | fap |
| Fgf1 | batch | fap | d4 | Fgfr1 | fap |
| Fgf1 | constitutive | per | d4 | Fgfr1 | fap |
| Gas6 | batch | fap | d4 | Axl | fap |
| Hbegf | batch | fap | d4 | Cd44 | fap |
| Hbegf | constitutive | ec | d4 | Cd44 | fap |
| Hbegf | constitutive | per | d4 | Cd44 | fap |
| Hbegf | batch | fap | d4 | Egfr | fap |
| Hbegf | constitutive | ec | d4 | Egfr | fap |
| Hbegf | constitutive | per | d4 | Egfr | fap |
| Hgf | batch | fap | d4 | Vtn | fap |
| Hgf | batch | ic | d4 | Vtn | fap |
| Igf1 | batch | ec | d4 | Igfbp3 | fap |
| Igf1 | batch | fap | d4 | Igfbp3 | fap |
| Igf1 | batch | ic | d4 | Igfbp3 | fap |
| Igf1 | batch | ec | d4 | Igfbp4 | fap |
| Igf1 | batch | fap | d4 | Igfbp4 | fap |
| Igf1 | batch | ic | d4 | Igfbp4 | fap |
| Igf1 | batch | ec | d4 | Igfbp5 | fap |
| Igf1 | batch | fap | d4 | Igfbp5 | fap |
| Igf1 | batch | ic | d4 | Igfbp5 | fap |
| Igf1 | batch | ec | d4 | Igfbp7 | fap |
| Igf1 | batch | fap | d4 | Igfbp7 | fap |
| Igf1 | batch | ic | d4 | Igfbp7 | fap |
| Igf1 | batch | ec | d4 | Igfbp2 | fap |
| Igf1 | batch | fap | d4 | Igfbp2 | fap |
| Igf1 | batch | ic | d4 | Igfbp2 | fap |
| Igf2 | batch | ec | d4 | Vtn | fap |
| Igf2 | batch | fap | d4 | Vtn | fap |
| Igf2 | batch | mp | d4 | Vtn | fap |
| Il1b | batch | fap | d4 | Il1r1 | fap |
| Il33 | batch | fap | d4 | Il1rl1 | fap |
| Il33 | batch | mp | d4 | Il1rl1 | fap |
| Mif | batch | fap | d4 | Cd74 | fap |
| Mmp13 | batch | fap | d4 | F2r | fap |
| Mmp13 | batch | ic | d4 | F2r | fap |
| Mmp13 | batch | mp | d4 | F2r | fap |
| Ntf3 | batch | ec | d4 | Ngfr | fap |
| Ntf3 | batch | fap | d4 | Ngfr | fap |
| Ntf3 | constitutive | per | d4 | Ngfr | fap |
| Pdgfa | batch | fap | d4 | Pdgfra | fap |
| Pdgfa | batch | ic | d4 | Pdgfra | fap |
| Pdgfa | constitutive | ec | d4 | Pdgfra | fap |
| Pdgfa | constitutive | mp | d4 | Pdgfra | fap |
| Pdgfa | constitutive | per | d4 | Pdgfra | fap |
| Pthlh | batch | fap | d4 | Pth1r | fap |
| Ptn | batch | fap | d4 | Ptprs | fap |
| Ptn | constitutive | per | d4 | Ptprs | fap |
| S100b | batch | fap | d4 | Fgfr1 | fap |
| S100b | constitutive | per | d4 | Fgfr1 | fap |
| Sfrp1 | batch | ec | d4 | Fzd2 | fap |
| Sfrp1 | batch | fap | d4 | Fzd2 | fap |
| Sfrp1 | constitutive | per | d4 | Fzd2 | fap |
| Sfrp2 | batch | ec | d4 | Fzd2 | fap |
| Sfrp2 | batch | fap | d4 | Fzd2 | fap |
| Sfrp4 | batch | fap | d4 | Fzd2 | fap |
| Spp1 | batch | ec | d4 | Itga5 | fap |
| Spp1 | batch | fap | d4 | Itga5 | fap |
| Spp1 | batch | ic | d4 | Itga5 | fap |
| Spp1 | batch | mp | d4 | Itga5 | fap |
| Spp1 | batch | ec | d4 | Vtn | fap |
| Spp1 | batch | fap | d4 | Vtn | fap |
| Spp1 | batch | ic | d4 | Vtn | fap |
| Spp1 | batch | mp | d4 | Vtn | fap |
| Spp1 | batch | ec | d4 | Itgb5 | fap |
| Spp1 | batch | fap | d4 | Itgb5 | fap |
| Spp1 | batch | ic | d4 | Itgb5 | fap |
| Spp1 | batch | mp | d4 | Itgb5 | fap |
| Spp1 | batch | ec | d4 | Itgb3 | fap |
| Spp1 | batch | fap | d4 | Itgb3 | fap |
| Spp1 | batch | ic | d4 | Itgb3 | fap |
| Spp1 | batch | mp | d4 | Itgb3 | fap |
| Spp1 | batch | ec | d4 | Itgav | fap |
| Spp1 | batch | fap | d4 | Itgav | fap |
| Spp1 | batch | ic | d4 | Itgav | fap |
| Spp1 | batch | mp | d4 | Itgav | fap |
| Spp1 | batch | ec | d4 | Itga9 | fap |
| Spp1 | batch | fap | d4 | Itga9 | fap |
| Spp1 | batch | ic | d4 | Itga9 | fap |
| Spp1 | batch | mp | d4 | Itga9 | fap |
| Tgfb3 | batch | fap | d4 | Acvrl1 | fap |
| Tgfb3 | total | ec | d4 | Acvrl1 | fap |
| Tgfb3 | total | per | d4 | Acvrl1 | fap |
| Tgfb3 | batch | fap | d4 | Tgfbr2 | fap |
| Tgfb3 | total | ec | d4 | Tgfbr2 | fap |
| Tgfb3 | total | per | d4 | Tgfbr2 | fap |
| Tgfb3 | batch | fap | d4 | Itgav | fap |
| Tgfb3 | total | ec | d4 | Itgav | fap |
| Tgfb3 | total | per | d4 | Itgav | fap |
| Tnfrsf11b | batch | fap | d4 | Vtn | fap |
| Dll1 | batch | ec | d4 | Notch1 | fap |
| Dll1 | total | mp | d4 | Notch1 | fap |
| Dll1 | batch | ec | d4 | Notch3 | fap |
| Dll1 | total | mp | d4 | Notch3 | fap |
| Dll1 | batch | ec | d4 | Notch2 | fap |
| Dll1 | total | mp | d4 | Notch2 | fap |
| Fgf9 | batch | ec | d4 | Fgfr2 | fap |
| Il15 | batch | ec | d4 | Il2rg | fap |
| Il15 | batch | ic | d4 | Il2rg | fap |
| Il15 | batch | ec | d4 | Il15ra | fap |
| Il15 | batch | ic | d4 | Il15ra | fap |
| Lgals3 | batch | ec | d4 | Lgals3bp | fap |
| Lgals3 | batch | ic | d4 | Lgals3bp | fap |
| Lgals3 | batch | mp | d4 | Lgals3bp | fap |
| Mmp12 | batch | ec | d4 | Plaur | fap |
| Mmp12 | batch | ic | d4 | Plaur | fap |
| Mmp12 | batch | mp | d4 | Plaur | fap |
| Osm | batch | ec | d4 | Osmr | fap |
| Osm | batch | ec | d4 | Il6st | fap |
| Tnfsf10 | batch | ec | d4 | Tnfrsf10b | fap |
| Vegfc | batch | ec | d4 | Nrp1 | fap |
| Vegfc | batch | mp | d4 | Nrp1 | fap |
| Vegfc | batch | ec | d4 | Itga9 | fap |
| Vegfc | batch | mp | d4 | Itga9 | fap |
| Areg | batch | mp | d4 | Egfr | fap |
| Gdnf | batch | mp | d4 | Gfra1 | fap |
| Gdnf | batch | mp | d4 | Gfra2 | fap |
| Gmfb | batch | mp | d4 | Egfr | fap |
| Pdap1 | batch | mp | d4 | Pdgfa | fap |
| Pdgfc | batch | mp | d4 | Pdgfra | fap |
| Pdgfc | constitutive | ic | d4 | Pdgfra | fap |
| Pdgfc | constitutive | per | d4 | Pdgfra | fap |
| Tgfb1 | batch | mp | d4 | Vtn | fap |
| Tgfb1 | batch | mp | d4 | Acvrl1 | fap |
| Tgfb1 | batch | mp | d4 | Tgfbr3 | fap |
| Tgfb1 | batch | mp | d4 | Tgfbr2 | fap |
| Tgfb1 | batch | mp | d4 | Itgav | fap |
| Tgfb2 | batch | mp | d4 | Vtn | fap |
| Tgfb2 | constitutive | ec | d4 | Vtn | fap |
| Tgfb2 | constitutive | fap | d4 | Vtn | fap |
| Tgfb2 | constitutive | per | d4 | Vtn | fap |
| Tgfb2 | batch | mp | d4 | Tgfbr3 | fap |
| Tgfb2 | constitutive | ec | d4 | Tgfbr3 | fap |
| Tgfb2 | constitutive | fap | d4 | Tgfbr3 | fap |
| Tgfb2 | constitutive | per | d4 | Tgfbr3 | fap |
| Tgfb2 | batch | mp | d4 | Tgfbr2 | fap |
| Tgfb2 | constitutive | ec | d4 | Tgfbr2 | fap |
| Tgfb2 | constitutive | fap | d4 | Tgfbr2 | fap |
| Tgfb2 | constitutive | per | d4 | Tgfbr2 | fap |
| Csf1 | batch | ic | d4 | Csf1r | fap |
| Pdgfb | batch | ic | d4 | Pdgfra | fap |
| Pdgfb | constitutive | ec | d4 | Pdgfra | fap |
| Pdgfb | batch | ic | d4 | Pdgfrb | fap |
| Pdgfb | constitutive | ec | d4 | Pdgfrb | fap |
| Tnfsf13 | batch | ic | d4 | Tnfrsf11b | fap |
| Tnfsf13 | batch | ic | d4 | Tnfrsf12a | fap |
| Tnfsf13 | batch | ic | d4 | Tnfrsf1a | fap |
| Fgf7 | constitutive | fap | d4 | Fgfr2 | fap |
| Fgf7 | constitutive | fap | d4 | Nrp1 | fap |
| Bmp5 | constitutive | ec | d4 | Bmpr1a | fap |
| Bmp5 | constitutive | fap | d4 | Bmpr1a | fap |
| Bmp5 | constitutive | per | d4 | Bmpr1a | fap |
| Inhbb | constitutive | fap | d4 | Acvr2a | fap |
| Inhbb | constitutive | per | d4 | Acvr2a | fap |
| Inhbb | constitutive | fap | d4 | Acvr1 | fap |
| Inhbb | constitutive | per | d4 | Acvr1 | fap |
| Jag2 | constitutive | ec | d4 | Notch1 | fap |
| Jag2 | constitutive | ec | d4 | Notch3 | fap |
| Jag2 | constitutive | ec | d4 | Notch2 | fap |
| Cmtm8 | constitutive | ec | d4 | Egfr | fap |
| Edn1 | constitutive | ec | d4 | Ednra | fap |
| Bmp4 | constitutive | ec | d4 | Bmpr1a | fap |
| Bmp4 | constitutive | per | d4 | Bmpr1a | fap |
| Tslp | constitutive | ec | d4 | Crlf2 | fap |
| Il18 | constitutive | ic | d4 | Il1rl2 | fap |
| Il18 | constitutive | mp | d4 | Il1rl2 | fap |
| Lif | constitutive | mp | d4 | Il6st | fap |
| Serpini1 | constitutive | per | d4 | Plat | fap |
| Ngf | constitutive | per | d4 | Ngfr | fap |
| Bmp6 | constitutive | per | d4 | Bmpr1a | fap |
| Bmp6 | constitutive | per | d4 | Acvr2a | fap |
| Pgf | constitutive | per | d4 | Nrp1 | fap |
| Jag1 | constitutive | per | d4 | Notch1 | fap |
| Jag1 | constitutive | per | d4 | Notch3 | fap |
| Jag1 | constitutive | per | d4 | Notch2 | fap |
| Bmp2 | constitutive | per | d4 | Bmpr1a | fap |
| Bmp2 | constitutive | per | d4 | Acvr2a | fap |
| Apln | batch | ec | d5 | Aplnr | fap |
| Apln | batch | fap | d5 | Aplnr | fap |
| Ccl5 | batch | fap | d5 | Sdc4 | fap |
| Cst3 | batch | fap | d5 | Tgfbr2 | fap |
| Cst3 | constitutive | ic | d5 | Tgfbr2 | fap |
| Cst3 | constitutive | per | d5 | Tgfbr2 | fap |
| Ctgf | batch | fap | d5 | Itga5 | fap |
| Ctgf | batch | ic | d5 | Itga5 | fap |
| Ctgf | constitutive | per | d5 | Itga5 | fap |
| Cxcl12 | batch | fap | d5 | Cxcr4 | fap |
| Cxcl12 | batch | ic | d5 | Cxcr4 | fap |
| Cxcl12 | constitutive | ec | d5 | Cxcr4 | fap |
| Cxcl12 | constitutive | per | d5 | Cxcr4 | fap |
| Efnb1 | batch | fap | d5 | Ephb2 | fap |
| Efnb1 | batch | ic | d5 | Ephb2 | fap |
| Efnb1 | batch | mp | d5 | Ephb2 | fap |
| Efnb1 | constitutive | ec | d5 | Ephb2 | fap |
| Efnb1 | constitutive | per | d5 | Ephb2 | fap |
| Fgf1 | batch | fap | d5 | Fgfr2 | fap |
| Fgf1 | constitutive | per | d5 | Fgfr2 | fap |
| Fgf1 | batch | fap | d5 | Fgfr1 | fap |
| Fgf1 | constitutive | per | d5 | Fgfr1 | fap |
| Gas6 | batch | fap | d5 | Axl | fap |
| Hbegf | batch | fap | d5 | Cd44 | fap |
| Hbegf | constitutive | ec | d5 | Cd44 | fap |
| Hbegf | constitutive | per | d5 | Cd44 | fap |
| Hbegf | batch | fap | d5 | Egfr | fap |
| Hbegf | constitutive | ec | d5 | Egfr | fap |
| Hbegf | constitutive | per | d5 | Egfr | fap |
| Hgf | batch | fap | d5 | Vtn | fap |
| Hgf | batch | ic | d5 | Vtn | fap |
| Igf1 | batch | ec | d5 | Igfbp3 | fap |
| Igf1 | batch | fap | d5 | Igfbp3 | fap |
| Igf1 | batch | ic | d5 | Igfbp3 | fap |
| Igf1 | batch | per | d5 | Igfbp3 | fap |
| Igf1 | batch | ec | d5 | Igfbp4 | fap |
| Igf1 | batch | fap | d5 | Igfbp4 | fap |
| Igf1 | batch | ic | d5 | Igfbp4 | fap |
| Igf1 | batch | per | d5 | Igfbp4 | fap |
| Igf1 | batch | ec | d5 | Igfbp5 | fap |
| Igf1 | batch | fap | d5 | Igfbp5 | fap |
| Igf1 | batch | ic | d5 | Igfbp5 | fap |
| Igf1 | batch | per | d5 | Igfbp5 | fap |
| Igf1 | batch | ec | d5 | Igfbp7 | fap |
| Igf1 | batch | fap | d5 | Igfbp7 | fap |
| Igf1 | batch | ic | d5 | Igfbp7 | fap |
| Igf1 | batch | per | d5 | Igfbp7 | fap |
| Igf1 | batch | ec | d5 | Igfbp2 | fap |
| Igf1 | batch | fap | d5 | Igfbp2 | fap |
| Igf1 | batch | ic | d5 | Igfbp2 | fap |
| Igf1 | batch | per | d5 | Igfbp2 | fap |
| Igf2 | batch | ec | d5 | Vtn | fap |
| Igf2 | batch | fap | d5 | Vtn | fap |
| Igf2 | batch | ic | d5 | Vtn | fap |
| Igf2 | batch | mp | d5 | Vtn | fap |
| Il1b | batch | fap | d5 | Il1r1 | fap |
| Il33 | batch | fap | d5 | Il1rl1 | fap |
| Il33 | batch | mp | d5 | Il1rl1 | fap |
| Il33 | batch | per | d5 | Il1rl1 | fap |
| Mif | batch | fap | d5 | Cd74 | fap |
| Mmp13 | batch | ec | d5 | F2r | fap |
| Mmp13 | batch | fap | d5 | F2r | fap |
| Mmp13 | batch | ic | d5 | F2r | fap |
| Mmp13 | batch | mp | d5 | F2r | fap |
| Mmp13 | batch | per | d5 | F2r | fap |
| Ntf3 | batch | ec | d5 | Ngfr | fap |
| Ntf3 | batch | fap | d5 | Ngfr | fap |
| Ntf3 | constitutive | per | d5 | Ngfr | fap |
| Pdgfa | batch | fap | d5 | Pdgfra | fap |
| Pdgfa | batch | ic | d5 | Pdgfra | fap |
| Pdgfa | constitutive | ec | d5 | Pdgfra | fap |
| Pdgfa | constitutive | mp | d5 | Pdgfra | fap |
| Pdgfa | constitutive | per | d5 | Pdgfra | fap |
| Pthlh | batch | fap | d5 | Pth1r | fap |
| Ptn | batch | fap | d5 | Ptprs | fap |
| Ptn | constitutive | per | d5 | Ptprs | fap |
| S100b | batch | fap | d5 | Fgfr1 | fap |
| S100b | constitutive | per | d5 | Fgfr1 | fap |
| Sfrp1 | batch | fap | d5 | Fzd2 | fap |
| Sfrp1 | constitutive | per | d5 | Fzd2 | fap |
| Sfrp2 | batch | fap | d5 | Fzd2 | fap |
| Sfrp2 | batch | per | d5 | Fzd2 | fap |
| Sfrp4 | batch | fap | d5 | Fzd2 | fap |
| Sfrp4 | batch | per | d5 | Fzd2 | fap |
| Spp1 | batch | fap | d5 | Itga5 | fap |
| Spp1 | batch | ic | d5 | Itga5 | fap |
| Spp1 | batch | mp | d5 | Itga5 | fap |
| Spp1 | batch | fap | d5 | Vtn | fap |
| Spp1 | batch | ic | d5 | Vtn | fap |
| Spp1 | batch | mp | d5 | Vtn | fap |
| Spp1 | batch | fap | d5 | Itgb5 | fap |
| Spp1 | batch | ic | d5 | Itgb5 | fap |
| Spp1 | batch | mp | d5 | Itgb5 | fap |
| Spp1 | batch | fap | d5 | Itgb3 | fap |
| Spp1 | batch | ic | d5 | Itgb3 | fap |
| Spp1 | batch | mp | d5 | Itgb3 | fap |
| Spp1 | batch | fap | d5 | Itgav | fap |
| Spp1 | batch | ic | d5 | Itgav | fap |
| Spp1 | batch | mp | d5 | Itgav | fap |
| Spp1 | batch | fap | d5 | Itga9 | fap |
| Spp1 | batch | ic | d5 | Itga9 | fap |
| Spp1 | batch | mp | d5 | Itga9 | fap |
| Tgfb3 | batch | fap | d5 | Acvrl1 | fap |
| Tgfb3 | total | ec | d5 | Acvrl1 | fap |
| Tgfb3 | total | per | d5 | Acvrl1 | fap |
| Tgfb3 | batch | fap | d5 | Tgfbr2 | fap |
| Tgfb3 | total | ec | d5 | Tgfbr2 | fap |
| Tgfb3 | total | per | d5 | Tgfbr2 | fap |
| Tgfb3 | batch | fap | d5 | Itgav | fap |
| Tgfb3 | total | ec | d5 | Itgav | fap |
| Tgfb3 | total | per | d5 | Itgav | fap |
| Tnfrsf11b | batch | fap | d5 | Vtn | fap |
| Dll1 | batch | ec | d5 | Notch1 | fap |
| Dll1 | total | mp | d5 | Notch1 | fap |
| Dll1 | batch | ec | d5 | Notch3 | fap |
| Dll1 | total | mp | d5 | Notch3 | fap |
| Dll1 | batch | ec | d5 | Notch2 | fap |
| Dll1 | total | mp | d5 | Notch2 | fap |
| Fgf9 | batch | ec | d5 | Fgfr2 | fap |
| Il15 | batch | ec | d5 | Il2rg | fap |
| Il15 | batch | ic | d5 | Il2rg | fap |
| Il15 | batch | ec | d5 | Il15ra | fap |
| Il15 | batch | ic | d5 | Il15ra | fap |
| Il6 | batch | ec | d5 | Il6st | fap |
| Il6 | batch | per | d5 | Il6st | fap |
| Inhbb | batch | ec | d5 | Acvr2a | fap |
| Inhbb | constitutive | fap | d5 | Acvr2a | fap |
| Inhbb | constitutive | per | d5 | Acvr2a | fap |
| Inhbb | batch | ec | d5 | Acvr1 | fap |
| Inhbb | constitutive | fap | d5 | Acvr1 | fap |
| Inhbb | constitutive | per | d5 | Acvr1 | fap |
| Tnfsf10 | batch | ec | d5 | Tnfrsf10b | fap |
| Vegfc | batch | ec | d5 | Nrp1 | fap |
| Vegfc | batch | mp | d5 | Nrp1 | fap |
| Vegfc | batch | ec | d5 | Itga9 | fap |
| Vegfc | batch | mp | d5 | Itga9 | fap |
| Areg | batch | mp | d5 | Egfr | fap |
| Gdnf | batch | mp | d5 | Gfra1 | fap |
| Gdnf | batch | per | d5 | Gfra1 | fap |
| Gdnf | batch | mp | d5 | Gfra2 | fap |
| Gdnf | batch | per | d5 | Gfra2 | fap |
| Gmfb | batch | mp | d5 | Egfr | fap |
| Pdap1 | batch | mp | d5 | Pdgfa | fap |
| Pdgfc | batch | mp | d5 | Pdgfra | fap |
| Pdgfc | constitutive | ic | d5 | Pdgfra | fap |
| Pdgfc | constitutive | per | d5 | Pdgfra | fap |
| Tgfb2 | batch | mp | d5 | Vtn | fap |
| Tgfb2 | constitutive | ec | d5 | Vtn | fap |
| Tgfb2 | constitutive | fap | d5 | Vtn | fap |
| Tgfb2 | constitutive | per | d5 | Vtn | fap |
| Tgfb2 | batch | mp | d5 | Tgfbr3 | fap |
| Tgfb2 | constitutive | ec | d5 | Tgfbr3 | fap |
| Tgfb2 | constitutive | fap | d5 | Tgfbr3 | fap |
| Tgfb2 | constitutive | per | d5 | Tgfbr3 | fap |
| Tgfb2 | batch | mp | d5 | Tgfbr2 | fap |
| Tgfb2 | constitutive | ec | d5 | Tgfbr2 | fap |
| Tgfb2 | constitutive | fap | d5 | Tgfbr2 | fap |
| Tgfb2 | constitutive | per | d5 | Tgfbr2 | fap |
| Csf1 | batch | ic | d5 | Csf1r | fap |
| Lgals3 | batch | ic | d5 | Lgals3bp | fap |
| Mmp12 | batch | ic | d5 | Plaur | fap |
| Mmp12 | batch | per | d5 | Plaur | fap |
| Pdgfb | batch | ic | d5 | Pdgfra | fap |
| Pdgfb | constitutive | ec | d5 | Pdgfra | fap |
| Pdgfb | batch | ic | d5 | Pdgfrb | fap |
| Pdgfb | constitutive | ec | d5 | Pdgfrb | fap |
| Tnfsf13 | batch | ic | d5 | Tnfrsf11b | fap |
| Tnfsf13 | batch | ic | d5 | Tnfrsf12a | fap |
| Tnfsf13 | batch | ic | d5 | Tnfrsf1a | fap |
| Btc | batch | per | d5 | Erbb2 | fap |
| Btc | batch | per | d5 | Egfr | fap |
| Il11 | batch | per | d5 | Il11ra1 | fap |
| Il11 | batch | per | d5 | Il6st | fap |
| Lif | batch | per | d5 | Il6st | fap |
| Lif | constitutive | mp | d5 | Il6st | fap |
| Nov | batch | per | d5 | Itga5 | fap |
| Nov | batch | per | d5 | Notch1 | fap |
| Nov | batch | per | d5 | Itgb3 | fap |
| Nov | batch | per | d5 | Itgav | fap |
| Sfrp5 | batch | per | d5 | Fzd2 | fap |
| Fgf7 | constitutive | fap | d5 | Fgfr2 | fap |
| Fgf7 | constitutive | fap | d5 | Nrp1 | fap |
| Bmp5 | constitutive | ec | d5 | Bmpr1a | fap |
| Bmp5 | constitutive | fap | d5 | Bmpr1a | fap |
| Bmp5 | constitutive | per | d5 | Bmpr1a | fap |
| Jag2 | constitutive | ec | d5 | Notch1 | fap |
| Jag2 | constitutive | ec | d5 | Notch3 | fap |
| Jag2 | constitutive | ec | d5 | Notch2 | fap |
| Cmtm8 | constitutive | ec | d5 | Egfr | fap |
| Edn1 | constitutive | ec | d5 | Ednra | fap |
| Bmp4 | constitutive | ec | d5 | Bmpr1a | fap |
| Bmp4 | constitutive | per | d5 | Bmpr1a | fap |
| Tslp | constitutive | ec | d5 | Crlf2 | fap |
| Il18 | constitutive | ic | d5 | Il1rl2 | fap |
| Il18 | constitutive | mp | d5 | Il1rl2 | fap |
| Serpini1 | constitutive | per | d5 | Plat | fap |
| Ngf | constitutive | per | d5 | Ngfr | fap |
| Bmp6 | constitutive | per | d5 | Bmpr1a | fap |
| Bmp6 | constitutive | per | d5 | Acvr2a | fap |
| Pgf | constitutive | per | d5 | Nrp1 | fap |
| Jag1 | constitutive | per | d5 | Notch1 | fap |
| Jag1 | constitutive | per | d5 | Notch3 | fap |
| Jag1 | constitutive | per | d5 | Notch2 | fap |
| Bmp2 | constitutive | per | d5 | Bmpr1a | fap |
| Bmp2 | constitutive | per | d5 | Acvr2a | fap |
| Ccl5 | batch | fap | d6 | Sdc4 | fap |
| Crlf1 | batch | fap | d6 | Ctf1 | fap |
| Crlf1 | batch | mp | d6 | Ctf1 | fap |
| Cst3 | batch | fap | d6 | Tgfbr2 | fap |
| Cst3 | constitutive | ic | d6 | Tgfbr2 | fap |
| Cst3 | constitutive | per | d6 | Tgfbr2 | fap |
| Cxcl12 | batch | fap | d6 | Cxcr4 | fap |
| Cxcl12 | batch | ic | d6 | Cxcr4 | fap |
| Cxcl12 | batch | mp | d6 | Cxcr4 | fap |
| Cxcl12 | constitutive | ec | d6 | Cxcr4 | fap |
| Cxcl12 | constitutive | per | d6 | Cxcr4 | fap |
| Efnb1 | batch | fap | d6 | Ephb2 | fap |
| Efnb1 | batch | ic | d6 | Ephb2 | fap |
| Efnb1 | batch | mp | d6 | Ephb2 | fap |
| Efnb1 | constitutive | ec | d6 | Ephb2 | fap |
| Efnb1 | constitutive | per | d6 | Ephb2 | fap |
| Fgf1 | batch | fap | d6 | Fgfr2 | fap |
| Fgf1 | constitutive | per | d6 | Fgfr2 | fap |
| Fgf1 | batch | fap | d6 | Fgfr1 | fap |
| Fgf1 | constitutive | per | d6 | Fgfr1 | fap |
| Gas6 | batch | fap | d6 | Axl | fap |
| Gas6 | batch | mp | d6 | Axl | fap |
| Hbegf | batch | fap | d6 | Egfr | fap |
| Hbegf | batch | mp | d6 | Egfr | fap |
| Hbegf | constitutive | ec | d6 | Egfr | fap |
| Hbegf | constitutive | per | d6 | Egfr | fap |
| Igf1 | batch | ec | d6 | Igfbp3 | fap |
| Igf1 | batch | fap | d6 | Igfbp3 | fap |
| Igf1 | batch | ic | d6 | Igfbp3 | fap |
| Igf1 | batch | ec | d6 | Igfbp4 | fap |
| Igf1 | batch | fap | d6 | Igfbp4 | fap |
| Igf1 | batch | ic | d6 | Igfbp4 | fap |
| Igf1 | batch | ec | d6 | Igfbp5 | fap |
| Igf1 | batch | fap | d6 | Igfbp5 | fap |
| Igf1 | batch | ic | d6 | Igfbp5 | fap |
| Igf1 | batch | ec | d6 | Igfbp7 | fap |
| Igf1 | batch | fap | d6 | Igfbp7 | fap |
| Igf1 | batch | ic | d6 | Igfbp7 | fap |
| Igf1 | batch | ec | d6 | Igfbp2 | fap |
| Igf1 | batch | fap | d6 | Igfbp2 | fap |
| Igf1 | batch | ic | d6 | Igfbp2 | fap |
| Igf2 | batch | ec | d6 | Vtn | fap |
| Igf2 | batch | fap | d6 | Vtn | fap |
| Igf2 | batch | ic | d6 | Vtn | fap |
| Igf2 | batch | mp | d6 | Vtn | fap |
| Mmp13 | batch | ec | d6 | F2r | fap |
| Mmp13 | batch | fap | d6 | F2r | fap |
| Mmp13 | batch | ic | d6 | F2r | fap |
| Mmp13 | batch | mp | d6 | F2r | fap |
| Ntf3 | batch | ec | d6 | Ngfr | fap |
| Ntf3 | batch | fap | d6 | Ngfr | fap |
| Ntf3 | batch | mp | d6 | Ngfr | fap |
| Ntf3 | constitutive | per | d6 | Ngfr | fap |
| Pdgfa | batch | fap | d6 | Pdgfra | fap |
| Pdgfa | batch | ic | d6 | Pdgfra | fap |
| Pdgfa | constitutive | ec | d6 | Pdgfra | fap |
| Pdgfa | constitutive | mp | d6 | Pdgfra | fap |
| Pdgfa | constitutive | per | d6 | Pdgfra | fap |
| Pthlh | batch | fap | d6 | Pth1r | fap |
| Ptn | batch | fap | d6 | Ptprs | fap |
| Ptn | constitutive | per | d6 | Ptprs | fap |
| S100b | batch | fap | d6 | Fgfr1 | fap |
| S100b | batch | mp | d6 | Fgfr1 | fap |
| S100b | constitutive | per | d6 | Fgfr1 | fap |
| Sfrp1 | batch | fap | d6 | Fzd2 | fap |
| Sfrp1 | constitutive | per | d6 | Fzd2 | fap |
| Sfrp2 | batch | fap | d6 | Fzd2 | fap |
| Sfrp4 | batch | fap | d6 | Fzd2 | fap |
| Sfrp4 | batch | mp | d6 | Fzd2 | fap |
| Spp1 | batch | fap | d6 | Vtn | fap |
| Spp1 | batch | ic | d6 | Vtn | fap |
| Spp1 | batch | mp | d6 | Vtn | fap |
| Spp1 | batch | fap | d6 | Itgb5 | fap |
| Spp1 | batch | ic | d6 | Itgb5 | fap |
| Spp1 | batch | mp | d6 | Itgb5 | fap |
| Spp1 | batch | fap | d6 | Itgb3 | fap |
| Spp1 | batch | ic | d6 | Itgb3 | fap |
| Spp1 | batch | mp | d6 | Itgb3 | fap |
| Spp1 | batch | fap | d6 | Itgav | fap |
| Spp1 | batch | ic | d6 | Itgav | fap |
| Spp1 | batch | mp | d6 | Itgav | fap |
| Spp1 | batch | fap | d6 | Itga9 | fap |
| Spp1 | batch | ic | d6 | Itga9 | fap |
| Spp1 | batch | mp | d6 | Itga9 | fap |
| Tgfb3 | batch | fap | d6 | Acvrl1 | fap |
| Tgfb3 | batch | mp | d6 | Acvrl1 | fap |
| Tgfb3 | total | ec | d6 | Acvrl1 | fap |
| Tgfb3 | total | per | d6 | Acvrl1 | fap |
| Tgfb3 | batch | fap | d6 | Tgfbr2 | fap |
| Tgfb3 | batch | mp | d6 | Tgfbr2 | fap |
| Tgfb3 | total | ec | d6 | Tgfbr2 | fap |
| Tgfb3 | total | per | d6 | Tgfbr2 | fap |
| Tgfb3 | batch | fap | d6 | Itgav | fap |
| Tgfb3 | batch | mp | d6 | Itgav | fap |
| Tgfb3 | total | ec | d6 | Itgav | fap |
| Tgfb3 | total | per | d6 | Itgav | fap |
| Apln | batch | ec | d6 | Aplnr | fap |
| Dll1 | batch | ec | d6 | Notch1 | fap |
| Dll1 | total | mp | d6 | Notch1 | fap |
| Dll1 | batch | ec | d6 | Notch3 | fap |
| Dll1 | total | mp | d6 | Notch3 | fap |
| Dll1 | batch | ec | d6 | Notch2 | fap |
| Dll1 | total | mp | d6 | Notch2 | fap |
| Fgf9 | batch | ec | d6 | Fgfr2 | fap |
| Il15 | batch | ec | d6 | Il2rg | fap |
| Il15 | batch | ic | d6 | Il2rg | fap |
| Il15 | batch | ec | d6 | Il15ra | fap |
| Il15 | batch | ic | d6 | Il15ra | fap |
| Il6 | batch | ec | d6 | Il6st | fap |
| Inhbb | batch | ec | d6 | Acvr2a | fap |
| Inhbb | batch | mp | d6 | Acvr2a | fap |
| Inhbb | constitutive | fap | d6 | Acvr2a | fap |
| Inhbb | constitutive | per | d6 | Acvr2a | fap |
| Inhbb | batch | ec | d6 | Acvr1 | fap |
| Inhbb | batch | mp | d6 | Acvr1 | fap |
| Inhbb | constitutive | fap | d6 | Acvr1 | fap |
| Inhbb | constitutive | per | d6 | Acvr1 | fap |
| Vegfc | batch | ec | d6 | Nrp1 | fap |
| Vegfc | batch | mp | d6 | Nrp1 | fap |
| Vegfc | batch | ec | d6 | Itga9 | fap |
| Vegfc | batch | mp | d6 | Itga9 | fap |
| Bmp2 | batch | mp | d6 | Bmpr1a | fap |
| Bmp2 | constitutive | per | d6 | Bmpr1a | fap |
| Bmp2 | batch | mp | d6 | Acvr2a | fap |
| Bmp2 | constitutive | per | d6 | Acvr2a | fap |
| Bmp4 | batch | mp | d6 | Bmpr1a | fap |
| Bmp4 | constitutive | ec | d6 | Bmpr1a | fap |
| Bmp4 | constitutive | per | d6 | Bmpr1a | fap |
| Bmp6 | batch | mp | d6 | Bmpr1a | fap |
| Bmp6 | constitutive | per | d6 | Bmpr1a | fap |
| Bmp6 | batch | mp | d6 | Acvr2a | fap |
| Bmp6 | constitutive | per | d6 | Acvr2a | fap |
| Clcf1 | batch | mp | d6 | Crlf1 | fap |
| Clcf1 | batch | mp | d6 | Cntfr | fap |
| Clcf1 | batch | mp | d6 | Esr1 | fap |
| Ctf1 | batch | mp | d6 | Il6st | fap |
| Edn3 | batch | mp | d6 | Ednra | fap |
| Jag1 | batch | mp | d6 | Notch1 | fap |
| Jag1 | constitutive | per | d6 | Notch1 | fap |
| Jag1 | batch | mp | d6 | Notch3 | fap |
| Jag1 | constitutive | per | d6 | Notch3 | fap |
| Jag1 | batch | mp | d6 | Notch2 | fap |
| Jag1 | constitutive | per | d6 | Notch2 | fap |
| Jag2 | batch | mp | d6 | Notch1 | fap |
| Jag2 | constitutive | ec | d6 | Notch1 | fap |
| Jag2 | batch | mp | d6 | Notch3 | fap |
| Jag2 | constitutive | ec | d6 | Notch3 | fap |
| Jag2 | batch | mp | d6 | Notch2 | fap |
| Jag2 | constitutive | ec | d6 | Notch2 | fap |
| Lgals3 | batch | ic | d6 | Lgals3bp | fap |
| Lgals3 | batch | mp | d6 | Lgals3bp | fap |
| Ngf | batch | mp | d6 | Ngfr | fap |
| Ngf | constitutive | per | d6 | Ngfr | fap |
| Nppc | batch | mp | d6 | Npr2 | fap |
| Pdap1 | batch | mp | d6 | Pdgfa | fap |
| Pdgfc | batch | mp | d6 | Pdgfra | fap |
| Pdgfc | constitutive | ic | d6 | Pdgfra | fap |
| Pdgfc | constitutive | per | d6 | Pdgfra | fap |
| Serpini1 | batch | mp | d6 | Plat | fap |
| Serpini1 | constitutive | per | d6 | Plat | fap |
| Sfrp5 | batch | mp | d6 | Fzd2 | fap |
| Tgfb1 | batch | mp | d6 | Vtn | fap |
| Tgfb1 | batch | mp | d6 | Acvrl1 | fap |
| Tgfb1 | batch | mp | d6 | Tgfbr3 | fap |
| Tgfb1 | batch | mp | d6 | Tgfbr2 | fap |
| Tgfb1 | batch | mp | d6 | Itgav | fap |
| Tgfb2 | batch | mp | d6 | Vtn | fap |
| Tgfb2 | constitutive | ec | d6 | Vtn | fap |
| Tgfb2 | constitutive | fap | d6 | Vtn | fap |
| Tgfb2 | constitutive | per | d6 | Vtn | fap |
| Tgfb2 | batch | mp | d6 | Tgfbr3 | fap |
| Tgfb2 | constitutive | ec | d6 | Tgfbr3 | fap |
| Tgfb2 | constitutive | fap | d6 | Tgfbr3 | fap |
| Tgfb2 | constitutive | per | d6 | Tgfbr3 | fap |
| Tgfb2 | batch | mp | d6 | Tgfbr2 | fap |
| Tgfb2 | constitutive | ec | d6 | Tgfbr2 | fap |
| Tgfb2 | constitutive | fap | d6 | Tgfbr2 | fap |
| Tgfb2 | constitutive | per | d6 | Tgfbr2 | fap |
| Tnfsf12 | batch | mp | d6 | Tnfrsf12a | fap |
| Tnfsf12 | batch | mp | d6 | Tnfrsf1a | fap |
| Vegfa | batch | mp | d6 | Vtn | fap |
| Vegfa | batch | mp | d6 | Nrp1 | fap |
| Vegfa | batch | mp | d6 | Ephb2 | fap |
| Vegfa | batch | mp | d6 | Itga9 | fap |
| Csf1 | batch | ic | d6 | Csf1r | fap |
| Hgf | batch | ic | d6 | Vtn | fap |
| Pdgfb | batch | ic | d6 | Pdgfra | fap |
| Pdgfb | constitutive | ec | d6 | Pdgfra | fap |
| Pdgfb | batch | ic | d6 | Pdgfrb | fap |
| Pdgfb | constitutive | ec | d6 | Pdgfrb | fap |
| Tnfsf13 | batch | ic | d6 | Tnfrsf12a | fap |
| Tnfsf13 | batch | ic | d6 | Tnfrsf1a | fap |
| Fgf7 | constitutive | fap | d6 | Fgfr2 | fap |
| Fgf7 | constitutive | fap | d6 | Nrp1 | fap |
| Bmp5 | constitutive | ec | d6 | Bmpr1a | fap |
| Bmp5 | constitutive | fap | d6 | Bmpr1a | fap |
| Bmp5 | constitutive | per | d6 | Bmpr1a | fap |
| Cmtm8 | constitutive | ec | d6 | Egfr | fap |
| Edn1 | constitutive | ec | d6 | Ednra | fap |
| Tslp | constitutive | ec | d6 | Crlf2 | fap |
| Il18 | constitutive | ic | d6 | Il1rl2 | fap |
| Il18 | constitutive | mp | d6 | Il1rl2 | fap |
| Lif | constitutive | mp | d6 | Il6st | fap |
| Pgf | constitutive | per | d6 | Nrp1 | fap |
| Bmp2 | batch | fap | d7 | Bmpr1a | fap |
| Bmp2 | batch | mp | d7 | Bmpr1a | fap |
| Bmp2 | constitutive | per | d7 | Bmpr1a | fap |
| Bmp2 | batch | fap | d7 | Acvr2a | fap |
| Bmp2 | batch | mp | d7 | Acvr2a | fap |
| Bmp2 | constitutive | per | d7 | Acvr2a | fap |
| Bmp4 | batch | fap | d7 | Bmpr1a | fap |
| Bmp4 | batch | mp | d7 | Bmpr1a | fap |
| Bmp4 | constitutive | ec | d7 | Bmpr1a | fap |
| Bmp4 | constitutive | per | d7 | Bmpr1a | fap |
| Bmp6 | batch | ec | d7 | Bmpr1a | fap |
| Bmp6 | batch | fap | d7 | Bmpr1a | fap |
| Bmp6 | batch | mp | d7 | Bmpr1a | fap |
| Bmp6 | constitutive | per | d7 | Bmpr1a | fap |
| Bmp6 | batch | ec | d7 | Acvr2a | fap |
| Bmp6 | batch | fap | d7 | Acvr2a | fap |
| Bmp6 | batch | mp | d7 | Acvr2a | fap |
| Bmp6 | constitutive | per | d7 | Acvr2a | fap |
| Bmp7 | batch | fap | d7 | Bmpr1a | fap |
| Bmp7 | batch | fap | d7 | Acvr2a | fap |
| Bmp7 | batch | fap | d7 | Acvr1 | fap |
| Ccl11 | batch | fap | d7 | Dpp4 | fap |
| Ccl11 | batch | mp | d7 | Dpp4 | fap |
| Clu | batch | ec | d7 | Vldlr | fap |
| Clu | batch | fap | d7 | Vldlr | fap |
| Clu | batch | ic | d7 | Vldlr | fap |
| Clu | batch | mp | d7 | Vldlr | fap |
| Crlf1 | batch | ec | d7 | Ctf1 | fap |
| Crlf1 | batch | fap | d7 | Ctf1 | fap |
| Crlf1 | batch | mp | d7 | Ctf1 | fap |
| Cst3 | batch | fap | d7 | Tgfbr2 | fap |
| Cst3 | constitutive | ic | d7 | Tgfbr2 | fap |
| Cst3 | constitutive | per | d7 | Tgfbr2 | fap |
| Edn1 | batch | fap | d7 | Ednra | fap |
| Edn1 | constitutive | ec | d7 | Ednra | fap |
| Fgf1 | batch | fap | d7 | Fgfr2 | fap |
| Fgf1 | constitutive | per | d7 | Fgfr2 | fap |
| Fgf1 | batch | fap | d7 | Fgfr1 | fap |
| Fgf1 | constitutive | per | d7 | Fgfr1 | fap |
| Gas6 | batch | fap | d7 | Axl | fap |
| Gas6 | batch | mp | d7 | Axl | fap |
| Hbegf | batch | fap | d7 | Egfr | fap |
| Hbegf | batch | mp | d7 | Egfr | fap |
| Hbegf | constitutive | ec | d7 | Egfr | fap |
| Hbegf | constitutive | per | d7 | Egfr | fap |
| Igf1 | batch | ec | d7 | Igfbp3 | fap |
| Igf1 | batch | fap | d7 | Igfbp3 | fap |
| Igf1 | batch | ic | d7 | Igfbp3 | fap |
| Igf1 | batch | ec | d7 | Igfbp4 | fap |
| Igf1 | batch | fap | d7 | Igfbp4 | fap |
| Igf1 | batch | ic | d7 | Igfbp4 | fap |
| Igf1 | batch | ec | d7 | Igfbp5 | fap |
| Igf1 | batch | fap | d7 | Igfbp5 | fap |
| Igf1 | batch | ic | d7 | Igfbp5 | fap |
| Igf1 | batch | ec | d7 | Igfbp6 | fap |
| Igf1 | batch | fap | d7 | Igfbp6 | fap |
| Igf1 | batch | ic | d7 | Igfbp6 | fap |
| Igf1 | batch | ec | d7 | Igfbp7 | fap |
| Igf1 | batch | fap | d7 | Igfbp7 | fap |
| Igf1 | batch | ic | d7 | Igfbp7 | fap |
| Igf1 | batch | ec | d7 | Igfbp2 | fap |
| Igf1 | batch | fap | d7 | Igfbp2 | fap |
| Igf1 | batch | ic | d7 | Igfbp2 | fap |
| Igf2 | batch | ec | d7 | Vtn | fap |
| Igf2 | batch | fap | d7 | Vtn | fap |
| Igf2 | batch | ic | d7 | Vtn | fap |
| Igf2 | batch | mp | d7 | Vtn | fap |
| Il18 | batch | fap | d7 | Il1rl2 | fap |
| Il18 | constitutive | ic | d7 | Il1rl2 | fap |
| Il18 | constitutive | mp | d7 | Il1rl2 | fap |
| Mmp13 | batch | ec | d7 | F2r | fap |
| Mmp13 | batch | fap | d7 | F2r | fap |
| Mmp13 | batch | mp | d7 | F2r | fap |
| Nov | batch | ec | d7 | Itgb3 | fap |
| Nov | batch | fap | d7 | Itgb3 | fap |
| Nov | batch | ec | d7 | Itgav | fap |
| Nov | batch | fap | d7 | Itgav | fap |
| Ntf3 | batch | ec | d7 | Ntrk2 | fap |
| Ntf3 | batch | fap | d7 | Ntrk2 | fap |
| Ntf3 | batch | mp | d7 | Ntrk2 | fap |
| Ntf3 | constitutive | per | d7 | Ntrk2 | fap |
| Ntf3 | batch | ec | d7 | Ngfr | fap |
| Ntf3 | batch | fap | d7 | Ngfr | fap |
| Ntf3 | batch | mp | d7 | Ngfr | fap |
| Ntf3 | constitutive | per | d7 | Ngfr | fap |
| Pdgfa | batch | fap | d7 | Pdgfra | fap |
| Pdgfa | constitutive | ec | d7 | Pdgfra | fap |
| Pdgfa | constitutive | mp | d7 | Pdgfra | fap |
| Pdgfa | constitutive | per | d7 | Pdgfra | fap |
| Pdgfc | batch | fap | d7 | Pdgfra | fap |
| Pdgfc | batch | mp | d7 | Pdgfra | fap |
| Pdgfc | constitutive | ic | d7 | Pdgfra | fap |
| Pdgfc | constitutive | per | d7 | Pdgfra | fap |
| Pgf | batch | fap | d7 | Nrp1 | fap |
| Pgf | constitutive | per | d7 | Nrp1 | fap |
| Pthlh | batch | fap | d7 | Pth1r | fap |
| Ptn | batch | fap | d7 | Ptprs | fap |
| Ptn | constitutive | per | d7 | Ptprs | fap |
| S100b | batch | fap | d7 | Fgfr1 | fap |
| S100b | batch | mp | d7 | Fgfr1 | fap |
| S100b | constitutive | per | d7 | Fgfr1 | fap |
| Sfrp1 | batch | fap | d7 | Fzd2 | fap |
| Sfrp1 | constitutive | per | d7 | Fzd2 | fap |
| Sfrp4 | batch | ec | d7 | Fzd2 | fap |
| Sfrp4 | batch | fap | d7 | Fzd2 | fap |
| Sfrp4 | batch | mp | d7 | Fzd2 | fap |
| Tgfb3 | batch | fap | d7 | Acvrl1 | fap |
| Tgfb3 | batch | mp | d7 | Acvrl1 | fap |
| Tgfb3 | total | ec | d7 | Acvrl1 | fap |
| Tgfb3 | total | per | d7 | Acvrl1 | fap |
| Tgfb3 | batch | fap | d7 | Tgfbr2 | fap |
| Tgfb3 | batch | mp | d7 | Tgfbr2 | fap |
| Tgfb3 | total | ec | d7 | Tgfbr2 | fap |
| Tgfb3 | total | per | d7 | Tgfbr2 | fap |
| Tgfb3 | batch | fap | d7 | Itgav | fap |
| Tgfb3 | batch | mp | d7 | Itgav | fap |
| Tgfb3 | total | ec | d7 | Itgav | fap |
| Tgfb3 | total | per | d7 | Itgav | fap |
| Vegfa | batch | fap | d7 | Vtn | fap |
| Vegfa | batch | mp | d7 | Vtn | fap |
| Vegfa | batch | fap | d7 | Nrp1 | fap |
| Vegfa | batch | mp | d7 | Nrp1 | fap |
| Vegfa | batch | fap | d7 | Ephb2 | fap |
| Vegfa | batch | mp | d7 | Ephb2 | fap |
| Vegfa | batch | fap | d7 | Itga9 | fap |
| Vegfa | batch | mp | d7 | Itga9 | fap |
| Wnt2 | batch | fap | d7 | Sfrp1 | fap |
| Wnt2 | batch | fap | d7 | Fzd1 | fap |
| Cxcl10 | batch | ec | d7 | Dpp4 | fap |
| Cxcl10 | batch | ic | d7 | Dpp4 | fap |
| Cxcl9 | batch | ec | d7 | Dpp4 | fap |
| Cxcl9 | batch | ic | d7 | Dpp4 | fap |
| Dll1 | batch | ec | d7 | Notch2 | fap |
| Dll1 | total | mp | d7 | Notch2 | fap |
| Fgf9 | batch | ec | d7 | Fgfr2 | fap |
| Il15 | batch | ec | d7 | Il15ra | fap |
| Il15 | batch | ic | d7 | Il15ra | fap |
| Sfrp5 | batch | ec | d7 | Fzd2 | fap |
| Sfrp5 | batch | mp | d7 | Fzd2 | fap |
| Vegfc | batch | ec | d7 | Nrp1 | fap |
| Vegfc | batch | mp | d7 | Nrp1 | fap |
| Vegfc | batch | ec | d7 | Itga9 | fap |
| Vegfc | batch | mp | d7 | Itga9 | fap |
| Areg | batch | mp | d7 | Egfr | fap |
| Clcf1 | batch | ic | d7 | Crlf1 | fap |
| Clcf1 | batch | mp | d7 | Crlf1 | fap |
| Clcf1 | batch | ic | d7 | Esr1 | fap |
| Clcf1 | batch | mp | d7 | Esr1 | fap |
| Ctf1 | batch | mp | d7 | Il6st | fap |
| Cxcl12 | batch | ic | d7 | Cxcr4 | fap |
| Cxcl12 | batch | mp | d7 | Cxcr4 | fap |
| Cxcl12 | constitutive | ec | d7 | Cxcr4 | fap |
| Cxcl12 | constitutive | per | d7 | Cxcr4 | fap |
| Cxcl12 | batch | ic | d7 | Dpp4 | fap |
| Cxcl12 | batch | mp | d7 | Dpp4 | fap |
| Cxcl12 | constitutive | ec | d7 | Dpp4 | fap |
| Cxcl12 | constitutive | per | d7 | Dpp4 | fap |
| Edn3 | batch | mp | d7 | Ednra | fap |
| Efnb1 | batch | ic | d7 | Ephb2 | fap |
| Efnb1 | batch | mp | d7 | Ephb2 | fap |
| Efnb1 | constitutive | ec | d7 | Ephb2 | fap |
| Efnb1 | constitutive | per | d7 | Ephb2 | fap |
| Gdnf | batch | mp | d7 | Gfra1 | fap |
| Gdnf | batch | mp | d7 | Ret | fap |
| Gdnf | batch | mp | d7 | Gfra2 | fap |
| Gmfb | batch | mp | d7 | Egfr | fap |
| Inhbb | batch | mp | d7 | Acvr2a | fap |
| Inhbb | constitutive | fap | d7 | Acvr2a | fap |
| Inhbb | constitutive | per | d7 | Acvr2a | fap |
| Inhbb | batch | mp | d7 | Acvr1 | fap |
| Inhbb | constitutive | fap | d7 | Acvr1 | fap |
| Inhbb | constitutive | per | d7 | Acvr1 | fap |
| Jag1 | batch | mp | d7 | Notch2 | fap |
| Jag1 | constitutive | per | d7 | Notch2 | fap |
| Jag2 | batch | mp | d7 | Notch2 | fap |
| Jag2 | constitutive | ec | d7 | Notch2 | fap |
| Ngf | batch | mp | d7 | Ngfr | fap |
| Ngf | constitutive | per | d7 | Ngfr | fap |
| Nppc | batch | mp | d7 | Npr2 | fap |
| Pdap1 | batch | mp | d7 | Pdgfa | fap |
| Serpini1 | batch | mp | d7 | Plat | fap |
| Serpini1 | constitutive | per | d7 | Plat | fap |
| Spp1 | batch | mp | d7 | Vtn | fap |
| Spp1 | batch | mp | d7 | Itgb5 | fap |
| Spp1 | batch | mp | d7 | Itgb3 | fap |
| Spp1 | batch | mp | d7 | Itgav | fap |
| Spp1 | batch | mp | d7 | Itga9 | fap |
| Tgfb1 | batch | mp | d7 | Vtn | fap |
| Tgfb1 | batch | mp | d7 | Acvrl1 | fap |
| Tgfb1 | batch | mp | d7 | Tgfbr3 | fap |
| Tgfb1 | batch | mp | d7 | Tgfbr2 | fap |
| Tgfb1 | batch | mp | d7 | Itgav | fap |
| Tgfb2 | batch | mp | d7 | Vtn | fap |
| Tgfb2 | constitutive | ec | d7 | Vtn | fap |
| Tgfb2 | constitutive | fap | d7 | Vtn | fap |
| Tgfb2 | constitutive | per | d7 | Vtn | fap |
| Tgfb2 | batch | mp | d7 | Tgfbr3 | fap |
| Tgfb2 | constitutive | ec | d7 | Tgfbr3 | fap |
| Tgfb2 | constitutive | fap | d7 | Tgfbr3 | fap |
| Tgfb2 | constitutive | per | d7 | Tgfbr3 | fap |
| Tgfb2 | batch | mp | d7 | Tgfbr2 | fap |
| Tgfb2 | constitutive | ec | d7 | Tgfbr2 | fap |
| Tgfb2 | constitutive | fap | d7 | Tgfbr2 | fap |
| Tgfb2 | constitutive | per | d7 | Tgfbr2 | fap |
| Tnfsf12 | batch | mp | d7 | Tnfrsf1a | fap |
| Ccl22 | batch | ic | d7 | Dpp4 | fap |
| Ifng | batch | ic | d7 | Ifngr1 | fap |
| Il6 | batch | ic | d7 | Il6st | fap |
| Ltb | batch | ic | d7 | Tnfrsf1a | fap |
| Pdgfb | batch | ic | d7 | Pdgfra | fap |
| Pdgfb | constitutive | ec | d7 | Pdgfra | fap |
| Pdgfb | batch | ic | d7 | Pdgfrb | fap |
| Pdgfb | constitutive | ec | d7 | Pdgfrb | fap |
| Angpt1 | constitutive | fap | d7 | Tek | fap |
| Fgf7 | constitutive | fap | d7 | Fgfr2 | fap |
| Fgf7 | constitutive | fap | d7 | Nrp1 | fap |
| Bmp5 | constitutive | ec | d7 | Bmpr1a | fap |
| Bmp5 | constitutive | fap | d7 | Bmpr1a | fap |
| Bmp5 | constitutive | per | d7 | Bmpr1a | fap |
| Cmtm8 | constitutive | ec | d7 | Egfr | fap |
| Angpt2 | constitutive | ec | d7 | Tek | fap |
| Angpt2 | constitutive | ic | d7 | Tek | fap |
| Angpt2 | constitutive | per | d7 | Tek | fap |
| Lif | constitutive | mp | d7 | Il6st | fap |
| Bmp2 | batch | fap | d10 | Bmpr1a | fap |
| Bmp2 | batch | mp | d10 | Bmpr1a | fap |
| Bmp2 | constitutive | per | d10 | Bmpr1a | fap |
| Bmp2 | batch | fap | d10 | Acvr2a | fap |
| Bmp2 | batch | mp | d10 | Acvr2a | fap |
| Bmp2 | constitutive | per | d10 | Acvr2a | fap |
| Bmp4 | batch | fap | d10 | Bmpr1a | fap |
| Bmp4 | batch | mp | d10 | Bmpr1a | fap |
| Bmp4 | constitutive | ec | d10 | Bmpr1a | fap |
| Bmp4 | constitutive | per | d10 | Bmpr1a | fap |
| Bmp6 | batch | fap | d10 | Bmpr1a | fap |
| Bmp6 | batch | mp | d10 | Bmpr1a | fap |
| Bmp6 | constitutive | per | d10 | Bmpr1a | fap |
| Bmp6 | batch | fap | d10 | Acvr2a | fap |
| Bmp6 | batch | mp | d10 | Acvr2a | fap |
| Bmp6 | constitutive | per | d10 | Acvr2a | fap |
| Bmp7 | batch | fap | d10 | Bmpr1a | fap |
| Bmp7 | batch | fap | d10 | Acvr2a | fap |
| Bmp7 | batch | fap | d10 | Acvr1 | fap |
| Ccl11 | batch | fap | d10 | Dpp4 | fap |
| Ccl11 | batch | mp | d10 | Dpp4 | fap |
| Ccl11 | batch | per | d10 | Dpp4 | fap |
| Clu | batch | ec | d10 | Vldlr | fap |
| Clu | batch | fap | d10 | Vldlr | fap |
| Clu | batch | mp | d10 | Vldlr | fap |
| Clu | total | per | d10 | Vldlr | fap |
| Crlf1 | batch | fap | d10 | Ctf1 | fap |
| Crlf1 | batch | mp | d10 | Ctf1 | fap |
| Cst3 | batch | fap | d10 | Tgfbr2 | fap |
| Cst3 | constitutive | ic | d10 | Tgfbr2 | fap |
| Cst3 | constitutive | per | d10 | Tgfbr2 | fap |
| Edn1 | batch | fap | d10 | Ednra | fap |
| Edn1 | constitutive | ec | d10 | Ednra | fap |
| Efnb1 | batch | fap | d10 | Ephb2 | fap |
| Efnb1 | batch | mp | d10 | Ephb2 | fap |
| Efnb1 | constitutive | ec | d10 | Ephb2 | fap |
| Efnb1 | constitutive | per | d10 | Ephb2 | fap |
| Fgf1 | batch | fap | d10 | Fgfr2 | fap |
| Fgf1 | batch | mp | d10 | Fgfr2 | fap |
| Fgf1 | constitutive | per | d10 | Fgfr2 | fap |
| Fgf1 | batch | fap | d10 | Fgfr1 | fap |
| Fgf1 | batch | mp | d10 | Fgfr1 | fap |
| Fgf1 | constitutive | per | d10 | Fgfr1 | fap |
| Gas6 | batch | fap | d10 | Axl | fap |
| Gas6 | batch | mp | d10 | Axl | fap |
| Hbegf | batch | fap | d10 | Egfr | fap |
| Hbegf | batch | mp | d10 | Egfr | fap |
| Hbegf | constitutive | ec | d10 | Egfr | fap |
| Hbegf | constitutive | per | d10 | Egfr | fap |
| Igf1 | batch | ec | d10 | Igfbp3 | fap |
| Igf1 | batch | fap | d10 | Igfbp3 | fap |
| Igf1 | batch | ic | d10 | Igfbp3 | fap |
| Igf1 | batch | ec | d10 | Igfbp4 | fap |
| Igf1 | batch | fap | d10 | Igfbp4 | fap |
| Igf1 | batch | ic | d10 | Igfbp4 | fap |
| Igf1 | batch | ec | d10 | Igfbp5 | fap |
| Igf1 | batch | fap | d10 | Igfbp5 | fap |
| Igf1 | batch | ic | d10 | Igfbp5 | fap |
| Igf1 | batch | ec | d10 | Igfbp6 | fap |
| Igf1 | batch | fap | d10 | Igfbp6 | fap |
| Igf1 | batch | ic | d10 | Igfbp6 | fap |
| Igf1 | batch | ec | d10 | Igfbp7 | fap |
| Igf1 | batch | fap | d10 | Igfbp7 | fap |
| Igf1 | batch | ic | d10 | Igfbp7 | fap |
| Igf1 | batch | ec | d10 | Igfbp2 | fap |
| Igf1 | batch | fap | d10 | Igfbp2 | fap |
| Igf1 | batch | ic | d10 | Igfbp2 | fap |
| Igf2 | batch | ec | d10 | Vtn | fap |
| Igf2 | batch | fap | d10 | Vtn | fap |
| Igf2 | batch | mp | d10 | Vtn | fap |
| Il18 | batch | fap | d10 | Il1rl2 | fap |
| Il18 | constitutive | ic | d10 | Il1rl2 | fap |
| Il18 | constitutive | mp | d10 | Il1rl2 | fap |
| Mdk | batch | fap | d10 | Lrp1 | fap |
| Mdk | batch | mp | d10 | Lrp1 | fap |
| Mmp13 | batch | ec | d10 | F2r | fap |
| Mmp13 | batch | fap | d10 | F2r | fap |
| Mmp13 | batch | mp | d10 | F2r | fap |
| Nov | batch | fap | d10 | Itgb3 | fap |
| Nov | batch | per | d10 | Itgb3 | fap |
| Nov | batch | fap | d10 | Itgav | fap |
| Nov | batch | per | d10 | Itgav | fap |
| Ntf3 | batch | ec | d10 | Ntrk2 | fap |
| Ntf3 | batch | fap | d10 | Ntrk2 | fap |
| Ntf3 | batch | mp | d10 | Ntrk2 | fap |
| Ntf3 | constitutive | per | d10 | Ntrk2 | fap |
| Ntf3 | batch | ec | d10 | Ngfr | fap |
| Ntf3 | batch | fap | d10 | Ngfr | fap |
| Ntf3 | batch | mp | d10 | Ngfr | fap |
| Ntf3 | constitutive | per | d10 | Ngfr | fap |
| Pdgfa | batch | fap | d10 | Pdgfra | fap |
| Pdgfa | constitutive | ec | d10 | Pdgfra | fap |
| Pdgfa | constitutive | mp | d10 | Pdgfra | fap |
| Pdgfa | constitutive | per | d10 | Pdgfra | fap |
| Pdgfc | batch | fap | d10 | Pdgfra | fap |
| Pdgfc | batch | mp | d10 | Pdgfra | fap |
| Pdgfc | constitutive | ic | d10 | Pdgfra | fap |
| Pdgfc | constitutive | per | d10 | Pdgfra | fap |
| Pgf | batch | fap | d10 | Nrp1 | fap |
| Pgf | batch | mp | d10 | Nrp1 | fap |
| Pgf | constitutive | per | d10 | Nrp1 | fap |
| Pthlh | batch | fap | d10 | Pth1r | fap |
| Ptn | batch | fap | d10 | Ptprs | fap |
| Ptn | batch | mp | d10 | Ptprs | fap |
| Ptn | constitutive | per | d10 | Ptprs | fap |
| S100b | batch | fap | d10 | Fgfr1 | fap |
| S100b | batch | mp | d10 | Fgfr1 | fap |
| S100b | constitutive | per | d10 | Fgfr1 | fap |
| Sfrp1 | batch | fap | d10 | Fzd2 | fap |
| Sfrp1 | constitutive | per | d10 | Fzd2 | fap |
| Sfrp4 | batch | fap | d10 | Fzd2 | fap |
| Sfrp4 | batch | mp | d10 | Fzd2 | fap |
| Tgfb3 | batch | fap | d10 | Acvrl1 | fap |
| Tgfb3 | batch | mp | d10 | Acvrl1 | fap |
| Tgfb3 | batch | fap | d10 | Tgfbr2 | fap |
| Tgfb3 | batch | mp | d10 | Tgfbr2 | fap |
| Tgfb3 | batch | fap | d10 | Itgav | fap |
| Tgfb3 | batch | mp | d10 | Itgav | fap |
| Tslp | batch | fap | d10 | Crlf2 | fap |
| Tslp | constitutive | ec | d10 | Crlf2 | fap |
| Vegfa | batch | fap | d10 | Vtn | fap |
| Vegfa | batch | ic | d10 | Vtn | fap |
| Vegfa | batch | mp | d10 | Vtn | fap |
| Vegfa | batch | fap | d10 | Nrp1 | fap |
| Vegfa | batch | ic | d10 | Nrp1 | fap |
| Vegfa | batch | mp | d10 | Nrp1 | fap |
| Vegfa | batch | fap | d10 | Ephb2 | fap |
| Vegfa | batch | ic | d10 | Ephb2 | fap |
| Vegfa | batch | mp | d10 | Ephb2 | fap |
| Vegfa | batch | fap | d10 | Itga9 | fap |
| Vegfa | batch | ic | d10 | Itga9 | fap |
| Vegfa | batch | mp | d10 | Itga9 | fap |
| Wnt2 | batch | fap | d10 | Sfrp1 | fap |
| Wnt2 | batch | fap | d10 | Fzd1 | fap |
| Cxcl10 | batch | ec | d10 | Dpp4 | fap |
| Cxcl10 | batch | ic | d10 | Dpp4 | fap |
| Cxcl9 | batch | ec | d10 | Dpp4 | fap |
| Cxcl9 | batch | ic | d10 | Dpp4 | fap |
| Dll1 | batch | ec | d10 | Notch2 | fap |
| Fgf9 | batch | ec | d10 | Fgfr2 | fap |
| Il15 | batch | ec | d10 | Il17ra | fap |
| Il15 | batch | ic | d10 | Il17ra | fap |
| Il15 | batch | ec | d10 | Il15ra | fap |
| Il15 | batch | ic | d10 | Il15ra | fap |
| Vegfc | batch | ec | d10 | Nrp1 | fap |
| Vegfc | batch | ec | d10 | Itga9 | fap |
| Clcf1 | batch | ic | d10 | Crlf1 | fap |
| Clcf1 | batch | mp | d10 | Crlf1 | fap |
| Clcf1 | batch | ic | d10 | Cntfr | fap |
| Clcf1 | batch | mp | d10 | Cntfr | fap |
| Clcf1 | batch | ic | d10 | Esr1 | fap |
| Clcf1 | batch | mp | d10 | Esr1 | fap |
| Ctf1 | batch | mp | d10 | Il6st | fap |
| Cxcl12 | batch | mp | d10 | Cxcr4 | fap |
| Cxcl12 | constitutive | ec | d10 | Cxcr4 | fap |
| Cxcl12 | constitutive | per | d10 | Cxcr4 | fap |
| Cxcl12 | batch | mp | d10 | Dpp4 | fap |
| Cxcl12 | constitutive | ec | d10 | Dpp4 | fap |
| Cxcl12 | constitutive | per | d10 | Dpp4 | fap |
| Edn3 | batch | mp | d10 | Ednra | fap |
| Inhbb | batch | mp | d10 | Acvr2a | fap |
| Inhbb | constitutive | fap | d10 | Acvr2a | fap |
| Inhbb | constitutive | per | d10 | Acvr2a | fap |
| Inhbb | batch | mp | d10 | Acvr1 | fap |
| Inhbb | constitutive | fap | d10 | Acvr1 | fap |
| Inhbb | constitutive | per | d10 | Acvr1 | fap |
| Jag1 | batch | ic | d10 | Notch2 | fap |
| Jag1 | batch | mp | d10 | Notch2 | fap |
| Jag1 | constitutive | per | d10 | Notch2 | fap |
| Jag2 | batch | mp | d10 | Notch2 | fap |
| Jag2 | constitutive | ec | d10 | Notch2 | fap |
| Ngf | batch | mp | d10 | Ngfr | fap |
| Ngf | constitutive | per | d10 | Ngfr | fap |
| Nppc | batch | mp | d10 | Npr2 | fap |
| Serpini1 | batch | mp | d10 | Plat | fap |
| Serpini1 | constitutive | per | d10 | Plat | fap |
| Sfrp2 | batch | mp | d10 | Fzd2 | fap |
| Sfrp5 | batch | mp | d10 | Fzd2 | fap |
| Sfrp5 | batch | per | d10 | Fzd2 | fap |
| Tgfb2 | batch | mp | d10 | Vtn | fap |
| Tgfb2 | constitutive | ec | d10 | Vtn | fap |
| Tgfb2 | constitutive | fap | d10 | Vtn | fap |
| Tgfb2 | constitutive | per | d10 | Vtn | fap |
| Tgfb2 | batch | mp | d10 | Tgfbr3 | fap |
| Tgfb2 | constitutive | ec | d10 | Tgfbr3 | fap |
| Tgfb2 | constitutive | fap | d10 | Tgfbr3 | fap |
| Tgfb2 | constitutive | per | d10 | Tgfbr3 | fap |
| Tgfb2 | batch | mp | d10 | Tgfbr2 | fap |
| Tgfb2 | constitutive | ec | d10 | Tgfbr2 | fap |
| Tgfb2 | constitutive | fap | d10 | Tgfbr2 | fap |
| Tgfb2 | constitutive | per | d10 | Tgfbr2 | fap |
| Tnfsf12 | batch | mp | d10 | Tnfrsf12a | fap |
| Tnfsf12 | total | ec | d10 | Tnfrsf12a | fap |
| Tnfsf12 | total | ic | d10 | Tnfrsf12a | fap |
| Tnfsf12 | total | per | d10 | Tnfrsf12a | fap |
| Tnfsf12 | batch | mp | d10 | Tnfrsf1a | fap |
| Tnfsf12 | total | ec | d10 | Tnfrsf1a | fap |
| Tnfsf12 | total | ic | d10 | Tnfrsf1a | fap |
| Tnfsf12 | total | per | d10 | Tnfrsf1a | fap |
| Vegfb | batch | mp | d10 | Nrp1 | fap |
| Areg | batch | ic | d10 | Egfr | fap |
| Ccl22 | batch | ic | d10 | Dpp4 | fap |
| Ccl5 | batch | ic | d10 | Sdc4 | fap |
| Cxcl2 | batch | ic | d10 | Dpp4 | fap |
| Ifng | batch | ic | d10 | Ifngr1 | fap |
| Il1a | batch | ic | d10 | Il1r2 | fap |
| Il1b | batch | ic | d10 | Il1r2 | fap |
| Il6 | batch | ic | d10 | Il6st | fap |
| Lif | batch | ic | d10 | Il6st | fap |
| Lif | constitutive | mp | d10 | Il6st | fap |
| Ltb | batch | ic | d10 | Tnfrsf1a | fap |
| Osm | batch | ic | d10 | Osmr | fap |
| Osm | batch | ic | d10 | Il6st | fap |
| Pdgfb | batch | ic | d10 | Lrp1 | fap |
| Pdgfb | constitutive | ec | d10 | Lrp1 | fap |
| Pdgfb | batch | ic | d10 | Pdgfra | fap |
| Pdgfb | constitutive | ec | d10 | Pdgfra | fap |
| Pdgfb | batch | ic | d10 | Pdgfrb | fap |
| Pdgfb | constitutive | ec | d10 | Pdgfrb | fap |
| Pf4 | batch | ic | d10 | Thbd | fap |
| Ppbp | batch | ic | d10 | Gabbr1 | fap |
| Ppbp | batch | ic | d10 | Slc1a5 | fap |
| Ppbp | batch | ic | d10 | Itgb5 | fap |
| Tnf | batch | ic | d10 | Tnfrsf1a | fap |
| Adm | batch | per | d10 | Calcrl | fap |
| Angpt4 | batch | per | d10 | Tek | fap |
| Gdnf | batch | per | d10 | Gfra1 | fap |
| Gdnf | batch | per | d10 | Ret | fap |
| Gdnf | batch | per | d10 | Gfra2 | fap |
| Tgfb1 | total | ec | d10 | Vtn | fap |
| Tgfb1 | total | ic | d10 | Vtn | fap |
| Tgfb1 | total | ec | d10 | Acvrl1 | fap |
| Tgfb1 | total | ic | d10 | Acvrl1 | fap |
| Tgfb1 | total | ec | d10 | Tgfbr3 | fap |
| Tgfb1 | total | ic | d10 | Tgfbr3 | fap |
| Tgfb1 | total | ec | d10 | Tgfbr2 | fap |
| Tgfb1 | total | ic | d10 | Tgfbr2 | fap |
| Tgfb1 | total | ec | d10 | Itgav | fap |
| Tgfb1 | total | ic | d10 | Itgav | fap |
| Angpt1 | constitutive | fap | d10 | Tek | fap |
| Fgf7 | constitutive | fap | d10 | Fgfr2 | fap |
| Fgf7 | constitutive | fap | d10 | Nrp1 | fap |
| Bmp5 | constitutive | ec | d10 | Bmpr1a | fap |
| Bmp5 | constitutive | fap | d10 | Bmpr1a | fap |
| Bmp5 | constitutive | per | d10 | Bmpr1a | fap |
| Cmtm8 | constitutive | ec | d10 | Egfr | fap |
| Angpt2 | constitutive | ec | d10 | Tek | fap |
| Angpt2 | constitutive | ic | d10 | Tek | fap |
| Angpt2 | constitutive | per | d10 | Tek | fap |
| Bmp2 | batch | fap | d14 | Bmpr1a | fap |
| Bmp2 | constitutive | per | d14 | Bmpr1a | fap |
| Bmp2 | batch | fap | d14 | Acvr2a | fap |
| Bmp2 | constitutive | per | d14 | Acvr2a | fap |
| Bmp4 | batch | fap | d14 | Bmpr1a | fap |
| Bmp4 | constitutive | ec | d14 | Bmpr1a | fap |
| Bmp4 | constitutive | per | d14 | Bmpr1a | fap |
| Bmp6 | batch | fap | d14 | Bmpr1a | fap |
| Bmp6 | constitutive | per | d14 | Bmpr1a | fap |
| Bmp6 | batch | fap | d14 | Acvr2a | fap |
| Bmp6 | constitutive | per | d14 | Acvr2a | fap |
| Bmp7 | batch | fap | d14 | Bmpr1a | fap |
| Bmp7 | batch | fap | d14 | Acvr2a | fap |
| Bmp7 | batch | fap | d14 | Acvr1 | fap |
| Ccl11 | batch | fap | d14 | Dpp4 | fap |
| Clu | batch | ec | d14 | Vldlr | fap |
| Clu | batch | fap | d14 | Vldlr | fap |
| Clu | batch | ic | d14 | Vldlr | fap |
| Cst3 | batch | fap | d14 | Tgfbr2 | fap |
| Cst3 | constitutive | ic | d14 | Tgfbr2 | fap |
| Cst3 | constitutive | per | d14 | Tgfbr2 | fap |
| Edn1 | batch | fap | d14 | Ednra | fap |
| Edn1 | constitutive | ec | d14 | Ednra | fap |
| Efnb1 | batch | fap | d14 | Ephb2 | fap |
| Efnb1 | batch | ic | d14 | Ephb2 | fap |
| Efnb1 | constitutive | ec | d14 | Ephb2 | fap |
| Efnb1 | constitutive | per | d14 | Ephb2 | fap |
| Fgf1 | batch | fap | d14 | Fgfr2 | fap |
| Fgf1 | constitutive | per | d14 | Fgfr2 | fap |
| Fgf1 | batch | fap | d14 | Fgfr1 | fap |
| Fgf1 | constitutive | per | d14 | Fgfr1 | fap |
| Gas6 | batch | fap | d14 | Axl | fap |
| Hbegf | batch | fap | d14 | Egfr | fap |
| Hbegf | constitutive | ec | d14 | Egfr | fap |
| Hbegf | constitutive | per | d14 | Egfr | fap |
| Igf1 | batch | ec | d14 | Igfbp3 | fap |
| Igf1 | batch | fap | d14 | Igfbp3 | fap |
| Igf1 | batch | ic | d14 | Igfbp3 | fap |
| Igf1 | batch | ec | d14 | Igfbp4 | fap |
| Igf1 | batch | fap | d14 | Igfbp4 | fap |
| Igf1 | batch | ic | d14 | Igfbp4 | fap |
| Igf1 | batch | ec | d14 | Igfbp5 | fap |
| Igf1 | batch | fap | d14 | Igfbp5 | fap |
| Igf1 | batch | ic | d14 | Igfbp5 | fap |
| Igf1 | batch | ec | d14 | Igfbp6 | fap |
| Igf1 | batch | fap | d14 | Igfbp6 | fap |
| Igf1 | batch | ic | d14 | Igfbp6 | fap |
| Igf1 | batch | ec | d14 | Igfbp7 | fap |
| Igf1 | batch | fap | d14 | Igfbp7 | fap |
| Igf1 | batch | ic | d14 | Igfbp7 | fap |
| Igf1 | batch | ec | d14 | Igfbp2 | fap |
| Igf1 | batch | fap | d14 | Igfbp2 | fap |
| Igf1 | batch | ic | d14 | Igfbp2 | fap |
| Igf2 | batch | ec | d14 | Vtn | fap |
| Igf2 | batch | fap | d14 | Vtn | fap |
| Igf2 | batch | ic | d14 | Vtn | fap |
| Il18 | batch | fap | d14 | Il1rl2 | fap |
| Il18 | constitutive | ic | d14 | Il1rl2 | fap |
| Il18 | constitutive | mp | d14 | Il1rl2 | fap |
| Mmp13 | batch | fap | d14 | F2r | fap |
| Nov | batch | fap | d14 | Itgb3 | fap |
| Nov | batch | fap | d14 | Itgav | fap |
| Ntf3 | batch | ec | d14 | Ntrk2 | fap |
| Ntf3 | batch | fap | d14 | Ntrk2 | fap |
| Ntf3 | constitutive | per | d14 | Ntrk2 | fap |
| Ntf3 | batch | ec | d14 | Ngfr | fap |
| Ntf3 | batch | fap | d14 | Ngfr | fap |
| Ntf3 | constitutive | per | d14 | Ngfr | fap |
| Pdgfa | batch | fap | d14 | Pdgfra | fap |
| Pdgfa | constitutive | ec | d14 | Pdgfra | fap |
| Pdgfa | constitutive | mp | d14 | Pdgfra | fap |
| Pdgfa | constitutive | per | d14 | Pdgfra | fap |
| Pdgfc | batch | fap | d14 | Pdgfra | fap |
| Pdgfc | constitutive | ic | d14 | Pdgfra | fap |
| Pdgfc | constitutive | per | d14 | Pdgfra | fap |
| Pgf | batch | fap | d14 | Nrp1 | fap |
| Pgf | constitutive | per | d14 | Nrp1 | fap |
| Pthlh | batch | fap | d14 | Pth1r | fap |
| Ptn | batch | fap | d14 | Ptprs | fap |
| Ptn | constitutive | per | d14 | Ptprs | fap |
| S100b | batch | fap | d14 | Fgfr1 | fap |
| S100b | constitutive | per | d14 | Fgfr1 | fap |
| Sfrp1 | batch | fap | d14 | Fzd2 | fap |
| Sfrp1 | constitutive | per | d14 | Fzd2 | fap |
| Sfrp4 | batch | fap | d14 | Fzd2 | fap |
| Tgfb3 | batch | fap | d14 | Acvrl1 | fap |
| Tgfb3 | batch | fap | d14 | Tgfbr2 | fap |
| Tgfb3 | batch | fap | d14 | Itgav | fap |
| Tslp | batch | fap | d14 | Crlf2 | fap |
| Tslp | constitutive | ec | d14 | Crlf2 | fap |
| Vegfa | batch | fap | d14 | Vtn | fap |
| Vegfa | batch | fap | d14 | Nrp1 | fap |
| Vegfa | batch | fap | d14 | Ephb2 | fap |
| Vegfa | batch | fap | d14 | Itga9 | fap |
| Wnt11 | batch | fap | d14 | Fzd4 | fap |
| Wnt2 | batch | fap | d14 | Sfrp1 | fap |
| Wnt2 | batch | fap | d14 | Fzd1 | fap |
| Cxcl10 | batch | ec | d14 | Dpp4 | fap |
| Cxcl10 | batch | ic | d14 | Dpp4 | fap |
| Cxcl9 | batch | ec | d14 | Dpp4 | fap |
| Cxcl9 | batch | ic | d14 | Dpp4 | fap |
| Dll1 | batch | ec | d14 | Notch2 | fap |
| Fgf9 | batch | ec | d14 | Fgfr2 | fap |
| Il15 | batch | ec | d14 | Il15ra | fap |
| Il15 | batch | ic | d14 | Il15ra | fap |
| Vegfc | batch | ec | d14 | Nrp1 | fap |
| Vegfc | batch | ec | d14 | Itga9 | fap |
| Cxcl12 | batch | ic | d14 | Cxcr4 | fap |
| Cxcl12 | constitutive | ec | d14 | Cxcr4 | fap |
| Cxcl12 | constitutive | per | d14 | Cxcr4 | fap |
| Cxcl12 | batch | ic | d14 | Dpp4 | fap |
| Cxcl12 | constitutive | ec | d14 | Dpp4 | fap |
| Cxcl12 | constitutive | per | d14 | Dpp4 | fap |
| Pdgfb | batch | ic | d14 | Pdgfra | fap |
| Pdgfb | constitutive | ec | d14 | Pdgfra | fap |
| Pdgfb | batch | ic | d14 | Pdgfrb | fap |
| Pdgfb | constitutive | ec | d14 | Pdgfrb | fap |
| Angpt1 | constitutive | fap | d14 | Tek | fap |
| Fgf7 | constitutive | fap | d14 | Fgfr2 | fap |
| Fgf7 | constitutive | fap | d14 | Nrp1 | fap |
| Tgfb2 | constitutive | ec | d14 | Vtn | fap |
| Tgfb2 | constitutive | fap | d14 | Vtn | fap |
| Tgfb2 | constitutive | per | d14 | Vtn | fap |
| Tgfb2 | constitutive | ec | d14 | Tgfbr3 | fap |
| Tgfb2 | constitutive | fap | d14 | Tgfbr3 | fap |
| Tgfb2 | constitutive | per | d14 | Tgfbr3 | fap |
| Tgfb2 | constitutive | ec | d14 | Tgfbr2 | fap |
| Tgfb2 | constitutive | fap | d14 | Tgfbr2 | fap |
| Tgfb2 | constitutive | per | d14 | Tgfbr2 | fap |
| Bmp5 | constitutive | ec | d14 | Bmpr1a | fap |
| Bmp5 | constitutive | fap | d14 | Bmpr1a | fap |
| Bmp5 | constitutive | per | d14 | Bmpr1a | fap |
| Inhbb | constitutive | fap | d14 | Acvr2a | fap |
| Inhbb | constitutive | per | d14 | Acvr2a | fap |
| Inhbb | constitutive | fap | d14 | Acvr1 | fap |
| Inhbb | constitutive | per | d14 | Acvr1 | fap |
| Jag2 | constitutive | ec | d14 | Notch2 | fap |
| Cmtm8 | constitutive | ec | d14 | Egfr | fap |
| Angpt2 | constitutive | ec | d14 | Tek | fap |
| Angpt2 | constitutive | ic | d14 | Tek | fap |
| Angpt2 | constitutive | per | d14 | Tek | fap |
| Lif | constitutive | mp | d14 | Il6st | fap |
| Serpini1 | constitutive | per | d14 | Plat | fap |
| Ngf | constitutive | per | d14 | Ngfr | fap |
| Jag1 | constitutive | per | d14 | Notch2 | fap |
| Bmp2 | batch | fap | d0 | Acvr2a | mp |
| Bmp2 | batch | mp | d0 | Acvr2a | mp |
| Bmp2 | constitutive | per | d0 | Acvr2a | mp |
| Bmp2 | batch | fap | d0 | Bmpr1a | mp |
| Bmp2 | batch | mp | d0 | Bmpr1a | mp |
| Bmp2 | constitutive | per | d0 | Bmpr1a | mp |
| Bmp2 | batch | fap | d0 | Eng | mp |
| Bmp2 | batch | mp | d0 | Eng | mp |
| Bmp2 | constitutive | per | d0 | Eng | mp |
| Bmp4 | batch | fap | d0 | Bmpr1a | mp |
| Bmp4 | batch | mp | d0 | Bmpr1a | mp |
| Bmp4 | constitutive | ec | d0 | Bmpr1a | mp |
| Bmp4 | constitutive | per | d0 | Bmpr1a | mp |
| Bmp6 | batch | ec | d0 | Acvr2a | mp |
| Bmp6 | batch | fap | d0 | Acvr2a | mp |
| Bmp6 | batch | mp | d0 | Acvr2a | mp |
| Bmp6 | constitutive | per | d0 | Acvr2a | mp |
| Bmp6 | batch | ec | d0 | Bmpr1a | mp |
| Bmp6 | batch | fap | d0 | Bmpr1a | mp |
| Bmp6 | batch | mp | d0 | Bmpr1a | mp |
| Bmp6 | constitutive | per | d0 | Bmpr1a | mp |
| Bmp7 | batch | fap | d0 | Acvr2a | mp |
| Bmp7 | batch | fap | d0 | Bmpr1a | mp |
| Bmp7 | batch | fap | d0 | Eng | mp |
| Clu | batch | ec | d0 | Vldlr | mp |
| Clu | batch | fap | d0 | Vldlr | mp |
| Clu | batch | ic | d0 | Vldlr | mp |
| Clu | batch | mp | d0 | Vldlr | mp |
| Crlf1 | batch | ec | d0 | Ctf1 | mp |
| Crlf1 | batch | fap | d0 | Ctf1 | mp |
| Crlf1 | batch | mp | d0 | Ctf1 | mp |
| Edn1 | batch | fap | d0 | Ednrb | mp |
| Edn1 | constitutive | ec | d0 | Ednrb | mp |
| Fgf1 | batch | fap | d0 | Fgfr1 | mp |
| Fgf1 | constitutive | per | d0 | Fgfr1 | mp |
| Fgf1 | batch | fap | d0 | Fgfr4 | mp |
| Fgf1 | constitutive | per | d0 | Fgfr4 | mp |
| Fgf18 | batch | fap | d0 | Fgfr4 | mp |
| Fgf18 | batch | per | d0 | Fgfr4 | mp |
| Gas6 | batch | fap | d0 | Axl | mp |
| Gas6 | batch | mp | d0 | Axl | mp |
| Igf1 | batch | ec | d0 | Igfbp4 | mp |
| Igf1 | batch | fap | d0 | Igfbp4 | mp |
| Igf1 | batch | ic | d0 | Igfbp4 | mp |
| Igf1 | batch | ec | d0 | Igf1r | mp |
| Igf1 | batch | fap | d0 | Igf1r | mp |
| Igf1 | batch | ic | d0 | Igf1r | mp |
| Kitl | batch | fap | d0 | Kit | mp |
| Kitl | batch | ic | d0 | Kit | mp |
| Kitl | batch | mp | d0 | Kit | mp |
| Kitl | constitutive | ec | d0 | Kit | mp |
| Kitl | constitutive | per | d0 | Kit | mp |
| Mdk | batch | fap | d0 | Ptprz1 | mp |
| Mdk | batch | mp | d0 | Ptprz1 | mp |
| Nov | batch | ec | d0 | Notch1 | mp |
| Nov | batch | fap | d0 | Notch1 | mp |
| Nov | batch | per | d0 | Notch1 | mp |
| Ntf3 | batch | ec | d0 | Ntrk2 | mp |
| Ntf3 | batch | fap | d0 | Ntrk2 | mp |
| Ntf3 | batch | mp | d0 | Ntrk2 | mp |
| Ntf3 | constitutive | per | d0 | Ntrk2 | mp |
| Ntf3 | batch | ec | d0 | Ngfr | mp |
| Ntf3 | batch | fap | d0 | Ngfr | mp |
| Ntf3 | batch | mp | d0 | Ngfr | mp |
| Ntf3 | constitutive | per | d0 | Ngfr | mp |
| Pgf | batch | fap | d0 | Flt1 | mp |
| Pgf | constitutive | per | d0 | Flt1 | mp |
| Pthlh | batch | fap | d0 | Pth1r | mp |
| Ptn | batch | fap | d0 | Ptprz1 | mp |
| Ptn | constitutive | per | d0 | Ptprz1 | mp |
| Rspo3 | batch | fap | d0 | Lrp6 | mp |
| S100b | batch | fap | d0 | Fgfr1 | mp |
| S100b | batch | mp | d0 | Fgfr1 | mp |
| S100b | constitutive | per | d0 | Fgfr1 | mp |
| Vegfa | batch | fap | d0 | Flt1 | mp |
| Vegfa | batch | mp | d0 | Flt1 | mp |
| Vegfa | batch | fap | d0 | Kdr | mp |
| Vegfa | batch | mp | d0 | Kdr | mp |
| Vegfa | batch | fap | d0 | Vtn | mp |
| Vegfa | batch | mp | d0 | Vtn | mp |
| Wnt11 | batch | fap | d0 | Fzd4 | mp |
| Wnt11 | batch | ic | d0 | Fzd4 | mp |
| Wnt2 | batch | fap | d0 | Sfrp1 | mp |
| Wnt2 | batch | fap | d0 | Fzd1 | mp |
| Dll1 | batch | ec | d0 | Notch1 | mp |
| Dll1 | batch | ec | d0 | Notch3 | mp |
| Dll1 | batch | ec | d0 | Notch2 | mp |
| Fgf9 | batch | ec | d0 | Fgfr4 | mp |
| Igf2 | batch | ec | d0 | Vtn | mp |
| Igf2 | batch | ec | d0 | Igf1r | mp |
| Il15 | batch | ec | d0 | Il17ra | mp |
| Il15 | batch | ic | d0 | Il17ra | mp |
| Vegfc | batch | ec | d0 | Kdr | mp |
| Vegfc | batch | mp | d0 | Kdr | mp |
| Areg | batch | mp | d0 | Egfr | mp |
| Clcf1 | batch | ic | d0 | Crlf1 | mp |
| Clcf1 | batch | mp | d0 | Crlf1 | mp |
| Clcf1 | batch | ic | d0 | Cntfr | mp |
| Clcf1 | batch | mp | d0 | Cntfr | mp |
| Clcf1 | batch | ic | d0 | Esr1 | mp |
| Clcf1 | batch | mp | d0 | Esr1 | mp |
| Ctsg | batch | ic | d0 | Met | mp |
| Ctsg | batch | mp | d0 | Met | mp |
| Edn3 | batch | mp | d0 | Ednrb | mp |
| Gdnf | batch | mp | d0 | Gfra1 | mp |
| Gdnf | batch | per | d0 | Gfra1 | mp |
| Gdnf | batch | mp | d0 | Gfra2 | mp |
| Gdnf | batch | per | d0 | Gfra2 | mp |
| Gdnf | batch | mp | d0 | Ret | mp |
| Gdnf | batch | per | d0 | Ret | mp |
| Gmfb | batch | mp | d0 | Egfr | mp |
| Hbegf | batch | mp | d0 | Egfr | mp |
| Hbegf | constitutive | ec | d0 | Egfr | mp |
| Hbegf | constitutive | per | d0 | Egfr | mp |
| Inhbb | batch | mp | d0 | Acvr2a | mp |
| Inhbb | constitutive | fap | d0 | Acvr2a | mp |
| Inhbb | constitutive | per | d0 | Acvr2a | mp |
| Jag1 | batch | mp | d0 | Notch1 | mp |
| Jag1 | constitutive | per | d0 | Notch1 | mp |
| Jag1 | batch | mp | d0 | Notch3 | mp |
| Jag1 | constitutive | per | d0 | Notch3 | mp |
| Jag1 | batch | mp | d0 | Notch2 | mp |
| Jag1 | constitutive | per | d0 | Notch2 | mp |
| Jag2 | batch | mp | d0 | Notch1 | mp |
| Jag2 | constitutive | ec | d0 | Notch1 | mp |
| Jag2 | batch | mp | d0 | Notch3 | mp |
| Jag2 | constitutive | ec | d0 | Notch3 | mp |
| Jag2 | batch | mp | d0 | Notch2 | mp |
| Jag2 | constitutive | ec | d0 | Notch2 | mp |
| Ngf | batch | mp | d0 | Ror1 | mp |
| Ngf | constitutive | per | d0 | Ror1 | mp |
| Ngf | batch | mp | d0 | Sort1 | mp |
| Ngf | constitutive | per | d0 | Sort1 | mp |
| Ngf | batch | mp | d0 | Ngfr | mp |
| Ngf | constitutive | per | d0 | Ngfr | mp |
| Nppc | batch | mp | d0 | Npr2 | mp |
| Tnfsf12 | batch | mp | d0 | Tnfrsf12a | mp |
| Ccl5 | batch | ic | d0 | Sdc4 | mp |
| Mmp9 | batch | ic | d0 | Flt1 | mp |
| Adm | batch | per | d0 | Calcrl | mp |
| Angpt4 | batch | per | d0 | Tek | mp |
| Dll4 | total | ec | d0 | Notch1 | mp |
| Angpt1 | constitutive | fap | d0 | Tek | mp |
| Fgf7 | constitutive | fap | d0 | Fgfr4 | mp |
| Tgfb2 | constitutive | ec | d0 | Tgfbr3 | mp |
| Tgfb2 | constitutive | fap | d0 | Tgfbr3 | mp |
| Tgfb2 | constitutive | per | d0 | Tgfbr3 | mp |
| Tgfb2 | constitutive | ec | d0 | Vtn | mp |
| Tgfb2 | constitutive | fap | d0 | Vtn | mp |
| Tgfb2 | constitutive | per | d0 | Vtn | mp |
| Bmp5 | constitutive | ec | d0 | Bmpr1a | mp |
| Bmp5 | constitutive | fap | d0 | Bmpr1a | mp |
| Bmp5 | constitutive | per | d0 | Bmpr1a | mp |
| Cmtm8 | constitutive | ec | d0 | Egfr | mp |
| Dhh | constitutive | ec | d0 | Ptch1 | mp |
| Dhh | constitutive | mp | d0 | Ptch1 | mp |
| Dhh | constitutive | per | d0 | Ptch1 | mp |
| Angpt2 | constitutive | ec | d0 | Tek | mp |
| Angpt2 | constitutive | ic | d0 | Tek | mp |
| Angpt2 | constitutive | per | d0 | Tek | mp |
| Bmp2 | batch | ec | d1 | Acvr2a | mp |
| Bmp2 | batch | fap | d1 | Acvr2a | mp |
| Bmp2 | batch | mp | d1 | Acvr2a | mp |
| Bmp2 | constitutive | per | d1 | Acvr2a | mp |
| Bmp2 | batch | ec | d1 | Bmpr1a | mp |
| Bmp2 | batch | fap | d1 | Bmpr1a | mp |
| Bmp2 | batch | mp | d1 | Bmpr1a | mp |
| Bmp2 | constitutive | per | d1 | Bmpr1a | mp |
| Bmp2 | batch | ec | d1 | Eng | mp |
| Bmp2 | batch | fap | d1 | Eng | mp |
| Bmp2 | batch | mp | d1 | Eng | mp |
| Bmp2 | constitutive | per | d1 | Eng | mp |
| Bmp6 | batch | fap | d1 | Acvr2a | mp |
| Bmp6 | constitutive | per | d1 | Acvr2a | mp |
| Bmp6 | batch | fap | d1 | Bmpr1a | mp |
| Bmp6 | constitutive | per | d1 | Bmpr1a | mp |
| Ccl2 | batch | ec | d1 | Ccr2 | mp |
| Ccl2 | batch | fap | d1 | Ccr2 | mp |
| Ccl2 | batch | ic | d1 | Ccr2 | mp |
| Ccl2 | batch | per | d1 | Ccr2 | mp |
| Hgf | batch | fap | d1 | Met | mp |
| Il18 | batch | fap | d1 | Il18rap | mp |
| Il18 | constitutive | ic | d1 | Il18rap | mp |
| Il18 | constitutive | mp | d1 | Il18rap | mp |
| Kitl | batch | fap | d1 | Kit | mp |
| Kitl | batch | ic | d1 | Kit | mp |
| Kitl | batch | mp | d1 | Kit | mp |
| Kitl | constitutive | ec | d1 | Kit | mp |
| Kitl | constitutive | per | d1 | Kit | mp |
| Pf4 | batch | fap | d1 | Ldlr | mp |
| Pf4 | batch | ic | d1 | Ldlr | mp |
| Pf4 | batch | per | d1 | Ldlr | mp |
| Pf4 | batch | fap | d1 | Thbd | mp |
| Pf4 | batch | ic | d1 | Thbd | mp |
| Pf4 | batch | per | d1 | Thbd | mp |
| Rspo3 | batch | fap | d1 | Lrp6 | mp |
| Wnt2 | batch | fap | d1 | Sfrp1 | mp |
| Wnt2 | batch | fap | d1 | Fzd1 | mp |
| Il1b | batch | ec | d1 | Il1rap | mp |
| Il1b | batch | ic | d1 | Il1rap | mp |
| Il1b | batch | per | d1 | Il1rap | mp |
| Il1b | batch | ec | d1 | Adrb2 | mp |
| Il1b | batch | ic | d1 | Adrb2 | mp |
| Il1b | batch | per | d1 | Adrb2 | mp |
| Inhbb | batch | ec | d1 | Acvr2a | mp |
| Inhbb | constitutive | fap | d1 | Acvr2a | mp |
| Inhbb | constitutive | per | d1 | Acvr2a | mp |
| Pgf | batch | ec | d1 | Flt1 | mp |
| Pgf | constitutive | per | d1 | Flt1 | mp |
| Pgf | batch | ec | d1 | Nrp1 | mp |
| Pgf | constitutive | per | d1 | Nrp1 | mp |
| Areg | batch | ic | d1 | Egfr | mp |
| Areg | batch | mp | d1 | Egfr | mp |
| Bmp4 | batch | mp | d1 | Bmpr1a | mp |
| Bmp4 | constitutive | ec | d1 | Bmpr1a | mp |
| Bmp4 | constitutive | per | d1 | Bmpr1a | mp |
| Clcf1 | batch | mp | d1 | Crlf1 | mp |
| Clcf1 | batch | mp | d1 | Cntfr | mp |
| Clcf1 | batch | mp | d1 | Esr1 | mp |
| Clu | batch | ic | d1 | Vldlr | mp |
| Clu | batch | mp | d1 | Vldlr | mp |
| Ctsg | batch | mp | d1 | Met | mp |
| Gas6 | batch | mp | d1 | Axl | mp |
| Gdnf | batch | mp | d1 | Gfra1 | mp |
| Gdnf | batch | per | d1 | Gfra1 | mp |
| Gdnf | batch | mp | d1 | Gfra2 | mp |
| Gdnf | batch | per | d1 | Gfra2 | mp |
| Gdnf | batch | mp | d1 | Ret | mp |
| Gdnf | batch | per | d1 | Ret | mp |
| Gmfb | batch | mp | d1 | Egfr | mp |
| Hbegf | batch | mp | d1 | Cd44 | mp |
| Hbegf | constitutive | ec | d1 | Cd44 | mp |
| Hbegf | constitutive | per | d1 | Cd44 | mp |
| Hbegf | batch | mp | d1 | Egfr | mp |
| Hbegf | constitutive | ec | d1 | Egfr | mp |
| Hbegf | constitutive | per | d1 | Egfr | mp |
| Jag2 | batch | mp | d1 | Notch2 | mp |
| Jag2 | constitutive | ec | d1 | Notch2 | mp |
| Ngf | batch | mp | d1 | Ror1 | mp |
| Ngf | constitutive | per | d1 | Ror1 | mp |
| Ngf | batch | mp | d1 | Ngfr | mp |
| Ngf | constitutive | per | d1 | Ngfr | mp |
| Pdap1 | batch | mp | d1 | Pdgfa | mp |
| Vegfa | batch | ic | d1 | Flt1 | mp |
| Vegfa | batch | mp | d1 | Flt1 | mp |
| Vegfa | batch | ic | d1 | Kdr | mp |
| Vegfa | batch | mp | d1 | Kdr | mp |
| Vegfa | batch | ic | d1 | Nrp1 | mp |
| Vegfa | batch | mp | d1 | Nrp1 | mp |
| Vegfc | batch | mp | d1 | Kdr | mp |
| Vegfc | batch | mp | d1 | Nrp1 | mp |
| Il1a | batch | ic | d1 | Il1rap | mp |
| Jag1 | batch | ic | d1 | Notch2 | mp |
| Jag1 | constitutive | per | d1 | Notch2 | mp |
| Osm | batch | ic | d1 | Osmr | mp |
| Osm | batch | per | d1 | Osmr | mp |
| Tnf | batch | ic | d1 | Ptprz1 | mp |
| Tnf | batch | per | d1 | Ptprz1 | mp |
| Angpt4 | batch | per | d1 | Tek | mp |
| Btc | batch | per | d1 | Egfr | mp |
| Il11 | batch | per | d1 | Il11ra1 | mp |
| Mmp9 | batch | per | d1 | Flt1 | mp |
| Angpt1 | constitutive | fap | d1 | Tek | mp |
| Fgf7 | constitutive | fap | d1 | Nrp1 | mp |
| Tgfb2 | constitutive | ec | d1 | Tgfbr3 | mp |
| Tgfb2 | constitutive | fap | d1 | Tgfbr3 | mp |
| Tgfb2 | constitutive | per | d1 | Tgfbr3 | mp |
| Bmp5 | constitutive | ec | d1 | Bmpr1a | mp |
| Bmp5 | constitutive | fap | d1 | Bmpr1a | mp |
| Bmp5 | constitutive | per | d1 | Bmpr1a | mp |
| Cmtm8 | constitutive | ec | d1 | Egfr | mp |
| Edn1 | constitutive | ec | d1 | Ednrb | mp |
| Dhh | constitutive | ec | d1 | Ptch1 | mp |
| Dhh | constitutive | mp | d1 | Ptch1 | mp |
| Dhh | constitutive | per | d1 | Ptch1 | mp |
| Angpt2 | constitutive | ec | d1 | Tek | mp |
| Angpt2 | constitutive | ic | d1 | Tek | mp |
| Angpt2 | constitutive | per | d1 | Tek | mp |
| Ptn | constitutive | per | d1 | Ptprz1 | mp |
| Ntf3 | constitutive | per | d1 | Ntrk2 | mp |
| Ntf3 | constitutive | per | d1 | Ngfr | mp |
| Apln | batch | ec | d2 | Aplnr | mp |
| Apln | batch | fap | d2 | Aplnr | mp |
| Bmp2 | batch | ec | d2 | Acvr2a | mp |
| Bmp2 | batch | fap | d2 | Acvr2a | mp |
| Bmp2 | batch | mp | d2 | Acvr2a | mp |
| Bmp2 | constitutive | per | d2 | Acvr2a | mp |
| Bmp2 | batch | ec | d2 | Bmpr1a | mp |
| Bmp2 | batch | fap | d2 | Bmpr1a | mp |
| Bmp2 | batch | mp | d2 | Bmpr1a | mp |
| Bmp2 | constitutive | per | d2 | Bmpr1a | mp |
| Bmp2 | batch | ec | d2 | Eng | mp |
| Bmp2 | batch | fap | d2 | Eng | mp |
| Bmp2 | batch | mp | d2 | Eng | mp |
| Bmp2 | constitutive | per | d2 | Eng | mp |
| Bmp6 | batch | fap | d2 | Acvr2a | mp |
| Bmp6 | constitutive | per | d2 | Acvr2a | mp |
| Bmp6 | batch | fap | d2 | Bmpr1a | mp |
| Bmp6 | constitutive | per | d2 | Bmpr1a | mp |
| Ccl2 | batch | ec | d2 | Ccr1 | mp |
| Ccl2 | batch | fap | d2 | Ccr1 | mp |
| Ccl2 | batch | ic | d2 | Ccr1 | mp |
| Ccl2 | batch | mp | d2 | Ccr1 | mp |
| Ccl2 | batch | per | d2 | Ccr1 | mp |
| Ccl2 | batch | ec | d2 | Ccr2 | mp |
| Ccl2 | batch | fap | d2 | Ccr2 | mp |
| Ccl2 | batch | ic | d2 | Ccr2 | mp |
| Ccl2 | batch | mp | d2 | Ccr2 | mp |
| Ccl2 | batch | per | d2 | Ccr2 | mp |
| Ccl2 | batch | ec | d2 | Ccr5 | mp |
| Ccl2 | batch | fap | d2 | Ccr5 | mp |
| Ccl2 | batch | ic | d2 | Ccr5 | mp |
| Ccl2 | batch | mp | d2 | Ccr5 | mp |
| Ccl2 | batch | per | d2 | Ccr5 | mp |
| Ccl3 | batch | ec | d2 | Ccr1 | mp |
| Ccl3 | batch | fap | d2 | Ccr1 | mp |
| Ccl3 | batch | ic | d2 | Ccr1 | mp |
| Ccl3 | batch | mp | d2 | Ccr1 | mp |
| Ccl3 | batch | per | d2 | Ccr1 | mp |
| Ccl3 | batch | ec | d2 | Ccr5 | mp |
| Ccl3 | batch | fap | d2 | Ccr5 | mp |
| Ccl3 | batch | ic | d2 | Ccr5 | mp |
| Ccl3 | batch | mp | d2 | Ccr5 | mp |
| Ccl3 | batch | per | d2 | Ccr5 | mp |
| Ccl4 | batch | fap | d2 | Ccr1 | mp |
| Ccl4 | batch | ic | d2 | Ccr1 | mp |
| Ccl4 | batch | mp | d2 | Ccr1 | mp |
| Ccl4 | batch | per | d2 | Ccr1 | mp |
| Ccl4 | batch | fap | d2 | Ccr5 | mp |
| Ccl4 | batch | ic | d2 | Ccr5 | mp |
| Ccl4 | batch | mp | d2 | Ccr5 | mp |
| Ccl4 | batch | per | d2 | Ccr5 | mp |
| Ccl5 | batch | ec | d2 | Ccr1 | mp |
| Ccl5 | batch | fap | d2 | Ccr1 | mp |
| Ccl5 | batch | ec | d2 | Ccr5 | mp |
| Ccl5 | batch | fap | d2 | Ccr5 | mp |
| Ccl5 | batch | ec | d2 | Sdc4 | mp |
| Ccl5 | batch | fap | d2 | Sdc4 | mp |
| Ccl6 | batch | fap | d2 | Ccr1 | mp |
| Ccl6 | batch | ic | d2 | Ccr1 | mp |
| Ccl6 | batch | mp | d2 | Ccr1 | mp |
| Ccl6 | batch | per | d2 | Ccr1 | mp |
| Ccl7 | batch | ec | d2 | Ccr1 | mp |
| Ccl7 | batch | fap | d2 | Ccr1 | mp |
| Ccl7 | batch | ic | d2 | Ccr1 | mp |
| Ccl7 | batch | mp | d2 | Ccr1 | mp |
| Ccl7 | batch | per | d2 | Ccr1 | mp |
| Ccl7 | batch | ec | d2 | Ccr5 | mp |
| Ccl7 | batch | fap | d2 | Ccr5 | mp |
| Ccl7 | batch | ic | d2 | Ccr5 | mp |
| Ccl7 | batch | mp | d2 | Ccr5 | mp |
| Ccl7 | batch | per | d2 | Ccr5 | mp |
| Csf1 | batch | fap | d2 | Csf1r | mp |
| Csf1 | batch | ic | d2 | Csf1r | mp |
| Csf1 | batch | mp | d2 | Csf1r | mp |
| Csf2 | batch | fap | d2 | Csf2ra | mp |
| Csf2 | batch | fap | d2 | Csf2rb | mp |
| Csf2 | batch | fap | d2 | Csf3r | mp |
| Hgf | batch | fap | d2 | Met | mp |
| Hgf | batch | ic | d2 | Met | mp |
| Il18 | batch | fap | d2 | Il18rap | mp |
| Il18 | constitutive | ic | d2 | Il18rap | mp |
| Il18 | constitutive | mp | d2 | Il18rap | mp |
| Il1b | batch | ec | d2 | Il1r2 | mp |
| Il1b | batch | fap | d2 | Il1r2 | mp |
| Il1b | batch | ic | d2 | Il1r2 | mp |
| Il1b | batch | mp | d2 | Il1r2 | mp |
| Il1b | batch | per | d2 | Il1r2 | mp |
| Il1b | batch | ec | d2 | Il1rap | mp |
| Il1b | batch | fap | d2 | Il1rap | mp |
| Il1b | batch | ic | d2 | Il1rap | mp |
| Il1b | batch | mp | d2 | Il1rap | mp |
| Il1b | batch | per | d2 | Il1rap | mp |
| Il1b | batch | ec | d2 | Adrb2 | mp |
| Il1b | batch | fap | d2 | Adrb2 | mp |
| Il1b | batch | ic | d2 | Adrb2 | mp |
| Il1b | batch | mp | d2 | Adrb2 | mp |
| Il1b | batch | per | d2 | Adrb2 | mp |
| Kitl | batch | fap | d2 | Kit | mp |
| Kitl | batch | mp | d2 | Kit | mp |
| Kitl | constitutive | ec | d2 | Kit | mp |
| Kitl | constitutive | per | d2 | Kit | mp |
| Mif | batch | ec | d2 | Cd74 | mp |
| Mif | batch | fap | d2 | Cd74 | mp |
| Mif | batch | mp | d2 | Cd74 | mp |
| Mmp9 | batch | fap | d2 | Flt1 | mp |
| Mmp9 | batch | mp | d2 | Flt1 | mp |
| Mmp9 | batch | per | d2 | Flt1 | mp |
| Pf4 | batch | ec | d2 | Ldlr | mp |
| Pf4 | batch | fap | d2 | Ldlr | mp |
| Pf4 | batch | ic | d2 | Ldlr | mp |
| Pf4 | batch | mp | d2 | Ldlr | mp |
| Pf4 | batch | per | d2 | Ldlr | mp |
| Pf4 | batch | ec | d2 | Thbd | mp |
| Pf4 | batch | fap | d2 | Thbd | mp |
| Pf4 | batch | ic | d2 | Thbd | mp |
| Pf4 | batch | mp | d2 | Thbd | mp |
| Pf4 | batch | per | d2 | Thbd | mp |
| Rspo3 | batch | fap | d2 | Lrp6 | mp |
| Spp1 | batch | ec | d2 | Itgb5 | mp |
| Spp1 | batch | fap | d2 | Itgb5 | mp |
| Spp1 | batch | ic | d2 | Itgb5 | mp |
| Spp1 | batch | mp | d2 | Itgb5 | mp |
| Spp1 | batch | per | d2 | Itgb5 | mp |
| Tslp | batch | fap | d2 | Crlf2 | mp |
| Tslp | constitutive | ec | d2 | Crlf2 | mp |
| Wnt2 | batch | fap | d2 | Sfrp1 | mp |
| Wnt2 | batch | fap | d2 | Fzd1 | mp |
| Igf1 | batch | ec | d2 | Igfbp2 | mp |
| Igf1 | batch | ic | d2 | Igfbp2 | mp |
| Igf1 | batch | mp | d2 | Igfbp2 | mp |
| Igf1 | batch | ec | d2 | Igf1r | mp |
| Igf1 | batch | ic | d2 | Igf1r | mp |
| Igf1 | batch | mp | d2 | Igf1r | mp |
| Il15 | batch | ec | d2 | Il17ra | mp |
| Il15 | batch | ic | d2 | Il17ra | mp |
| Il15 | batch | ec | d2 | Il2rg | mp |
| Il15 | batch | ic | d2 | Il2rg | mp |
| Inhbb | batch | ec | d2 | Acvr2a | mp |
| Inhbb | constitutive | fap | d2 | Acvr2a | mp |
| Inhbb | constitutive | per | d2 | Acvr2a | mp |
| Lgals3 | batch | ec | d2 | Lgals3bp | mp |
| Lgals3 | batch | ic | d2 | Lgals3bp | mp |
| Lgals3 | batch | mp | d2 | Lgals3bp | mp |
| Lgals3 | batch | per | d2 | Lgals3bp | mp |
| Mmp12 | batch | ec | d2 | Plaur | mp |
| Mmp12 | batch | ic | d2 | Plaur | mp |
| Mmp12 | batch | mp | d2 | Plaur | mp |
| Mmp12 | batch | per | d2 | Plaur | mp |
| Mmp13 | batch | ec | d2 | F2r | mp |
| Mmp13 | batch | ic | d2 | F2r | mp |
| Mmp13 | batch | per | d2 | F2r | mp |
| Osm | batch | ec | d2 | Osmr | mp |
| Osm | batch | ic | d2 | Osmr | mp |
| Osm | batch | mp | d2 | Osmr | mp |
| Osm | batch | per | d2 | Osmr | mp |
| Pgf | batch | ec | d2 | Flt1 | mp |
| Pgf | constitutive | per | d2 | Flt1 | mp |
| Pgf | batch | ec | d2 | Nrp1 | mp |
| Pgf | constitutive | per | d2 | Nrp1 | mp |
| Pgf | batch | ec | d2 | Nrp2 | mp |
| Pgf | constitutive | per | d2 | Nrp2 | mp |
| Areg | batch | ic | d2 | Egfr | mp |
| Areg | batch | mp | d2 | Egfr | mp |
| Bmp4 | batch | mp | d2 | Bmpr1a | mp |
| Bmp4 | constitutive | ec | d2 | Bmpr1a | mp |
| Bmp4 | constitutive | per | d2 | Bmpr1a | mp |
| C4b | batch | mp | d2 | C3ar1 | mp |
| C4b | constitutive | per | d2 | C3ar1 | mp |
| Ccl11 | batch | mp | d2 | Ccr5 | mp |
| Ccl11 | batch | per | d2 | Ccr5 | mp |
| Ccl8 | batch | mp | d2 | Ccr1 | mp |
| Ccl8 | total | fap | d2 | Ccr1 | mp |
| Ccl8 | total | ic | d2 | Ccr1 | mp |
| Ccl8 | batch | mp | d2 | Ccr2 | mp |
| Ccl8 | total | fap | d2 | Ccr2 | mp |
| Ccl8 | total | ic | d2 | Ccr2 | mp |
| Ccl8 | batch | mp | d2 | Ccr5 | mp |
| Ccl8 | total | fap | d2 | Ccr5 | mp |
| Ccl8 | total | ic | d2 | Ccr5 | mp |
| Clcf1 | batch | mp | d2 | Crlf1 | mp |
| Clcf1 | batch | mp | d2 | Cntfr | mp |
| Clcf1 | batch | mp | d2 | Esr1 | mp |
| Clu | batch | mp | d2 | Vldlr | mp |
| Clu | total | per | d2 | Vldlr | mp |
| Ctsg | batch | mp | d2 | Met | mp |
| Cx3cl1 | batch | mp | d2 | Cx3cr1 | mp |
| Esm1 | batch | mp | d2 | Itgal | mp |
| Gas6 | batch | mp | d2 | Axl | mp |
| Gdnf | batch | mp | d2 | Gfra1 | mp |
| Gdnf | batch | per | d2 | Gfra1 | mp |
| Gdnf | batch | mp | d2 | Gfra2 | mp |
| Gdnf | batch | per | d2 | Gfra2 | mp |
| Gdnf | batch | mp | d2 | Ret | mp |
| Gdnf | batch | per | d2 | Ret | mp |
| Gmfb | batch | mp | d2 | Egfr | mp |
| Hbegf | batch | mp | d2 | Cd44 | mp |
| Hbegf | constitutive | ec | d2 | Cd44 | mp |
| Hbegf | constitutive | per | d2 | Cd44 | mp |
| Hbegf | batch | mp | d2 | Egfr | mp |
| Hbegf | constitutive | ec | d2 | Egfr | mp |
| Hbegf | constitutive | per | d2 | Egfr | mp |
| Il10 | batch | ic | d2 | Il10ra | mp |
| Il10 | batch | mp | d2 | Il10ra | mp |
| Il10 | batch | ic | d2 | Il10rb | mp |
| Il10 | batch | mp | d2 | Il10rb | mp |
| Jag2 | batch | mp | d2 | Notch2 | mp |
| Jag2 | constitutive | ec | d2 | Notch2 | mp |
| Ngf | batch | mp | d2 | Ror1 | mp |
| Ngf | constitutive | per | d2 | Ror1 | mp |
| Ngf | batch | mp | d2 | Ngfr | mp |
| Ngf | constitutive | per | d2 | Ngfr | mp |
| Pdap1 | batch | mp | d2 | Pdgfa | mp |
| Tgfb1 | batch | mp | d2 | Eng | mp |
| Tgfb1 | total | ec | d2 | Eng | mp |
| Tgfb1 | total | ic | d2 | Eng | mp |
| Tgfb1 | batch | mp | d2 | Tgfbr3 | mp |
| Tgfb1 | total | ec | d2 | Tgfbr3 | mp |
| Tgfb1 | total | ic | d2 | Tgfbr3 | mp |
| Tnf | batch | ic | d2 | Ptprz1 | mp |
| Tnf | batch | mp | d2 | Ptprz1 | mp |
| Tnf | batch | per | d2 | Ptprz1 | mp |
| Tnf | batch | ic | d2 | Tnfrsf1b | mp |
| Tnf | batch | mp | d2 | Tnfrsf1b | mp |
| Tnf | batch | per | d2 | Tnfrsf1b | mp |
| Vegfa | batch | ic | d2 | Flt1 | mp |
| Vegfa | batch | mp | d2 | Flt1 | mp |
| Vegfa | batch | ic | d2 | Kdr | mp |
| Vegfa | batch | mp | d2 | Kdr | mp |
| Vegfa | batch | ic | d2 | Nrp1 | mp |
| Vegfa | batch | mp | d2 | Nrp1 | mp |
| Vegfa | batch | ic | d2 | Nrp2 | mp |
| Vegfa | batch | mp | d2 | Nrp2 | mp |
| Vegfc | batch | mp | d2 | Kdr | mp |
| Vegfc | batch | mp | d2 | Nrp1 | mp |
| Vegfc | batch | mp | d2 | Nrp2 | mp |
| Il1a | batch | ic | d2 | Il1r2 | mp |
| Il1a | batch | ic | d2 | Il1rap | mp |
| Jag1 | batch | ic | d2 | Notch2 | mp |
| Jag1 | constitutive | per | d2 | Notch2 | mp |
| Ppbp | batch | ic | d2 | Itgb5 | mp |
| Tnfsf13 | batch | ic | d2 | Tnfrsf11b | mp |
| Tnfsf13 | batch | ic | d2 | Tnfrsf12a | mp |
| Angpt4 | batch | per | d2 | Tek | mp |
| Btc | batch | per | d2 | Egfr | mp |
| Il11 | batch | per | d2 | Il11ra1 | mp |
| Ucn2 | batch | per | d2 | Il10rb | mp |
| Tnfsf12 | total | ec | d2 | Tnfrsf11b | mp |
| Tnfsf12 | total | ic | d2 | Tnfrsf11b | mp |
| Tnfsf12 | total | per | d2 | Tnfrsf11b | mp |
| Tnfsf12 | total | ec | d2 | Tnfrsf12a | mp |
| Tnfsf12 | total | ic | d2 | Tnfrsf12a | mp |
| Tnfsf12 | total | per | d2 | Tnfrsf12a | mp |
| Fstl1 | constitutive | fap | d2 | Cd14 | mp |
| Angpt1 | constitutive | fap | d2 | Tek | mp |
| Fgf7 | constitutive | fap | d2 | Nrp1 | mp |
| Tgfb2 | constitutive | ec | d2 | Tgfbr3 | mp |
| Tgfb2 | constitutive | fap | d2 | Tgfbr3 | mp |
| Tgfb2 | constitutive | per | d2 | Tgfbr3 | mp |
| Bmp5 | constitutive | ec | d2 | Bmpr1a | mp |
| Bmp5 | constitutive | fap | d2 | Bmpr1a | mp |
| Bmp5 | constitutive | per | d2 | Bmpr1a | mp |
| Cmtm8 | constitutive | ec | d2 | Egfr | mp |
| Edn1 | constitutive | ec | d2 | Ednrb | mp |
| Dhh | constitutive | ec | d2 | Ptch1 | mp |
| Dhh | constitutive | mp | d2 | Ptch1 | mp |
| Dhh | constitutive | per | d2 | Ptch1 | mp |
| Angpt2 | constitutive | ec | d2 | Tek | mp |
| Angpt2 | constitutive | ic | d2 | Tek | mp |
| Angpt2 | constitutive | per | d2 | Tek | mp |
| Ptn | constitutive | per | d2 | Ptprz1 | mp |
| Ntf3 | constitutive | per | d2 | Ntrk2 | mp |
| Ntf3 | constitutive | per | d2 | Ngfr | mp |
| Apln | batch | ec | d3 | Aplnr | mp |
| Apln | batch | fap | d3 | Aplnr | mp |
| C4b | batch | fap | d3 | C3ar1 | mp |
| C4b | constitutive | per | d3 | C3ar1 | mp |
| Ccl3 | batch | ec | d3 | Ccr1 | mp |
| Ccl3 | batch | fap | d3 | Ccr1 | mp |
| Ccl3 | batch | mp | d3 | Ccr1 | mp |
| Ccl3 | batch | per | d3 | Ccr1 | mp |
| Ccl3 | batch | ec | d3 | Ccr5 | mp |
| Ccl3 | batch | fap | d3 | Ccr5 | mp |
| Ccl3 | batch | mp | d3 | Ccr5 | mp |
| Ccl3 | batch | per | d3 | Ccr5 | mp |
| Ccl4 | batch | fap | d3 | Ccr1 | mp |
| Ccl4 | batch | mp | d3 | Ccr1 | mp |
| Ccl4 | batch | per | d3 | Ccr1 | mp |
| Ccl4 | batch | fap | d3 | Ccr5 | mp |
| Ccl4 | batch | mp | d3 | Ccr5 | mp |
| Ccl4 | batch | per | d3 | Ccr5 | mp |
| Ccl5 | batch | ec | d3 | Ccr1 | mp |
| Ccl5 | batch | fap | d3 | Ccr1 | mp |
| Ccl5 | batch | ec | d3 | Ccr5 | mp |
| Ccl5 | batch | fap | d3 | Ccr5 | mp |
| Ccl6 | batch | fap | d3 | Ccr1 | mp |
| Ccl6 | batch | mp | d3 | Ccr1 | mp |
| Ccl6 | batch | per | d3 | Ccr1 | mp |
| Csf2 | batch | fap | d3 | Csf2ra | mp |
| Csf2 | batch | fap | d3 | Csf2rb | mp |
| Csf2 | batch | fap | d3 | Csf3r | mp |
| Cx3cl1 | batch | fap | d3 | Cx3cr1 | mp |
| Cx3cl1 | total | ec | d3 | Cx3cr1 | mp |
| Cx3cl1 | total | per | d3 | Cx3cr1 | mp |
| Gas6 | batch | fap | d3 | Axl | mp |
| Hbegf | batch | fap | d3 | Cd44 | mp |
| Hbegf | constitutive | ec | d3 | Cd44 | mp |
| Hbegf | constitutive | per | d3 | Cd44 | mp |
| Igf1 | batch | ec | d3 | Igfbp2 | mp |
| Igf1 | batch | fap | d3 | Igfbp2 | mp |
| Igf1 | batch | ic | d3 | Igfbp2 | mp |
| Igf1 | batch | mp | d3 | Igfbp2 | mp |
| Igf1 | batch | ec | d3 | Igf1r | mp |
| Igf1 | batch | fap | d3 | Igf1r | mp |
| Igf1 | batch | ic | d3 | Igf1r | mp |
| Igf1 | batch | mp | d3 | Igf1r | mp |
| Igf2 | batch | fap | d3 | Igf1r | mp |
| Il18 | batch | fap | d3 | Il18rap | mp |
| Il18 | constitutive | ic | d3 | Il18rap | mp |
| Il18 | constitutive | mp | d3 | Il18rap | mp |
| Il1b | batch | fap | d3 | Il1r2 | mp |
| Il1b | batch | mp | d3 | Il1r2 | mp |
| Il1b | batch | per | d3 | Il1r2 | mp |
| Il1b | batch | fap | d3 | Adrb2 | mp |
| Il1b | batch | mp | d3 | Adrb2 | mp |
| Il1b | batch | per | d3 | Adrb2 | mp |
| Kitl | batch | fap | d3 | Kit | mp |
| Kitl | constitutive | ec | d3 | Kit | mp |
| Kitl | constitutive | per | d3 | Kit | mp |
| Mif | batch | fap | d3 | Cd74 | mp |
| Mif | batch | mp | d3 | Cd74 | mp |
| Ntf3 | batch | ec | d3 | Ngfr | mp |
| Ntf3 | batch | fap | d3 | Ngfr | mp |
| Ntf3 | constitutive | per | d3 | Ngfr | mp |
| Spp1 | batch | ec | d3 | Itgb5 | mp |
| Spp1 | batch | fap | d3 | Itgb5 | mp |
| Spp1 | batch | ic | d3 | Itgb5 | mp |
| Spp1 | batch | mp | d3 | Itgb5 | mp |
| Spp1 | batch | per | d3 | Itgb5 | mp |
| Tslp | batch | fap | d3 | Crlf2 | mp |
| Tslp | constitutive | ec | d3 | Crlf2 | mp |
| Wnt2 | batch | fap | d3 | Sfrp1 | mp |
| Il15 | batch | ec | d3 | Il2rg | mp |
| Il15 | batch | ic | d3 | Il2rg | mp |
| Lgals3 | batch | ec | d3 | Lgals3bp | mp |
| Lgals3 | batch | ic | d3 | Lgals3bp | mp |
| Lgals3 | batch | mp | d3 | Lgals3bp | mp |
| Lgals3 | batch | per | d3 | Lgals3bp | mp |
| Mmp12 | batch | ec | d3 | Plaur | mp |
| Mmp12 | batch | ic | d3 | Plaur | mp |
| Mmp12 | batch | mp | d3 | Plaur | mp |
| Mmp12 | batch | per | d3 | Plaur | mp |
| Vegfc | batch | ec | d3 | Nrp1 | mp |
| Vegfc | batch | ec | d3 | Nrp2 | mp |
| Ccl2 | batch | mp | d3 | Ccr1 | mp |
| Ccl2 | batch | per | d3 | Ccr1 | mp |
| Ccl2 | batch | mp | d3 | Ccr2 | mp |
| Ccl2 | batch | per | d3 | Ccr2 | mp |
| Ccl2 | batch | mp | d3 | Ccr5 | mp |
| Ccl2 | batch | per | d3 | Ccr5 | mp |
| Ccl7 | batch | mp | d3 | Ccr1 | mp |
| Ccl7 | batch | per | d3 | Ccr1 | mp |
| Ccl7 | batch | mp | d3 | Ccr5 | mp |
| Ccl7 | batch | per | d3 | Ccr5 | mp |
| Ccl8 | batch | mp | d3 | Ccr1 | mp |
| Ccl8 | total | fap | d3 | Ccr1 | mp |
| Ccl8 | total | ic | d3 | Ccr1 | mp |
| Ccl8 | batch | mp | d3 | Ccr2 | mp |
| Ccl8 | total | fap | d3 | Ccr2 | mp |
| Ccl8 | total | ic | d3 | Ccr2 | mp |
| Ccl8 | batch | mp | d3 | Ccr5 | mp |
| Ccl8 | total | fap | d3 | Ccr5 | mp |
| Ccl8 | total | ic | d3 | Ccr5 | mp |
| Csf1 | batch | ic | d3 | Csf1r | mp |
| Csf1 | batch | mp | d3 | Csf1r | mp |
| Il10 | batch | ic | d3 | Il10ra | mp |
| Il10 | batch | mp | d3 | Il10ra | mp |
| Il10 | batch | ic | d3 | Il10rb | mp |
| Il10 | batch | mp | d3 | Il10rb | mp |
| Pdap1 | batch | mp | d3 | Pdgfa | mp |
| Tnf | batch | mp | d3 | Tnfrsf1b | mp |
| Tnf | batch | per | d3 | Tnfrsf1b | mp |
| Tnfsf13 | batch | ic | d3 | Tnfrsf11b | mp |
| Btc | batch | per | d3 | Erbb2 | mp |
| Ucn2 | batch | per | d3 | Il10rb | mp |
| Dll1 | total | mp | d3 | Notch2 | mp |
| Fstl1 | constitutive | fap | d3 | Cd14 | mp |
| Fgf7 | constitutive | fap | d3 | Nrp1 | mp |
| Jag2 | constitutive | ec | d3 | Notch2 | mp |
| Ngf | constitutive | per | d3 | Ngfr | mp |
| Pgf | constitutive | per | d3 | Nrp1 | mp |
| Pgf | constitutive | per | d3 | Nrp2 | mp |
| Jag1 | constitutive | per | d3 | Notch2 | mp |
| C4b | batch | fap | d4 | C3ar1 | mp |
| C4b | batch | ic | d4 | C3ar1 | mp |
| C4b | constitutive | per | d4 | C3ar1 | mp |
| Ccl3 | batch | ec | d4 | Ccr5 | mp |
| Ccl3 | batch | fap | d4 | Ccr5 | mp |
| Ccl3 | batch | mp | d4 | Ccr5 | mp |
| Ccl4 | batch | fap | d4 | Ccr5 | mp |
| Ccl4 | batch | mp | d4 | Ccr5 | mp |
| Ccl5 | batch | ec | d4 | Ccr5 | mp |
| Ccl5 | batch | fap | d4 | Ccr5 | mp |
| Cx3cl1 | batch | fap | d4 | Cx3cr1 | mp |
| Cx3cl1 | batch | mp | d4 | Cx3cr1 | mp |
| Cx3cl1 | total | ec | d4 | Cx3cr1 | mp |
| Cx3cl1 | total | per | d4 | Cx3cr1 | mp |
| Gas6 | batch | fap | d4 | Axl | mp |
| Hbegf | batch | fap | d4 | Cd44 | mp |
| Hbegf | constitutive | ec | d4 | Cd44 | mp |
| Hbegf | constitutive | per | d4 | Cd44 | mp |
| Hbegf | batch | fap | d4 | Egfr | mp |
| Hbegf | constitutive | ec | d4 | Egfr | mp |
| Hbegf | constitutive | per | d4 | Egfr | mp |
| Hgf | batch | fap | d4 | Met | mp |
| Hgf | batch | ic | d4 | Met | mp |
| Igf1 | batch | ec | d4 | Igfbp2 | mp |
| Igf1 | batch | fap | d4 | Igfbp2 | mp |
| Igf1 | batch | ic | d4 | Igfbp2 | mp |
| Igf1 | batch | ec | d4 | Igfbp3 | mp |
| Igf1 | batch | fap | d4 | Igfbp3 | mp |
| Igf1 | batch | ic | d4 | Igfbp3 | mp |
| Igf1 | batch | ec | d4 | Igfbp5 | mp |
| Igf1 | batch | fap | d4 | Igfbp5 | mp |
| Igf1 | batch | ic | d4 | Igfbp5 | mp |
| Igf1 | batch | ec | d4 | Igf1r | mp |
| Igf1 | batch | fap | d4 | Igf1r | mp |
| Igf1 | batch | ic | d4 | Igf1r | mp |
| Igf2 | batch | ec | d4 | Igf2r | mp |
| Igf2 | batch | fap | d4 | Igf2r | mp |
| Igf2 | batch | mp | d4 | Igf2r | mp |
| Igf2 | batch | ec | d4 | Igf1r | mp |
| Igf2 | batch | fap | d4 | Igf1r | mp |
| Igf2 | batch | mp | d4 | Igf1r | mp |
| Il1b | batch | fap | d4 | Il1rap | mp |
| Il1b | batch | fap | d4 | Adrb2 | mp |
| Kitl | batch | fap | d4 | Kit | mp |
| Kitl | batch | mp | d4 | Kit | mp |
| Kitl | constitutive | ec | d4 | Kit | mp |
| Kitl | constitutive | per | d4 | Kit | mp |
| Mif | batch | fap | d4 | Cd74 | mp |
| Mmp13 | batch | fap | d4 | F2r | mp |
| Mmp13 | batch | ic | d4 | F2r | mp |
| Mmp13 | batch | mp | d4 | F2r | mp |
| Ntf3 | batch | ec | d4 | Ngfr | mp |
| Ntf3 | batch | fap | d4 | Ngfr | mp |
| Ntf3 | constitutive | per | d4 | Ngfr | mp |
| Pf4 | batch | ec | d4 | Thbd | mp |
| Pf4 | batch | fap | d4 | Thbd | mp |
| Rspo3 | batch | fap | d4 | Lrp6 | mp |
| Sfrp1 | batch | ec | d4 | Fzd2 | mp |
| Sfrp1 | batch | fap | d4 | Fzd2 | mp |
| Sfrp1 | constitutive | per | d4 | Fzd2 | mp |
| Sfrp2 | batch | ec | d4 | Fzd2 | mp |
| Sfrp2 | batch | fap | d4 | Fzd2 | mp |
| Sfrp4 | batch | fap | d4 | Fzd2 | mp |
| Spp1 | batch | ec | d4 | Itgb5 | mp |
| Spp1 | batch | fap | d4 | Itgb5 | mp |
| Spp1 | batch | ic | d4 | Itgb5 | mp |
| Spp1 | batch | mp | d4 | Itgb5 | mp |
| Clu | batch | ec | d4 | Vldlr | mp |
| Dll1 | batch | ec | d4 | Notch2 | mp |
| Dll1 | total | mp | d4 | Notch2 | mp |
| Il15 | batch | ec | d4 | Il2rg | mp |
| Il15 | batch | ic | d4 | Il2rg | mp |
| Lgals3 | batch | ec | d4 | Lgals3bp | mp |
| Lgals3 | batch | ic | d4 | Lgals3bp | mp |
| Lgals3 | batch | mp | d4 | Lgals3bp | mp |
| Osm | batch | ec | d4 | Osmr | mp |
| Vegfc | batch | ec | d4 | Nrp1 | mp |
| Vegfc | batch | mp | d4 | Nrp1 | mp |
| Areg | batch | mp | d4 | Egfr | mp |
| Ccl8 | batch | mp | d4 | Ccr2 | mp |
| Ccl8 | total | fap | d4 | Ccr2 | mp |
| Ccl8 | total | ic | d4 | Ccr2 | mp |
| Ccl8 | batch | mp | d4 | Ccr5 | mp |
| Ccl8 | total | fap | d4 | Ccr5 | mp |
| Ccl8 | total | ic | d4 | Ccr5 | mp |
| Gdnf | batch | mp | d4 | Gfra1 | mp |
| Gdnf | batch | mp | d4 | Ret | mp |
| Gmfb | batch | mp | d4 | Egfr | mp |
| Il10 | batch | ic | d4 | Il10ra | mp |
| Il10 | batch | mp | d4 | Il10ra | mp |
| Il10 | batch | ic | d4 | Il10rb | mp |
| Il10 | batch | mp | d4 | Il10rb | mp |
| Pdap1 | batch | mp | d4 | Pdgfa | mp |
| Csf1 | batch | ic | d4 | Csf1r | mp |
| Tnfsf13 | batch | ic | d4 | Tnfrsf11b | mp |
| Angpt1 | constitutive | fap | d4 | Tek | mp |
| Fgf7 | constitutive | fap | d4 | Nrp1 | mp |
| Bmp5 | constitutive | ec | d4 | Bmpr1a | mp |
| Bmp5 | constitutive | fap | d4 | Bmpr1a | mp |
| Bmp5 | constitutive | per | d4 | Bmpr1a | mp |
| Inhbb | constitutive | fap | d4 | Acvr2a | mp |
| Inhbb | constitutive | per | d4 | Acvr2a | mp |
| Jag2 | constitutive | ec | d4 | Notch2 | mp |
| Efnb2 | constitutive | ec | d4 | Ephb1 | mp |
| Cmtm8 | constitutive | ec | d4 | Egfr | mp |
| Angpt2 | constitutive | ec | d4 | Tek | mp |
| Angpt2 | constitutive | ic | d4 | Tek | mp |
| Angpt2 | constitutive | per | d4 | Tek | mp |
| Bmp4 | constitutive | ec | d4 | Bmpr1a | mp |
| Bmp4 | constitutive | per | d4 | Bmpr1a | mp |
| Il18 | constitutive | ic | d4 | Il18rap | mp |
| Il18 | constitutive | mp | d4 | Il18rap | mp |
| Serpini1 | constitutive | per | d4 | Plat | mp |
| Ngf | constitutive | per | d4 | Ror1 | mp |
| Ngf | constitutive | per | d4 | Ngfr | mp |
| Bmp6 | constitutive | per | d4 | Acvr2a | mp |
| Bmp6 | constitutive | per | d4 | Bmpr1a | mp |
| Pgf | constitutive | per | d4 | Nrp1 | mp |
| Jag1 | constitutive | per | d4 | Notch2 | mp |
| Bmp2 | constitutive | per | d4 | Acvr2a | mp |
| Bmp2 | constitutive | per | d4 | Bmpr1a | mp |
| Gas6 | batch | fap | d5 | Axl | mp |
| Hbegf | batch | fap | d5 | Cd44 | mp |
| Hbegf | constitutive | ec | d5 | Cd44 | mp |
| Hbegf | constitutive | per | d5 | Cd44 | mp |
| Hbegf | batch | fap | d5 | Egfr | mp |
| Hbegf | constitutive | ec | d5 | Egfr | mp |
| Hbegf | constitutive | per | d5 | Egfr | mp |
| Hgf | batch | fap | d5 | Met | mp |
| Hgf | batch | ic | d5 | Met | mp |
| Igf1 | batch | ec | d5 | Igfbp2 | mp |
| Igf1 | batch | fap | d5 | Igfbp2 | mp |
| Igf1 | batch | ic | d5 | Igfbp2 | mp |
| Igf1 | batch | per | d5 | Igfbp2 | mp |
| Igf1 | batch | ec | d5 | Igfbp3 | mp |
| Igf1 | batch | fap | d5 | Igfbp3 | mp |
| Igf1 | batch | ic | d5 | Igfbp3 | mp |
| Igf1 | batch | per | d5 | Igfbp3 | mp |
| Igf1 | batch | ec | d5 | Igfbp5 | mp |
| Igf1 | batch | fap | d5 | Igfbp5 | mp |
| Igf1 | batch | ic | d5 | Igfbp5 | mp |
| Igf1 | batch | per | d5 | Igfbp5 | mp |
| Igf1 | batch | ec | d5 | Igf1r | mp |
| Igf1 | batch | fap | d5 | Igf1r | mp |
| Igf1 | batch | ic | d5 | Igf1r | mp |
| Igf1 | batch | per | d5 | Igf1r | mp |
| Igf2 | batch | ec | d5 | Igf2r | mp |
| Igf2 | batch | fap | d5 | Igf2r | mp |
| Igf2 | batch | ic | d5 | Igf2r | mp |
| Igf2 | batch | mp | d5 | Igf2r | mp |
| Igf2 | batch | ec | d5 | Igf1r | mp |
| Igf2 | batch | fap | d5 | Igf1r | mp |
| Igf2 | batch | ic | d5 | Igf1r | mp |
| Igf2 | batch | mp | d5 | Igf1r | mp |
| Il1b | batch | fap | d5 | Il1rap | mp |
| Il1b | batch | fap | d5 | Adrb2 | mp |
| Kitl | batch | fap | d5 | Kit | mp |
| Kitl | batch | ic | d5 | Kit | mp |
| Kitl | batch | mp | d5 | Kit | mp |
| Kitl | constitutive | ec | d5 | Kit | mp |
| Kitl | constitutive | per | d5 | Kit | mp |
| Mmp13 | batch | ec | d5 | F2r | mp |
| Mmp13 | batch | fap | d5 | F2r | mp |
| Mmp13 | batch | ic | d5 | F2r | mp |
| Mmp13 | batch | mp | d5 | F2r | mp |
| Mmp13 | batch | per | d5 | F2r | mp |
| Ntf3 | batch | ec | d5 | Ngfr | mp |
| Ntf3 | batch | fap | d5 | Ngfr | mp |
| Ntf3 | constitutive | per | d5 | Ngfr | mp |
| Pf4 | batch | fap | d5 | Thbd | mp |
| Rspo3 | batch | fap | d5 | Lrp6 | mp |
| Sfrp1 | batch | fap | d5 | Fzd2 | mp |
| Sfrp1 | constitutive | per | d5 | Fzd2 | mp |
| Sfrp2 | batch | fap | d5 | Fzd2 | mp |
| Sfrp2 | batch | per | d5 | Fzd2 | mp |
| Sfrp4 | batch | fap | d5 | Fzd2 | mp |
| Sfrp4 | batch | per | d5 | Fzd2 | mp |
| Clu | batch | ec | d5 | Vldlr | mp |
| Clu | batch | ic | d5 | Vldlr | mp |
| Dll1 | batch | ec | d5 | Notch2 | mp |
| Dll1 | total | mp | d5 | Notch2 | mp |
| Inhbb | batch | ec | d5 | Acvr2a | mp |
| Inhbb | constitutive | fap | d5 | Acvr2a | mp |
| Inhbb | constitutive | per | d5 | Acvr2a | mp |
| Vegfc | batch | ec | d5 | Nrp1 | mp |
| Vegfc | batch | mp | d5 | Nrp1 | mp |
| Areg | batch | mp | d5 | Egfr | mp |
| Gdnf | batch | mp | d5 | Gfra1 | mp |
| Gdnf | batch | per | d5 | Gfra1 | mp |
| Gdnf | batch | mp | d5 | Ret | mp |
| Gdnf | batch | per | d5 | Ret | mp |
| Gmfb | batch | mp | d5 | Egfr | mp |
| Pdap1 | batch | mp | d5 | Pdgfa | mp |
| Tnfsf13 | batch | ic | d5 | Tnfrsf11b | mp |
| Angpt4 | batch | per | d5 | Tek | mp |
| Btc | batch | per | d5 | Egfr | mp |
| Btc | batch | per | d5 | Erbb2 | mp |
| Ccl2 | batch | per | d5 | Ccr2 | mp |
| Sfrp5 | batch | per | d5 | Fzd2 | mp |
| Ccl8 | total | fap | d5 | Ccr2 | mp |
| Ccl8 | total | ic | d5 | Ccr2 | mp |
| Angpt1 | constitutive | fap | d5 | Tek | mp |
| Fgf7 | constitutive | fap | d5 | Nrp1 | mp |
| Bmp5 | constitutive | ec | d5 | Bmpr1a | mp |
| Bmp5 | constitutive | fap | d5 | Bmpr1a | mp |
| Bmp5 | constitutive | per | d5 | Bmpr1a | mp |
| Jag2 | constitutive | ec | d5 | Notch2 | mp |
| Efnb2 | constitutive | ec | d5 | Ephb1 | mp |
| Cmtm8 | constitutive | ec | d5 | Egfr | mp |
| Angpt2 | constitutive | ec | d5 | Tek | mp |
| Angpt2 | constitutive | ic | d5 | Tek | mp |
| Angpt2 | constitutive | per | d5 | Tek | mp |
| Bmp4 | constitutive | ec | d5 | Bmpr1a | mp |
| Bmp4 | constitutive | per | d5 | Bmpr1a | mp |
| Il18 | constitutive | ic | d5 | Il18rap | mp |
| Il18 | constitutive | mp | d5 | Il18rap | mp |
| Serpini1 | constitutive | per | d5 | Plat | mp |
| Ngf | constitutive | per | d5 | Ror1 | mp |
| Ngf | constitutive | per | d5 | Ngfr | mp |
| Bmp6 | constitutive | per | d5 | Acvr2a | mp |
| Bmp6 | constitutive | per | d5 | Bmpr1a | mp |
| Pgf | constitutive | per | d5 | Nrp1 | mp |
| Jag1 | constitutive | per | d5 | Notch2 | mp |
| Bmp2 | constitutive | per | d5 | Acvr2a | mp |
| Bmp2 | constitutive | per | d5 | Bmpr1a | mp |
| C4b | batch | fap | d6 | C3ar1 | mp |
| C4b | batch | ic | d6 | C3ar1 | mp |
| C4b | batch | mp | d6 | C3ar1 | mp |
| C4b | constitutive | per | d6 | C3ar1 | mp |
| Ccl3 | batch | fap | d6 | Ccr5 | mp |
| Ccl3 | batch | mp | d6 | Ccr5 | mp |
| Ccl4 | batch | fap | d6 | Ccr5 | mp |
| Ccl4 | batch | mp | d6 | Ccr5 | mp |
| Ccl5 | batch | fap | d6 | Ccr5 | mp |
| Ccl5 | batch | fap | d6 | Sdc4 | mp |
| Crlf1 | batch | fap | d6 | Ctf1 | mp |
| Crlf1 | batch | mp | d6 | Ctf1 | mp |
| Cx3cl1 | batch | fap | d6 | Cx3cr1 | mp |
| Cx3cl1 | batch | mp | d6 | Cx3cr1 | mp |
| Cx3cl1 | total | ec | d6 | Cx3cr1 | mp |
| Cx3cl1 | total | per | d6 | Cx3cr1 | mp |
| Fgf1 | batch | fap | d6 | Fgfr1 | mp |
| Fgf1 | constitutive | per | d6 | Fgfr1 | mp |
| Fgf1 | batch | fap | d6 | Fgfr4 | mp |
| Fgf1 | constitutive | per | d6 | Fgfr4 | mp |
| Gas6 | batch | fap | d6 | Axl | mp |
| Gas6 | batch | mp | d6 | Axl | mp |
| Hbegf | batch | fap | d6 | Cd44 | mp |
| Hbegf | batch | mp | d6 | Cd44 | mp |
| Hbegf | constitutive | ec | d6 | Cd44 | mp |
| Hbegf | constitutive | per | d6 | Cd44 | mp |
| Hbegf | batch | fap | d6 | Egfr | mp |
| Hbegf | batch | mp | d6 | Egfr | mp |
| Hbegf | constitutive | ec | d6 | Egfr | mp |
| Hbegf | constitutive | per | d6 | Egfr | mp |
| Igf1 | batch | ec | d6 | Igfbp2 | mp |
| Igf1 | batch | fap | d6 | Igfbp2 | mp |
| Igf1 | batch | ic | d6 | Igfbp2 | mp |
| Igf1 | batch | ec | d6 | Igfbp3 | mp |
| Igf1 | batch | fap | d6 | Igfbp3 | mp |
| Igf1 | batch | ic | d6 | Igfbp3 | mp |
| Igf1 | batch | ec | d6 | Igfbp4 | mp |
| Igf1 | batch | fap | d6 | Igfbp4 | mp |
| Igf1 | batch | ic | d6 | Igfbp4 | mp |
| Igf1 | batch | ec | d6 | Igfbp5 | mp |
| Igf1 | batch | fap | d6 | Igfbp5 | mp |
| Igf1 | batch | ic | d6 | Igfbp5 | mp |
| Igf1 | batch | ec | d6 | Igf1r | mp |
| Igf1 | batch | fap | d6 | Igf1r | mp |
| Igf1 | batch | ic | d6 | Igf1r | mp |
| Igf2 | batch | ec | d6 | Igf2r | mp |
| Igf2 | batch | fap | d6 | Igf2r | mp |
| Igf2 | batch | ic | d6 | Igf2r | mp |
| Igf2 | batch | mp | d6 | Igf2r | mp |
| Igf2 | batch | ec | d6 | Vtn | mp |
| Igf2 | batch | fap | d6 | Vtn | mp |
| Igf2 | batch | ic | d6 | Vtn | mp |
| Igf2 | batch | mp | d6 | Vtn | mp |
| Igf2 | batch | ec | d6 | Igf1r | mp |
| Igf2 | batch | fap | d6 | Igf1r | mp |
| Igf2 | batch | ic | d6 | Igf1r | mp |
| Igf2 | batch | mp | d6 | Igf1r | mp |
| Il1b | batch | fap | d6 | Adrb2 | mp |
| Kitl | batch | fap | d6 | Kit | mp |
| Kitl | batch | ic | d6 | Kit | mp |
| Kitl | batch | mp | d6 | Kit | mp |
| Kitl | constitutive | ec | d6 | Kit | mp |
| Kitl | constitutive | per | d6 | Kit | mp |
| Mdk | batch | ec | d6 | Ptprz1 | mp |
| Mdk | batch | fap | d6 | Ptprz1 | mp |
| Mdk | batch | ic | d6 | Ptprz1 | mp |
| Mdk | batch | mp | d6 | Ptprz1 | mp |
| Mdk | total | per | d6 | Ptprz1 | mp |
| Mmp13 | batch | ec | d6 | F2r | mp |
| Mmp13 | batch | fap | d6 | F2r | mp |
| Mmp13 | batch | ic | d6 | F2r | mp |
| Mmp13 | batch | mp | d6 | F2r | mp |
| Mmp9 | batch | fap | d6 | Flt1 | mp |
| Mmp9 | batch | mp | d6 | Flt1 | mp |
| Ntf3 | batch | ec | d6 | Ntrk2 | mp |
| Ntf3 | batch | fap | d6 | Ntrk2 | mp |
| Ntf3 | batch | mp | d6 | Ntrk2 | mp |
| Ntf3 | constitutive | per | d6 | Ntrk2 | mp |
| Ntf3 | batch | ec | d6 | Ngfr | mp |
| Ntf3 | batch | fap | d6 | Ngfr | mp |
| Ntf3 | batch | mp | d6 | Ngfr | mp |
| Ntf3 | constitutive | per | d6 | Ngfr | mp |
| Pthlh | batch | fap | d6 | Pth1r | mp |
| Ptn | batch | fap | d6 | Ptprz1 | mp |
| Ptn | constitutive | per | d6 | Ptprz1 | mp |
| Rspo3 | batch | fap | d6 | Lrp6 | mp |
| S100b | batch | fap | d6 | Fgfr1 | mp |
| S100b | batch | mp | d6 | Fgfr1 | mp |
| S100b | constitutive | per | d6 | Fgfr1 | mp |
| Sfrp1 | batch | fap | d6 | Fzd2 | mp |
| Sfrp1 | constitutive | per | d6 | Fzd2 | mp |
| Sfrp2 | batch | fap | d6 | Fzd2 | mp |
| Sfrp4 | batch | fap | d6 | Fzd2 | mp |
| Sfrp4 | batch | mp | d6 | Fzd2 | mp |
| Spp1 | batch | fap | d6 | Itgb5 | mp |
| Spp1 | batch | ic | d6 | Itgb5 | mp |
| Spp1 | batch | mp | d6 | Itgb5 | mp |
| Spp1 | batch | fap | d6 | Vtn | mp |
| Spp1 | batch | ic | d6 | Vtn | mp |
| Spp1 | batch | mp | d6 | Vtn | mp |
| Clu | batch | ec | d6 | Vldlr | mp |
| Clu | batch | ic | d6 | Vldlr | mp |
| Clu | batch | mp | d6 | Vldlr | mp |
| Dll1 | batch | ec | d6 | Notch1 | mp |
| Dll1 | total | mp | d6 | Notch1 | mp |
| Dll1 | batch | ec | d6 | Notch3 | mp |
| Dll1 | total | mp | d6 | Notch3 | mp |
| Dll1 | batch | ec | d6 | Notch2 | mp |
| Dll1 | total | mp | d6 | Notch2 | mp |
| Fgf9 | batch | ec | d6 | Fgfr4 | mp |
| Il15 | batch | ec | d6 | Il17ra | mp |
| Il15 | batch | ic | d6 | Il17ra | mp |
| Il15 | batch | ec | d6 | Il2rg | mp |
| Il15 | batch | ic | d6 | Il2rg | mp |
| Inhbb | batch | ec | d6 | Acvr2a | mp |
| Inhbb | batch | mp | d6 | Acvr2a | mp |
| Inhbb | constitutive | fap | d6 | Acvr2a | mp |
| Inhbb | constitutive | per | d6 | Acvr2a | mp |
| Vegfc | batch | ec | d6 | Kdr | mp |
| Vegfc | batch | mp | d6 | Kdr | mp |
| Vegfc | batch | ec | d6 | Nrp1 | mp |
| Vegfc | batch | mp | d6 | Nrp1 | mp |
| Bmp2 | batch | mp | d6 | Acvr2a | mp |
| Bmp2 | constitutive | per | d6 | Acvr2a | mp |
| Bmp2 | batch | mp | d6 | Eng | mp |
| Bmp2 | constitutive | per | d6 | Eng | mp |
| Bmp6 | batch | mp | d6 | Acvr2a | mp |
| Bmp6 | constitutive | per | d6 | Acvr2a | mp |
| Ccl11 | batch | mp | d6 | Ccr5 | mp |
| Ccl8 | batch | mp | d6 | Ccr2 | mp |
| Ccl8 | total | fap | d6 | Ccr2 | mp |
| Ccl8 | total | ic | d6 | Ccr2 | mp |
| Ccl8 | batch | mp | d6 | Ccr5 | mp |
| Ccl8 | total | fap | d6 | Ccr5 | mp |
| Ccl8 | total | ic | d6 | Ccr5 | mp |
| Clcf1 | batch | mp | d6 | Crlf1 | mp |
| Clcf1 | batch | mp | d6 | Cntfr | mp |
| Clcf1 | batch | mp | d6 | Esr1 | mp |
| Edn3 | batch | mp | d6 | Ednrb | mp |
| Il10 | batch | ic | d6 | Il10ra | mp |
| Il10 | batch | mp | d6 | Il10ra | mp |
| Il10 | batch | ic | d6 | Il10rb | mp |
| Il10 | batch | mp | d6 | Il10rb | mp |
| Jag1 | batch | mp | d6 | Notch1 | mp |
| Jag1 | constitutive | per | d6 | Notch1 | mp |
| Jag1 | batch | mp | d6 | Notch3 | mp |
| Jag1 | constitutive | per | d6 | Notch3 | mp |
| Jag1 | batch | mp | d6 | Notch2 | mp |
| Jag1 | constitutive | per | d6 | Notch2 | mp |
| Jag2 | batch | mp | d6 | Notch1 | mp |
| Jag2 | constitutive | ec | d6 | Notch1 | mp |
| Jag2 | batch | mp | d6 | Notch3 | mp |
| Jag2 | constitutive | ec | d6 | Notch3 | mp |
| Jag2 | batch | mp | d6 | Notch2 | mp |
| Jag2 | constitutive | ec | d6 | Notch2 | mp |
| Lgals3 | batch | ic | d6 | Lgals3bp | mp |
| Lgals3 | batch | mp | d6 | Lgals3bp | mp |
| Ngf | batch | mp | d6 | Ror1 | mp |
| Ngf | constitutive | per | d6 | Ror1 | mp |
| Ngf | batch | mp | d6 | Sort1 | mp |
| Ngf | constitutive | per | d6 | Sort1 | mp |
| Ngf | batch | mp | d6 | Ngfr | mp |
| Ngf | constitutive | per | d6 | Ngfr | mp |
| Nppc | batch | mp | d6 | Npr2 | mp |
| Pdap1 | batch | mp | d6 | Pdgfa | mp |
| Serpini1 | batch | mp | d6 | Plat | mp |
| Serpini1 | constitutive | per | d6 | Plat | mp |
| Sfrp5 | batch | mp | d6 | Fzd2 | mp |
| Tgfb1 | batch | mp | d6 | Eng | mp |
| Tgfb1 | batch | mp | d6 | Tgfbr3 | mp |
| Tgfb1 | batch | mp | d6 | Vtn | mp |
| Tgfb2 | batch | mp | d6 | Tgfbr3 | mp |
| Tgfb2 | constitutive | ec | d6 | Tgfbr3 | mp |
| Tgfb2 | constitutive | fap | d6 | Tgfbr3 | mp |
| Tgfb2 | constitutive | per | d6 | Tgfbr3 | mp |
| Tgfb2 | batch | mp | d6 | Vtn | mp |
| Tgfb2 | constitutive | ec | d6 | Vtn | mp |
| Tgfb2 | constitutive | fap | d6 | Vtn | mp |
| Tgfb2 | constitutive | per | d6 | Vtn | mp |
| Tnfsf12 | batch | mp | d6 | Tnfrsf12a | mp |
| Vegfa | batch | mp | d6 | Flt1 | mp |
| Vegfa | batch | mp | d6 | Kdr | mp |
| Vegfa | batch | mp | d6 | Nrp1 | mp |
| Vegfa | batch | mp | d6 | Vtn | mp |
| Csf1 | batch | ic | d6 | Csf1r | mp |
| Hgf | batch | ic | d6 | Vtn | mp |
| Tnfsf13 | batch | ic | d6 | Tnfrsf12a | mp |
| Angpt1 | constitutive | fap | d6 | Tek | mp |
| Fgf7 | constitutive | fap | d6 | Fgfr4 | mp |
| Fgf7 | constitutive | fap | d6 | Nrp1 | mp |
| Efnb2 | constitutive | ec | d6 | Ephb1 | mp |
| Cmtm8 | constitutive | ec | d6 | Egfr | mp |
| Edn1 | constitutive | ec | d6 | Ednrb | mp |
| Dhh | constitutive | ec | d6 | Ptch1 | mp |
| Dhh | constitutive | mp | d6 | Ptch1 | mp |
| Dhh | constitutive | per | d6 | Ptch1 | mp |
| Angpt2 | constitutive | ec | d6 | Tek | mp |
| Angpt2 | constitutive | ic | d6 | Tek | mp |
| Angpt2 | constitutive | per | d6 | Tek | mp |
| Il18 | constitutive | ic | d6 | Il18rap | mp |
| Il18 | constitutive | mp | d6 | Il18rap | mp |
| Pgf | constitutive | per | d6 | Flt1 | mp |
| Pgf | constitutive | per | d6 | Nrp1 | mp |
| Bmp2 | batch | fap | d7 | Acvr2a | mp |
| Bmp2 | batch | mp | d7 | Acvr2a | mp |
| Bmp2 | constitutive | per | d7 | Acvr2a | mp |
| Bmp2 | batch | fap | d7 | Bmpr1a | mp |
| Bmp2 | batch | mp | d7 | Bmpr1a | mp |
| Bmp2 | constitutive | per | d7 | Bmpr1a | mp |
| Bmp2 | batch | fap | d7 | Eng | mp |
| Bmp2 | batch | mp | d7 | Eng | mp |
| Bmp2 | constitutive | per | d7 | Eng | mp |
| Bmp4 | batch | fap | d7 | Bmpr1a | mp |
| Bmp4 | batch | mp | d7 | Bmpr1a | mp |
| Bmp4 | constitutive | ec | d7 | Bmpr1a | mp |
| Bmp4 | constitutive | per | d7 | Bmpr1a | mp |
| Bmp6 | batch | ec | d7 | Acvr2a | mp |
| Bmp6 | batch | fap | d7 | Acvr2a | mp |
| Bmp6 | batch | mp | d7 | Acvr2a | mp |
| Bmp6 | constitutive | per | d7 | Acvr2a | mp |
| Bmp6 | batch | ec | d7 | Bmpr1a | mp |
| Bmp6 | batch | fap | d7 | Bmpr1a | mp |
| Bmp6 | batch | mp | d7 | Bmpr1a | mp |
| Bmp6 | constitutive | per | d7 | Bmpr1a | mp |
| Bmp7 | batch | fap | d7 | Acvr2a | mp |
| Bmp7 | batch | fap | d7 | Bmpr1a | mp |
| Bmp7 | batch | fap | d7 | Eng | mp |
| C4b | batch | fap | d7 | C3ar1 | mp |
| C4b | batch | ic | d7 | C3ar1 | mp |
| C4b | batch | mp | d7 | C3ar1 | mp |
| C4b | constitutive | per | d7 | C3ar1 | mp |
| Ccl11 | batch | fap | d7 | Ccr5 | mp |
| Ccl11 | batch | mp | d7 | Ccr5 | mp |
| Clu | batch | ec | d7 | Vldlr | mp |
| Clu | batch | fap | d7 | Vldlr | mp |
| Clu | batch | ic | d7 | Vldlr | mp |
| Clu | batch | mp | d7 | Vldlr | mp |
| Crlf1 | batch | ec | d7 | Ctf1 | mp |
| Crlf1 | batch | fap | d7 | Ctf1 | mp |
| Crlf1 | batch | mp | d7 | Ctf1 | mp |
| Csf2 | batch | fap | d7 | Csf2ra | mp |
| Csf2 | batch | fap | d7 | Csf2rb | mp |
| Cx3cl1 | batch | fap | d7 | Cx3cr1 | mp |
| Cx3cl1 | batch | mp | d7 | Cx3cr1 | mp |
| Cx3cl1 | total | ec | d7 | Cx3cr1 | mp |
| Cx3cl1 | total | per | d7 | Cx3cr1 | mp |
| Edn1 | batch | fap | d7 | Ednrb | mp |
| Edn1 | constitutive | ec | d7 | Ednrb | mp |
| Fgf1 | batch | fap | d7 | Fgfr1 | mp |
| Fgf1 | constitutive | per | d7 | Fgfr1 | mp |
| Fgf1 | batch | fap | d7 | Fgfr4 | mp |
| Fgf1 | constitutive | per | d7 | Fgfr4 | mp |
| Fgf18 | batch | fap | d7 | Fgfr4 | mp |
| Gas6 | batch | fap | d7 | Axl | mp |
| Gas6 | batch | mp | d7 | Axl | mp |
| Hbegf | batch | fap | d7 | Cd44 | mp |
| Hbegf | batch | mp | d7 | Cd44 | mp |
| Hbegf | constitutive | ec | d7 | Cd44 | mp |
| Hbegf | constitutive | per | d7 | Cd44 | mp |
| Hbegf | batch | fap | d7 | Egfr | mp |
| Hbegf | batch | mp | d7 | Egfr | mp |
| Hbegf | constitutive | ec | d7 | Egfr | mp |
| Hbegf | constitutive | per | d7 | Egfr | mp |
| Igf1 | batch | ec | d7 | Igfbp2 | mp |
| Igf1 | batch | fap | d7 | Igfbp2 | mp |
| Igf1 | batch | ic | d7 | Igfbp2 | mp |
| Igf1 | batch | ec | d7 | Igfbp3 | mp |
| Igf1 | batch | fap | d7 | Igfbp3 | mp |
| Igf1 | batch | ic | d7 | Igfbp3 | mp |
| Igf1 | batch | ec | d7 | Igfbp4 | mp |
| Igf1 | batch | fap | d7 | Igfbp4 | mp |
| Igf1 | batch | ic | d7 | Igfbp4 | mp |
| Igf1 | batch | ec | d7 | Igfbp5 | mp |
| Igf1 | batch | fap | d7 | Igfbp5 | mp |
| Igf1 | batch | ic | d7 | Igfbp5 | mp |
| Igf1 | batch | ec | d7 | Igf1r | mp |
| Igf1 | batch | fap | d7 | Igf1r | mp |
| Igf1 | batch | ic | d7 | Igf1r | mp |
| Igf2 | batch | ec | d7 | Igf2r | mp |
| Igf2 | batch | fap | d7 | Igf2r | mp |
| Igf2 | batch | ic | d7 | Igf2r | mp |
| Igf2 | batch | mp | d7 | Igf2r | mp |
| Igf2 | batch | ec | d7 | Vtn | mp |
| Igf2 | batch | fap | d7 | Vtn | mp |
| Igf2 | batch | ic | d7 | Vtn | mp |
| Igf2 | batch | mp | d7 | Vtn | mp |
| Igf2 | batch | ec | d7 | Igf1r | mp |
| Igf2 | batch | fap | d7 | Igf1r | mp |
| Igf2 | batch | ic | d7 | Igf1r | mp |
| Igf2 | batch | mp | d7 | Igf1r | mp |
| Il18 | batch | fap | d7 | Il18rap | mp |
| Il18 | constitutive | ic | d7 | Il18rap | mp |
| Il18 | constitutive | mp | d7 | Il18rap | mp |
| Mdk | batch | ec | d7 | Ptprz1 | mp |
| Mdk | batch | fap | d7 | Ptprz1 | mp |
| Mdk | batch | ic | d7 | Ptprz1 | mp |
| Mdk | batch | mp | d7 | Ptprz1 | mp |
| Mdk | total | per | d7 | Ptprz1 | mp |
| Mmp13 | batch | ec | d7 | F2r | mp |
| Mmp13 | batch | fap | d7 | F2r | mp |
| Mmp13 | batch | mp | d7 | F2r | mp |
| Nov | batch | ec | d7 | Notch1 | mp |
| Nov | batch | fap | d7 | Notch1 | mp |
| Ntf3 | batch | ec | d7 | Ntrk2 | mp |
| Ntf3 | batch | fap | d7 | Ntrk2 | mp |
| Ntf3 | batch | mp | d7 | Ntrk2 | mp |
| Ntf3 | constitutive | per | d7 | Ntrk2 | mp |
| Ntf3 | batch | ec | d7 | Ngfr | mp |
| Ntf3 | batch | fap | d7 | Ngfr | mp |
| Ntf3 | batch | mp | d7 | Ngfr | mp |
| Ntf3 | constitutive | per | d7 | Ngfr | mp |
| Pgf | batch | fap | d7 | Flt1 | mp |
| Pgf | constitutive | per | d7 | Flt1 | mp |
| Pgf | batch | fap | d7 | Nrp1 | mp |
| Pgf | constitutive | per | d7 | Nrp1 | mp |
| Pthlh | batch | fap | d7 | Pth1r | mp |
| Ptn | batch | fap | d7 | Ptprz1 | mp |
| Ptn | constitutive | per | d7 | Ptprz1 | mp |
| S100b | batch | fap | d7 | Fgfr1 | mp |
| S100b | batch | mp | d7 | Fgfr1 | mp |
| S100b | constitutive | per | d7 | Fgfr1 | mp |
| Sfrp1 | batch | fap | d7 | Fzd2 | mp |
| Sfrp1 | constitutive | per | d7 | Fzd2 | mp |
| Sfrp4 | batch | ec | d7 | Fzd2 | mp |
| Sfrp4 | batch | fap | d7 | Fzd2 | mp |
| Sfrp4 | batch | mp | d7 | Fzd2 | mp |
| Vegfa | batch | fap | d7 | Flt1 | mp |
| Vegfa | batch | mp | d7 | Flt1 | mp |
| Vegfa | batch | fap | d7 | Kdr | mp |
| Vegfa | batch | mp | d7 | Kdr | mp |
| Vegfa | batch | fap | d7 | Nrp1 | mp |
| Vegfa | batch | mp | d7 | Nrp1 | mp |
| Vegfa | batch | fap | d7 | Vtn | mp |
| Vegfa | batch | mp | d7 | Vtn | mp |
| Wnt11 | batch | fap | d7 | Fzd4 | mp |
| Wnt11 | batch | ic | d7 | Fzd4 | mp |
| Wnt2 | batch | fap | d7 | Sfrp1 | mp |
| Wnt2 | batch | fap | d7 | Fzd1 | mp |
| Dll1 | batch | ec | d7 | Notch1 | mp |
| Dll1 | total | mp | d7 | Notch1 | mp |
| Dll1 | batch | ec | d7 | Notch3 | mp |
| Dll1 | total | mp | d7 | Notch3 | mp |
| Dll1 | batch | ec | d7 | Notch2 | mp |
| Dll1 | total | mp | d7 | Notch2 | mp |
| Fgf9 | batch | ec | d7 | Fgfr4 | mp |
| Il15 | batch | ec | d7 | Il17ra | mp |
| Il15 | batch | ic | d7 | Il17ra | mp |
| Il15 | batch | ec | d7 | Il2rg | mp |
| Il15 | batch | ic | d7 | Il2rg | mp |
| Sfrp5 | batch | ec | d7 | Fzd2 | mp |
| Sfrp5 | batch | mp | d7 | Fzd2 | mp |
| Vegfc | batch | ec | d7 | Kdr | mp |
| Vegfc | batch | mp | d7 | Kdr | mp |
| Vegfc | batch | ec | d7 | Nrp1 | mp |
| Vegfc | batch | mp | d7 | Nrp1 | mp |
| Areg | batch | mp | d7 | Egfr | mp |
| Ccl3 | batch | mp | d7 | Ccr5 | mp |
| Ccl4 | batch | mp | d7 | Ccr5 | mp |
| Ccl8 | batch | mp | d7 | Ccr2 | mp |
| Ccl8 | total | fap | d7 | Ccr2 | mp |
| Ccl8 | total | ic | d7 | Ccr2 | mp |
| Ccl8 | batch | mp | d7 | Ccr5 | mp |
| Ccl8 | total | fap | d7 | Ccr5 | mp |
| Ccl8 | total | ic | d7 | Ccr5 | mp |
| Clcf1 | batch | ic | d7 | Crlf1 | mp |
| Clcf1 | batch | mp | d7 | Crlf1 | mp |
| Clcf1 | batch | ic | d7 | Cntfr | mp |
| Clcf1 | batch | mp | d7 | Cntfr | mp |
| Clcf1 | batch | ic | d7 | Esr1 | mp |
| Clcf1 | batch | mp | d7 | Esr1 | mp |
| Ctsg | batch | ic | d7 | Met | mp |
| Ctsg | batch | mp | d7 | Met | mp |
| Edn3 | batch | mp | d7 | Ednrb | mp |
| Gdnf | batch | mp | d7 | Gfra1 | mp |
| Gdnf | batch | mp | d7 | Gfra2 | mp |
| Gdnf | batch | mp | d7 | Ret | mp |
| Gmfb | batch | mp | d7 | Egfr | mp |
| Il10 | batch | ic | d7 | Il10ra | mp |
| Il10 | batch | mp | d7 | Il10ra | mp |
| Il10 | batch | ic | d7 | Il10rb | mp |
| Il10 | batch | mp | d7 | Il10rb | mp |
| Inhbb | batch | mp | d7 | Acvr2a | mp |
| Inhbb | constitutive | fap | d7 | Acvr2a | mp |
| Inhbb | constitutive | per | d7 | Acvr2a | mp |
| Jag1 | batch | mp | d7 | Notch1 | mp |
| Jag1 | constitutive | per | d7 | Notch1 | mp |
| Jag1 | batch | mp | d7 | Notch3 | mp |
| Jag1 | constitutive | per | d7 | Notch3 | mp |
| Jag1 | batch | mp | d7 | Notch2 | mp |
| Jag1 | constitutive | per | d7 | Notch2 | mp |
| Jag2 | batch | mp | d7 | Notch1 | mp |
| Jag2 | constitutive | ec | d7 | Notch1 | mp |
| Jag2 | batch | mp | d7 | Notch3 | mp |
| Jag2 | constitutive | ec | d7 | Notch3 | mp |
| Jag2 | batch | mp | d7 | Notch2 | mp |
| Jag2 | constitutive | ec | d7 | Notch2 | mp |
| Kitl | batch | ic | d7 | Kit | mp |
| Kitl | batch | mp | d7 | Kit | mp |
| Kitl | constitutive | ec | d7 | Kit | mp |
| Kitl | constitutive | per | d7 | Kit | mp |
| Lgals3 | batch | ic | d7 | Lgals3bp | mp |
| Lgals3 | batch | mp | d7 | Lgals3bp | mp |
| Mmp9 | batch | ic | d7 | Flt1 | mp |
| Mmp9 | batch | mp | d7 | Flt1 | mp |
| Ngf | batch | mp | d7 | Ror1 | mp |
| Ngf | constitutive | per | d7 | Ror1 | mp |
| Ngf | batch | mp | d7 | Sort1 | mp |
| Ngf | constitutive | per | d7 | Sort1 | mp |
| Ngf | batch | mp | d7 | Ngfr | mp |
| Ngf | constitutive | per | d7 | Ngfr | mp |
| Nppc | batch | mp | d7 | Npr2 | mp |
| Pdap1 | batch | mp | d7 | Pdgfa | mp |
| Serpini1 | batch | mp | d7 | Plat | mp |
| Serpini1 | constitutive | per | d7 | Plat | mp |
| Spp1 | batch | mp | d7 | Itgb5 | mp |
| Spp1 | batch | mp | d7 | Vtn | mp |
| Tgfb1 | batch | mp | d7 | Eng | mp |
| Tgfb1 | batch | mp | d7 | Tgfbr3 | mp |
| Tgfb1 | batch | mp | d7 | Vtn | mp |
| Tgfb2 | batch | mp | d7 | Tgfbr3 | mp |
| Tgfb2 | constitutive | ec | d7 | Tgfbr3 | mp |
| Tgfb2 | constitutive | fap | d7 | Tgfbr3 | mp |
| Tgfb2 | constitutive | per | d7 | Tgfbr3 | mp |
| Tgfb2 | batch | mp | d7 | Vtn | mp |
| Tgfb2 | constitutive | ec | d7 | Vtn | mp |
| Tgfb2 | constitutive | fap | d7 | Vtn | mp |
| Tgfb2 | constitutive | per | d7 | Vtn | mp |
| Tnfsf12 | batch | mp | d7 | Tnfrsf12a | mp |
| Ccl5 | batch | ic | d7 | Ccr5 | mp |
| Ccl5 | batch | ic | d7 | Sdc4 | mp |
| Csf1 | batch | ic | d7 | Csf1r | mp |
| Angpt1 | constitutive | fap | d7 | Tek | mp |
| Fgf7 | constitutive | fap | d7 | Fgfr4 | mp |
| Fgf7 | constitutive | fap | d7 | Nrp1 | mp |
| Bmp5 | constitutive | ec | d7 | Bmpr1a | mp |
| Bmp5 | constitutive | fap | d7 | Bmpr1a | mp |
| Bmp5 | constitutive | per | d7 | Bmpr1a | mp |
| Efnb2 | constitutive | ec | d7 | Ephb1 | mp |
| Cmtm8 | constitutive | ec | d7 | Egfr | mp |
| Dhh | constitutive | ec | d7 | Ptch1 | mp |
| Dhh | constitutive | mp | d7 | Ptch1 | mp |
| Dhh | constitutive | per | d7 | Ptch1 | mp |
| Angpt2 | constitutive | ec | d7 | Tek | mp |
| Angpt2 | constitutive | ic | d7 | Tek | mp |
| Angpt2 | constitutive | per | d7 | Tek | mp |
| Bmp2 | batch | fap | d10 | Eng | mp |
| Bmp2 | batch | mp | d10 | Eng | mp |
| Bmp2 | constitutive | per | d10 | Eng | mp |
| Bmp7 | batch | fap | d10 | Eng | mp |
| Crlf1 | batch | fap | d10 | Ctf1 | mp |
| Crlf1 | batch | mp | d10 | Ctf1 | mp |
| Edn1 | batch | fap | d10 | Ednrb | mp |
| Edn1 | constitutive | ec | d10 | Ednrb | mp |
| Fgf1 | batch | fap | d10 | Fgfr1 | mp |
| Fgf1 | batch | mp | d10 | Fgfr1 | mp |
| Fgf1 | constitutive | per | d10 | Fgfr1 | mp |
| Fgf1 | batch | fap | d10 | Fgfr4 | mp |
| Fgf1 | batch | mp | d10 | Fgfr4 | mp |
| Fgf1 | constitutive | per | d10 | Fgfr4 | mp |
| Fgf18 | batch | fap | d10 | Fgfr4 | mp |
| Fgf18 | batch | per | d10 | Fgfr4 | mp |
| Gas6 | batch | fap | d10 | Axl | mp |
| Gas6 | batch | mp | d10 | Axl | mp |
| Igf1 | batch | ec | d10 | Igfbp3 | mp |
| Igf1 | batch | fap | d10 | Igfbp3 | mp |
| Igf1 | batch | ic | d10 | Igfbp3 | mp |
| Igf1 | batch | ec | d10 | Igfbp4 | mp |
| Igf1 | batch | fap | d10 | Igfbp4 | mp |
| Igf1 | batch | ic | d10 | Igfbp4 | mp |
| Igf1 | batch | ec | d10 | Igfbp5 | mp |
| Igf1 | batch | fap | d10 | Igfbp5 | mp |
| Igf1 | batch | ic | d10 | Igfbp5 | mp |
| Igf1 | batch | ec | d10 | Igfbp6 | mp |
| Igf1 | batch | fap | d10 | Igfbp6 | mp |
| Igf1 | batch | ic | d10 | Igfbp6 | mp |
| Igf1 | batch | ec | d10 | Igfbp7 | mp |
| Igf1 | batch | fap | d10 | Igfbp7 | mp |
| Igf1 | batch | ic | d10 | Igfbp7 | mp |
| Igf1 | batch | ec | d10 | Igf1r | mp |
| Igf1 | batch | fap | d10 | Igf1r | mp |
| Igf1 | batch | ic | d10 | Igf1r | mp |
| Igf2 | batch | ec | d10 | Igf2r | mp |
| Igf2 | batch | fap | d10 | Igf2r | mp |
| Igf2 | batch | mp | d10 | Igf2r | mp |
| Igf2 | batch | ec | d10 | Vtn | mp |
| Igf2 | batch | fap | d10 | Vtn | mp |
| Igf2 | batch | mp | d10 | Vtn | mp |
| Igf2 | batch | ec | d10 | Igf1r | mp |
| Igf2 | batch | fap | d10 | Igf1r | mp |
| Igf2 | batch | mp | d10 | Igf1r | mp |
| Kitl | batch | fap | d10 | Kit | mp |
| Kitl | constitutive | ec | d10 | Kit | mp |
| Kitl | constitutive | per | d10 | Kit | mp |
| Mdk | batch | fap | d10 | Lrp1 | mp |
| Mdk | batch | mp | d10 | Lrp1 | mp |
| Mdk | batch | fap | d10 | Ptprz1 | mp |
| Mdk | batch | mp | d10 | Ptprz1 | mp |
| Nov | batch | fap | d10 | Notch1 | mp |
| Nov | batch | per | d10 | Notch1 | mp |
| Ntf3 | batch | ec | d10 | Ntrk2 | mp |
| Ntf3 | batch | fap | d10 | Ntrk2 | mp |
| Ntf3 | batch | mp | d10 | Ntrk2 | mp |
| Ntf3 | constitutive | per | d10 | Ntrk2 | mp |
| Ntf3 | batch | ec | d10 | Ngfr | mp |
| Ntf3 | batch | fap | d10 | Ngfr | mp |
| Ntf3 | batch | mp | d10 | Ngfr | mp |
| Ntf3 | constitutive | per | d10 | Ngfr | mp |
| Pgf | batch | fap | d10 | Flt1 | mp |
| Pgf | batch | mp | d10 | Flt1 | mp |
| Pgf | constitutive | per | d10 | Flt1 | mp |
| Pthlh | batch | fap | d10 | Pth1r | mp |
| Ptn | batch | fap | d10 | Ptprs | mp |
| Ptn | batch | mp | d10 | Ptprs | mp |
| Ptn | constitutive | per | d10 | Ptprs | mp |
| Ptn | batch | fap | d10 | Ptprz1 | mp |
| Ptn | batch | mp | d10 | Ptprz1 | mp |
| Ptn | constitutive | per | d10 | Ptprz1 | mp |
| S100b | batch | fap | d10 | Fgfr1 | mp |
| S100b | batch | mp | d10 | Fgfr1 | mp |
| S100b | constitutive | per | d10 | Fgfr1 | mp |
| Sfrp1 | batch | fap | d10 | Fzd2 | mp |
| Sfrp1 | constitutive | per | d10 | Fzd2 | mp |
| Sfrp4 | batch | fap | d10 | Fzd2 | mp |
| Sfrp4 | batch | mp | d10 | Fzd2 | mp |
| Vegfa | batch | fap | d10 | Flt1 | mp |
| Vegfa | batch | ic | d10 | Flt1 | mp |
| Vegfa | batch | mp | d10 | Flt1 | mp |
| Vegfa | batch | fap | d10 | Kdr | mp |
| Vegfa | batch | ic | d10 | Kdr | mp |
| Vegfa | batch | mp | d10 | Kdr | mp |
| Vegfa | batch | fap | d10 | Vtn | mp |
| Vegfa | batch | ic | d10 | Vtn | mp |
| Vegfa | batch | mp | d10 | Vtn | mp |
| Wnt2 | batch | fap | d10 | Sfrp1 | mp |
| Wnt2 | batch | fap | d10 | Fzd1 | mp |
| Dll1 | batch | ec | d10 | Notch1 | mp |
| Dll1 | batch | ec | d10 | Notch3 | mp |
| Dll1 | batch | ec | d10 | Notch2 | mp |
| Fgf9 | batch | ec | d10 | Fgfr4 | mp |
| Il15 | batch | ec | d10 | Il17ra | mp |
| Il15 | batch | ic | d10 | Il17ra | mp |
| Vegfc | batch | ec | d10 | Kdr | mp |
| Clcf1 | batch | ic | d10 | Crlf1 | mp |
| Clcf1 | batch | mp | d10 | Crlf1 | mp |
| Clcf1 | batch | ic | d10 | Cntfr | mp |
| Clcf1 | batch | mp | d10 | Cntfr | mp |
| Edn3 | batch | mp | d10 | Ednrb | mp |
| Jag1 | batch | ic | d10 | Notch1 | mp |
| Jag1 | batch | mp | d10 | Notch1 | mp |
| Jag1 | constitutive | per | d10 | Notch1 | mp |
| Jag1 | batch | ic | d10 | Notch3 | mp |
| Jag1 | batch | mp | d10 | Notch3 | mp |
| Jag1 | constitutive | per | d10 | Notch3 | mp |
| Jag1 | batch | ic | d10 | Notch2 | mp |
| Jag1 | batch | mp | d10 | Notch2 | mp |
| Jag1 | constitutive | per | d10 | Notch2 | mp |
| Jag2 | batch | mp | d10 | Notch1 | mp |
| Jag2 | constitutive | ec | d10 | Notch1 | mp |
| Jag2 | batch | mp | d10 | Notch3 | mp |
| Jag2 | constitutive | ec | d10 | Notch3 | mp |
| Jag2 | batch | mp | d10 | Notch2 | mp |
| Jag2 | constitutive | ec | d10 | Notch2 | mp |
| Ngf | batch | mp | d10 | Sort1 | mp |
| Ngf | constitutive | per | d10 | Sort1 | mp |
| Ngf | batch | mp | d10 | Ngfr | mp |
| Ngf | constitutive | per | d10 | Ngfr | mp |
| Nppc | batch | mp | d10 | Npr2 | mp |
| Serpini1 | batch | mp | d10 | Plat | mp |
| Serpini1 | constitutive | per | d10 | Plat | mp |
| Sfrp2 | batch | mp | d10 | Fzd2 | mp |
| Sfrp5 | batch | mp | d10 | Fzd2 | mp |
| Sfrp5 | batch | per | d10 | Fzd2 | mp |
| Tgfb2 | batch | mp | d10 | Tgfbr3 | mp |
| Tgfb2 | constitutive | ec | d10 | Tgfbr3 | mp |
| Tgfb2 | constitutive | fap | d10 | Tgfbr3 | mp |
| Tgfb2 | constitutive | per | d10 | Tgfbr3 | mp |
| Tgfb2 | batch | mp | d10 | Vtn | mp |
| Tgfb2 | constitutive | ec | d10 | Vtn | mp |
| Tgfb2 | constitutive | fap | d10 | Vtn | mp |
| Tgfb2 | constitutive | per | d10 | Vtn | mp |
| Tnfsf12 | batch | mp | d10 | Tnfrsf12a | mp |
| Tnfsf12 | total | ec | d10 | Tnfrsf12a | mp |
| Tnfsf12 | total | ic | d10 | Tnfrsf12a | mp |
| Tnfsf12 | total | per | d10 | Tnfrsf12a | mp |
| Vegfb | batch | mp | d10 | Flt1 | mp |
| Ccl5 | batch | ic | d10 | Sdc4 | mp |
| Il1b | batch | ic | d10 | Adrb2 | mp |
| Mmp9 | batch | ic | d10 | Flt1 | mp |
| Pdgfb | batch | ic | d10 | Lrp1 | mp |
| Pdgfb | constitutive | ec | d10 | Lrp1 | mp |
| Pdgfb | batch | ic | d10 | Pdgfrb | mp |
| Pdgfb | constitutive | ec | d10 | Pdgfrb | mp |
| Pf4 | batch | ic | d10 | Ldlr | mp |
| Ppbp | batch | ic | d10 | Gabbr1 | mp |
| Ppbp | batch | ic | d10 | Slc1a5 | mp |
| Tnf | batch | ic | d10 | Ptprz1 | mp |
| Adm | batch | per | d10 | Calcrl | mp |
| Gdnf | batch | per | d10 | Gfra2 | mp |
| Tgfb1 | total | ec | d10 | Eng | mp |
| Tgfb1 | total | ic | d10 | Eng | mp |
| Tgfb1 | total | ec | d10 | Tgfbr3 | mp |
| Tgfb1 | total | ic | d10 | Tgfbr3 | mp |
| Tgfb1 | total | ec | d10 | Vtn | mp |
| Tgfb1 | total | ic | d10 | Vtn | mp |
| Fgf7 | constitutive | fap | d10 | Fgfr4 | mp |
| Efnb2 | constitutive | ec | d10 | Ephb1 | mp |
| Dhh | constitutive | ec | d10 | Ptch1 | mp |
| Dhh | constitutive | mp | d10 | Ptch1 | mp |
| Dhh | constitutive | per | d10 | Ptch1 | mp |
| Ccl11 | batch | fap | d0 | Ccr3 | ic |
| Ccl11 | batch | mp | d0 | Ccr3 | ic |
| Ccl11 | batch | per | d0 | Ccr3 | ic |
| Ccl11 | batch | fap | d0 | Cxcr3 | ic |
| Ccl11 | batch | mp | d0 | Cxcr3 | ic |
| Ccl11 | batch | per | d0 | Cxcr3 | ic |
| Ccl11 | batch | fap | d0 | Dpp4 | ic |
| Ccl11 | batch | mp | d0 | Dpp4 | ic |
| Ccl11 | batch | per | d0 | Dpp4 | ic |
| Crlf1 | batch | ec | d0 | Ctf1 | ic |
| Crlf1 | batch | fap | d0 | Ctf1 | ic |
| Crlf1 | batch | mp | d0 | Ctf1 | ic |
| Cxcl13 | batch | fap | d0 | Cxcr3 | ic |
| Cxcl13 | batch | fap | d0 | Cxcr5 | ic |
| Edn1 | batch | fap | d0 | Ednrb | ic |
| Edn1 | constitutive | ec | d0 | Ednrb | ic |
| Fgf1 | batch | fap | d0 | Fgfr1 | ic |
| Fgf1 | constitutive | per | d0 | Fgfr1 | ic |
| Gas6 | batch | fap | d0 | Axl | ic |
| Gas6 | batch | mp | d0 | Axl | ic |
| Igf1 | batch | ec | d0 | Igfbp3 | ic |
| Igf1 | batch | fap | d0 | Igfbp3 | ic |
| Igf1 | batch | ic | d0 | Igfbp3 | ic |
| Igf1 | batch | ec | d0 | Igfbp6 | ic |
| Igf1 | batch | fap | d0 | Igfbp6 | ic |
| Igf1 | batch | ic | d0 | Igfbp6 | ic |
| Igf1 | batch | ec | d0 | Igfbp7 | ic |
| Igf1 | batch | fap | d0 | Igfbp7 | ic |
| Igf1 | batch | ic | d0 | Igfbp7 | ic |
| Il16 | batch | fap | d0 | Cd4 | ic |
| Il16 | batch | ic | d0 | Cd4 | ic |
| Il18 | batch | fap | d0 | Il18rap | ic |
| Il18 | constitutive | ic | d0 | Il18rap | ic |
| Il18 | constitutive | mp | d0 | Il18rap | ic |
| Kitl | batch | fap | d0 | Kit | ic |
| Kitl | batch | ic | d0 | Kit | ic |
| Kitl | batch | mp | d0 | Kit | ic |
| Kitl | constitutive | ec | d0 | Kit | ic |
| Kitl | constitutive | per | d0 | Kit | ic |
| Nov | batch | ec | d0 | Itgb3 | ic |
| Nov | batch | fap | d0 | Itgb3 | ic |
| Nov | batch | per | d0 | Itgb3 | ic |
| Pgf | batch | fap | d0 | Nrp2 | ic |
| Pgf | constitutive | per | d0 | Nrp2 | ic |
| Pgf | batch | fap | d0 | Nrp1 | ic |
| Pgf | constitutive | per | d0 | Nrp1 | ic |
| Pthlh | batch | fap | d0 | Pth1r | ic |
| Ptn | batch | fap | d0 | Ptprs | ic |
| Ptn | constitutive | per | d0 | Ptprs | ic |
| S100b | batch | fap | d0 | Fgfr1 | ic |
| S100b | batch | mp | d0 | Fgfr1 | ic |
| S100b | constitutive | per | d0 | Fgfr1 | ic |
| Vegfa | batch | fap | d0 | Itga9 | ic |
| Vegfa | batch | mp | d0 | Itga9 | ic |
| Vegfa | batch | fap | d0 | Nrp2 | ic |
| Vegfa | batch | mp | d0 | Nrp2 | ic |
| Vegfa | batch | fap | d0 | Nrp1 | ic |
| Vegfa | batch | mp | d0 | Nrp1 | ic |
| Wnt2 | batch | fap | d0 | Sfrp1 | ic |
| Cxcl10 | batch | ec | d0 | Ccr3 | ic |
| Cxcl10 | batch | ec | d0 | Cxcr3 | ic |
| Cxcl10 | batch | ec | d0 | Dpp4 | ic |
| Cxcl9 | batch | ec | d0 | Ccr3 | ic |
| Cxcl9 | batch | ic | d0 | Ccr3 | ic |
| Cxcl9 | batch | ec | d0 | Cxcr3 | ic |
| Cxcl9 | batch | ic | d0 | Cxcr3 | ic |
| Cxcl9 | batch | ec | d0 | Dpp4 | ic |
| Cxcl9 | batch | ic | d0 | Dpp4 | ic |
| Dll1 | batch | ec | d0 | Notch2 | ic |
| Esm1 | batch | ec | d0 | Itgal | ic |
| Esm1 | batch | mp | d0 | Itgal | ic |
| Il15 | batch | ec | d0 | Il2rb | ic |
| Il15 | batch | ic | d0 | Il2rb | ic |
| Vegfc | batch | ec | d0 | Itga9 | ic |
| Vegfc | batch | mp | d0 | Itga9 | ic |
| Vegfc | batch | ec | d0 | Nrp2 | ic |
| Vegfc | batch | mp | d0 | Nrp2 | ic |
| Vegfc | batch | ec | d0 | Nrp1 | ic |
| Vegfc | batch | mp | d0 | Nrp1 | ic |
| Clcf1 | batch | ic | d0 | Crlf1 | ic |
| Clcf1 | batch | mp | d0 | Crlf1 | ic |
| Clcf1 | batch | ic | d0 | Ncoa3 | ic |
| Clcf1 | batch | mp | d0 | Ncoa3 | ic |
| Cx3cl1 | batch | mp | d0 | Cx3cr1 | ic |
| Cxcl12 | batch | ic | d0 | Dpp4 | ic |
| Cxcl12 | batch | mp | d0 | Dpp4 | ic |
| Cxcl12 | constitutive | ec | d0 | Dpp4 | ic |
| Cxcl12 | constitutive | per | d0 | Dpp4 | ic |
| Edn3 | batch | mp | d0 | Ednrb | ic |
| Gdnf | batch | mp | d0 | Gfra2 | ic |
| Gdnf | batch | per | d0 | Gfra2 | ic |
| Gmfb | batch | mp | d0 | Itpr3 | ic |
| Jag1 | batch | mp | d0 | Notch2 | ic |
| Jag1 | constitutive | per | d0 | Notch2 | ic |
| Jag2 | batch | mp | d0 | Notch2 | ic |
| Jag2 | constitutive | ec | d0 | Notch2 | ic |
| Rabep1 | batch | mp | d0 | Lsr | ic |
| Tgfb3 | batch | mp | d0 | Acvrl1 | ic |
| Tnfsf12 | batch | mp | d0 | Tnfrsf14 | ic |
| Btla | batch | ic | d0 | Tnfrsf14 | ic |
| Ccl22 | batch | ic | d0 | Dpp4 | ic |
| Ccl24 | batch | ic | d0 | Ccr3 | ic |
| Ccl5 | batch | ic | d0 | Ccr3 | ic |
| Ccl5 | batch | ic | d0 | Cxcr3 | ic |
| Ccl5 | batch | ic | d0 | Cxcr5 | ic |
| Cxcl16 | batch | ic | d0 | Cxcr6 | ic |
| Ifng | batch | ic | d0 | Ifngr2 | ic |
| Ifng | batch | ic | d0 | Ifngr1 | ic |
| Il6 | batch | ic | d0 | Il6ra | ic |
| Pdgfb | batch | ic | d0 | Pdgfrb | ic |
| Pdgfb | constitutive | ec | d0 | Pdgfrb | ic |
| Fgf7 | constitutive | fap | d0 | Nrp1 | ic |
| Tgfb2 | constitutive | ec | d0 | Tgfbr3 | ic |
| Tgfb2 | constitutive | fap | d0 | Tgfbr3 | ic |
| Tgfb2 | constitutive | per | d0 | Tgfbr3 | ic |
| Lif | constitutive | mp | d0 | Lifr | ic |
| Ebi3 | constitutive | ic | d0 | Il27ra | ic |
| Nampt | constitutive | ic | d0 | Adora2a | ic |
| Apln | batch | ec | d1 | Aplnr | ic |
| Apln | batch | fap | d1 | Aplnr | ic |
| Ccl2 | batch | ec | d1 | Ccr1 | ic |
| Ccl2 | batch | fap | d1 | Ccr1 | ic |
| Ccl2 | batch | ic | d1 | Ccr1 | ic |
| Ccl2 | batch | per | d1 | Ccr1 | ic |
| Ccl2 | batch | ec | d1 | Ccr5 | ic |
| Ccl2 | batch | fap | d1 | Ccr5 | ic |
| Ccl2 | batch | ic | d1 | Ccr5 | ic |
| Ccl2 | batch | per | d1 | Ccr5 | ic |
| Ccl7 | batch | ec | d1 | Ccr1 | ic |
| Ccl7 | batch | fap | d1 | Ccr1 | ic |
| Ccl7 | batch | ic | d1 | Ccr1 | ic |
| Ccl7 | batch | per | d1 | Ccr1 | ic |
| Ccl7 | batch | ec | d1 | Ccr5 | ic |
| Ccl7 | batch | fap | d1 | Ccr5 | ic |
| Ccl7 | batch | ic | d1 | Ccr5 | ic |
| Ccl7 | batch | per | d1 | Ccr5 | ic |
| Csf2 | batch | fap | d1 | Csf3r | ic |
| Cxcl1 | batch | ec | d1 | Cxcr2 | ic |
| Cxcl1 | batch | fap | d1 | Cxcr2 | ic |
| Cxcl1 | batch | ic | d1 | Cxcr2 | ic |
| Cxcl1 | batch | mp | d1 | Cxcr2 | ic |
| Cxcl1 | batch | per | d1 | Cxcr2 | ic |
| Cxcl2 | batch | ec | d1 | Cxcr2 | ic |
| Cxcl2 | batch | fap | d1 | Cxcr2 | ic |
| Cxcl2 | batch | ic | d1 | Cxcr2 | ic |
| Cxcl2 | batch | per | d1 | Cxcr2 | ic |
| Cxcl3 | batch | fap | d1 | Cxcr2 | ic |
| Cxcl3 | batch | ic | d1 | Cxcr2 | ic |
| Cxcl3 | batch | mp | d1 | Cxcr2 | ic |
| Cxcl5 | batch | ec | d1 | Cxcr2 | ic |
| Cxcl5 | batch | fap | d1 | Cxcr2 | ic |
| Il6 | batch | ec | d1 | Il6ra | ic |
| Il6 | batch | fap | d1 | Il6ra | ic |
| Il6 | batch | per | d1 | Il6ra | ic |
| Mif | batch | ec | d1 | Tnfrsf14 | ic |
| Mif | batch | fap | d1 | Tnfrsf14 | ic |
| Pf4 | batch | fap | d1 | Ldlr | ic |
| Pf4 | batch | ic | d1 | Ldlr | ic |
| Pf4 | batch | per | d1 | Ldlr | ic |
| Wnt2 | batch | fap | d1 | Sfrp1 | ic |
| Il1b | batch | ec | d1 | Il1r2 | ic |
| Il1b | batch | ic | d1 | Il1r2 | ic |
| Il1b | batch | per | d1 | Il1r2 | ic |
| Pgf | batch | ec | d1 | Nrp2 | ic |
| Pgf | constitutive | per | d1 | Nrp2 | ic |
| Pgf | batch | ec | d1 | Nrp1 | ic |
| Pgf | constitutive | per | d1 | Nrp1 | ic |
| Ccl11 | batch | mp | d1 | Ccr5 | ic |
| Ccl11 | batch | per | d1 | Ccr5 | ic |
| Clcf1 | batch | mp | d1 | Crlf1 | ic |
| Clcf1 | batch | mp | d1 | Ncoa3 | ic |
| Ctgf | batch | ic | d1 | Itga5 | ic |
| Ctgf | batch | mp | d1 | Itga5 | ic |
| Ctgf | constitutive | per | d1 | Itga5 | ic |
| Esm1 | batch | mp | d1 | Itgal | ic |
| Hbegf | batch | mp | d1 | Cd44 | ic |
| Hbegf | constitutive | ec | d1 | Cd44 | ic |
| Hbegf | constitutive | per | d1 | Cd44 | ic |
| Jag2 | batch | mp | d1 | Notch2 | ic |
| Jag2 | constitutive | ec | d1 | Notch2 | ic |
| Pdap1 | batch | mp | d1 | Pdgfa | ic |
| Rabep1 | batch | mp | d1 | Lsr | ic |
| Spp1 | batch | mp | d1 | Itga5 | ic |
| Spp1 | batch | per | d1 | Itga5 | ic |
| Vegfa | batch | ic | d1 | Nrp2 | ic |
| Vegfa | batch | mp | d1 | Nrp2 | ic |
| Vegfa | batch | ic | d1 | Nrp1 | ic |
| Vegfa | batch | mp | d1 | Nrp1 | ic |
| Vegfc | batch | mp | d1 | Nrp2 | ic |
| Vegfc | batch | mp | d1 | Nrp1 | ic |
| Ccl3 | batch | ic | d1 | Ccr1 | ic |
| Ccl3 | batch | per | d1 | Ccr1 | ic |
| Ccl3 | batch | ic | d1 | Ccr5 | ic |
| Ccl3 | batch | per | d1 | Ccr5 | ic |
| Ccl4 | batch | ic | d1 | Ccr1 | ic |
| Ccl4 | batch | per | d1 | Ccr1 | ic |
| Ccl4 | batch | ic | d1 | Ccr5 | ic |
| Ccl4 | batch | per | d1 | Ccr5 | ic |
| Ccl6 | batch | ic | d1 | Ccr1 | ic |
| Ccl6 | batch | per | d1 | Ccr1 | ic |
| Il1a | batch | ic | d1 | Il1r2 | ic |
| Jag1 | batch | ic | d1 | Notch2 | ic |
| Jag1 | constitutive | per | d1 | Notch2 | ic |
| Ppbp | batch | ic | d1 | Cxcr2 | ic |
| Ppbp | batch | ic | d1 | Ncoa3 | ic |
| Tnfsf14 | batch | ic | d1 | Tnfrsf14 | ic |
| Nov | batch | per | d1 | Itga5 | ic |
| Fstl1 | constitutive | fap | d1 | Cd14 | ic |
| Fgf7 | constitutive | fap | d1 | Nrp1 | ic |
| Tgfb2 | constitutive | ec | d1 | Tgfbr3 | ic |
| Tgfb2 | constitutive | fap | d1 | Tgfbr3 | ic |
| Tgfb2 | constitutive | per | d1 | Tgfbr3 | ic |
| Pdgfb | constitutive | ec | d1 | Pdgfrb | ic |
| Ccl2 | batch | ec | d2 | Ccr1 | ic |
| Ccl2 | batch | fap | d2 | Ccr1 | ic |
| Ccl2 | batch | ic | d2 | Ccr1 | ic |
| Ccl2 | batch | mp | d2 | Ccr1 | ic |
| Ccl2 | batch | per | d2 | Ccr1 | ic |
| Ccl2 | batch | ec | d2 | Ccr5 | ic |
| Ccl2 | batch | fap | d2 | Ccr5 | ic |
| Ccl2 | batch | ic | d2 | Ccr5 | ic |
| Ccl2 | batch | mp | d2 | Ccr5 | ic |
| Ccl2 | batch | per | d2 | Ccr5 | ic |
| Ccl2 | batch | ec | d2 | Ccr2 | ic |
| Ccl2 | batch | fap | d2 | Ccr2 | ic |
| Ccl2 | batch | ic | d2 | Ccr2 | ic |
| Ccl2 | batch | mp | d2 | Ccr2 | ic |
| Ccl2 | batch | per | d2 | Ccr2 | ic |
| Ccl3 | batch | ec | d2 | Ccr1 | ic |
| Ccl3 | batch | fap | d2 | Ccr1 | ic |
| Ccl3 | batch | ic | d2 | Ccr1 | ic |
| Ccl3 | batch | mp | d2 | Ccr1 | ic |
| Ccl3 | batch | per | d2 | Ccr1 | ic |
| Ccl3 | batch | ec | d2 | Ccr5 | ic |
| Ccl3 | batch | fap | d2 | Ccr5 | ic |
| Ccl3 | batch | ic | d2 | Ccr5 | ic |
| Ccl3 | batch | mp | d2 | Ccr5 | ic |
| Ccl3 | batch | per | d2 | Ccr5 | ic |
| Ccl4 | batch | fap | d2 | Ccr1 | ic |
| Ccl4 | batch | ic | d2 | Ccr1 | ic |
| Ccl4 | batch | mp | d2 | Ccr1 | ic |
| Ccl4 | batch | per | d2 | Ccr1 | ic |
| Ccl4 | batch | fap | d2 | Ccr5 | ic |
| Ccl4 | batch | ic | d2 | Ccr5 | ic |
| Ccl4 | batch | mp | d2 | Ccr5 | ic |
| Ccl4 | batch | per | d2 | Ccr5 | ic |
| Ccl5 | batch | ec | d2 | Ccr1 | ic |
| Ccl5 | batch | fap | d2 | Ccr1 | ic |
| Ccl5 | batch | ec | d2 | Ccr5 | ic |
| Ccl5 | batch | fap | d2 | Ccr5 | ic |
| Ccl5 | batch | ec | d2 | Sdc4 | ic |
| Ccl5 | batch | fap | d2 | Sdc4 | ic |
| Ccl6 | batch | fap | d2 | Ccr1 | ic |
| Ccl6 | batch | ic | d2 | Ccr1 | ic |
| Ccl6 | batch | mp | d2 | Ccr1 | ic |
| Ccl6 | batch | per | d2 | Ccr1 | ic |
| Ccl7 | batch | ec | d2 | Ccr1 | ic |
| Ccl7 | batch | fap | d2 | Ccr1 | ic |
| Ccl7 | batch | ic | d2 | Ccr1 | ic |
| Ccl7 | batch | mp | d2 | Ccr1 | ic |
| Ccl7 | batch | per | d2 | Ccr1 | ic |
| Ccl7 | batch | ec | d2 | Ccr5 | ic |
| Ccl7 | batch | fap | d2 | Ccr5 | ic |
| Ccl7 | batch | ic | d2 | Ccr5 | ic |
| Ccl7 | batch | mp | d2 | Ccr5 | ic |
| Ccl7 | batch | per | d2 | Ccr5 | ic |
| Csf1 | batch | fap | d2 | Csf1r | ic |
| Csf1 | batch | ic | d2 | Csf1r | ic |
| Csf1 | batch | mp | d2 | Csf1r | ic |
| Csf2 | batch | fap | d2 | Csf3r | ic |
| Csf2 | batch | fap | d2 | Il3ra | ic |
| Csf2 | batch | fap | d2 | Csf2rb | ic |
| Csf2 | batch | fap | d2 | Csf2ra | ic |
| Cxcl1 | batch | ec | d2 | Cxcr2 | ic |
| Cxcl1 | batch | fap | d2 | Cxcr2 | ic |
| Cxcl1 | batch | ic | d2 | Cxcr2 | ic |
| Cxcl1 | batch | mp | d2 | Cxcr2 | ic |
| Cxcl1 | batch | per | d2 | Cxcr2 | ic |
| Cxcl12 | batch | fap | d2 | Cxcr4 | ic |
| Cxcl12 | batch | mp | d2 | Cxcr4 | ic |
| Cxcl12 | constitutive | ec | d2 | Cxcr4 | ic |
| Cxcl12 | constitutive | per | d2 | Cxcr4 | ic |
| Cxcl2 | batch | ec | d2 | Cxcr2 | ic |
| Cxcl2 | batch | fap | d2 | Cxcr2 | ic |
| Cxcl2 | batch | ic | d2 | Cxcr2 | ic |
| Cxcl2 | batch | mp | d2 | Cxcr2 | ic |
| Cxcl2 | batch | per | d2 | Cxcr2 | ic |
| Cxcl3 | batch | fap | d2 | Cxcr2 | ic |
| Cxcl3 | batch | ic | d2 | Cxcr2 | ic |
| Cxcl3 | batch | mp | d2 | Cxcr2 | ic |
| Cxcl5 | batch | ec | d2 | Cxcr2 | ic |
| Cxcl5 | batch | fap | d2 | Cxcr2 | ic |
| Gdf10 | batch | fap | d2 | Tgfbr1 | ic |
| Il1b | batch | ec | d2 | Il1r2 | ic |
| Il1b | batch | fap | d2 | Il1r2 | ic |
| Il1b | batch | ic | d2 | Il1r2 | ic |
| Il1b | batch | mp | d2 | Il1r2 | ic |
| Il1b | batch | per | d2 | Il1r2 | ic |
| Il6 | batch | ec | d2 | Il6ra | ic |
| Il6 | batch | fap | d2 | Il6ra | ic |
| Il6 | batch | per | d2 | Il6ra | ic |
| Mif | batch | ec | d2 | Tnfrsf14 | ic |
| Mif | batch | fap | d2 | Tnfrsf14 | ic |
| Mif | batch | mp | d2 | Tnfrsf14 | ic |
| Pf4 | batch | ec | d2 | Ldlr | ic |
| Pf4 | batch | fap | d2 | Ldlr | ic |
| Pf4 | batch | ic | d2 | Ldlr | ic |
| Pf4 | batch | mp | d2 | Ldlr | ic |
| Pf4 | batch | per | d2 | Ldlr | ic |
| Spp1 | batch | ec | d2 | Itga5 | ic |
| Spp1 | batch | fap | d2 | Itga5 | ic |
| Spp1 | batch | ic | d2 | Itga5 | ic |
| Spp1 | batch | mp | d2 | Itga5 | ic |
| Spp1 | batch | per | d2 | Itga5 | ic |
| Spp1 | batch | ec | d2 | Itga9 | ic |
| Spp1 | batch | fap | d2 | Itga9 | ic |
| Spp1 | batch | ic | d2 | Itga9 | ic |
| Spp1 | batch | mp | d2 | Itga9 | ic |
| Spp1 | batch | per | d2 | Itga9 | ic |
| Spp1 | batch | ec | d2 | Itgav | ic |
| Spp1 | batch | fap | d2 | Itgav | ic |
| Spp1 | batch | ic | d2 | Itgav | ic |
| Spp1 | batch | mp | d2 | Itgav | ic |
| Spp1 | batch | per | d2 | Itgav | ic |
| Spp1 | batch | ec | d2 | Itgb5 | ic |
| Spp1 | batch | fap | d2 | Itgb5 | ic |
| Spp1 | batch | ic | d2 | Itgb5 | ic |
| Spp1 | batch | mp | d2 | Itgb5 | ic |
| Spp1 | batch | per | d2 | Itgb5 | ic |
| Tslp | batch | fap | d2 | Il7r | ic |
| Tslp | constitutive | ec | d2 | Il7r | ic |
| Tslp | batch | fap | d2 | Crlf2 | ic |
| Tslp | constitutive | ec | d2 | Crlf2 | ic |
| Wnt2 | batch | fap | d2 | Sfrp1 | ic |
| Il15 | batch | ec | d2 | Il17ra | ic |
| Il15 | batch | ic | d2 | Il17ra | ic |
| Il15 | batch | ec | d2 | Il2rg | ic |
| Il15 | batch | ic | d2 | Il2rg | ic |
| Lgals3 | batch | ec | d2 | Lgals3bp | ic |
| Lgals3 | batch | ic | d2 | Lgals3bp | ic |
| Lgals3 | batch | mp | d2 | Lgals3bp | ic |
| Lgals3 | batch | per | d2 | Lgals3bp | ic |
| Mmp12 | batch | ec | d2 | Plaur | ic |
| Mmp12 | batch | ic | d2 | Plaur | ic |
| Mmp12 | batch | mp | d2 | Plaur | ic |
| Mmp12 | batch | per | d2 | Plaur | ic |
| Pgf | batch | ec | d2 | Nrp2 | ic |
| Pgf | constitutive | per | d2 | Nrp2 | ic |
| Pgf | batch | ec | d2 | Nrp1 | ic |
| Pgf | constitutive | per | d2 | Nrp1 | ic |
| C4b | batch | mp | d2 | C3ar1 | ic |
| C4b | constitutive | per | d2 | C3ar1 | ic |
| Ccl11 | batch | mp | d2 | Ccr5 | ic |
| Ccl11 | batch | per | d2 | Ccr5 | ic |
| Ccl8 | batch | mp | d2 | Ccr1 | ic |
| Ccl8 | total | fap | d2 | Ccr1 | ic |
| Ccl8 | total | ic | d2 | Ccr1 | ic |
| Ccl8 | batch | mp | d2 | Ccr5 | ic |
| Ccl8 | total | fap | d2 | Ccr5 | ic |
| Ccl8 | total | ic | d2 | Ccr5 | ic |
| Ccl8 | batch | mp | d2 | Ccr2 | ic |
| Ccl8 | total | fap | d2 | Ccr2 | ic |
| Ccl8 | total | ic | d2 | Ccr2 | ic |
| Clcf1 | batch | mp | d2 | Crlf1 | ic |
| Clcf1 | batch | mp | d2 | Ncoa3 | ic |
| Ctgf | batch | mp | d2 | Itga5 | ic |
| Ctgf | constitutive | per | d2 | Itga5 | ic |
| Cx3cl1 | batch | mp | d2 | Cx3cr1 | ic |
| Esm1 | batch | mp | d2 | Itgal | ic |
| Gas6 | batch | mp | d2 | Axl | ic |
| Hbegf | batch | mp | d2 | Cd44 | ic |
| Hbegf | constitutive | ec | d2 | Cd44 | ic |
| Hbegf | constitutive | per | d2 | Cd44 | ic |
| Il10 | batch | ic | d2 | Ifnar2 | ic |
| Il10 | batch | mp | d2 | Ifnar2 | ic |
| Il10 | batch | ic | d2 | Il10rb | ic |
| Il10 | batch | mp | d2 | Il10rb | ic |
| Il10 | batch | ic | d2 | Il10ra | ic |
| Il10 | batch | mp | d2 | Il10ra | ic |
| Jag2 | batch | mp | d2 | Notch2 | ic |
| Jag2 | constitutive | ec | d2 | Notch2 | ic |
| Ngf | batch | mp | d2 | Sort1 | ic |
| Ngf | constitutive | per | d2 | Sort1 | ic |
| Pdap1 | batch | mp | d2 | Pdgfa | ic |
| Rabep1 | batch | mp | d2 | Lsr | ic |
| Tgfb1 | batch | mp | d2 | Itgav | ic |
| Tgfb1 | total | ec | d2 | Itgav | ic |
| Tgfb1 | total | ic | d2 | Itgav | ic |
| Tgfb1 | batch | mp | d2 | Tgfbr1 | ic |
| Tgfb1 | total | ec | d2 | Tgfbr1 | ic |
| Tgfb1 | total | ic | d2 | Tgfbr1 | ic |
| Tgfb1 | batch | mp | d2 | Acvrl1 | ic |
| Tgfb1 | total | ec | d2 | Acvrl1 | ic |
| Tgfb1 | total | ic | d2 | Acvrl1 | ic |
| Tnf | batch | ic | d2 | Tnfrsf1b | ic |
| Tnf | batch | mp | d2 | Tnfrsf1b | ic |
| Tnf | batch | per | d2 | Tnfrsf1b | ic |
| Vegfa | batch | ic | d2 | Itga9 | ic |
| Vegfa | batch | mp | d2 | Itga9 | ic |
| Vegfa | batch | ic | d2 | Nrp2 | ic |
| Vegfa | batch | mp | d2 | Nrp2 | ic |
| Vegfa | batch | ic | d2 | Nrp1 | ic |
| Vegfa | batch | mp | d2 | Nrp1 | ic |
| Vegfc | batch | mp | d2 | Itga9 | ic |
| Vegfc | batch | mp | d2 | Nrp2 | ic |
| Vegfc | batch | mp | d2 | Nrp1 | ic |
| Ifnb1 | batch | ic | d2 | Ifnar2 | ic |
| Il1a | batch | ic | d2 | Il1r2 | ic |
| Jag1 | batch | ic | d2 | Notch2 | ic |
| Jag1 | constitutive | per | d2 | Notch2 | ic |
| Ppbp | batch | ic | d2 | Cxcr2 | ic |
| Ppbp | batch | ic | d2 | Itgb5 | ic |
| Ppbp | batch | ic | d2 | Il7r | ic |
| Ppbp | batch | ic | d2 | Ncoa3 | ic |
| Tnfsf13 | batch | ic | d2 | Tnfrsf11b | ic |
| Tnfsf13 | batch | ic | d2 | Tnfrsf14 | ic |
| Tnfsf14 | batch | ic | d2 | Tnfrsf14 | ic |
| Nov | batch | per | d2 | Itga5 | ic |
| Nov | batch | per | d2 | Itgav | ic |
| Ucn2 | batch | per | d2 | Il10rb | ic |
| Tnfsf12 | total | ec | d2 | Tnfrsf11b | ic |
| Tnfsf12 | total | ic | d2 | Tnfrsf11b | ic |
| Tnfsf12 | total | per | d2 | Tnfrsf11b | ic |
| Tnfsf12 | total | ec | d2 | Tnfrsf14 | ic |
| Tnfsf12 | total | ic | d2 | Tnfrsf14 | ic |
| Tnfsf12 | total | per | d2 | Tnfrsf14 | ic |
| Fstl1 | constitutive | fap | d2 | Cd14 | ic |
| Fgf7 | constitutive | fap | d2 | Nrp1 | ic |
| Tgfb2 | constitutive | ec | d2 | Tgfbr1 | ic |
| Tgfb2 | constitutive | fap | d2 | Tgfbr1 | ic |
| Tgfb2 | constitutive | per | d2 | Tgfbr1 | ic |
| Pdgfb | constitutive | ec | d2 | Lrp1 | ic |
| Ptn | constitutive | per | d2 | Plxnb2 | ic |
| C4b | batch | fap | d3 | C3ar1 | ic |
| C4b | constitutive | per | d3 | C3ar1 | ic |
| Ccl3 | batch | ec | d3 | Ccr5 | ic |
| Ccl3 | batch | fap | d3 | Ccr5 | ic |
| Ccl3 | batch | mp | d3 | Ccr5 | ic |
| Ccl3 | batch | per | d3 | Ccr5 | ic |
| Ccl4 | batch | fap | d3 | Ccr5 | ic |
| Ccl4 | batch | mp | d3 | Ccr5 | ic |
| Ccl4 | batch | per | d3 | Ccr5 | ic |
| Ccl5 | batch | ec | d3 | Ccr5 | ic |
| Ccl5 | batch | fap | d3 | Ccr5 | ic |
| Csf2 | batch | fap | d3 | Il3ra | ic |
| Csf2 | batch | fap | d3 | Csf2rb | ic |
| Ctgf | batch | fap | d3 | Itga5 | ic |
| Ctgf | constitutive | per | d3 | Itga5 | ic |
| Cx3cl1 | batch | fap | d3 | Cx3cr1 | ic |
| Cx3cl1 | total | ec | d3 | Cx3cr1 | ic |
| Cx3cl1 | total | per | d3 | Cx3cr1 | ic |
| Cxcl12 | batch | fap | d3 | Cxcr4 | ic |
| Cxcl12 | constitutive | ec | d3 | Cxcr4 | ic |
| Cxcl12 | constitutive | per | d3 | Cxcr4 | ic |
| Gas6 | batch | fap | d3 | Axl | ic |
| Gdf10 | batch | fap | d3 | Tgfbr1 | ic |
| Mif | batch | fap | d3 | Tnfrsf14 | ic |
| Mif | batch | mp | d3 | Tnfrsf14 | ic |
| Ptn | batch | fap | d3 | Plxnb2 | ic |
| Ptn | constitutive | per | d3 | Plxnb2 | ic |
| Spp1 | batch | ec | d3 | Itga5 | ic |
| Spp1 | batch | fap | d3 | Itga5 | ic |
| Spp1 | batch | ic | d3 | Itga5 | ic |
| Spp1 | batch | mp | d3 | Itga5 | ic |
| Spp1 | batch | per | d3 | Itga5 | ic |
| Spp1 | batch | ec | d3 | Itga9 | ic |
| Spp1 | batch | fap | d3 | Itga9 | ic |
| Spp1 | batch | ic | d3 | Itga9 | ic |
| Spp1 | batch | mp | d3 | Itga9 | ic |
| Spp1 | batch | per | d3 | Itga9 | ic |
| Spp1 | batch | ec | d3 | Itgav | ic |
| Spp1 | batch | fap | d3 | Itgav | ic |
| Spp1 | batch | ic | d3 | Itgav | ic |
| Spp1 | batch | mp | d3 | Itgav | ic |
| Spp1 | batch | per | d3 | Itgav | ic |
| Spp1 | batch | ec | d3 | Itgb5 | ic |
| Spp1 | batch | fap | d3 | Itgb5 | ic |
| Spp1 | batch | ic | d3 | Itgb5 | ic |
| Spp1 | batch | mp | d3 | Itgb5 | ic |
| Spp1 | batch | per | d3 | Itgb5 | ic |
| Tgfb3 | batch | fap | d3 | Itgav | ic |
| Tgfb3 | total | ec | d3 | Itgav | ic |
| Tgfb3 | total | per | d3 | Itgav | ic |
| Tgfb3 | batch | fap | d3 | Tgfbr1 | ic |
| Tgfb3 | total | ec | d3 | Tgfbr1 | ic |
| Tgfb3 | total | per | d3 | Tgfbr1 | ic |
| Tgfb3 | batch | fap | d3 | Acvrl1 | ic |
| Tgfb3 | total | ec | d3 | Acvrl1 | ic |
| Tgfb3 | total | per | d3 | Acvrl1 | ic |
| Tslp | batch | fap | d3 | Il7r | ic |
| Tslp | constitutive | ec | d3 | Il7r | ic |
| Wnt2 | batch | fap | d3 | Sfrp1 | ic |
| Il15 | batch | ec | d3 | Il17ra | ic |
| Il15 | batch | ic | d3 | Il17ra | ic |
| Il15 | batch | ec | d3 | Il2rg | ic |
| Il15 | batch | ic | d3 | Il2rg | ic |
| Il6 | batch | ec | d3 | Il6ra | ic |
| Il6 | batch | per | d3 | Il6ra | ic |
| Lgals3 | batch | ec | d3 | Lgals3bp | ic |
| Lgals3 | batch | ic | d3 | Lgals3bp | ic |
| Lgals3 | batch | mp | d3 | Lgals3bp | ic |
| Lgals3 | batch | per | d3 | Lgals3bp | ic |
| Vegfc | batch | ec | d3 | Itga9 | ic |
| Vegfc | batch | ec | d3 | Nrp2 | ic |
| Vegfc | batch | ec | d3 | Nrp1 | ic |
| Ccl2 | batch | mp | d3 | Ccr5 | ic |
| Ccl2 | batch | per | d3 | Ccr5 | ic |
| Ccl2 | batch | mp | d3 | Ccr2 | ic |
| Ccl2 | batch | per | d3 | Ccr2 | ic |
| Ccl7 | batch | mp | d3 | Ccr5 | ic |
| Ccl7 | batch | per | d3 | Ccr5 | ic |
| Ccl8 | batch | mp | d3 | Ccr5 | ic |
| Ccl8 | total | fap | d3 | Ccr5 | ic |
| Ccl8 | total | ic | d3 | Ccr5 | ic |
| Ccl8 | batch | mp | d3 | Ccr2 | ic |
| Ccl8 | total | fap | d3 | Ccr2 | ic |
| Ccl8 | total | ic | d3 | Ccr2 | ic |
| Csf1 | batch | ic | d3 | Csf1r | ic |
| Csf1 | batch | mp | d3 | Csf1r | ic |
| Il10 | batch | ic | d3 | Ifnar2 | ic |
| Il10 | batch | mp | d3 | Ifnar2 | ic |
| Il10 | batch | ic | d3 | Il10ra | ic |
| Il10 | batch | mp | d3 | Il10ra | ic |
| Pdap1 | batch | mp | d3 | Pdgfa | ic |
| Tgfb1 | batch | mp | d3 | Itgav | ic |
| Tgfb1 | total | ec | d3 | Itgav | ic |
| Tgfb1 | total | ic | d3 | Itgav | ic |
| Tgfb1 | batch | mp | d3 | Tgfbr1 | ic |
| Tgfb1 | total | ec | d3 | Tgfbr1 | ic |
| Tgfb1 | total | ic | d3 | Tgfbr1 | ic |
| Tgfb1 | batch | mp | d3 | Acvrl1 | ic |
| Tgfb1 | total | ec | d3 | Acvrl1 | ic |
| Tgfb1 | total | ic | d3 | Acvrl1 | ic |
| Tnf | batch | mp | d3 | Tnfrsf1b | ic |
| Tnf | batch | per | d3 | Tnfrsf1b | ic |
| Tnfsf13 | batch | ic | d3 | Tnfrsf11b | ic |
| Tnfsf13 | batch | ic | d3 | Tnfrsf14 | ic |
| Dll1 | total | mp | d3 | Notch2 | ic |
| Fgf7 | constitutive | fap | d3 | Nrp1 | ic |
| Tgfb2 | constitutive | ec | d3 | Tgfbr1 | ic |
| Tgfb2 | constitutive | fap | d3 | Tgfbr1 | ic |
| Tgfb2 | constitutive | per | d3 | Tgfbr1 | ic |
| Jag2 | constitutive | ec | d3 | Notch2 | ic |
| Ngf | constitutive | per | d3 | Sort1 | ic |
| Pgf | constitutive | per | d3 | Nrp2 | ic |
| Pgf | constitutive | per | d3 | Nrp1 | ic |
| Jag1 | constitutive | per | d3 | Notch2 | ic |
| C4b | batch | fap | d4 | C3ar1 | ic |
| C4b | batch | ic | d4 | C3ar1 | ic |
| C4b | constitutive | per | d4 | C3ar1 | ic |
| Ccl3 | batch | ec | d4 | Ccr5 | ic |
| Ccl3 | batch | fap | d4 | Ccr5 | ic |
| Ccl3 | batch | mp | d4 | Ccr5 | ic |
| Ccl4 | batch | fap | d4 | Ccr5 | ic |
| Ccl4 | batch | mp | d4 | Ccr5 | ic |
| Ccl5 | batch | ec | d4 | Ccr5 | ic |
| Ccl5 | batch | fap | d4 | Ccr5 | ic |
| Ccl5 | batch | ec | d4 | Cxcr3 | ic |
| Ccl5 | batch | fap | d4 | Cxcr3 | ic |
| Ctgf | batch | fap | d4 | Itga5 | ic |
| Ctgf | constitutive | per | d4 | Itga5 | ic |
| Cx3cl1 | batch | fap | d4 | Cx3cr1 | ic |
| Cx3cl1 | batch | mp | d4 | Cx3cr1 | ic |
| Cx3cl1 | total | ec | d4 | Cx3cr1 | ic |
| Cx3cl1 | total | per | d4 | Cx3cr1 | ic |
| Cxcl10 | batch | ec | d4 | Cxcr3 | ic |
| Cxcl10 | batch | fap | d4 | Cxcr3 | ic |
| Cxcl10 | batch | ic | d4 | Cxcr3 | ic |
| Cxcl9 | batch | ec | d4 | Cxcr3 | ic |
| Cxcl9 | batch | fap | d4 | Cxcr3 | ic |
| Cxcl9 | batch | ic | d4 | Cxcr3 | ic |
| Gas6 | batch | fap | d4 | Axl | ic |
| Il16 | batch | fap | d4 | Cd4 | ic |
| Il16 | batch | ic | d4 | Cd4 | ic |
| Mif | batch | fap | d4 | Cd74 | ic |
| Mif | batch | fap | d4 | Tnfrsf14 | ic |
| Pf4 | batch | ec | d4 | Cxcr3 | ic |
| Pf4 | batch | fap | d4 | Cxcr3 | ic |
| Ptn | batch | fap | d4 | Ptprs | ic |
| Ptn | constitutive | per | d4 | Ptprs | ic |
| Spp1 | batch | ec | d4 | Itga5 | ic |
| Spp1 | batch | fap | d4 | Itga5 | ic |
| Spp1 | batch | ic | d4 | Itga5 | ic |
| Spp1 | batch | mp | d4 | Itga5 | ic |
| Spp1 | batch | ec | d4 | Itga9 | ic |
| Spp1 | batch | fap | d4 | Itga9 | ic |
| Spp1 | batch | ic | d4 | Itga9 | ic |
| Spp1 | batch | mp | d4 | Itga9 | ic |
| Spp1 | batch | ec | d4 | Itgav | ic |
| Spp1 | batch | fap | d4 | Itgav | ic |
| Spp1 | batch | ic | d4 | Itgav | ic |
| Spp1 | batch | mp | d4 | Itgav | ic |
| Spp1 | batch | ec | d4 | Itgb5 | ic |
| Spp1 | batch | fap | d4 | Itgb5 | ic |
| Spp1 | batch | ic | d4 | Itgb5 | ic |
| Spp1 | batch | mp | d4 | Itgb5 | ic |
| Tgfb3 | batch | fap | d4 | Itgav | ic |
| Tgfb3 | total | ec | d4 | Itgav | ic |
| Tgfb3 | total | per | d4 | Itgav | ic |
| Tgfb3 | batch | fap | d4 | Tgfbr1 | ic |
| Tgfb3 | total | ec | d4 | Tgfbr1 | ic |
| Tgfb3 | total | per | d4 | Tgfbr1 | ic |
| Tgfb3 | batch | fap | d4 | Acvrl1 | ic |
| Tgfb3 | total | ec | d4 | Acvrl1 | ic |
| Tgfb3 | total | per | d4 | Acvrl1 | ic |
| Dll1 | batch | ec | d4 | Notch2 | ic |
| Dll1 | total | mp | d4 | Notch2 | ic |
| Lgals3 | batch | ec | d4 | Lgals3bp | ic |
| Lgals3 | batch | ic | d4 | Lgals3bp | ic |
| Lgals3 | batch | mp | d4 | Lgals3bp | ic |
| Vegfc | batch | ec | d4 | Itga9 | ic |
| Vegfc | batch | mp | d4 | Itga9 | ic |
| Vegfc | batch | ec | d4 | Nrp2 | ic |
| Vegfc | batch | mp | d4 | Nrp2 | ic |
| Vegfc | batch | ec | d4 | Nrp1 | ic |
| Vegfc | batch | mp | d4 | Nrp1 | ic |
| Ccl8 | batch | mp | d4 | Ccr5 | ic |
| Ccl8 | total | fap | d4 | Ccr5 | ic |
| Ccl8 | total | ic | d4 | Ccr5 | ic |
| Esm1 | batch | mp | d4 | Itgal | ic |
| Il10 | batch | ic | d4 | Il10ra | ic |
| Il10 | batch | mp | d4 | Il10ra | ic |
| Pdap1 | batch | mp | d4 | Pdgfa | ic |
| Rabep1 | batch | mp | d4 | Lsr | ic |
| Tgfb1 | batch | mp | d4 | Itgav | ic |
| Tgfb1 | batch | mp | d4 | Tgfbr1 | ic |
| Tgfb1 | batch | mp | d4 | Acvrl1 | ic |
| Tgfb2 | batch | mp | d4 | Tgfbr1 | ic |
| Tgfb2 | constitutive | ec | d4 | Tgfbr1 | ic |
| Tgfb2 | constitutive | fap | d4 | Tgfbr1 | ic |
| Tgfb2 | constitutive | per | d4 | Tgfbr1 | ic |
| Tnfsf13 | batch | ic | d4 | Tnfrsf11b | ic |
| Tnfsf13 | batch | ic | d4 | Tnfrsf14 | ic |
| Fgf7 | constitutive | fap | d4 | Nrp1 | ic |
| Jag2 | constitutive | ec | d4 | Notch2 | ic |
| Ngf | constitutive | per | d4 | Sort1 | ic |
| Pgf | constitutive | per | d4 | Nrp2 | ic |
| Pgf | constitutive | per | d4 | Nrp1 | ic |
| Jag1 | constitutive | per | d4 | Notch2 | ic |
| Apln | batch | ec | d5 | Aplnr | ic |
| Apln | batch | fap | d5 | Aplnr | ic |
| C4b | batch | fap | d5 | C3ar1 | ic |
| C4b | batch | ic | d5 | C3ar1 | ic |
| C4b | constitutive | per | d5 | C3ar1 | ic |
| Ccl3 | batch | fap | d5 | Ccr5 | ic |
| Ccl4 | batch | fap | d5 | Ccr5 | ic |
| Ccl5 | batch | fap | d5 | Ccr5 | ic |
| Ccl5 | batch | fap | d5 | Cxcr3 | ic |
| Ctgf | batch | fap | d5 | Itga5 | ic |
| Ctgf | batch | ic | d5 | Itga5 | ic |
| Ctgf | constitutive | per | d5 | Itga5 | ic |
| Cx3cl1 | batch | fap | d5 | Cx3cr1 | ic |
| Cx3cl1 | batch | mp | d5 | Cx3cr1 | ic |
| Cx3cl1 | total | ec | d5 | Cx3cr1 | ic |
| Cx3cl1 | total | per | d5 | Cx3cr1 | ic |
| Cxcl10 | batch | ec | d5 | Cxcr3 | ic |
| Cxcl10 | batch | fap | d5 | Cxcr3 | ic |
| Cxcl10 | batch | ic | d5 | Cxcr3 | ic |
| Cxcl10 | batch | per | d5 | Cxcr3 | ic |
| Cxcl9 | batch | ec | d5 | Cxcr3 | ic |
| Cxcl9 | batch | fap | d5 | Cxcr3 | ic |
| Cxcl9 | batch | ic | d5 | Cxcr3 | ic |
| Gas6 | batch | fap | d5 | Axl | ic |
| Hgf | batch | fap | d5 | Vtn | ic |
| Hgf | batch | ic | d5 | Vtn | ic |
| Igf1 | batch | ec | d5 | Igfbp3 | ic |
| Igf1 | batch | fap | d5 | Igfbp3 | ic |
| Igf1 | batch | ic | d5 | Igfbp3 | ic |
| Igf1 | batch | per | d5 | Igfbp3 | ic |
| Igf1 | batch | ec | d5 | Igfbp4 | ic |
| Igf1 | batch | fap | d5 | Igfbp4 | ic |
| Igf1 | batch | ic | d5 | Igfbp4 | ic |
| Igf1 | batch | per | d5 | Igfbp4 | ic |
| Igf1 | batch | ec | d5 | Igfbp5 | ic |
| Igf1 | batch | fap | d5 | Igfbp5 | ic |
| Igf1 | batch | ic | d5 | Igfbp5 | ic |
| Igf1 | batch | per | d5 | Igfbp5 | ic |
| Igf1 | batch | ec | d5 | Igfbp6 | ic |
| Igf1 | batch | fap | d5 | Igfbp6 | ic |
| Igf1 | batch | ic | d5 | Igfbp6 | ic |
| Igf1 | batch | per | d5 | Igfbp6 | ic |
| Igf1 | batch | ec | d5 | Igfbp7 | ic |
| Igf1 | batch | fap | d5 | Igfbp7 | ic |
| Igf1 | batch | ic | d5 | Igfbp7 | ic |
| Igf1 | batch | per | d5 | Igfbp7 | ic |
| Igf2 | batch | ec | d5 | Vtn | ic |
| Igf2 | batch | fap | d5 | Vtn | ic |
| Igf2 | batch | ic | d5 | Vtn | ic |
| Igf2 | batch | mp | d5 | Vtn | ic |
| Il16 | batch | fap | d5 | Cd4 | ic |
| Il16 | batch | ic | d5 | Cd4 | ic |
| Mif | batch | fap | d5 | Cd74 | ic |
| Mif | batch | fap | d5 | Tnfrsf14 | ic |
| Pf4 | batch | fap | d5 | Cxcr3 | ic |
| Pthlh | batch | fap | d5 | Pth1r | ic |
| Ptn | batch | fap | d5 | Ptprs | ic |
| Ptn | constitutive | per | d5 | Ptprs | ic |
| Spp1 | batch | fap | d5 | Itga5 | ic |
| Spp1 | batch | ic | d5 | Itga5 | ic |
| Spp1 | batch | mp | d5 | Itga5 | ic |
| Spp1 | batch | fap | d5 | Itga9 | ic |
| Spp1 | batch | ic | d5 | Itga9 | ic |
| Spp1 | batch | mp | d5 | Itga9 | ic |
| Spp1 | batch | fap | d5 | Itgav | ic |
| Spp1 | batch | ic | d5 | Itgav | ic |
| Spp1 | batch | mp | d5 | Itgav | ic |
| Spp1 | batch | fap | d5 | Itgb5 | ic |
| Spp1 | batch | ic | d5 | Itgb5 | ic |
| Spp1 | batch | mp | d5 | Itgb5 | ic |
| Spp1 | batch | fap | d5 | Vtn | ic |
| Spp1 | batch | ic | d5 | Vtn | ic |
| Spp1 | batch | mp | d5 | Vtn | ic |
| Tgfb3 | batch | fap | d5 | Itgav | ic |
| Tgfb3 | total | ec | d5 | Itgav | ic |
| Tgfb3 | total | per | d5 | Itgav | ic |
| Tgfb3 | batch | fap | d5 | Tgfbr1 | ic |
| Tgfb3 | total | ec | d5 | Tgfbr1 | ic |
| Tgfb3 | total | per | d5 | Tgfbr1 | ic |
| Tgfb3 | batch | fap | d5 | Acvrl1 | ic |
| Tgfb3 | total | ec | d5 | Acvrl1 | ic |
| Tgfb3 | total | per | d5 | Acvrl1 | ic |
| Tnfrsf11b | batch | fap | d5 | Vtn | ic |
| Dll1 | batch | ec | d5 | Notch2 | ic |
| Dll1 | total | mp | d5 | Notch2 | ic |
| Il6 | batch | ec | d5 | Il6ra | ic |
| Il6 | batch | per | d5 | Il6ra | ic |
| Vegfc | batch | ec | d5 | Itga9 | ic |
| Vegfc | batch | mp | d5 | Itga9 | ic |
| Vegfc | batch | ec | d5 | Nrp2 | ic |
| Vegfc | batch | mp | d5 | Nrp2 | ic |
| Vegfc | batch | ec | d5 | Nrp1 | ic |
| Vegfc | batch | mp | d5 | Nrp1 | ic |
| Esm1 | batch | mp | d5 | Itgal | ic |
| Pdap1 | batch | mp | d5 | Pdgfa | ic |
| Rabep1 | batch | mp | d5 | Lsr | ic |
| Tgfb2 | batch | mp | d5 | Tgfbr1 | ic |
| Tgfb2 | constitutive | ec | d5 | Tgfbr1 | ic |
| Tgfb2 | constitutive | fap | d5 | Tgfbr1 | ic |
| Tgfb2 | constitutive | per | d5 | Tgfbr1 | ic |
| Tgfb2 | batch | mp | d5 | Tgfbr3 | ic |
| Tgfb2 | constitutive | ec | d5 | Tgfbr3 | ic |
| Tgfb2 | constitutive | fap | d5 | Tgfbr3 | ic |
| Tgfb2 | constitutive | per | d5 | Tgfbr3 | ic |
| Tgfb2 | batch | mp | d5 | Vtn | ic |
| Tgfb2 | constitutive | ec | d5 | Vtn | ic |
| Tgfb2 | constitutive | fap | d5 | Vtn | ic |
| Tgfb2 | constitutive | per | d5 | Vtn | ic |
| Il10 | batch | ic | d5 | Il10ra | ic |
| Lgals3 | batch | ic | d5 | Lgals3bp | ic |
| Pdgfb | batch | ic | d5 | Pdgfrb | ic |
| Pdgfb | constitutive | ec | d5 | Pdgfrb | ic |
| Tnfsf13 | batch | ic | d5 | Tnfrsf11b | ic |
| Tnfsf13 | batch | ic | d5 | Tnfrsf14 | ic |
| Ccl11 | batch | per | d5 | Ccr5 | ic |
| Ccl11 | batch | per | d5 | Cxcr3 | ic |
| Ccl2 | batch | per | d5 | Ccr5 | ic |
| Ccl7 | batch | per | d5 | Ccr5 | ic |
| Ccl7 | batch | per | d5 | Cxcr3 | ic |
| Gdf10 | batch | per | d5 | Tgfbr1 | ic |
| Nov | batch | per | d5 | Itga5 | ic |
| Nov | batch | per | d5 | Itgav | ic |
| Ccl8 | total | fap | d5 | Ccr5 | ic |
| Ccl8 | total | ic | d5 | Ccr5 | ic |
| Fgf7 | constitutive | fap | d5 | Nrp1 | ic |
| Jag2 | constitutive | ec | d5 | Notch2 | ic |
| Ngf | constitutive | per | d5 | Sort1 | ic |
| Pgf | constitutive | per | d5 | Nrp2 | ic |
| Pgf | constitutive | per | d5 | Nrp1 | ic |
| Jag1 | constitutive | per | d5 | Notch2 | ic |
| C4b | batch | fap | d6 | C3ar1 | ic |
| C4b | batch | ic | d6 | C3ar1 | ic |
| C4b | batch | mp | d6 | C3ar1 | ic |
| C4b | constitutive | per | d6 | C3ar1 | ic |
| Ccl3 | batch | fap | d6 | Ccr5 | ic |
| Ccl3 | batch | mp | d6 | Ccr5 | ic |
| Ccl4 | batch | fap | d6 | Ccr5 | ic |
| Ccl4 | batch | mp | d6 | Ccr5 | ic |
| Ccl5 | batch | fap | d6 | Ccr5 | ic |
| Ccl5 | batch | fap | d6 | Cxcr3 | ic |
| Crlf1 | batch | fap | d6 | Ctf1 | ic |
| Crlf1 | batch | mp | d6 | Ctf1 | ic |
| Ctgf | batch | fap | d6 | Itga5 | ic |
| Ctgf | batch | ic | d6 | Itga5 | ic |
| Ctgf | batch | mp | d6 | Itga5 | ic |
| Ctgf | constitutive | per | d6 | Itga5 | ic |
| Cx3cl1 | batch | fap | d6 | Cx3cr1 | ic |
| Cx3cl1 | batch | mp | d6 | Cx3cr1 | ic |
| Cx3cl1 | total | ec | d6 | Cx3cr1 | ic |
| Cx3cl1 | total | per | d6 | Cx3cr1 | ic |
| Cxcl10 | batch | ec | d6 | Cxcr3 | ic |
| Cxcl10 | batch | fap | d6 | Cxcr3 | ic |
| Cxcl10 | batch | ic | d6 | Cxcr3 | ic |
| Cxcl9 | batch | ec | d6 | Cxcr3 | ic |
| Cxcl9 | batch | fap | d6 | Cxcr3 | ic |
| Cxcl9 | batch | ic | d6 | Cxcr3 | ic |
| Gas6 | batch | fap | d6 | Axl | ic |
| Gas6 | batch | mp | d6 | Axl | ic |
| Igf1 | batch | ec | d6 | Igfbp3 | ic |
| Igf1 | batch | fap | d6 | Igfbp3 | ic |
| Igf1 | batch | ic | d6 | Igfbp3 | ic |
| Igf1 | batch | ec | d6 | Igfbp4 | ic |
| Igf1 | batch | fap | d6 | Igfbp4 | ic |
| Igf1 | batch | ic | d6 | Igfbp4 | ic |
| Igf1 | batch | ec | d6 | Igfbp5 | ic |
| Igf1 | batch | fap | d6 | Igfbp5 | ic |
| Igf1 | batch | ic | d6 | Igfbp5 | ic |
| Igf1 | batch | ec | d6 | Igfbp6 | ic |
| Igf1 | batch | fap | d6 | Igfbp6 | ic |
| Igf1 | batch | ic | d6 | Igfbp6 | ic |
| Igf1 | batch | ec | d6 | Igfbp7 | ic |
| Igf1 | batch | fap | d6 | Igfbp7 | ic |
| Igf1 | batch | ic | d6 | Igfbp7 | ic |
| Igf2 | batch | ec | d6 | Vtn | ic |
| Igf2 | batch | fap | d6 | Vtn | ic |
| Igf2 | batch | ic | d6 | Vtn | ic |
| Igf2 | batch | mp | d6 | Vtn | ic |
| Il16 | batch | fap | d6 | Cd4 | ic |
| Il16 | batch | ic | d6 | Cd4 | ic |
| Pthlh | batch | fap | d6 | Pth1r | ic |
| Ptn | batch | fap | d6 | Ptprs | ic |
| Ptn | constitutive | per | d6 | Ptprs | ic |
| Spp1 | batch | fap | d6 | Itga5 | ic |
| Spp1 | batch | ic | d6 | Itga5 | ic |
| Spp1 | batch | mp | d6 | Itga5 | ic |
| Spp1 | batch | fap | d6 | Itga9 | ic |
| Spp1 | batch | ic | d6 | Itga9 | ic |
| Spp1 | batch | mp | d6 | Itga9 | ic |
| Spp1 | batch | fap | d6 | Itgav | ic |
| Spp1 | batch | ic | d6 | Itgav | ic |
| Spp1 | batch | mp | d6 | Itgav | ic |
| Spp1 | batch | fap | d6 | Itgb5 | ic |
| Spp1 | batch | ic | d6 | Itgb5 | ic |
| Spp1 | batch | mp | d6 | Itgb5 | ic |
| Spp1 | batch | fap | d6 | Vtn | ic |
| Spp1 | batch | ic | d6 | Vtn | ic |
| Spp1 | batch | mp | d6 | Vtn | ic |
| Tgfb3 | batch | fap | d6 | Itgav | ic |
| Tgfb3 | batch | mp | d6 | Itgav | ic |
| Tgfb3 | total | ec | d6 | Itgav | ic |
| Tgfb3 | total | per | d6 | Itgav | ic |
| Tgfb3 | batch | fap | d6 | Tgfbr1 | ic |
| Tgfb3 | batch | mp | d6 | Tgfbr1 | ic |
| Tgfb3 | total | ec | d6 | Tgfbr1 | ic |
| Tgfb3 | total | per | d6 | Tgfbr1 | ic |
| Tgfb3 | batch | fap | d6 | Acvrl1 | ic |
| Tgfb3 | batch | mp | d6 | Acvrl1 | ic |
| Tgfb3 | total | ec | d6 | Acvrl1 | ic |
| Tgfb3 | total | per | d6 | Acvrl1 | ic |
| Apln | batch | ec | d6 | Aplnr | ic |
| Dll1 | batch | ec | d6 | Notch2 | ic |
| Dll1 | total | mp | d6 | Notch2 | ic |
| Il6 | batch | ec | d6 | Il6ra | ic |
| Vegfc | batch | ec | d6 | Itga9 | ic |
| Vegfc | batch | mp | d6 | Itga9 | ic |
| Vegfc | batch | ec | d6 | Nrp2 | ic |
| Vegfc | batch | mp | d6 | Nrp2 | ic |
| Vegfc | batch | ec | d6 | Nrp1 | ic |
| Vegfc | batch | mp | d6 | Nrp1 | ic |
| Ccl11 | batch | mp | d6 | Ccr5 | ic |
| Ccl11 | batch | mp | d6 | Cxcr3 | ic |
| Ccl8 | batch | mp | d6 | Ccr5 | ic |
| Ccl8 | total | fap | d6 | Ccr5 | ic |
| Ccl8 | total | ic | d6 | Ccr5 | ic |
| Clcf1 | batch | mp | d6 | Crlf1 | ic |
| Clcf1 | batch | mp | d6 | Ncoa3 | ic |
| Il10 | batch | ic | d6 | Il10ra | ic |
| Il10 | batch | mp | d6 | Il10ra | ic |
| Jag1 | batch | mp | d6 | Notch2 | ic |
| Jag1 | constitutive | per | d6 | Notch2 | ic |
| Jag2 | batch | mp | d6 | Notch2 | ic |
| Jag2 | constitutive | ec | d6 | Notch2 | ic |
| Lgals3 | batch | ic | d6 | Lgals3bp | ic |
| Lgals3 | batch | mp | d6 | Lgals3bp | ic |
| Ngf | batch | mp | d6 | Sort1 | ic |
| Ngf | constitutive | per | d6 | Sort1 | ic |
| Pdap1 | batch | mp | d6 | Pdgfa | ic |
| Tgfb1 | batch | mp | d6 | Itgav | ic |
| Tgfb1 | batch | mp | d6 | Tgfbr1 | ic |
| Tgfb1 | batch | mp | d6 | Tgfbr3 | ic |
| Tgfb1 | batch | mp | d6 | Vtn | ic |
| Tgfb1 | batch | mp | d6 | Acvrl1 | ic |
| Tgfb2 | batch | mp | d6 | Tgfbr1 | ic |
| Tgfb2 | constitutive | ec | d6 | Tgfbr1 | ic |
| Tgfb2 | constitutive | fap | d6 | Tgfbr1 | ic |
| Tgfb2 | constitutive | per | d6 | Tgfbr1 | ic |
| Tgfb2 | batch | mp | d6 | Tgfbr3 | ic |
| Tgfb2 | constitutive | ec | d6 | Tgfbr3 | ic |
| Tgfb2 | constitutive | fap | d6 | Tgfbr3 | ic |
| Tgfb2 | constitutive | per | d6 | Tgfbr3 | ic |
| Tgfb2 | batch | mp | d6 | Vtn | ic |
| Tgfb2 | constitutive | ec | d6 | Vtn | ic |
| Tgfb2 | constitutive | fap | d6 | Vtn | ic |
| Tgfb2 | constitutive | per | d6 | Vtn | ic |
| Tnfsf12 | batch | mp | d6 | Tnfrsf14 | ic |
| Vegfa | batch | mp | d6 | Itga9 | ic |
| Vegfa | batch | mp | d6 | Vtn | ic |
| Vegfa | batch | mp | d6 | Nrp2 | ic |
| Vegfa | batch | mp | d6 | Nrp1 | ic |
| Hgf | batch | ic | d6 | Vtn | ic |
| Pdgfb | batch | ic | d6 | Pdgfrb | ic |
| Pdgfb | constitutive | ec | d6 | Pdgfrb | ic |
| Tnfsf13 | batch | ic | d6 | Tnfrsf14 | ic |
| Fgf7 | constitutive | fap | d6 | Nrp1 | ic |
| Pgf | constitutive | per | d6 | Nrp2 | ic |
| Pgf | constitutive | per | d6 | Nrp1 | ic |
| Ccl11 | batch | fap | d7 | Ccr3 | ic |
| Ccl11 | batch | mp | d7 | Ccr3 | ic |
| Ccl11 | batch | fap | d7 | Ccr5 | ic |
| Ccl11 | batch | mp | d7 | Ccr5 | ic |
| Ccl11 | batch | fap | d7 | Cxcr3 | ic |
| Ccl11 | batch | mp | d7 | Cxcr3 | ic |
| Crlf1 | batch | ec | d7 | Ctf1 | ic |
| Crlf1 | batch | fap | d7 | Ctf1 | ic |
| Crlf1 | batch | mp | d7 | Ctf1 | ic |
| Ctgf | batch | ec | d7 | Itga5 | ic |
| Ctgf | batch | fap | d7 | Itga5 | ic |
| Ctgf | batch | ic | d7 | Itga5 | ic |
| Ctgf | batch | mp | d7 | Itga5 | ic |
| Ctgf | constitutive | per | d7 | Itga5 | ic |
| Cx3cl1 | batch | fap | d7 | Cx3cr1 | ic |
| Cx3cl1 | batch | mp | d7 | Cx3cr1 | ic |
| Cx3cl1 | total | ec | d7 | Cx3cr1 | ic |
| Cx3cl1 | total | per | d7 | Cx3cr1 | ic |
| Cxcl13 | batch | fap | d7 | Cxcr3 | ic |
| Cxcl13 | batch | fap | d7 | Cxcr5 | ic |
| Fgf1 | batch | fap | d7 | Fgfr1 | ic |
| Fgf1 | constitutive | per | d7 | Fgfr1 | ic |
| Gas6 | batch | fap | d7 | Axl | ic |
| Gas6 | batch | mp | d7 | Axl | ic |
| Igf1 | batch | ec | d7 | Igfbp3 | ic |
| Igf1 | batch | fap | d7 | Igfbp3 | ic |
| Igf1 | batch | ic | d7 | Igfbp3 | ic |
| Igf1 | batch | ec | d7 | Igfbp4 | ic |
| Igf1 | batch | fap | d7 | Igfbp4 | ic |
| Igf1 | batch | ic | d7 | Igfbp4 | ic |
| Igf1 | batch | ec | d7 | Igfbp5 | ic |
| Igf1 | batch | fap | d7 | Igfbp5 | ic |
| Igf1 | batch | ic | d7 | Igfbp5 | ic |
| Igf1 | batch | ec | d7 | Igfbp6 | ic |
| Igf1 | batch | fap | d7 | Igfbp6 | ic |
| Igf1 | batch | ic | d7 | Igfbp6 | ic |
| Igf1 | batch | ec | d7 | Igfbp7 | ic |
| Igf1 | batch | fap | d7 | Igfbp7 | ic |
| Igf1 | batch | ic | d7 | Igfbp7 | ic |
| Igf2 | batch | ec | d7 | Vtn | ic |
| Igf2 | batch | fap | d7 | Vtn | ic |
| Igf2 | batch | ic | d7 | Vtn | ic |
| Igf2 | batch | mp | d7 | Vtn | ic |
| Il16 | batch | fap | d7 | Cd4 | ic |
| Il16 | batch | ic | d7 | Cd4 | ic |
| Il18 | batch | fap | d7 | Il18rap | ic |
| Il18 | constitutive | ic | d7 | Il18rap | ic |
| Il18 | constitutive | mp | d7 | Il18rap | ic |
| Nov | batch | ec | d7 | Itga5 | ic |
| Nov | batch | fap | d7 | Itga5 | ic |
| Nov | batch | ec | d7 | Itgb3 | ic |
| Nov | batch | fap | d7 | Itgb3 | ic |
| Pgf | batch | fap | d7 | Nrp2 | ic |
| Pgf | constitutive | per | d7 | Nrp2 | ic |
| Pgf | batch | fap | d7 | Nrp1 | ic |
| Pgf | constitutive | per | d7 | Nrp1 | ic |
| Pthlh | batch | fap | d7 | Pth1r | ic |
| Ptn | batch | fap | d7 | Ptprs | ic |
| Ptn | constitutive | per | d7 | Ptprs | ic |
| S100b | batch | fap | d7 | Fgfr1 | ic |
| S100b | batch | mp | d7 | Fgfr1 | ic |
| S100b | constitutive | per | d7 | Fgfr1 | ic |
| Tgfb3 | batch | fap | d7 | Acvrl1 | ic |
| Tgfb3 | batch | mp | d7 | Acvrl1 | ic |
| Tgfb3 | total | ec | d7 | Acvrl1 | ic |
| Tgfb3 | total | per | d7 | Acvrl1 | ic |
| Vegfa | batch | fap | d7 | Itga9 | ic |
| Vegfa | batch | mp | d7 | Itga9 | ic |
| Vegfa | batch | fap | d7 | Vtn | ic |
| Vegfa | batch | mp | d7 | Vtn | ic |
| Vegfa | batch | fap | d7 | Nrp2 | ic |
| Vegfa | batch | mp | d7 | Nrp2 | ic |
| Vegfa | batch | fap | d7 | Nrp1 | ic |
| Vegfa | batch | mp | d7 | Nrp1 | ic |
| Wnt2 | batch | fap | d7 | Sfrp1 | ic |
| Cxcl10 | batch | ec | d7 | Ccr3 | ic |
| Cxcl10 | batch | ic | d7 | Ccr3 | ic |
| Cxcl10 | batch | ec | d7 | Cxcr3 | ic |
| Cxcl10 | batch | ic | d7 | Cxcr3 | ic |
| Cxcl9 | batch | ec | d7 | Ccr3 | ic |
| Cxcl9 | batch | ic | d7 | Ccr3 | ic |
| Cxcl9 | batch | ec | d7 | Cxcr3 | ic |
| Cxcl9 | batch | ic | d7 | Cxcr3 | ic |
| Dll1 | batch | ec | d7 | Notch2 | ic |
| Dll1 | total | mp | d7 | Notch2 | ic |
| Esm1 | batch | ec | d7 | Itgal | ic |
| Esm1 | batch | mp | d7 | Itgal | ic |
| Il15 | batch | ec | d7 | Il2rb | ic |
| Il15 | batch | ic | d7 | Il2rb | ic |
| Vegfc | batch | ec | d7 | Itga9 | ic |
| Vegfc | batch | mp | d7 | Itga9 | ic |
| Vegfc | batch | ec | d7 | Nrp2 | ic |
| Vegfc | batch | mp | d7 | Nrp2 | ic |
| Vegfc | batch | ec | d7 | Nrp1 | ic |
| Vegfc | batch | mp | d7 | Nrp1 | ic |
| Ccl3 | batch | mp | d7 | Ccr3 | ic |
| Ccl3 | batch | mp | d7 | Ccr5 | ic |
| Ccl3 | batch | mp | d7 | Cxcr5 | ic |
| Ccl4 | batch | mp | d7 | Ccr3 | ic |
| Ccl4 | batch | mp | d7 | Ccr5 | ic |
| Ccl4 | batch | mp | d7 | Cxcr5 | ic |
| Ccl8 | batch | mp | d7 | Ccr3 | ic |
| Ccl8 | total | fap | d7 | Ccr3 | ic |
| Ccl8 | total | ic | d7 | Ccr3 | ic |
| Ccl8 | batch | mp | d7 | Ccr5 | ic |
| Ccl8 | total | fap | d7 | Ccr5 | ic |
| Ccl8 | total | ic | d7 | Ccr5 | ic |
| Clcf1 | batch | ic | d7 | Crlf1 | ic |
| Clcf1 | batch | mp | d7 | Crlf1 | ic |
| Clcf1 | batch | ic | d7 | Ncoa3 | ic |
| Clcf1 | batch | mp | d7 | Ncoa3 | ic |
| Cxcl16 | batch | ic | d7 | Cxcr6 | ic |
| Cxcl16 | batch | mp | d7 | Cxcr6 | ic |
| Ebi3 | batch | mp | d7 | Il27ra | ic |
| Ebi3 | constitutive | ic | d7 | Il27ra | ic |
| Gdnf | batch | mp | d7 | Gfra2 | ic |
| Il10 | batch | ic | d7 | Il10ra | ic |
| Il10 | batch | mp | d7 | Il10ra | ic |
| Il33 | batch | mp | d7 | Il1rl1 | ic |
| Jag1 | batch | mp | d7 | Notch2 | ic |
| Jag1 | constitutive | per | d7 | Notch2 | ic |
| Jag2 | batch | mp | d7 | Notch2 | ic |
| Jag2 | constitutive | ec | d7 | Notch2 | ic |
| Kitl | batch | ic | d7 | Kit | ic |
| Kitl | batch | mp | d7 | Kit | ic |
| Kitl | constitutive | ec | d7 | Kit | ic |
| Kitl | constitutive | per | d7 | Kit | ic |
| Pdap1 | batch | mp | d7 | Pdgfa | ic |
| Rabep1 | batch | mp | d7 | Lsr | ic |
| Spp1 | batch | mp | d7 | Itga5 | ic |
| Spp1 | batch | mp | d7 | Itga9 | ic |
| Spp1 | batch | mp | d7 | Itgb3 | ic |
| Spp1 | batch | mp | d7 | Vtn | ic |
| Tgfb1 | batch | mp | d7 | Tgfbr3 | ic |
| Tgfb1 | batch | mp | d7 | Vtn | ic |
| Tgfb1 | batch | mp | d7 | Acvrl1 | ic |
| Tgfb2 | batch | mp | d7 | Tgfbr3 | ic |
| Tgfb2 | constitutive | ec | d7 | Tgfbr3 | ic |
| Tgfb2 | constitutive | fap | d7 | Tgfbr3 | ic |
| Tgfb2 | constitutive | per | d7 | Tgfbr3 | ic |
| Tgfb2 | batch | mp | d7 | Vtn | ic |
| Tgfb2 | constitutive | ec | d7 | Vtn | ic |
| Tgfb2 | constitutive | fap | d7 | Vtn | ic |
| Tgfb2 | constitutive | per | d7 | Vtn | ic |
| Tnfsf12 | batch | mp | d7 | Tnfrsf14 | ic |
| Btla | batch | ic | d7 | Tnfrsf14 | ic |
| Ccl5 | batch | ic | d7 | Ccr3 | ic |
| Ccl5 | batch | ic | d7 | Ccr5 | ic |
| Ccl5 | batch | ic | d7 | Cxcr3 | ic |
| Ccl5 | batch | ic | d7 | Cxcr5 | ic |
| Ifng | batch | ic | d7 | Ifngr2 | ic |
| Ifng | batch | ic | d7 | Ifngr1 | ic |
| Il6 | batch | ic | d7 | Il6ra | ic |
| Pdgfb | batch | ic | d7 | Pdgfrb | ic |
| Pdgfb | constitutive | ec | d7 | Pdgfrb | ic |
| Fgf7 | constitutive | fap | d7 | Nrp1 | ic |
| Lif | constitutive | mp | d7 | Lifr | ic |
| Nampt | constitutive | ic | d7 | Adora2a | ic |
| Ccl11 | batch | fap | d10 | Ccr3 | ic |
| Ccl11 | batch | mp | d10 | Ccr3 | ic |
| Ccl11 | batch | per | d10 | Ccr3 | ic |
| Ccl11 | batch | fap | d10 | Ccr5 | ic |
| Ccl11 | batch | mp | d10 | Ccr5 | ic |
| Ccl11 | batch | per | d10 | Ccr5 | ic |
| Ccl11 | batch | fap | d10 | Cxcr3 | ic |
| Ccl11 | batch | mp | d10 | Cxcr3 | ic |
| Ccl11 | batch | per | d10 | Cxcr3 | ic |
| Crlf1 | batch | fap | d10 | Ctf1 | ic |
| Crlf1 | batch | mp | d10 | Ctf1 | ic |
| Csf2 | batch | fap | d10 | Csf3r | ic |
| Csf2 | batch | fap | d10 | Il3ra | ic |
| Csf2 | batch | fap | d10 | Csf2rb | ic |
| Csf2 | batch | fap | d10 | Csf2ra | ic |
| Ctgf | batch | fap | d10 | Itga5 | ic |
| Ctgf | batch | mp | d10 | Itga5 | ic |
| Ctgf | constitutive | per | d10 | Itga5 | ic |
| Cx3cl1 | batch | fap | d10 | Cx3cr1 | ic |
| Cxcl13 | batch | fap | d10 | Cxcr3 | ic |
| Cxcl13 | batch | fap | d10 | Cxcr5 | ic |
| Fgf1 | batch | fap | d10 | Fgfr1 | ic |
| Fgf1 | batch | mp | d10 | Fgfr1 | ic |
| Fgf1 | constitutive | per | d10 | Fgfr1 | ic |
| Gas6 | batch | fap | d10 | Axl | ic |
| Gas6 | batch | mp | d10 | Axl | ic |
| Hbegf | batch | fap | d10 | Cd44 | ic |
| Hbegf | batch | mp | d10 | Cd44 | ic |
| Hbegf | constitutive | ec | d10 | Cd44 | ic |
| Hbegf | constitutive | per | d10 | Cd44 | ic |
| Il16 | batch | fap | d10 | Cd4 | ic |
| Il16 | batch | ic | d10 | Cd4 | ic |
| Il18 | batch | fap | d10 | Il18rap | ic |
| Il18 | constitutive | ic | d10 | Il18rap | ic |
| Il18 | constitutive | mp | d10 | Il18rap | ic |
| Kitl | batch | fap | d10 | Kit | ic |
| Kitl | constitutive | ec | d10 | Kit | ic |
| Kitl | constitutive | per | d10 | Kit | ic |
| Mdk | batch | fap | d10 | Lrp1 | ic |
| Mdk | batch | mp | d10 | Lrp1 | ic |
| Nov | batch | fap | d10 | Itga5 | ic |
| Nov | batch | per | d10 | Itga5 | ic |
| Nov | batch | fap | d10 | Itgb3 | ic |
| Nov | batch | per | d10 | Itgb3 | ic |
| Pgf | batch | fap | d10 | Nrp2 | ic |
| Pgf | batch | mp | d10 | Nrp2 | ic |
| Pgf | constitutive | per | d10 | Nrp2 | ic |
| Pgf | batch | fap | d10 | Nrp1 | ic |
| Pgf | batch | mp | d10 | Nrp1 | ic |
| Pgf | constitutive | per | d10 | Nrp1 | ic |
| Ptn | batch | fap | d10 | Ptprs | ic |
| Ptn | batch | mp | d10 | Ptprs | ic |
| Ptn | constitutive | per | d10 | Ptprs | ic |
| Ptn | batch | fap | d10 | Plxnb2 | ic |
| Ptn | batch | mp | d10 | Plxnb2 | ic |
| Ptn | constitutive | per | d10 | Plxnb2 | ic |
| S100b | batch | fap | d10 | Fgfr1 | ic |
| S100b | batch | mp | d10 | Fgfr1 | ic |
| S100b | constitutive | per | d10 | Fgfr1 | ic |
| Tgfb3 | batch | fap | d10 | Acvrl1 | ic |
| Tgfb3 | batch | mp | d10 | Acvrl1 | ic |
| Tslp | batch | fap | d10 | Il7r | ic |
| Tslp | constitutive | ec | d10 | Il7r | ic |
| Tslp | batch | fap | d10 | Crlf2 | ic |
| Tslp | constitutive | ec | d10 | Crlf2 | ic |
| Vegfa | batch | fap | d10 | Itga9 | ic |
| Vegfa | batch | ic | d10 | Itga9 | ic |
| Vegfa | batch | mp | d10 | Itga9 | ic |
| Vegfa | batch | fap | d10 | Nrp2 | ic |
| Vegfa | batch | ic | d10 | Nrp2 | ic |
| Vegfa | batch | mp | d10 | Nrp2 | ic |
| Vegfa | batch | fap | d10 | Nrp1 | ic |
| Vegfa | batch | ic | d10 | Nrp1 | ic |
| Vegfa | batch | mp | d10 | Nrp1 | ic |
| Wnt2 | batch | fap | d10 | Sfrp1 | ic |
| Csf3 | batch | ec | d10 | Csf3r | ic |
| Cxcl10 | batch | ec | d10 | Ccr3 | ic |
| Cxcl10 | batch | ic | d10 | Ccr3 | ic |
| Cxcl10 | batch | ec | d10 | Cxcr3 | ic |
| Cxcl10 | batch | ic | d10 | Cxcr3 | ic |
| Cxcl9 | batch | ec | d10 | Ccr3 | ic |
| Cxcl9 | batch | ic | d10 | Ccr3 | ic |
| Cxcl9 | batch | ec | d10 | Cxcr3 | ic |
| Cxcl9 | batch | ic | d10 | Cxcr3 | ic |
| Dll1 | batch | ec | d10 | Notch2 | ic |
| Il15 | batch | ec | d10 | Il2rb | ic |
| Il15 | batch | ic | d10 | Il2rb | ic |
| Il15 | batch | ec | d10 | Il17ra | ic |
| Il15 | batch | ic | d10 | Il17ra | ic |
| Il15 | batch | ec | d10 | Il2rg | ic |
| Il15 | batch | ic | d10 | Il2rg | ic |
| Vegfc | batch | ec | d10 | Itga9 | ic |
| Vegfc | batch | ec | d10 | Nrp2 | ic |
| Vegfc | batch | ec | d10 | Nrp1 | ic |
| Clcf1 | batch | ic | d10 | Crlf1 | ic |
| Clcf1 | batch | mp | d10 | Crlf1 | ic |
| Clcf1 | batch | ic | d10 | Ncoa3 | ic |
| Clcf1 | batch | mp | d10 | Ncoa3 | ic |
| Cxcl1 | batch | ic | d10 | Cxcr2 | ic |
| Cxcl1 | batch | mp | d10 | Cxcr2 | ic |
| Cxcl12 | batch | mp | d10 | Cxcr4 | ic |
| Cxcl12 | constitutive | ec | d10 | Cxcr4 | ic |
| Cxcl12 | constitutive | per | d10 | Cxcr4 | ic |
| Fstl1 | batch | mp | d10 | Cd14 | ic |
| Fstl1 | constitutive | fap | d10 | Cd14 | ic |
| Jag1 | batch | ic | d10 | Notch2 | ic |
| Jag1 | batch | mp | d10 | Notch2 | ic |
| Jag1 | constitutive | per | d10 | Notch2 | ic |
| Jag2 | batch | mp | d10 | Notch2 | ic |
| Jag2 | constitutive | ec | d10 | Notch2 | ic |
| Tnfsf12 | batch | mp | d10 | Tnfrsf14 | ic |
| Tnfsf12 | total | ec | d10 | Tnfrsf14 | ic |
| Tnfsf12 | total | ic | d10 | Tnfrsf14 | ic |
| Tnfsf12 | total | per | d10 | Tnfrsf14 | ic |
| Vegfb | batch | mp | d10 | Nrp1 | ic |
| Btla | batch | ic | d10 | Tnfrsf14 | ic |
| Ccl2 | batch | ic | d10 | Ccr1 | ic |
| Ccl2 | batch | ic | d10 | Ccr3 | ic |
| Ccl2 | batch | ic | d10 | Ccr5 | ic |
| Ccl2 | batch | ic | d10 | Ccr2 | ic |
| Ccl3 | batch | ic | d10 | Ccr1 | ic |
| Ccl3 | batch | ic | d10 | Ccr3 | ic |
| Ccl3 | batch | ic | d10 | Ccr5 | ic |
| Ccl3 | batch | ic | d10 | Cxcr5 | ic |
| Ccl4 | batch | ic | d10 | Ccr1 | ic |
| Ccl4 | batch | ic | d10 | Ccr3 | ic |
| Ccl4 | batch | ic | d10 | Ccr5 | ic |
| Ccl4 | batch | ic | d10 | Cxcr5 | ic |
| Ccl5 | batch | ic | d10 | Ccr1 | ic |
| Ccl5 | batch | ic | d10 | Ccr3 | ic |
| Ccl5 | batch | ic | d10 | Ccr5 | ic |
| Ccl5 | batch | ic | d10 | Cxcr3 | ic |
| Ccl5 | batch | ic | d10 | Cxcr5 | ic |
| Ccl5 | batch | ic | d10 | Sdc4 | ic |
| Ccl6 | batch | ic | d10 | Ccr1 | ic |
| Ccl7 | batch | ic | d10 | Ccr1 | ic |
| Ccl7 | batch | ic | d10 | Ccr3 | ic |
| Ccl7 | batch | ic | d10 | Ccr5 | ic |
| Ccl7 | batch | ic | d10 | Cxcr3 | ic |
| Csf1 | batch | ic | d10 | Csf1r | ic |
| Cxcl16 | batch | ic | d10 | Cxcr6 | ic |
| Cxcl2 | batch | ic | d10 | Cxcr2 | ic |
| Cxcl3 | batch | ic | d10 | Cxcr2 | ic |
| Ifnb1 | batch | ic | d10 | Ifnar2 | ic |
| Ifng | batch | ic | d10 | Ifngr2 | ic |
| Ifng | batch | ic | d10 | Ifngr1 | ic |
| Il10 | batch | ic | d10 | Ifnar2 | ic |
| Il10 | batch | ic | d10 | Il10rb | ic |
| Il10 | batch | ic | d10 | Il10ra | ic |
| Il1a | batch | ic | d10 | Il1r2 | ic |
| Il1b | batch | ic | d10 | Adrb2 | ic |
| Il1b | batch | ic | d10 | Il1r2 | ic |
| Il6 | batch | ic | d10 | Il6ra | ic |
| Lif | batch | ic | d10 | Lifr | ic |
| Lif | constitutive | mp | d10 | Lifr | ic |
| Mmp12 | batch | ic | d10 | Plaur | ic |
| Osm | batch | ic | d10 | Lifr | ic |
| Pdgfb | batch | ic | d10 | Lrp1 | ic |
| Pdgfb | constitutive | ec | d10 | Lrp1 | ic |
| Pf4 | batch | ic | d10 | Cxcr3 | ic |
| Pf4 | batch | ic | d10 | Ldlr | ic |
| Ppbp | batch | ic | d10 | Cxcr2 | ic |
| Ppbp | batch | ic | d10 | Il7r | ic |
| Ppbp | batch | ic | d10 | Ncoa3 | ic |
| Tnf | batch | ic | d10 | Tnfrsf1b | ic |
| Tnfsf14 | batch | ic | d10 | Tnfrsf14 | ic |
| Gdnf | batch | per | d10 | Gfra2 | ic |
| Ccl8 | total | fap | d10 | Ccr1 | ic |
| Ccl8 | total | ic | d10 | Ccr1 | ic |
| Ccl8 | total | fap | d10 | Ccr3 | ic |
| Ccl8 | total | ic | d10 | Ccr3 | ic |
| Ccl8 | total | fap | d10 | Ccr5 | ic |
| Ccl8 | total | ic | d10 | Ccr5 | ic |
| Ccl8 | total | fap | d10 | Ccr2 | ic |
| Ccl8 | total | ic | d10 | Ccr2 | ic |
| Tgfb1 | total | ec | d10 | Acvrl1 | ic |
| Tgfb1 | total | ic | d10 | Acvrl1 | ic |
| Fgf7 | constitutive | fap | d10 | Nrp1 | ic |
| Ebi3 | constitutive | ic | d10 | Il27ra | ic |
| Nampt | constitutive | ic | d10 | Adora2a | ic |
| Ccl11 | batch | fap | d14 | Ccr5 | ic |
| Ccl11 | batch | fap | d14 | Cxcr3 | ic |
| Ctgf | batch | fap | d14 | Itga5 | ic |
| Ctgf | batch | ic | d14 | Itga5 | ic |
| Ctgf | constitutive | per | d14 | Itga5 | ic |
| Cx3cl1 | batch | fap | d14 | Cx3cr1 | ic |
| Cx3cl1 | total | ec | d14 | Cx3cr1 | ic |
| Cx3cl1 | total | per | d14 | Cx3cr1 | ic |
| Cxcl13 | batch | fap | d14 | Cxcr3 | ic |
| Gas6 | batch | fap | d14 | Axl | ic |
| Igf1 | batch | ec | d14 | Igfbp3 | ic |
| Igf1 | batch | fap | d14 | Igfbp3 | ic |
| Igf1 | batch | ic | d14 | Igfbp3 | ic |
| Igf1 | batch | ec | d14 | Igfbp4 | ic |
| Igf1 | batch | fap | d14 | Igfbp4 | ic |
| Igf1 | batch | ic | d14 | Igfbp4 | ic |
| Igf1 | batch | ec | d14 | Igfbp5 | ic |
| Igf1 | batch | fap | d14 | Igfbp5 | ic |
| Igf1 | batch | ic | d14 | Igfbp5 | ic |
| Igf1 | batch | ec | d14 | Igfbp6 | ic |
| Igf1 | batch | fap | d14 | Igfbp6 | ic |
| Igf1 | batch | ic | d14 | Igfbp6 | ic |
| Igf1 | batch | ec | d14 | Igfbp7 | ic |
| Igf1 | batch | fap | d14 | Igfbp7 | ic |
| Igf1 | batch | ic | d14 | Igfbp7 | ic |
| Igf2 | batch | ec | d14 | Vtn | ic |
| Igf2 | batch | fap | d14 | Vtn | ic |
| Igf2 | batch | ic | d14 | Vtn | ic |
| Il16 | batch | fap | d14 | Cd4 | ic |
| Il16 | batch | ic | d14 | Cd4 | ic |
| Nov | batch | fap | d14 | Itga5 | ic |
| Pgf | batch | fap | d14 | Nrp2 | ic |
| Pgf | constitutive | per | d14 | Nrp2 | ic |
| Pgf | batch | fap | d14 | Nrp1 | ic |
| Pgf | constitutive | per | d14 | Nrp1 | ic |
| Pthlh | batch | fap | d14 | Pth1r | ic |
| Ptn | batch | fap | d14 | Ptprs | ic |
| Ptn | constitutive | per | d14 | Ptprs | ic |
| Tgfb3 | batch | fap | d14 | Acvrl1 | ic |
| Vegfa | batch | fap | d14 | Itga9 | ic |
| Vegfa | batch | fap | d14 | Vtn | ic |
| Vegfa | batch | fap | d14 | Nrp2 | ic |
| Vegfa | batch | fap | d14 | Nrp1 | ic |
| Wnt2 | batch | fap | d14 | Sfrp1 | ic |
| Cxcl10 | batch | ec | d14 | Cxcr3 | ic |
| Cxcl10 | batch | ic | d14 | Cxcr3 | ic |
| Cxcl9 | batch | ec | d14 | Cxcr3 | ic |
| Cxcl9 | batch | ic | d14 | Cxcr3 | ic |
| Dll1 | batch | ec | d14 | Notch2 | ic |
| Vegfc | batch | ec | d14 | Itga9 | ic |
| Vegfc | batch | ec | d14 | Nrp2 | ic |
| Vegfc | batch | ec | d14 | Nrp1 | ic |
| Il10 | batch | ic | d14 | Il10ra | ic |
| Pdgfb | batch | ic | d14 | Pdgfrb | ic |
| Pdgfb | constitutive | ec | d14 | Pdgfrb | ic |
| Fgf7 | constitutive | fap | d14 | Nrp1 | ic |
| Tgfb2 | constitutive | ec | d14 | Tgfbr3 | ic |
| Tgfb2 | constitutive | fap | d14 | Tgfbr3 | ic |
| Tgfb2 | constitutive | per | d14 | Tgfbr3 | ic |
| Tgfb2 | constitutive | ec | d14 | Vtn | ic |
| Tgfb2 | constitutive | fap | d14 | Vtn | ic |
| Tgfb2 | constitutive | per | d14 | Vtn | ic |
| Jag2 | constitutive | ec | d14 | Notch2 | ic |
| Jag1 | constitutive | per | d14 | Notch2 | ic |
| Bmp2 | batch | fap | d0 | Bmpr1a | per |
| Bmp2 | batch | mp | d0 | Bmpr1a | per |
| Bmp2 | constitutive | per | d0 | Bmpr1a | per |
| Bmp4 | batch | fap | d0 | Bmpr1a | per |
| Bmp4 | batch | mp | d0 | Bmpr1a | per |
| Bmp4 | constitutive | ec | d0 | Bmpr1a | per |
| Bmp4 | constitutive | per | d0 | Bmpr1a | per |
| Bmp6 | batch | ec | d0 | Bmpr1a | per |
| Bmp6 | batch | fap | d0 | Bmpr1a | per |
| Bmp6 | batch | mp | d0 | Bmpr1a | per |
| Bmp6 | constitutive | per | d0 | Bmpr1a | per |
| Bmp7 | batch | fap | d0 | Bmpr1a | per |
| Clu | batch | ec | d0 | Vldlr | per |
| Clu | batch | fap | d0 | Vldlr | per |
| Clu | batch | ic | d0 | Vldlr | per |
| Clu | batch | mp | d0 | Vldlr | per |
| Crlf1 | batch | ec | d0 | Ctf1 | per |
| Crlf1 | batch | fap | d0 | Ctf1 | per |
| Crlf1 | batch | mp | d0 | Ctf1 | per |
| Edn1 | batch | fap | d0 | Ednra | per |
| Edn1 | constitutive | ec | d0 | Ednra | per |
| Efnb1 | batch | fap | d0 | Ephb2 | per |
| Efnb1 | batch | ic | d0 | Ephb2 | per |
| Efnb1 | constitutive | ec | d0 | Ephb2 | per |
| Efnb1 | constitutive | per | d0 | Ephb2 | per |
| Fgf1 | batch | fap | d0 | Fgfr2 | per |
| Fgf1 | constitutive | per | d0 | Fgfr2 | per |
| Fgf1 | batch | fap | d0 | Fgfr1 | per |
| Fgf1 | constitutive | per | d0 | Fgfr1 | per |
| Gas6 | batch | fap | d0 | Axl | per |
| Gas6 | batch | mp | d0 | Axl | per |
| Igf1 | batch | ec | d0 | Igfbp7 | per |
| Igf1 | batch | fap | d0 | Igfbp7 | per |
| Igf1 | batch | ic | d0 | Igfbp7 | per |
| Kitl | batch | fap | d0 | Kit | per |
| Kitl | batch | ic | d0 | Kit | per |
| Kitl | batch | mp | d0 | Kit | per |
| Kitl | constitutive | ec | d0 | Kit | per |
| Kitl | constitutive | per | d0 | Kit | per |
| Nov | batch | ec | d0 | Notch1 | per |
| Nov | batch | fap | d0 | Notch1 | per |
| Nov | batch | per | d0 | Notch1 | per |
| Nov | batch | ec | d0 | Itgb3 | per |
| Nov | batch | fap | d0 | Itgb3 | per |
| Nov | batch | per | d0 | Itgb3 | per |
| Nov | batch | ec | d0 | Itgav | per |
| Nov | batch | fap | d0 | Itgav | per |
| Nov | batch | per | d0 | Itgav | per |
| Ntf3 | batch | ec | d0 | Ntrk2 | per |
| Ntf3 | batch | fap | d0 | Ntrk2 | per |
| Ntf3 | batch | mp | d0 | Ntrk2 | per |
| Ntf3 | constitutive | per | d0 | Ntrk2 | per |
| Pthlh | batch | fap | d0 | Pth1r | per |
| Ptn | batch | fap | d0 | Ptprs | per |
| Ptn | constitutive | per | d0 | Ptprs | per |
| S100b | batch | fap | d0 | Fgfr1 | per |
| S100b | batch | mp | d0 | Fgfr1 | per |
| S100b | constitutive | per | d0 | Fgfr1 | per |
| Sfrp4 | batch | ec | d0 | Fzd2 | per |
| Sfrp4 | batch | fap | d0 | Fzd2 | per |
| Sfrp4 | batch | mp | d0 | Fzd2 | per |
| Vegfa | batch | fap | d0 | Ephb2 | per |
| Vegfa | batch | mp | d0 | Ephb2 | per |
| Vegfa | batch | fap | d0 | Vtn | per |
| Vegfa | batch | mp | d0 | Vtn | per |
| Vegfa | batch | fap | d0 | Itga9 | per |
| Vegfa | batch | mp | d0 | Itga9 | per |
| Wnt11 | batch | fap | d0 | Fzd4 | per |
| Wnt11 | batch | ic | d0 | Fzd4 | per |
| Wnt2 | batch | fap | d0 | Sfrp1 | per |
| Wnt2 | batch | fap | d0 | Fzd1 | per |
| Dll1 | batch | ec | d0 | Notch3 | per |
| Dll1 | batch | ec | d0 | Notch1 | per |
| Dll1 | batch | ec | d0 | Notch2 | per |
| Fgf9 | batch | ec | d0 | Fgfr2 | per |
| Igf2 | batch | ec | d0 | Vtn | per |
| Il15 | batch | ec | d0 | Il15ra | per |
| Il15 | batch | ic | d0 | Il15ra | per |
| Sfrp5 | batch | ec | d0 | Fzd2 | per |
| Sfrp5 | batch | mp | d0 | Fzd2 | per |
| Sfrp5 | batch | per | d0 | Fzd2 | per |
| Vegfc | batch | ec | d0 | Itga9 | per |
| Vegfc | batch | mp | d0 | Itga9 | per |
| Areg | batch | mp | d0 | Egfr | per |
| Clcf1 | batch | ic | d0 | Crlf1 | per |
| Clcf1 | batch | mp | d0 | Crlf1 | per |
| Clcf1 | batch | ic | d0 | Cntfr | per |
| Clcf1 | batch | mp | d0 | Cntfr | per |
| Ctf1 | batch | mp | d0 | Il6st | per |
| Edn3 | batch | mp | d0 | Ednra | per |
| Gdnf | batch | mp | d0 | Gfra1 | per |
| Gdnf | batch | per | d0 | Gfra1 | per |
| Gdnf | batch | mp | d0 | Gfra2 | per |
| Gdnf | batch | per | d0 | Gfra2 | per |
| Gmfb | batch | mp | d0 | Egfr | per |
| Gmfb | batch | mp | d0 | Itpr3 | per |
| Hbegf | batch | mp | d0 | Egfr | per |
| Hbegf | constitutive | ec | d0 | Egfr | per |
| Hbegf | constitutive | per | d0 | Egfr | per |
| Jag1 | batch | mp | d0 | Notch3 | per |
| Jag1 | constitutive | per | d0 | Notch3 | per |
| Jag1 | batch | mp | d0 | Notch1 | per |
| Jag1 | constitutive | per | d0 | Notch1 | per |
| Jag1 | batch | mp | d0 | Notch2 | per |
| Jag1 | constitutive | per | d0 | Notch2 | per |
| Jag2 | batch | mp | d0 | Notch3 | per |
| Jag2 | constitutive | ec | d0 | Notch3 | per |
| Jag2 | batch | mp | d0 | Notch1 | per |
| Jag2 | constitutive | ec | d0 | Notch1 | per |
| Jag2 | batch | mp | d0 | Notch2 | per |
| Jag2 | constitutive | ec | d0 | Notch2 | per |
| Ngf | batch | mp | d0 | Sort1 | per |
| Ngf | constitutive | per | d0 | Sort1 | per |
| Nppc | batch | mp | d0 | Npr2 | per |
| Serpini1 | batch | mp | d0 | Plat | per |
| Serpini1 | constitutive | per | d0 | Plat | per |
| Tgfb3 | batch | mp | d0 | Itgav | per |
| Il6 | batch | ic | d0 | Il6st | per |
| Pdgfb | batch | ic | d0 | Pdgfrb | per |
| Pdgfb | constitutive | ec | d0 | Pdgfrb | per |
| Adm | batch | per | d0 | Calcrl | per |
| Dll4 | total | ec | d0 | Notch1 | per |
| Fgf7 | constitutive | fap | d0 | Fgfr2 | per |
| Tgfb2 | constitutive | ec | d0 | Tgfbr3 | per |
| Tgfb2 | constitutive | fap | d0 | Tgfbr3 | per |
| Tgfb2 | constitutive | per | d0 | Tgfbr3 | per |
| Tgfb2 | constitutive | ec | d0 | Vtn | per |
| Tgfb2 | constitutive | fap | d0 | Vtn | per |
| Tgfb2 | constitutive | per | d0 | Vtn | per |
| Bmp5 | constitutive | ec | d0 | Bmpr1a | per |
| Bmp5 | constitutive | fap | d0 | Bmpr1a | per |
| Bmp5 | constitutive | per | d0 | Bmpr1a | per |
| Cmtm8 | constitutive | ec | d0 | Egfr | per |
| Lif | constitutive | mp | d0 | Il6st | per |
| Nampt | constitutive | ic | d0 | Adora2a | per |
| Sfrp1 | constitutive | per | d0 | Fzd2 | per |
| Adm | batch | fap | d1 | Calcrl | per |
| Adm | batch | per | d1 | Calcrl | per |
| Bmp2 | batch | ec | d1 | Bmpr1a | per |
| Bmp2 | batch | fap | d1 | Bmpr1a | per |
| Bmp2 | batch | mp | d1 | Bmpr1a | per |
| Bmp2 | constitutive | per | d1 | Bmpr1a | per |
| Bmp6 | batch | fap | d1 | Bmpr1a | per |
| Bmp6 | constitutive | per | d1 | Bmpr1a | per |
| Ccl2 | batch | ec | d1 | Ccr2 | per |
| Ccl2 | batch | fap | d1 | Ccr2 | per |
| Ccl2 | batch | ic | d1 | Ccr2 | per |
| Ccl2 | batch | per | d1 | Ccr2 | per |
| Ccl2 | batch | ec | d1 | Ccr5 | per |
| Ccl2 | batch | fap | d1 | Ccr5 | per |
| Ccl2 | batch | ic | d1 | Ccr5 | per |
| Ccl2 | batch | per | d1 | Ccr5 | per |
| Ccl7 | batch | ec | d1 | Ccr5 | per |
| Ccl7 | batch | fap | d1 | Ccr5 | per |
| Ccl7 | batch | ic | d1 | Ccr5 | per |
| Ccl7 | batch | per | d1 | Ccr5 | per |
| Efnb1 | batch | fap | d1 | Ephb2 | per |
| Efnb1 | batch | ic | d1 | Ephb2 | per |
| Efnb1 | constitutive | ec | d1 | Ephb2 | per |
| Efnb1 | constitutive | per | d1 | Ephb2 | per |
| Hgf | batch | fap | d1 | Vtn | per |
| Il6 | batch | ec | d1 | Il6st | per |
| Il6 | batch | fap | d1 | Il6st | per |
| Il6 | batch | per | d1 | Il6st | per |
| Kitl | batch | fap | d1 | Kit | per |
| Kitl | batch | ic | d1 | Kit | per |
| Kitl | batch | mp | d1 | Kit | per |
| Kitl | constitutive | ec | d1 | Kit | per |
| Kitl | constitutive | per | d1 | Kit | per |
| Lif | batch | fap | d1 | Il6st | per |
| Lif | batch | ic | d1 | Il6st | per |
| Lif | batch | per | d1 | Il6st | per |
| Lif | constitutive | mp | d1 | Il6st | per |
| Pf4 | batch | fap | d1 | Ldlr | per |
| Pf4 | batch | ic | d1 | Ldlr | per |
| Pf4 | batch | per | d1 | Ldlr | per |
| Tnfrsf11b | batch | fap | d1 | Vtn | per |
| Wnt2 | batch | fap | d1 | Sfrp1 | per |
| Wnt2 | batch | fap | d1 | Fzd1 | per |
| Il1b | batch | ec | d1 | Il1rap | per |
| Il1b | batch | ic | d1 | Il1rap | per |
| Il1b | batch | per | d1 | Il1rap | per |
| Il1b | batch | ec | d1 | Adrb2 | per |
| Il1b | batch | ic | d1 | Adrb2 | per |
| Il1b | batch | per | d1 | Adrb2 | per |
| Areg | batch | ic | d1 | Egfr | per |
| Areg | batch | mp | d1 | Egfr | per |
| Bmp4 | batch | mp | d1 | Bmpr1a | per |
| Bmp4 | constitutive | ec | d1 | Bmpr1a | per |
| Bmp4 | constitutive | per | d1 | Bmpr1a | per |
| C4b | batch | mp | d1 | C3ar1 | per |
| C4b | constitutive | per | d1 | C3ar1 | per |
| Ccl11 | batch | mp | d1 | Ccr5 | per |
| Ccl11 | batch | per | d1 | Ccr5 | per |
| Clcf1 | batch | mp | d1 | Crlf1 | per |
| Clu | batch | ic | d1 | Vldlr | per |
| Clu | batch | mp | d1 | Vldlr | per |
| Gas6 | batch | mp | d1 | Axl | per |
| Gdnf | batch | mp | d1 | Gfra1 | per |
| Gdnf | batch | per | d1 | Gfra1 | per |
| Gdnf | batch | mp | d1 | Gfra2 | per |
| Gdnf | batch | per | d1 | Gfra2 | per |
| Gmfb | batch | mp | d1 | Egfr | per |
| Gmfb | batch | mp | d1 | Itpr3 | per |
| Hbegf | batch | mp | d1 | Egfr | per |
| Hbegf | constitutive | ec | d1 | Egfr | per |
| Hbegf | constitutive | per | d1 | Egfr | per |
| Il33 | batch | mp | d1 | Il1rl1 | per |
| Jag2 | batch | mp | d1 | Notch3 | per |
| Jag2 | constitutive | ec | d1 | Notch3 | per |
| Jag2 | batch | mp | d1 | Notch1 | per |
| Jag2 | constitutive | ec | d1 | Notch1 | per |
| Jag2 | batch | mp | d1 | Notch2 | per |
| Jag2 | constitutive | ec | d1 | Notch2 | per |
| Ngf | batch | mp | d1 | Ngfr | per |
| Ngf | constitutive | per | d1 | Ngfr | per |
| Ngf | batch | mp | d1 | Sort1 | per |
| Ngf | constitutive | per | d1 | Sort1 | per |
| Nppc | batch | mp | d1 | Npr2 | per |
| Pdap1 | batch | mp | d1 | Pdgfa | per |
| S100b | batch | mp | d1 | Fgfr1 | per |
| S100b | constitutive | per | d1 | Fgfr1 | per |
| Serpini1 | batch | mp | d1 | Plat | per |
| Serpini1 | constitutive | per | d1 | Plat | per |
| Sfrp4 | batch | mp | d1 | Fzd2 | per |
| Sfrp5 | batch | mp | d1 | Fzd2 | per |
| Spp1 | batch | mp | d1 | Itgb5 | per |
| Spp1 | batch | per | d1 | Itgb5 | per |
| Spp1 | batch | mp | d1 | Vtn | per |
| Spp1 | batch | per | d1 | Vtn | per |
| Spp1 | batch | mp | d1 | Itgb3 | per |
| Spp1 | batch | per | d1 | Itgb3 | per |
| Spp1 | batch | mp | d1 | Itgav | per |
| Spp1 | batch | per | d1 | Itgav | per |
| Spp1 | batch | mp | d1 | Itga9 | per |
| Spp1 | batch | per | d1 | Itga9 | per |
| Vegfa | batch | ic | d1 | Ephb2 | per |
| Vegfa | batch | mp | d1 | Ephb2 | per |
| Vegfa | batch | ic | d1 | Vtn | per |
| Vegfa | batch | mp | d1 | Vtn | per |
| Vegfa | batch | ic | d1 | Itga9 | per |
| Vegfa | batch | mp | d1 | Itga9 | per |
| Vegfc | batch | mp | d1 | Itga9 | per |
| Ccl3 | batch | ic | d1 | Ccr5 | per |
| Ccl3 | batch | per | d1 | Ccr5 | per |
| Ccl4 | batch | ic | d1 | Ccr5 | per |
| Ccl4 | batch | per | d1 | Ccr5 | per |
| Il1a | batch | ic | d1 | Il1rap | per |
| Jag1 | batch | ic | d1 | Notch3 | per |
| Jag1 | constitutive | per | d1 | Notch3 | per |
| Jag1 | batch | ic | d1 | Notch1 | per |
| Jag1 | constitutive | per | d1 | Notch1 | per |
| Jag1 | batch | ic | d1 | Notch2 | per |
| Jag1 | constitutive | per | d1 | Notch2 | per |
| Osm | batch | ic | d1 | Osmr | per |
| Osm | batch | per | d1 | Osmr | per |
| Osm | batch | ic | d1 | Il6st | per |
| Osm | batch | per | d1 | Il6st | per |
| Ppbp | batch | ic | d1 | Itgb5 | per |
| Btc | batch | per | d1 | Egfr | per |
| Il11 | batch | per | d1 | Il6st | per |
| Nov | batch | per | d1 | Notch1 | per |
| Nov | batch | per | d1 | Itgb3 | per |
| Nov | batch | per | d1 | Itgav | per |
| Sfrp2 | batch | per | d1 | Fzd2 | per |
| Fstl1 | constitutive | fap | d1 | Cd14 | per |
| Fgf7 | constitutive | fap | d1 | Fgfr2 | per |
| Tgfb2 | constitutive | ec | d1 | Tgfbr3 | per |
| Tgfb2 | constitutive | fap | d1 | Tgfbr3 | per |
| Tgfb2 | constitutive | per | d1 | Tgfbr3 | per |
| Tgfb2 | constitutive | ec | d1 | Vtn | per |
| Tgfb2 | constitutive | fap | d1 | Vtn | per |
| Tgfb2 | constitutive | per | d1 | Vtn | per |
| Bmp5 | constitutive | ec | d1 | Bmpr1a | per |
| Bmp5 | constitutive | fap | d1 | Bmpr1a | per |
| Bmp5 | constitutive | per | d1 | Bmpr1a | per |
| Pdgfb | constitutive | ec | d1 | Pdgfrb | per |
| Cmtm8 | constitutive | ec | d1 | Egfr | per |
| Edn1 | constitutive | ec | d1 | Ednrb | per |
| Edn1 | constitutive | ec | d1 | Ednra | per |
| Nampt | constitutive | ic | d1 | Adora2a | per |
| Fgf1 | constitutive | per | d1 | Fgfr2 | per |
| Fgf1 | constitutive | per | d1 | Fgfr1 | per |
| Ptn | constitutive | per | d1 | Ptprs | per |
| Sfrp1 | constitutive | per | d1 | Fzd2 | per |
| Ntf3 | constitutive | per | d1 | Ngfr | per |
| Ntf3 | constitutive | per | d1 | Ntrk2 | per |
| Adm | batch | fap | d2 | Calcrl | per |
| Adm | batch | per | d2 | Calcrl | per |
| Bmp2 | batch | ec | d2 | Bmpr1a | per |
| Bmp2 | batch | fap | d2 | Bmpr1a | per |
| Bmp2 | batch | mp | d2 | Bmpr1a | per |
| Bmp2 | constitutive | per | d2 | Bmpr1a | per |
| Bmp6 | batch | fap | d2 | Bmpr1a | per |
| Bmp6 | constitutive | per | d2 | Bmpr1a | per |
| Ccl2 | batch | ec | d2 | Ccr2 | per |
| Ccl2 | batch | fap | d2 | Ccr2 | per |
| Ccl2 | batch | ic | d2 | Ccr2 | per |
| Ccl2 | batch | mp | d2 | Ccr2 | per |
| Ccl2 | batch | per | d2 | Ccr2 | per |
| Ccl2 | batch | ec | d2 | Ccr5 | per |
| Ccl2 | batch | fap | d2 | Ccr5 | per |
| Ccl2 | batch | ic | d2 | Ccr5 | per |
| Ccl2 | batch | mp | d2 | Ccr5 | per |
| Ccl2 | batch | per | d2 | Ccr5 | per |
| Ccl3 | batch | ec | d2 | Ccr5 | per |
| Ccl3 | batch | fap | d2 | Ccr5 | per |
| Ccl3 | batch | ic | d2 | Ccr5 | per |
| Ccl3 | batch | mp | d2 | Ccr5 | per |
| Ccl3 | batch | per | d2 | Ccr5 | per |
| Ccl4 | batch | fap | d2 | Ccr5 | per |
| Ccl4 | batch | ic | d2 | Ccr5 | per |
| Ccl4 | batch | mp | d2 | Ccr5 | per |
| Ccl4 | batch | per | d2 | Ccr5 | per |
| Ccl5 | batch | ec | d2 | Ccr5 | per |
| Ccl5 | batch | fap | d2 | Ccr5 | per |
| Ccl7 | batch | ec | d2 | Ccr5 | per |
| Ccl7 | batch | fap | d2 | Ccr5 | per |
| Ccl7 | batch | ic | d2 | Ccr5 | per |
| Ccl7 | batch | mp | d2 | Ccr5 | per |
| Ccl7 | batch | per | d2 | Ccr5 | per |
| Efnb1 | batch | fap | d2 | Ephb2 | per |
| Efnb1 | constitutive | ec | d2 | Ephb2 | per |
| Efnb1 | constitutive | per | d2 | Ephb2 | per |
| Hgf | batch | fap | d2 | Vtn | per |
| Hgf | batch | ic | d2 | Vtn | per |
| Il1b | batch | ec | d2 | Il1rap | per |
| Il1b | batch | fap | d2 | Il1rap | per |
| Il1b | batch | ic | d2 | Il1rap | per |
| Il1b | batch | mp | d2 | Il1rap | per |
| Il1b | batch | per | d2 | Il1rap | per |
| Il1b | batch | ec | d2 | Adrb2 | per |
| Il1b | batch | fap | d2 | Adrb2 | per |
| Il1b | batch | ic | d2 | Adrb2 | per |
| Il1b | batch | mp | d2 | Adrb2 | per |
| Il1b | batch | per | d2 | Adrb2 | per |
| Il33 | batch | fap | d2 | Il1rl1 | per |
| Il33 | batch | mp | d2 | Il1rl1 | per |
| Il6 | batch | ec | d2 | Il6st | per |
| Il6 | batch | fap | d2 | Il6st | per |
| Il6 | batch | per | d2 | Il6st | per |
| Kitl | batch | fap | d2 | Kit | per |
| Kitl | batch | mp | d2 | Kit | per |
| Kitl | constitutive | ec | d2 | Kit | per |
| Kitl | constitutive | per | d2 | Kit | per |
| Lif | batch | fap | d2 | Il6st | per |
| Lif | batch | ic | d2 | Il6st | per |
| Lif | batch | per | d2 | Il6st | per |
| Lif | constitutive | mp | d2 | Il6st | per |
| Pf4 | batch | ec | d2 | Ldlr | per |
| Pf4 | batch | fap | d2 | Ldlr | per |
| Pf4 | batch | ic | d2 | Ldlr | per |
| Pf4 | batch | mp | d2 | Ldlr | per |
| Pf4 | batch | per | d2 | Ldlr | per |
| Sfrp2 | batch | ec | d2 | Fzd2 | per |
| Sfrp2 | batch | fap | d2 | Fzd2 | per |
| Sfrp2 | batch | per | d2 | Fzd2 | per |
| Spp1 | batch | ec | d2 | Itgb5 | per |
| Spp1 | batch | fap | d2 | Itgb5 | per |
| Spp1 | batch | ic | d2 | Itgb5 | per |
| Spp1 | batch | mp | d2 | Itgb5 | per |
| Spp1 | batch | per | d2 | Itgb5 | per |
| Spp1 | batch | ec | d2 | Vtn | per |
| Spp1 | batch | fap | d2 | Vtn | per |
| Spp1 | batch | ic | d2 | Vtn | per |
| Spp1 | batch | mp | d2 | Vtn | per |
| Spp1 | batch | per | d2 | Vtn | per |
| Spp1 | batch | ec | d2 | Itgb3 | per |
| Spp1 | batch | fap | d2 | Itgb3 | per |
| Spp1 | batch | ic | d2 | Itgb3 | per |
| Spp1 | batch | mp | d2 | Itgb3 | per |
| Spp1 | batch | per | d2 | Itgb3 | per |
| Spp1 | batch | ec | d2 | Itgav | per |
| Spp1 | batch | fap | d2 | Itgav | per |
| Spp1 | batch | ic | d2 | Itgav | per |
| Spp1 | batch | mp | d2 | Itgav | per |
| Spp1 | batch | per | d2 | Itgav | per |
| Spp1 | batch | ec | d2 | Itga9 | per |
| Spp1 | batch | fap | d2 | Itga9 | per |
| Spp1 | batch | ic | d2 | Itga9 | per |
| Spp1 | batch | mp | d2 | Itga9 | per |
| Spp1 | batch | per | d2 | Itga9 | per |
| Tnfrsf11b | batch | fap | d2 | Vtn | per |
| Wnt2 | batch | fap | d2 | Sfrp1 | per |
| Wnt2 | batch | fap | d2 | Fzd1 | per |
| Igf1 | batch | ec | d2 | Igfbp2 | per |
| Igf1 | batch | ic | d2 | Igfbp2 | per |
| Igf1 | batch | mp | d2 | Igfbp2 | per |
| Igf1 | batch | ec | d2 | Igfbp7 | per |
| Igf1 | batch | ic | d2 | Igfbp7 | per |
| Igf1 | batch | mp | d2 | Igfbp7 | per |
| Il15 | batch | ec | d2 | Il17ra | per |
| Il15 | batch | ic | d2 | Il17ra | per |
| Il15 | batch | ec | d2 | Il2rg | per |
| Il15 | batch | ic | d2 | Il2rg | per |
| Il15 | batch | ec | d2 | Il15ra | per |
| Il15 | batch | ic | d2 | Il15ra | per |
| Lgals3 | batch | ec | d2 | Lgals3bp | per |
| Lgals3 | batch | ic | d2 | Lgals3bp | per |
| Lgals3 | batch | mp | d2 | Lgals3bp | per |
| Lgals3 | batch | per | d2 | Lgals3bp | per |
| Mmp12 | batch | ec | d2 | Plaur | per |
| Mmp12 | batch | ic | d2 | Plaur | per |
| Mmp12 | batch | mp | d2 | Plaur | per |
| Mmp12 | batch | per | d2 | Plaur | per |
| Osm | batch | ec | d2 | Osmr | per |
| Osm | batch | ic | d2 | Osmr | per |
| Osm | batch | mp | d2 | Osmr | per |
| Osm | batch | per | d2 | Osmr | per |
| Osm | batch | ec | d2 | Il6st | per |
| Osm | batch | ic | d2 | Il6st | per |
| Osm | batch | mp | d2 | Il6st | per |
| Osm | batch | per | d2 | Il6st | per |
| Sfrp1 | batch | ec | d2 | Fzd2 | per |
| Sfrp1 | constitutive | per | d2 | Fzd2 | per |
| Areg | batch | ic | d2 | Egfr | per |
| Areg | batch | mp | d2 | Egfr | per |
| Bmp4 | batch | mp | d2 | Bmpr1a | per |
| Bmp4 | constitutive | ec | d2 | Bmpr1a | per |
| Bmp4 | constitutive | per | d2 | Bmpr1a | per |
| C4b | batch | mp | d2 | C3ar1 | per |
| C4b | constitutive | per | d2 | C3ar1 | per |
| Ccl11 | batch | mp | d2 | Ccr5 | per |
| Ccl11 | batch | per | d2 | Ccr5 | per |
| Ccl8 | batch | mp | d2 | Ccr2 | per |
| Ccl8 | total | fap | d2 | Ccr2 | per |
| Ccl8 | total | ic | d2 | Ccr2 | per |
| Ccl8 | batch | mp | d2 | Ccr5 | per |
| Ccl8 | total | fap | d2 | Ccr5 | per |
| Ccl8 | total | ic | d2 | Ccr5 | per |
| Clcf1 | batch | mp | d2 | Crlf1 | per |
| Clu | batch | mp | d2 | Vldlr | per |
| Clu | total | per | d2 | Vldlr | per |
| Gas6 | batch | mp | d2 | Axl | per |
| Gdnf | batch | mp | d2 | Gfra1 | per |
| Gdnf | batch | per | d2 | Gfra1 | per |
| Gdnf | batch | mp | d2 | Gfra2 | per |
| Gdnf | batch | per | d2 | Gfra2 | per |
| Gmfb | batch | mp | d2 | Egfr | per |
| Gmfb | batch | mp | d2 | Itpr3 | per |
| Hbegf | batch | mp | d2 | Egfr | per |
| Hbegf | constitutive | ec | d2 | Egfr | per |
| Hbegf | constitutive | per | d2 | Egfr | per |
| Jag2 | batch | mp | d2 | Notch3 | per |
| Jag2 | constitutive | ec | d2 | Notch3 | per |
| Jag2 | batch | mp | d2 | Notch1 | per |
| Jag2 | constitutive | ec | d2 | Notch1 | per |
| Jag2 | batch | mp | d2 | Notch2 | per |
| Jag2 | constitutive | ec | d2 | Notch2 | per |
| Ngf | batch | mp | d2 | Ngfr | per |
| Ngf | constitutive | per | d2 | Ngfr | per |
| Ngf | batch | mp | d2 | Sort1 | per |
| Ngf | constitutive | per | d2 | Sort1 | per |
| Nppc | batch | mp | d2 | Npr2 | per |
| Pdap1 | batch | mp | d2 | Pdgfa | per |
| S100b | batch | mp | d2 | Fgfr1 | per |
| S100b | constitutive | per | d2 | Fgfr1 | per |
| Serpini1 | batch | mp | d2 | Plat | per |
| Serpini1 | constitutive | per | d2 | Plat | per |
| Sfrp4 | batch | mp | d2 | Fzd2 | per |
| Sfrp5 | batch | mp | d2 | Fzd2 | per |
| Tgfb1 | batch | mp | d2 | Tgfbr3 | per |
| Tgfb1 | total | ec | d2 | Tgfbr3 | per |
| Tgfb1 | total | ic | d2 | Tgfbr3 | per |
| Tgfb1 | batch | mp | d2 | Vtn | per |
| Tgfb1 | total | ec | d2 | Vtn | per |
| Tgfb1 | total | ic | d2 | Vtn | per |
| Tgfb1 | batch | mp | d2 | Itgav | per |
| Tgfb1 | total | ec | d2 | Itgav | per |
| Tgfb1 | total | ic | d2 | Itgav | per |
| Vegfa | batch | ic | d2 | Ephb2 | per |
| Vegfa | batch | mp | d2 | Ephb2 | per |
| Vegfa | batch | ic | d2 | Vtn | per |
| Vegfa | batch | mp | d2 | Vtn | per |
| Vegfa | batch | ic | d2 | Itga9 | per |
| Vegfa | batch | mp | d2 | Itga9 | per |
| Vegfc | batch | mp | d2 | Itga9 | per |
| Il1a | batch | ic | d2 | Il1rap | per |
| Jag1 | batch | ic | d2 | Notch3 | per |
| Jag1 | constitutive | per | d2 | Notch3 | per |
| Jag1 | batch | ic | d2 | Notch1 | per |
| Jag1 | constitutive | per | d2 | Notch1 | per |
| Jag1 | batch | ic | d2 | Notch2 | per |
| Jag1 | constitutive | per | d2 | Notch2 | per |
| Npy | batch | ic | d2 | Npy1r | per |
| Ppbp | batch | ic | d2 | Itgb5 | per |
| Tnfsf13 | batch | ic | d2 | Tnfrsf11b | per |
| Btc | batch | per | d2 | Egfr | per |
| Il11 | batch | per | d2 | Il6st | per |
| Nov | batch | per | d2 | Notch1 | per |
| Nov | batch | per | d2 | Itgb3 | per |
| Nov | batch | per | d2 | Itgav | per |
| Tnfsf12 | total | ec | d2 | Tnfrsf11b | per |
| Tnfsf12 | total | ic | d2 | Tnfrsf11b | per |
| Tnfsf12 | total | per | d2 | Tnfrsf11b | per |
| Fstl1 | constitutive | fap | d2 | Cd14 | per |
| Fgf7 | constitutive | fap | d2 | Fgfr2 | per |
| Tgfb2 | constitutive | ec | d2 | Tgfbr3 | per |
| Tgfb2 | constitutive | fap | d2 | Tgfbr3 | per |
| Tgfb2 | constitutive | per | d2 | Tgfbr3 | per |
| Tgfb2 | constitutive | ec | d2 | Vtn | per |
| Tgfb2 | constitutive | fap | d2 | Vtn | per |
| Tgfb2 | constitutive | per | d2 | Vtn | per |
| Bmp5 | constitutive | ec | d2 | Bmpr1a | per |
| Bmp5 | constitutive | fap | d2 | Bmpr1a | per |
| Bmp5 | constitutive | per | d2 | Bmpr1a | per |
| Pdgfb | constitutive | ec | d2 | Lrp1 | per |
| Pdgfb | constitutive | ec | d2 | Pdgfrb | per |
| Cmtm8 | constitutive | ec | d2 | Egfr | per |
| Edn1 | constitutive | ec | d2 | Ednrb | per |
| Edn1 | constitutive | ec | d2 | Ednra | per |
| Nampt | constitutive | ic | d2 | Adora2a | per |
| Fgf1 | constitutive | per | d2 | Fgfr2 | per |
| Fgf1 | constitutive | per | d2 | Fgfr1 | per |
| Ptn | constitutive | per | d2 | Ptprs | per |
| Ntf3 | constitutive | per | d2 | Ngfr | per |
| Ntf3 | constitutive | per | d2 | Ntrk2 | per |
| Bmp2 | batch | fap | d3 | Bmpr1a | per |
| Bmp2 | constitutive | per | d3 | Bmpr1a | per |
| Bmp6 | batch | fap | d3 | Bmpr1a | per |
| Bmp6 | constitutive | per | d3 | Bmpr1a | per |
| C4b | batch | fap | d3 | C3ar1 | per |
| C4b | constitutive | per | d3 | C3ar1 | per |
| Ccl3 | batch | ec | d3 | Ccr5 | per |
| Ccl3 | batch | fap | d3 | Ccr5 | per |
| Ccl3 | batch | mp | d3 | Ccr5 | per |
| Ccl3 | batch | per | d3 | Ccr5 | per |
| Ccl4 | batch | fap | d3 | Ccr5 | per |
| Ccl4 | batch | mp | d3 | Ccr5 | per |
| Ccl4 | batch | per | d3 | Ccr5 | per |
| Ccl5 | batch | ec | d3 | Ccr5 | per |
| Ccl5 | batch | fap | d3 | Ccr5 | per |
| Csf2 | batch | fap | d3 | Csf2ra | per |
| Cx3cl1 | batch | fap | d3 | Cx3cr1 | per |
| Cx3cl1 | total | ec | d3 | Cx3cr1 | per |
| Cx3cl1 | total | per | d3 | Cx3cr1 | per |
| Cxcl12 | batch | fap | d3 | Cxcr4 | per |
| Cxcl12 | constitutive | ec | d3 | Cxcr4 | per |
| Cxcl12 | constitutive | per | d3 | Cxcr4 | per |
| Efnb1 | batch | fap | d3 | Ephb2 | per |
| Efnb1 | constitutive | ec | d3 | Ephb2 | per |
| Efnb1 | constitutive | per | d3 | Ephb2 | per |
| Fgf1 | batch | fap | d3 | Fgfr1 | per |
| Fgf1 | constitutive | per | d3 | Fgfr1 | per |
| Gas6 | batch | fap | d3 | Axl | per |
| Hbegf | batch | fap | d3 | Egfr | per |
| Hbegf | constitutive | ec | d3 | Egfr | per |
| Hbegf | constitutive | per | d3 | Egfr | per |
| Hgf | batch | fap | d3 | Vtn | per |
| Hgf | batch | ic | d3 | Vtn | per |
| Igf1 | batch | ec | d3 | Igfbp2 | per |
| Igf1 | batch | fap | d3 | Igfbp2 | per |
| Igf1 | batch | ic | d3 | Igfbp2 | per |
| Igf1 | batch | mp | d3 | Igfbp2 | per |
| Igf1 | batch | ec | d3 | Igfbp5 | per |
| Igf1 | batch | fap | d3 | Igfbp5 | per |
| Igf1 | batch | ic | d3 | Igfbp5 | per |
| Igf1 | batch | mp | d3 | Igfbp5 | per |
| Igf1 | batch | ec | d3 | Igfbp4 | per |
| Igf1 | batch | fap | d3 | Igfbp4 | per |
| Igf1 | batch | ic | d3 | Igfbp4 | per |
| Igf1 | batch | mp | d3 | Igfbp4 | per |
| Igf1 | batch | ec | d3 | Igfbp7 | per |
| Igf1 | batch | fap | d3 | Igfbp7 | per |
| Igf1 | batch | ic | d3 | Igfbp7 | per |
| Igf1 | batch | mp | d3 | Igfbp7 | per |
| Igf2 | batch | fap | d3 | Vtn | per |
| Il1b | batch | fap | d3 | Il1rap | per |
| Il1b | batch | mp | d3 | Il1rap | per |
| Il1b | batch | per | d3 | Il1rap | per |
| Il1b | batch | fap | d3 | Adrb2 | per |
| Il1b | batch | mp | d3 | Adrb2 | per |
| Il1b | batch | per | d3 | Adrb2 | per |
| Il33 | batch | fap | d3 | Il1rl1 | per |
| Il33 | batch | mp | d3 | Il1rl1 | per |
| Kitl | batch | fap | d3 | Kit | per |
| Kitl | constitutive | ec | d3 | Kit | per |
| Kitl | constitutive | per | d3 | Kit | per |
| Mif | batch | fap | d3 | Cd74 | per |
| Mif | batch | mp | d3 | Cd74 | per |
| Ntf3 | batch | ec | d3 | Ngfr | per |
| Ntf3 | batch | fap | d3 | Ngfr | per |
| Ntf3 | constitutive | per | d3 | Ngfr | per |
| Ntf3 | batch | ec | d3 | Ntrk2 | per |
| Ntf3 | batch | fap | d3 | Ntrk2 | per |
| Ntf3 | constitutive | per | d3 | Ntrk2 | per |
| Pthlh | batch | fap | d3 | Pth1r | per |
| Ptn | batch | fap | d3 | Ptprs | per |
| Ptn | constitutive | per | d3 | Ptprs | per |
| S100b | batch | fap | d3 | Fgfr1 | per |
| S100b | constitutive | per | d3 | Fgfr1 | per |
| Sfrp1 | batch | ec | d3 | Fzd2 | per |
| Sfrp1 | batch | fap | d3 | Fzd2 | per |
| Sfrp1 | constitutive | per | d3 | Fzd2 | per |
| Sfrp2 | batch | ec | d3 | Fzd2 | per |
| Sfrp2 | batch | fap | d3 | Fzd2 | per |
| Sfrp2 | batch | per | d3 | Fzd2 | per |
| Sfrp4 | batch | fap | d3 | Fzd2 | per |
| Spp1 | batch | ec | d3 | Itgb5 | per |
| Spp1 | batch | fap | d3 | Itgb5 | per |
| Spp1 | batch | ic | d3 | Itgb5 | per |
| Spp1 | batch | mp | d3 | Itgb5 | per |
| Spp1 | batch | per | d3 | Itgb5 | per |
| Spp1 | batch | ec | d3 | Vtn | per |
| Spp1 | batch | fap | d3 | Vtn | per |
| Spp1 | batch | ic | d3 | Vtn | per |
| Spp1 | batch | mp | d3 | Vtn | per |
| Spp1 | batch | per | d3 | Vtn | per |
| Spp1 | batch | ec | d3 | Itgb3 | per |
| Spp1 | batch | fap | d3 | Itgb3 | per |
| Spp1 | batch | ic | d3 | Itgb3 | per |
| Spp1 | batch | mp | d3 | Itgb3 | per |
| Spp1 | batch | per | d3 | Itgb3 | per |
| Spp1 | batch | ec | d3 | Itgav | per |
| Spp1 | batch | fap | d3 | Itgav | per |
| Spp1 | batch | ic | d3 | Itgav | per |
| Spp1 | batch | mp | d3 | Itgav | per |
| Spp1 | batch | per | d3 | Itgav | per |
| Spp1 | batch | ec | d3 | Itga9 | per |
| Spp1 | batch | fap | d3 | Itga9 | per |
| Spp1 | batch | ic | d3 | Itga9 | per |
| Spp1 | batch | mp | d3 | Itga9 | per |
| Spp1 | batch | per | d3 | Itga9 | per |
| Tgfb3 | batch | fap | d3 | Itgav | per |
| Tgfb3 | total | ec | d3 | Itgav | per |
| Tgfb3 | total | per | d3 | Itgav | per |
| Tnfrsf11b | batch | fap | d3 | Vtn | per |
| Tslp | batch | fap | d3 | Crlf2 | per |
| Tslp | constitutive | ec | d3 | Crlf2 | per |
| Wnt2 | batch | fap | d3 | Sfrp1 | per |
| Wnt2 | batch | fap | d3 | Fzd1 | per |
| Il15 | batch | ec | d3 | Il17ra | per |
| Il15 | batch | ic | d3 | Il17ra | per |
| Il15 | batch | ec | d3 | Il2rg | per |
| Il15 | batch | ic | d3 | Il2rg | per |
| Il15 | batch | ec | d3 | Il15ra | per |
| Il15 | batch | ic | d3 | Il15ra | per |
| Il6 | batch | ec | d3 | Il6st | per |
| Il6 | batch | per | d3 | Il6st | per |
| Lgals3 | batch | ec | d3 | Lgals3bp | per |
| Lgals3 | batch | ic | d3 | Lgals3bp | per |
| Lgals3 | batch | mp | d3 | Lgals3bp | per |
| Lgals3 | batch | per | d3 | Lgals3bp | per |
| Mmp12 | batch | ec | d3 | Plaur | per |
| Mmp12 | batch | ic | d3 | Plaur | per |
| Mmp12 | batch | mp | d3 | Plaur | per |
| Mmp12 | batch | per | d3 | Plaur | per |
| Osm | batch | ec | d3 | Osmr | per |
| Osm | batch | mp | d3 | Osmr | per |
| Osm | batch | per | d3 | Osmr | per |
| Osm | batch | ec | d3 | Il6st | per |
| Osm | batch | mp | d3 | Il6st | per |
| Osm | batch | per | d3 | Il6st | per |
| Vegfc | batch | ec | d3 | Itga9 | per |
| Ccl2 | batch | mp | d3 | Ccr2 | per |
| Ccl2 | batch | per | d3 | Ccr2 | per |
| Ccl2 | batch | mp | d3 | Ccr5 | per |
| Ccl2 | batch | per | d3 | Ccr5 | per |
| Ccl7 | batch | mp | d3 | Ccr5 | per |
| Ccl7 | batch | per | d3 | Ccr5 | per |
| Ccl8 | batch | mp | d3 | Ccr2 | per |
| Ccl8 | total | fap | d3 | Ccr2 | per |
| Ccl8 | total | ic | d3 | Ccr2 | per |
| Ccl8 | batch | mp | d3 | Ccr5 | per |
| Ccl8 | total | fap | d3 | Ccr5 | per |
| Ccl8 | total | ic | d3 | Ccr5 | per |
| Csf1 | batch | ic | d3 | Csf1r | per |
| Csf1 | batch | mp | d3 | Csf1r | per |
| Pdap1 | batch | mp | d3 | Pdgfa | per |
| Tgfb1 | batch | mp | d3 | Tgfbr3 | per |
| Tgfb1 | total | ec | d3 | Tgfbr3 | per |
| Tgfb1 | total | ic | d3 | Tgfbr3 | per |
| Tgfb1 | batch | mp | d3 | Vtn | per |
| Tgfb1 | total | ec | d3 | Vtn | per |
| Tgfb1 | total | ic | d3 | Vtn | per |
| Tgfb1 | batch | mp | d3 | Itgav | per |
| Tgfb1 | total | ec | d3 | Itgav | per |
| Tgfb1 | total | ic | d3 | Itgav | per |
| Tnf | batch | mp | d3 | Tnfrsf1b | per |
| Tnf | batch | per | d3 | Tnfrsf1b | per |
| Npy | batch | ic | d3 | Npy1r | per |
| Tnfsf13 | batch | ic | d3 | Tnfrsf11b | per |
| Btc | batch | per | d3 | Erbb2 | per |
| Btc | batch | per | d3 | Egfr | per |
| Il11 | batch | per | d3 | Il6st | per |
| Lif | batch | per | d3 | Il6st | per |
| Lif | constitutive | mp | d3 | Il6st | per |
| Dll1 | total | mp | d3 | Notch3 | per |
| Dll1 | total | mp | d3 | Notch1 | per |
| Dll1 | total | mp | d3 | Notch2 | per |
| Fstl1 | constitutive | fap | d3 | Cd14 | per |
| Tgfb2 | constitutive | ec | d3 | Tgfbr3 | per |
| Tgfb2 | constitutive | fap | d3 | Tgfbr3 | per |
| Tgfb2 | constitutive | per | d3 | Tgfbr3 | per |
| Tgfb2 | constitutive | ec | d3 | Vtn | per |
| Tgfb2 | constitutive | fap | d3 | Vtn | per |
| Tgfb2 | constitutive | per | d3 | Vtn | per |
| Bmp5 | constitutive | ec | d3 | Bmpr1a | per |
| Bmp5 | constitutive | fap | d3 | Bmpr1a | per |
| Bmp5 | constitutive | per | d3 | Bmpr1a | per |
| Jag2 | constitutive | ec | d3 | Notch3 | per |
| Jag2 | constitutive | ec | d3 | Notch1 | per |
| Jag2 | constitutive | ec | d3 | Notch2 | per |
| Pdgfb | constitutive | ec | d3 | Pdgfrb | per |
| Cmtm8 | constitutive | ec | d3 | Egfr | per |
| Edn1 | constitutive | ec | d3 | Ednrb | per |
| Edn1 | constitutive | ec | d3 | Ednra | per |
| Bmp4 | constitutive | ec | d3 | Bmpr1a | per |
| Bmp4 | constitutive | per | d3 | Bmpr1a | per |
| Nampt | constitutive | ic | d3 | Adora2a | per |
| Serpini1 | constitutive | per | d3 | Plat | per |
| Ngf | constitutive | per | d3 | Ngfr | per |
| Ngf | constitutive | per | d3 | Sort1 | per |
| Jag1 | constitutive | per | d3 | Notch3 | per |
| Jag1 | constitutive | per | d3 | Notch1 | per |
| Jag1 | constitutive | per | d3 | Notch2 | per |
| Apln | batch | ec | d5 | Aplnr | per |
| Apln | batch | fap | d5 | Aplnr | per |
| Efnb1 | batch | fap | d5 | Ephb2 | per |
| Efnb1 | batch | ic | d5 | Ephb2 | per |
| Efnb1 | batch | mp | d5 | Ephb2 | per |
| Efnb1 | constitutive | ec | d5 | Ephb2 | per |
| Efnb1 | constitutive | per | d5 | Ephb2 | per |
| Fgf1 | batch | fap | d5 | Fgfr2 | per |
| Fgf1 | constitutive | per | d5 | Fgfr2 | per |
| Fgf1 | batch | fap | d5 | Fgfr1 | per |
| Fgf1 | constitutive | per | d5 | Fgfr1 | per |
| Gas6 | batch | fap | d5 | Axl | per |
| Hbegf | batch | fap | d5 | Egfr | per |
| Hbegf | constitutive | ec | d5 | Egfr | per |
| Hbegf | constitutive | per | d5 | Egfr | per |
| Hgf | batch | fap | d5 | Vtn | per |
| Hgf | batch | ic | d5 | Vtn | per |
| Igf1 | batch | ec | d5 | Igfbp3 | per |
| Igf1 | batch | fap | d5 | Igfbp3 | per |
| Igf1 | batch | ic | d5 | Igfbp3 | per |
| Igf1 | batch | per | d5 | Igfbp3 | per |
| Igf1 | batch | ec | d5 | Igfbp5 | per |
| Igf1 | batch | fap | d5 | Igfbp5 | per |
| Igf1 | batch | ic | d5 | Igfbp5 | per |
| Igf1 | batch | per | d5 | Igfbp5 | per |
| Igf1 | batch | ec | d5 | Igfbp4 | per |
| Igf1 | batch | fap | d5 | Igfbp4 | per |
| Igf1 | batch | ic | d5 | Igfbp4 | per |
| Igf1 | batch | per | d5 | Igfbp4 | per |
| Igf1 | batch | ec | d5 | Igfbp7 | per |
| Igf1 | batch | fap | d5 | Igfbp7 | per |
| Igf1 | batch | ic | d5 | Igfbp7 | per |
| Igf1 | batch | per | d5 | Igfbp7 | per |
| Igf2 | batch | ec | d5 | Vtn | per |
| Igf2 | batch | fap | d5 | Vtn | per |
| Igf2 | batch | ic | d5 | Vtn | per |
| Igf2 | batch | mp | d5 | Vtn | per |
| Il1b | batch | fap | d5 | Il1rap | per |
| Il1b | batch | fap | d5 | Adrb2 | per |
| Il33 | batch | fap | d5 | Il1rl1 | per |
| Il33 | batch | mp | d5 | Il1rl1 | per |
| Il33 | batch | per | d5 | Il1rl1 | per |
| Kitl | batch | fap | d5 | Kit | per |
| Kitl | batch | ic | d5 | Kit | per |
| Kitl | batch | mp | d5 | Kit | per |
| Kitl | constitutive | ec | d5 | Kit | per |
| Kitl | constitutive | per | d5 | Kit | per |
| Mmp13 | batch | ec | d5 | F2r | per |
| Mmp13 | batch | fap | d5 | F2r | per |
| Mmp13 | batch | ic | d5 | F2r | per |
| Mmp13 | batch | mp | d5 | F2r | per |
| Mmp13 | batch | per | d5 | F2r | per |
| Ntf3 | batch | ec | d5 | Ngfr | per |
| Ntf3 | batch | fap | d5 | Ngfr | per |
| Ntf3 | constitutive | per | d5 | Ngfr | per |
| Ntf3 | batch | ec | d5 | Ntrk2 | per |
| Ntf3 | batch | fap | d5 | Ntrk2 | per |
| Ntf3 | constitutive | per | d5 | Ntrk2 | per |
| Pf4 | batch | fap | d5 | Ldlr | per |
| Pthlh | batch | fap | d5 | Pth1r | per |
| Ptn | batch | fap | d5 | Ptprs | per |
| Ptn | constitutive | per | d5 | Ptprs | per |
| S100b | batch | fap | d5 | Fgfr1 | per |
| S100b | constitutive | per | d5 | Fgfr1 | per |
| Sfrp1 | batch | fap | d5 | Fzd2 | per |
| Sfrp1 | constitutive | per | d5 | Fzd2 | per |
| Sfrp2 | batch | fap | d5 | Fzd2 | per |
| Sfrp2 | batch | per | d5 | Fzd2 | per |
| Sfrp4 | batch | fap | d5 | Fzd2 | per |
| Sfrp4 | batch | per | d5 | Fzd2 | per |
| Spp1 | batch | fap | d5 | Itgb5 | per |
| Spp1 | batch | ic | d5 | Itgb5 | per |
| Spp1 | batch | mp | d5 | Itgb5 | per |
| Spp1 | batch | fap | d5 | Vtn | per |
| Spp1 | batch | ic | d5 | Vtn | per |
| Spp1 | batch | mp | d5 | Vtn | per |
| Spp1 | batch | fap | d5 | Itgb3 | per |
| Spp1 | batch | ic | d5 | Itgb3 | per |
| Spp1 | batch | mp | d5 | Itgb3 | per |
| Spp1 | batch | fap | d5 | Itgav | per |
| Spp1 | batch | ic | d5 | Itgav | per |
| Spp1 | batch | mp | d5 | Itgav | per |
| Spp1 | batch | fap | d5 | Itga9 | per |
| Spp1 | batch | ic | d5 | Itga9 | per |
| Spp1 | batch | mp | d5 | Itga9 | per |
| Tgfb3 | batch | fap | d5 | Itgav | per |
| Tgfb3 | total | ec | d5 | Itgav | per |
| Tgfb3 | total | per | d5 | Itgav | per |
| Tnfrsf11b | batch | fap | d5 | Vtn | per |
| Clu | batch | ec | d5 | Vldlr | per |
| Clu | batch | ic | d5 | Vldlr | per |
| Dll1 | batch | ec | d5 | Notch3 | per |
| Dll1 | total | mp | d5 | Notch3 | per |
| Dll1 | batch | ec | d5 | Notch1 | per |
| Dll1 | total | mp | d5 | Notch1 | per |
| Dll1 | batch | ec | d5 | Notch2 | per |
| Dll1 | total | mp | d5 | Notch2 | per |
| Fgf9 | batch | ec | d5 | Fgfr2 | per |
| Il15 | batch | ec | d5 | Il2rg | per |
| Il15 | batch | ic | d5 | Il2rg | per |
| Il15 | batch | ec | d5 | Il15ra | per |
| Il15 | batch | ic | d5 | Il15ra | per |
| Il6 | batch | ec | d5 | Il6st | per |
| Il6 | batch | per | d5 | Il6st | per |
| Vegfc | batch | ec | d5 | Itga9 | per |
| Vegfc | batch | mp | d5 | Itga9 | per |
| Areg | batch | mp | d5 | Egfr | per |
| Gdnf | batch | mp | d5 | Gfra1 | per |
| Gdnf | batch | per | d5 | Gfra1 | per |
| Gdnf | batch | mp | d5 | Gfra2 | per |
| Gdnf | batch | per | d5 | Gfra2 | per |
| Gmfb | batch | mp | d5 | Egfr | per |
| Gmfb | batch | mp | d5 | Itpr3 | per |
| Pdap1 | batch | mp | d5 | Pdgfa | per |
| Tgfb2 | batch | mp | d5 | Tgfbr3 | per |
| Tgfb2 | constitutive | ec | d5 | Tgfbr3 | per |
| Tgfb2 | constitutive | fap | d5 | Tgfbr3 | per |
| Tgfb2 | constitutive | per | d5 | Tgfbr3 | per |
| Tgfb2 | batch | mp | d5 | Vtn | per |
| Tgfb2 | constitutive | ec | d5 | Vtn | per |
| Tgfb2 | constitutive | fap | d5 | Vtn | per |
| Tgfb2 | constitutive | per | d5 | Vtn | per |
| Lgals3 | batch | ic | d5 | Lgals3bp | per |
| Npy | batch | ic | d5 | Npy1r | per |
| Pdgfb | batch | ic | d5 | Pdgfrb | per |
| Pdgfb | constitutive | ec | d5 | Pdgfrb | per |
| Tnfsf13 | batch | ic | d5 | Tnfrsf11b | per |
| Adm | batch | per | d5 | Calcrl | per |
| Btc | batch | per | d5 | Erbb2 | per |
| Btc | batch | per | d5 | Egfr | per |
| Il11 | batch | per | d5 | Il6st | per |
| Lif | batch | per | d5 | Il6st | per |
| Lif | constitutive | mp | d5 | Il6st | per |
| Nov | batch | per | d5 | Notch1 | per |
| Nov | batch | per | d5 | Itgb3 | per |
| Nov | batch | per | d5 | Itgav | per |
| Sfrp5 | batch | per | d5 | Fzd2 | per |
| Fgf7 | constitutive | fap | d5 | Fgfr2 | per |
| Bmp5 | constitutive | ec | d5 | Bmpr1a | per |
| Bmp5 | constitutive | fap | d5 | Bmpr1a | per |
| Bmp5 | constitutive | per | d5 | Bmpr1a | per |
| Jag2 | constitutive | ec | d5 | Notch3 | per |
| Jag2 | constitutive | ec | d5 | Notch1 | per |
| Jag2 | constitutive | ec | d5 | Notch2 | per |
| Cmtm8 | constitutive | ec | d5 | Egfr | per |
| Edn1 | constitutive | ec | d5 | Ednrb | per |
| Edn1 | constitutive | ec | d5 | Ednra | per |
| Bmp4 | constitutive | ec | d5 | Bmpr1a | per |
| Bmp4 | constitutive | per | d5 | Bmpr1a | per |
| Nampt | constitutive | ic | d5 | Adora2a | per |
| Serpini1 | constitutive | per | d5 | Plat | per |
| Ngf | constitutive | per | d5 | Ngfr | per |
| Ngf | constitutive | per | d5 | Sort1 | per |
| Bmp6 | constitutive | per | d5 | Bmpr1a | per |
| Jag1 | constitutive | per | d5 | Notch3 | per |
| Jag1 | constitutive | per | d5 | Notch1 | per |
| Jag1 | constitutive | per | d5 | Notch2 | per |
| Bmp2 | constitutive | per | d5 | Bmpr1a | per |
| Bmp2 | batch | fap | d10 | Bmpr1a | per |
| Bmp2 | batch | mp | d10 | Bmpr1a | per |
| Bmp2 | constitutive | per | d10 | Bmpr1a | per |
| Bmp4 | batch | fap | d10 | Bmpr1a | per |
| Bmp4 | batch | mp | d10 | Bmpr1a | per |
| Bmp4 | constitutive | ec | d10 | Bmpr1a | per |
| Bmp4 | constitutive | per | d10 | Bmpr1a | per |
| Bmp6 | batch | fap | d10 | Bmpr1a | per |
| Bmp6 | batch | mp | d10 | Bmpr1a | per |
| Bmp6 | constitutive | per | d10 | Bmpr1a | per |
| Bmp7 | batch | fap | d10 | Bmpr1a | per |
| Clu | batch | ec | d10 | Vldlr | per |
| Clu | batch | fap | d10 | Vldlr | per |
| Clu | batch | mp | d10 | Vldlr | per |
| Clu | total | per | d10 | Vldlr | per |
| Crlf1 | batch | fap | d10 | Ctf1 | per |
| Crlf1 | batch | mp | d10 | Ctf1 | per |
| Edn1 | batch | fap | d10 | Ednra | per |
| Edn1 | constitutive | ec | d10 | Ednra | per |
| Efnb1 | batch | fap | d10 | Ephb2 | per |
| Efnb1 | batch | mp | d10 | Ephb2 | per |
| Efnb1 | constitutive | ec | d10 | Ephb2 | per |
| Efnb1 | constitutive | per | d10 | Ephb2 | per |
| Fgf1 | batch | fap | d10 | Fgfr2 | per |
| Fgf1 | batch | mp | d10 | Fgfr2 | per |
| Fgf1 | constitutive | per | d10 | Fgfr2 | per |
| Fgf1 | batch | fap | d10 | Fgfr1 | per |
| Fgf1 | batch | mp | d10 | Fgfr1 | per |
| Fgf1 | constitutive | per | d10 | Fgfr1 | per |
| Gas6 | batch | fap | d10 | Axl | per |
| Gas6 | batch | mp | d10 | Axl | per |
| Hbegf | batch | fap | d10 | Egfr | per |
| Hbegf | batch | mp | d10 | Egfr | per |
| Hbegf | constitutive | ec | d10 | Egfr | per |
| Hbegf | constitutive | per | d10 | Egfr | per |
| Igf1 | batch | ec | d10 | Igfbp7 | per |
| Igf1 | batch | fap | d10 | Igfbp7 | per |
| Igf1 | batch | ic | d10 | Igfbp7 | per |
| Igf2 | batch | ec | d10 | Vtn | per |
| Igf2 | batch | fap | d10 | Vtn | per |
| Igf2 | batch | mp | d10 | Vtn | per |
| Kitl | batch | fap | d10 | Kit | per |
| Kitl | constitutive | ec | d10 | Kit | per |
| Kitl | constitutive | per | d10 | Kit | per |
| Mdk | batch | fap | d10 | Lrp1 | per |
| Mdk | batch | mp | d10 | Lrp1 | per |
| Nov | batch | fap | d10 | Notch1 | per |
| Nov | batch | per | d10 | Notch1 | per |
| Nov | batch | fap | d10 | Itgb3 | per |
| Nov | batch | per | d10 | Itgb3 | per |
| Nov | batch | fap | d10 | Itgav | per |
| Nov | batch | per | d10 | Itgav | per |
| Ntf3 | batch | ec | d10 | Ntrk2 | per |
| Ntf3 | batch | fap | d10 | Ntrk2 | per |
| Ntf3 | batch | mp | d10 | Ntrk2 | per |
| Ntf3 | constitutive | per | d10 | Ntrk2 | per |
| Pthlh | batch | fap | d10 | Pth1r | per |
| Ptn | batch | fap | d10 | Ptprs | per |
| Ptn | batch | mp | d10 | Ptprs | per |
| Ptn | constitutive | per | d10 | Ptprs | per |
| S100b | batch | fap | d10 | Fgfr1 | per |
| S100b | batch | mp | d10 | Fgfr1 | per |
| S100b | constitutive | per | d10 | Fgfr1 | per |
| Sfrp1 | batch | fap | d10 | Fzd2 | per |
| Sfrp1 | constitutive | per | d10 | Fzd2 | per |
| Sfrp4 | batch | fap | d10 | Fzd2 | per |
| Sfrp4 | batch | mp | d10 | Fzd2 | per |
| Tgfb3 | batch | fap | d10 | Itgav | per |
| Tgfb3 | batch | mp | d10 | Itgav | per |
| Vegfa | batch | fap | d10 | Ephb2 | per |
| Vegfa | batch | ic | d10 | Ephb2 | per |
| Vegfa | batch | mp | d10 | Ephb2 | per |
| Vegfa | batch | fap | d10 | Vtn | per |
| Vegfa | batch | ic | d10 | Vtn | per |
| Vegfa | batch | mp | d10 | Vtn | per |
| Vegfa | batch | fap | d10 | Itga9 | per |
| Vegfa | batch | ic | d10 | Itga9 | per |
| Vegfa | batch | mp | d10 | Itga9 | per |
| Wnt2 | batch | fap | d10 | Sfrp1 | per |
| Wnt2 | batch | fap | d10 | Fzd1 | per |
| Dll1 | batch | ec | d10 | Notch3 | per |
| Dll1 | batch | ec | d10 | Notch1 | per |
| Dll1 | batch | ec | d10 | Notch2 | per |
| Fgf9 | batch | ec | d10 | Fgfr2 | per |
| Il15 | batch | ec | d10 | Il15ra | per |
| Il15 | batch | ic | d10 | Il15ra | per |
| Vegfc | batch | ec | d10 | Itga9 | per |
| Clcf1 | batch | ic | d10 | Crlf1 | per |
| Clcf1 | batch | mp | d10 | Crlf1 | per |
| Ctf1 | batch | mp | d10 | Il6st | per |
| Edn3 | batch | mp | d10 | Ednra | per |
| Jag1 | batch | ic | d10 | Notch3 | per |
| Jag1 | batch | mp | d10 | Notch3 | per |
| Jag1 | constitutive | per | d10 | Notch3 | per |
| Jag1 | batch | ic | d10 | Notch1 | per |
| Jag1 | batch | mp | d10 | Notch1 | per |
| Jag1 | constitutive | per | d10 | Notch1 | per |
| Jag1 | batch | ic | d10 | Notch2 | per |
| Jag1 | batch | mp | d10 | Notch2 | per |
| Jag1 | constitutive | per | d10 | Notch2 | per |
| Jag2 | batch | mp | d10 | Notch3 | per |
| Jag2 | constitutive | ec | d10 | Notch3 | per |
| Jag2 | batch | mp | d10 | Notch1 | per |
| Jag2 | constitutive | ec | d10 | Notch1 | per |
| Jag2 | batch | mp | d10 | Notch2 | per |
| Jag2 | constitutive | ec | d10 | Notch2 | per |
| Ngf | batch | mp | d10 | Sort1 | per |
| Ngf | constitutive | per | d10 | Sort1 | per |
| Nppc | batch | mp | d10 | Npr2 | per |
| Serpini1 | batch | mp | d10 | Plat | per |
| Serpini1 | constitutive | per | d10 | Plat | per |
| Sfrp2 | batch | mp | d10 | Fzd2 | per |
| Sfrp5 | batch | mp | d10 | Fzd2 | per |
| Sfrp5 | batch | per | d10 | Fzd2 | per |
| Tgfb2 | batch | mp | d10 | Tgfbr3 | per |
| Tgfb2 | constitutive | ec | d10 | Tgfbr3 | per |
| Tgfb2 | constitutive | fap | d10 | Tgfbr3 | per |
| Tgfb2 | constitutive | per | d10 | Tgfbr3 | per |
| Tgfb2 | batch | mp | d10 | Vtn | per |
| Tgfb2 | constitutive | ec | d10 | Vtn | per |
| Tgfb2 | constitutive | fap | d10 | Vtn | per |
| Tgfb2 | constitutive | per | d10 | Vtn | per |
| Areg | batch | ic | d10 | Egfr | per |
| Il1a | batch | ic | d10 | Il1rap | per |
| Il1b | batch | ic | d10 | Il1rap | per |
| Il1b | batch | ic | d10 | Adrb2 | per |
| Il6 | batch | ic | d10 | Il6st | per |
| Lif | batch | ic | d10 | Il6st | per |
| Lif | constitutive | mp | d10 | Il6st | per |
| Osm | batch | ic | d10 | Osmr | per |
| Osm | batch | ic | d10 | Il6st | per |
| Pdgfb | batch | ic | d10 | Lrp1 | per |
| Pdgfb | constitutive | ec | d10 | Lrp1 | per |
| Pdgfb | batch | ic | d10 | Pdgfrb | per |
| Pdgfb | constitutive | ec | d10 | Pdgfrb | per |
| Pf4 | batch | ic | d10 | Ldlr | per |
| Ppbp | batch | ic | d10 | Itgb5 | per |
| Adm | batch | per | d10 | Calcrl | per |
| Gdnf | batch | per | d10 | Gfra1 | per |
| Gdnf | batch | per | d10 | Gfra2 | per |
| Tgfb1 | total | ec | d10 | Tgfbr3 | per |
| Tgfb1 | total | ic | d10 | Tgfbr3 | per |
| Tgfb1 | total | ec | d10 | Vtn | per |
| Tgfb1 | total | ic | d10 | Vtn | per |
| Tgfb1 | total | ec | d10 | Itgav | per |
| Tgfb1 | total | ic | d10 | Itgav | per |
| Fgf7 | constitutive | fap | d10 | Fgfr2 | per |
| Bmp5 | constitutive | ec | d10 | Bmpr1a | per |
| Bmp5 | constitutive | fap | d10 | Bmpr1a | per |
| Bmp5 | constitutive | per | d10 | Bmpr1a | per |
| Cmtm8 | constitutive | ec | d10 | Egfr | per |
| Nampt | constitutive | ic | d10 | Adora2a | per |
