## Supplementary figures and images for "Spatial compartmentalization of signalling imparts source-specific functions on secreted factors"

### SupFigure1.png

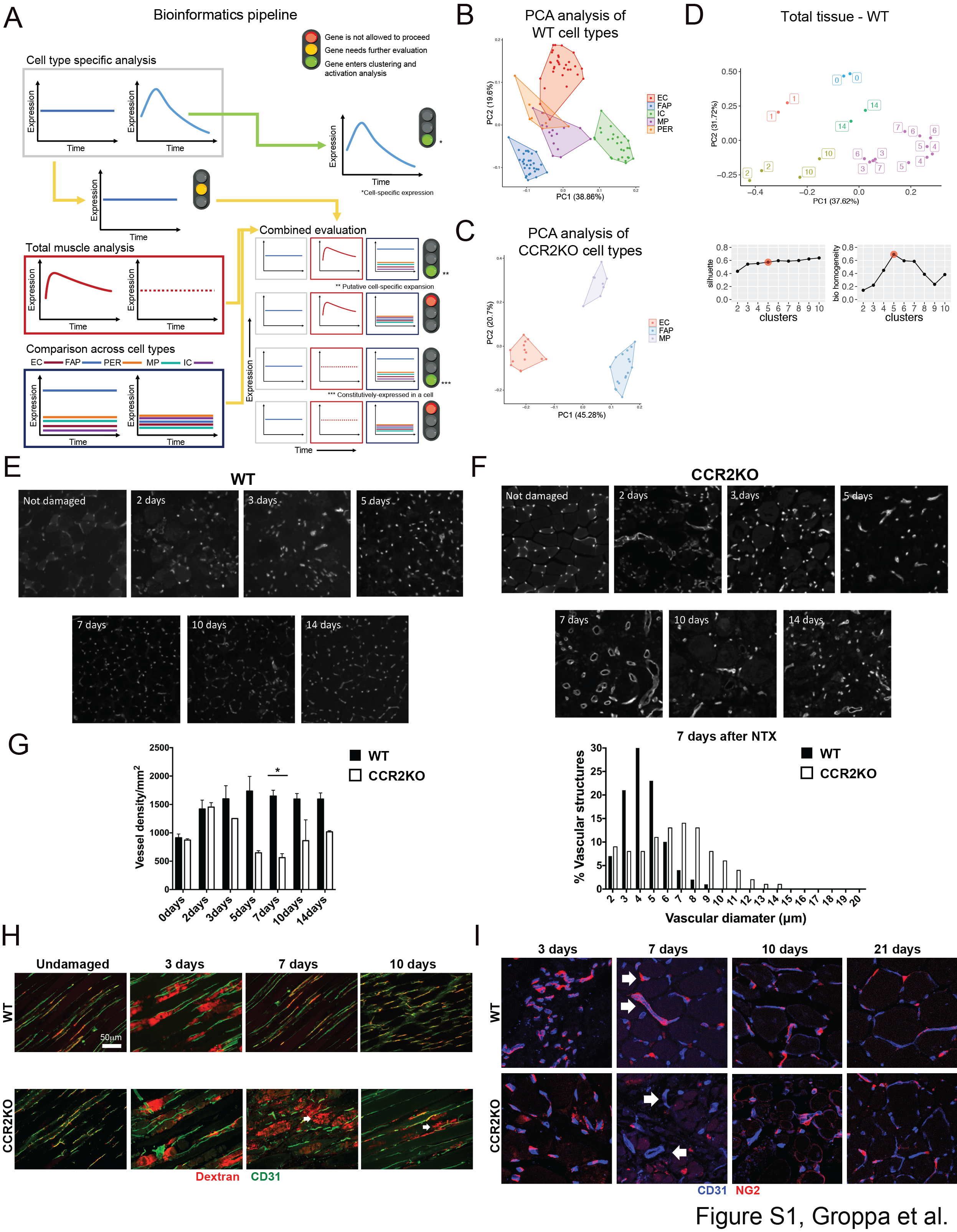

### SupFigure2.png

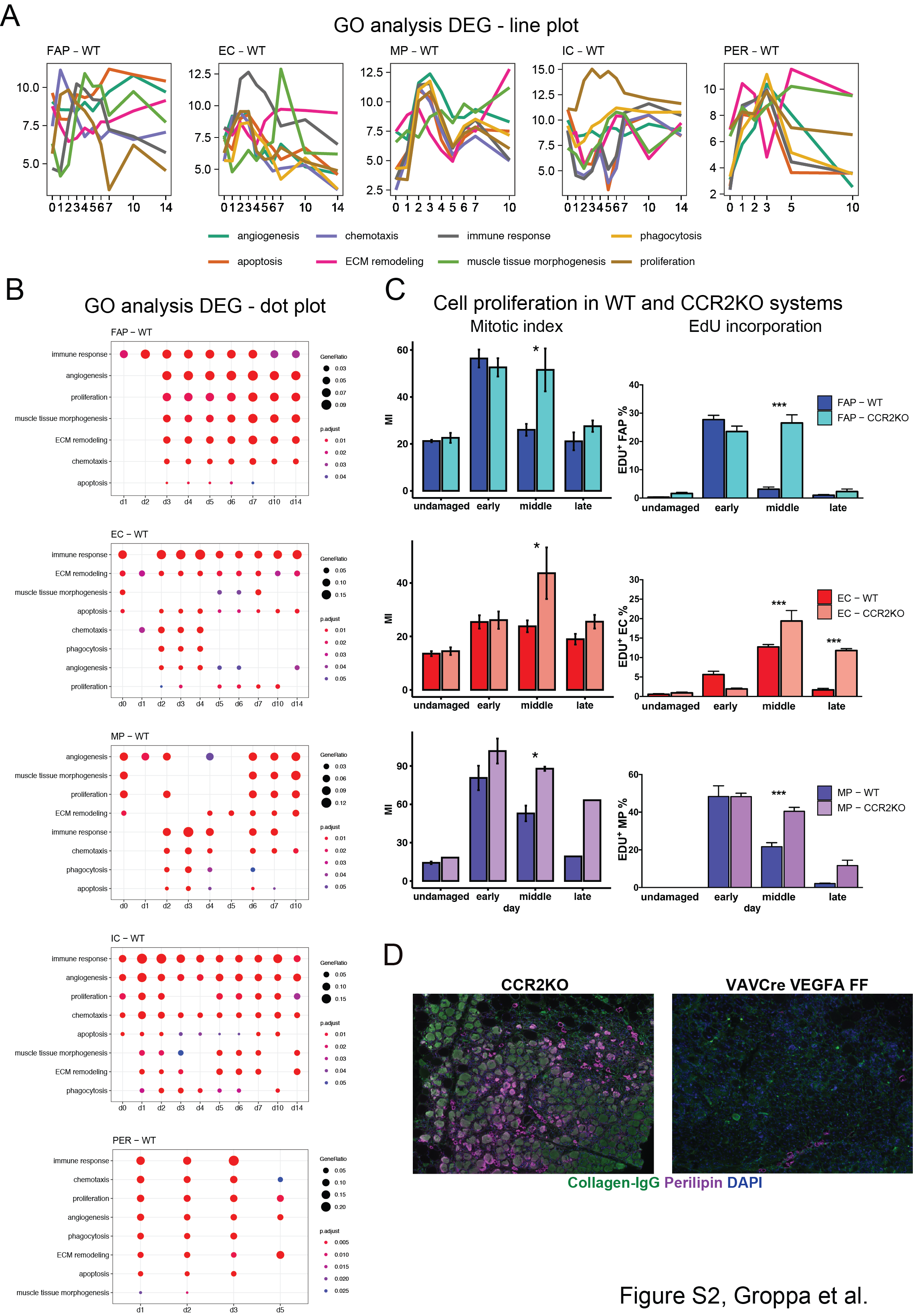

### SupFigure3.png

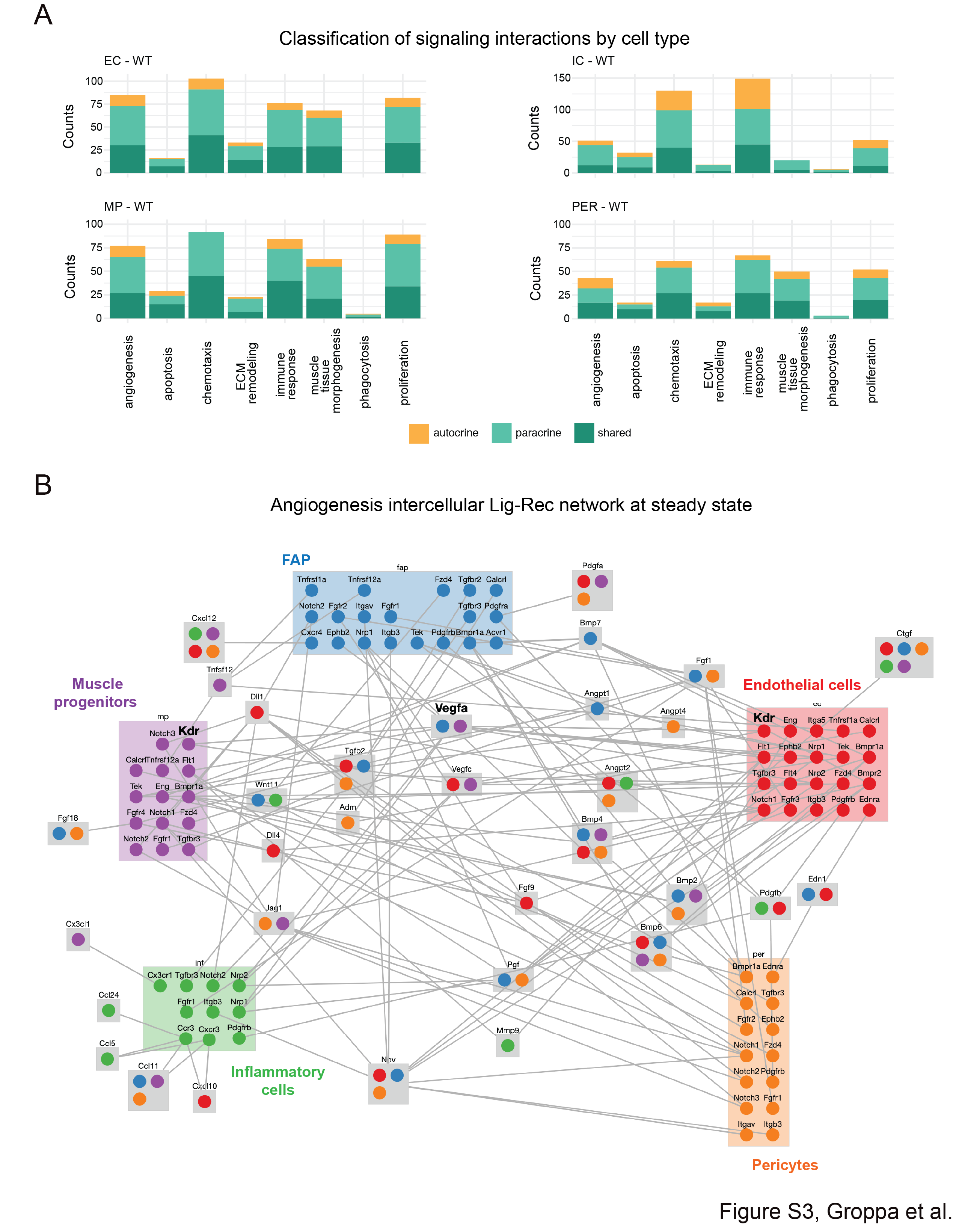

### SupFigure4.png

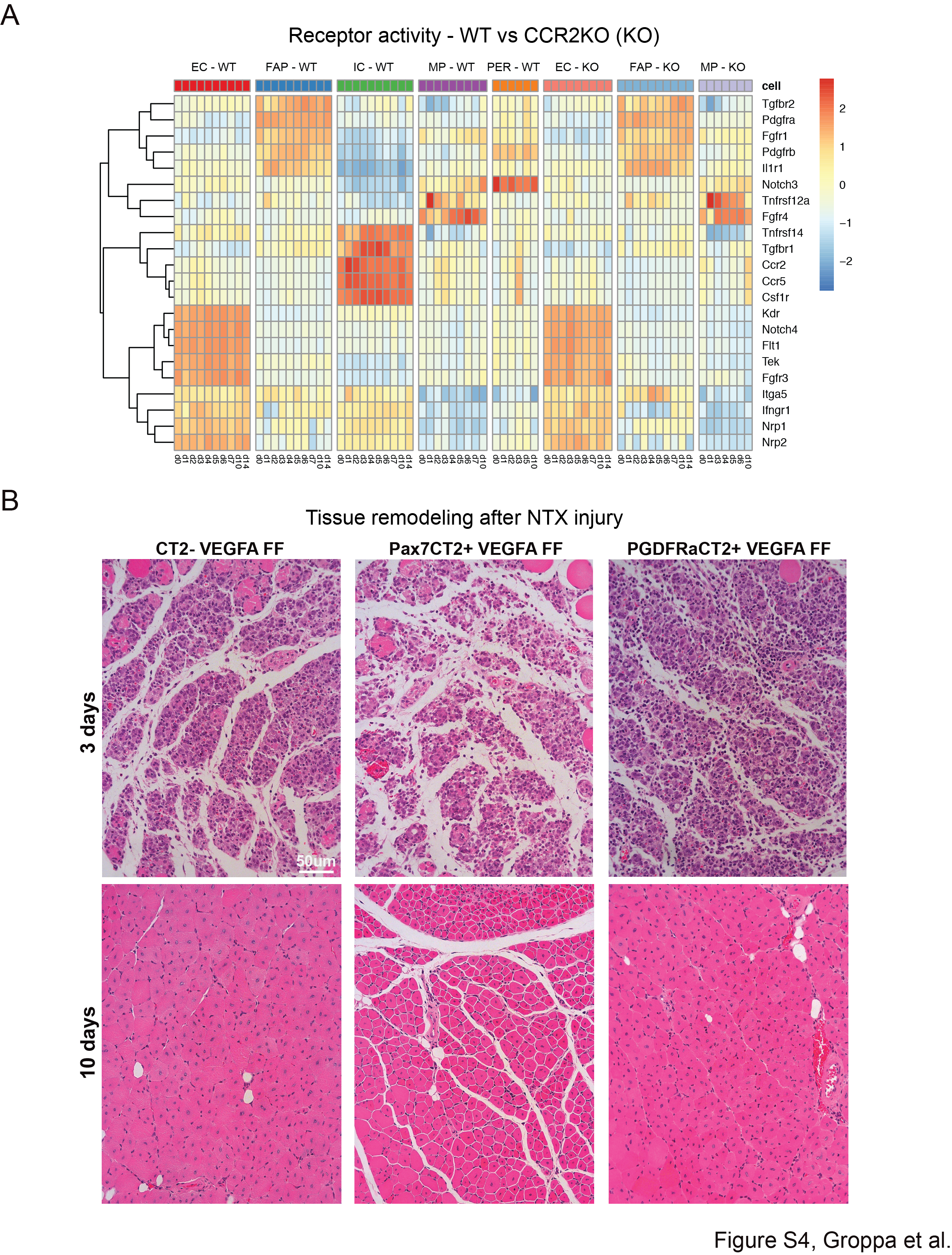

### SupFigure5.png

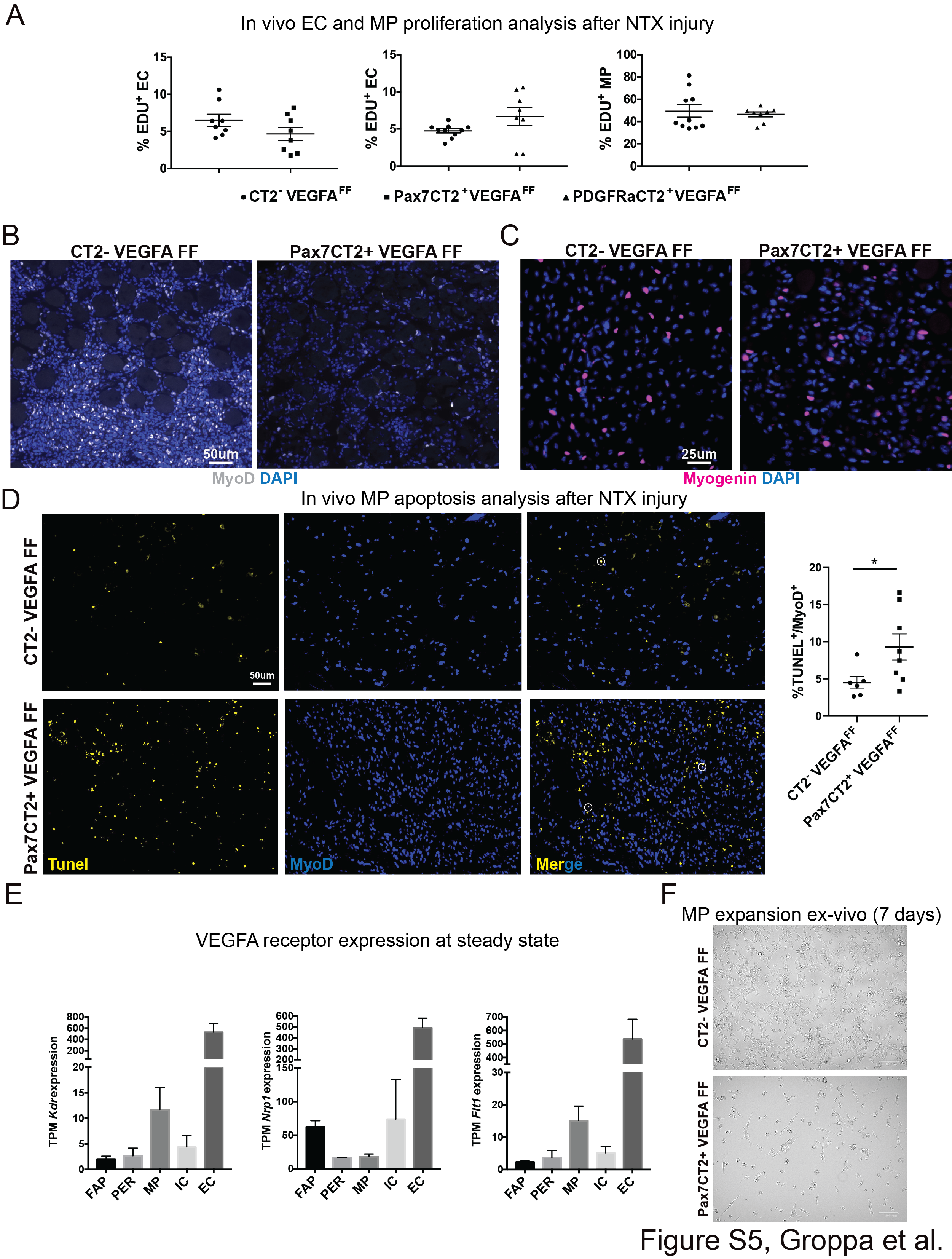

### SupFigure6.png

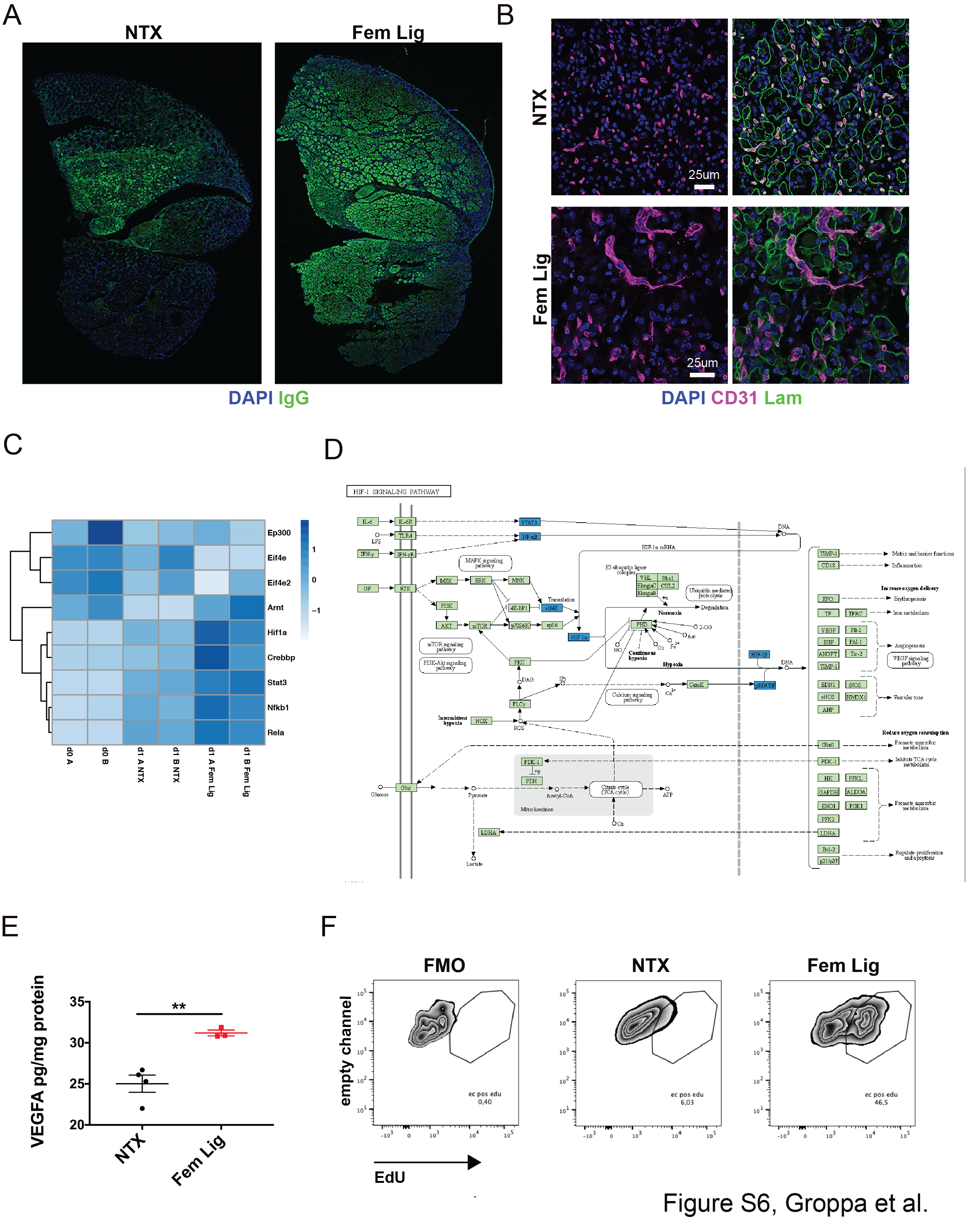

### SupFigure7.png

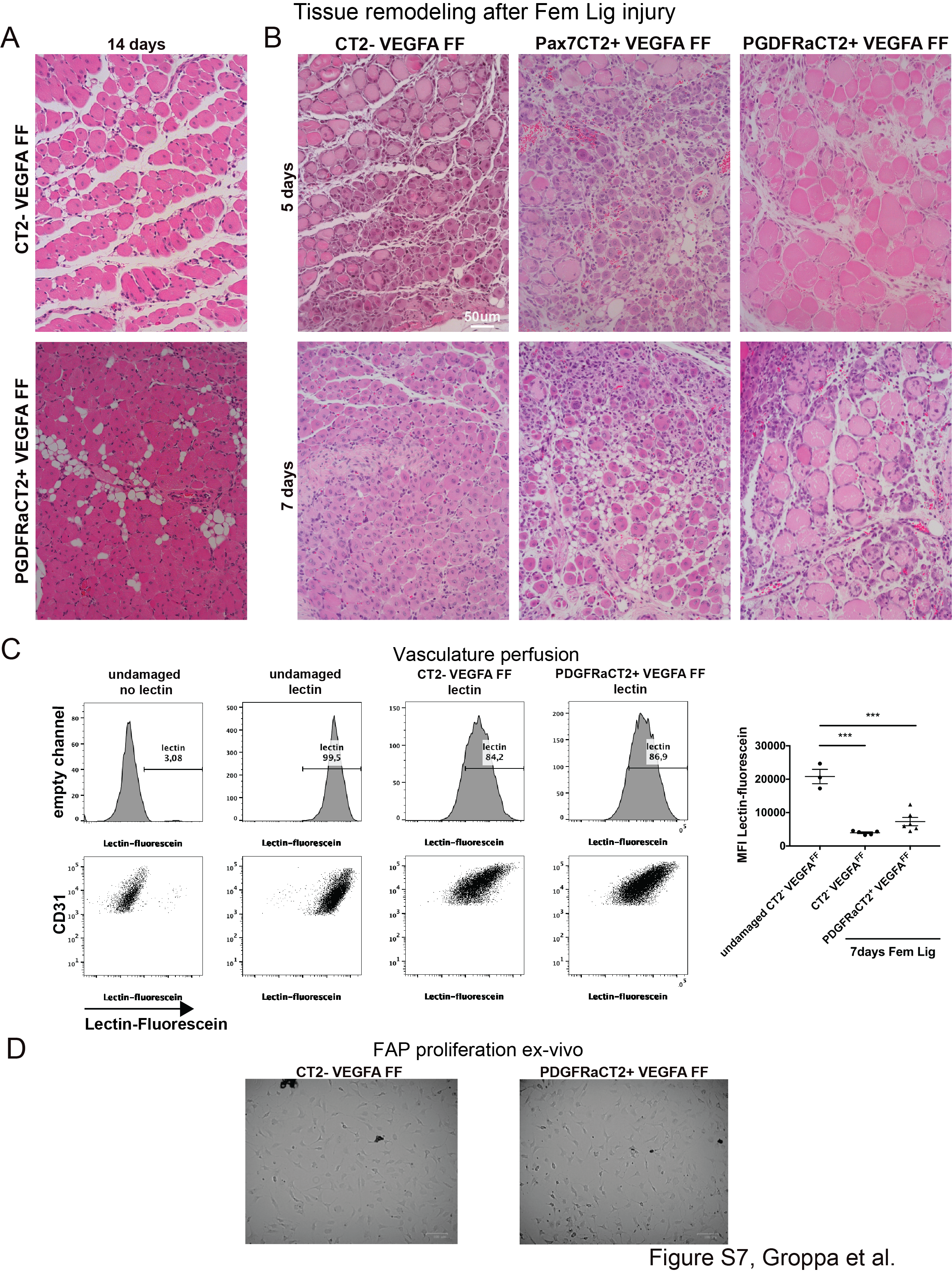
